# Power spectra reveal distinct BOLD resting‐state time courses in white matter

**DOI:** 10.1101/2021.02.24.432346

**Authors:** Muwei Li, Yurui Gao, Zhaohua Ding, John C. Gore

## Abstract

Accurate characterization of the time courses of BOLD signal changes is crucial for the analysis and interpretation of functional MRI data. While several studies have shown that white matter (WM) exhibits distinct BOLD responses evoked by tasks, there have been no comprehensive investigations into the time courses of spontaneous signal fluctuations in WM. We measured the power spectra of the resting‐state time courses in a set of regions within WM identified as showing synchronous signals using independent components analysis. In each component, a clear separation between voxels into two categories was evident, based on their power spectra: one group exhibited a single peak, the other had an additional peak at a higher frequency. Their groupings are location‐specific, and their distributions reflect unique neurovascular and anatomical configurations. Importantly, the two categories of voxels differed in their engagement in functional integration, revealed by differences in the number of inter‐regional connections based on the two categories separately. Moreover, the power spectral measurements in voxels with two peaks in specific components predict specific human behaviors. Taken together, these findings suggest WM signals are heterogeneous in nature and depend on local structural‐vascular‐functional associations.

## Introduction

Functional magnetic resonance imaging (fMRI) has become the leading technique for mapping neural activities by detecting changes in blood‐oxygen‐level‐dependent (BOLD) signals in the brain. Mathematical analyses have been developed to identify regions of activation where BOLD signals rise significantly above the baseline in response to a stimulus ^1^, and to identify functional circuits, by detecting synchronous BOLD signals in segregated gray matter (GM) regions during task‐free, resting states ^2,3^. White matter (WM) makes up over half of the brain volume, and exhibits a comparable oxygen extraction profile to that in GM ^4^, but its engagement is rarely reported in either task or resting‐state fMRI studies. In practice, the average BOLD fluctuations within WM have often been regressed out as a nuisance covariate ^5,6^. In recent years, a growing body of literature has recognized that changes in BOLD signals in WM may reflect neural activities ^7–9^ and, by using appropriate methods, BOLD changes associated with external stimuli can be reliably detected with conventional fMRI ^10–15^. However, the sensitivity of detecting WM activation is often much lower compared to GM, possibly due to incorrect assumptions made about the time courses of responses that are incorporated into regression models for detection ^16^. According to our recent studies, WM tracts that are involved in the processing of external stimuli may exhibit distinct time courses different from GM, with reduced magnitudes and delayed peaks, reflecting quantitative differences in hemodynamic conditions between GM and WM ^17–19^. These findings highlight the importance of characterizing the temporal profiles of BOLD signals so that neurovascular coupling in WM may be better understood and incorporated appropriately into analyses.

To date, efforts to characterize the BOLD response of WM have mostly focused on task‐evoked BOLD signals where the onset of each time course is locked to known events (or stimuli). Then the statistical significances of measurements of, for example, the magnitudes and times to peak of the responses can be rigorously evaluated based on averaging and comparing across trials, runs, sessions, and subjects. However, the lack of reference events affecting intrinsic BOLD signals makes it difficult to quantify or compare time courses among individuals or regions. The existing literature usually reports data based on inter‐regional correlations, which are reproducible over scans or subjects ^20–25^, in which, however, characteristics of the time courses themselves were disregarded. Spectral analysis of signals represents a complementary approach to identifying features of interest, and BOLD effects that appear to be random and small over time may reflect a distinct pattern of component frequencies. In fact, power spectral analysis has been frequently used to characterize spontaneous activities in GM, identifying significant differences in BOLD changes between different cortical areas, different cognitive states, or different frequency bands ^26–28^. Though shown to provide robust statistical descriptions of signals ^29–31^, such analyses have been limited to GM, and there remains a paucity of information regarding resting‐state time courses in WM. Our recent studies suggest that, compared to GM, WM signals have a comparable frequency range and exhibit similar patterns in their powers as a function of frequency ^32^ so power spectra may provide additional insight into the nature of WM signal fluctuations.

Here we report our detailed analysis of the power spectra of WM time courses in a set of regions identified from a data‐driven derivation of distinct functional activities. We observed that, while each region exhibited similar patterns of BOLD fluctuations, closer analyses showed the voxels within each region were readily clustered into two groups based on their distinct power spectra: one group had spectra with single peaks (SP voxels), and the other showed dual peaks (DP voxels), a spectral shape rarely observed in GM voxels. We then explored the possible origins of the second peak in the power spectra by their relationships with hemodynamic and anatomical features in the same locations. We observed that DP voxels exhibit more prominent initial dips than SP voxels in their hemodynamic response functions (HRFs), suggesting that the presence of the higher frequency peaks reflect the local hemodynamic conditions. Based on diffusion imaging data, we observed significantly more crossing fibers in DP voxels than in SP voxels, providing an anatomical basis for the distinct patterns. Next, we compared the inter‐region functional connectivities calculated separately from the correlations between the time courses of entire areas, of only the SP portions, and of only the DP portions, and observed that DP sub‐regions yielded the greatest number of inter‐region connections, suggesting that SP and DP voxels are engaged to different degrees in resting‐state functional networks. Finally, we observed significant correlations between specific behavioral scores and power spectral measurement in DP voxels in selected regions of interest. These findings together suggest there may be distinct time courses in WM BOLD signals during a resting state that reflect underlying variations in the anatomical, neurovascular and functional couplings in different populations of voxels.

## Results

### Spatial distributions of Independent Components and their characteristic power spectra

WM was decomposed into 80 independent components (ICs) using the group ICA approach (see methods for details) based on images acquired from 199 subjects. Each IC represents a cluster of voxels that exhibit similar patterns of BOLD signals over time. The power spectral frequency distributions of the signals from within the voxels in each IC were calculated by a Fourier transform. Figure 1 shows selected WM ICs and their power spectra in separate panels. For each panel, the first figure (I) shows the IC in three orthogonal planes in MNI space (spatial distributions of all 80 ICs can be found in supporting file 1). The second figure (II) in each panel shows the power spectra of the voxels that compose the IC where each line represents the mean power spectra averaged over 199 subjects for each voxel. Based on the observed patterns of power over frequencies, these lines were clustered using a k‐means algorithm into two groups; SP voxels exhibit single peaks at around 0.015 Hz, while DP voxels show an additional peak at around 0.065 Hz, as shown in the third figure (III) in each panel. The spatial distributions of SP and DP voxels within the IC in transaxial sections are displayed in the fourth figure (IV) in different colors.

**Figure 1.**
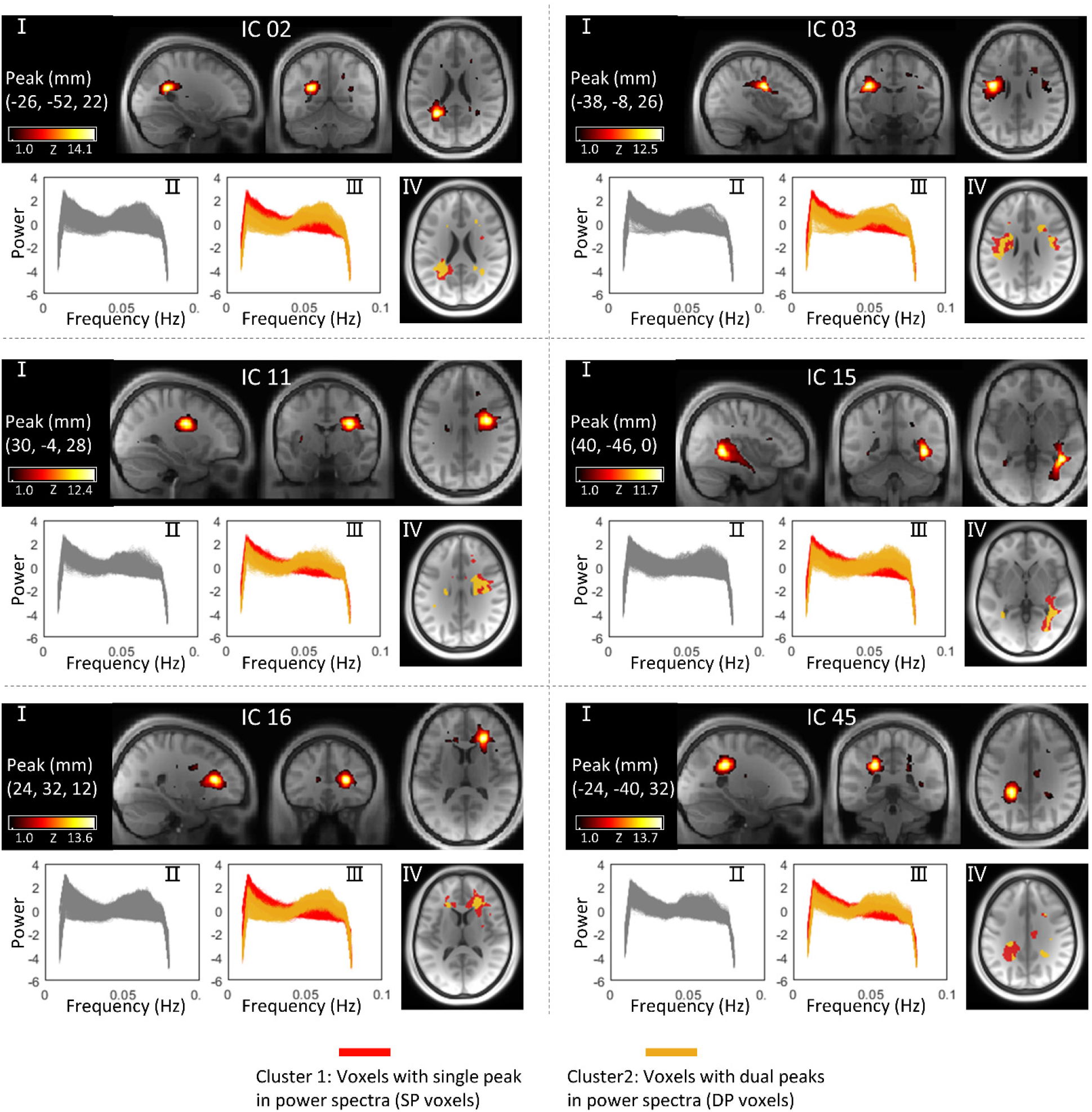
Spatial distributions of selected WM ICs and their power spectra patterns. For each panel: (I) visualization of the WM IC in three orthogonal planes. (II) power spectra of the voxels within the IC. Each line represents the mean power spectra over 199 subjects at the same voxel. (III) Two clusters of voxels (SP voxels and DP voxels) that exhibit distinct patterns of power spectra within the IC. (IV) Spatial distributions of SP voxels and DP voxels in different colors.

We observed the presence of DP voxels in 80 out of 80 WM ICs (see support file 2 for details. Note that the power spectra were not normalized to unit variance in support file 1 in which the second peaks were not visually obvious for a few ICs). To assess whether the DP voxels are unique to WM, we applied the same workflow to GM but observed no DP voxels in any of the 80 GM ICs (see details in support file 3). To examine the reproducibility of this work, the classifications were replicated on two resting sessions acquired from the same 199 subjects on different days. As shown in Figure 2, the spatial distributions of SP and DP voxels were in close agreement between the tests in WM ICs. Figure 3 shows global maps of SP and DP distributions, reconstructed by combining voxels in each category for the 80 ICs, and their reproducibilities were evaluated in terms of the Dice coefficients between the test and retest results. The Dice coefficients were 0.95 for SP voxels and 0.78 for DP voxels.

**Figure 2.**
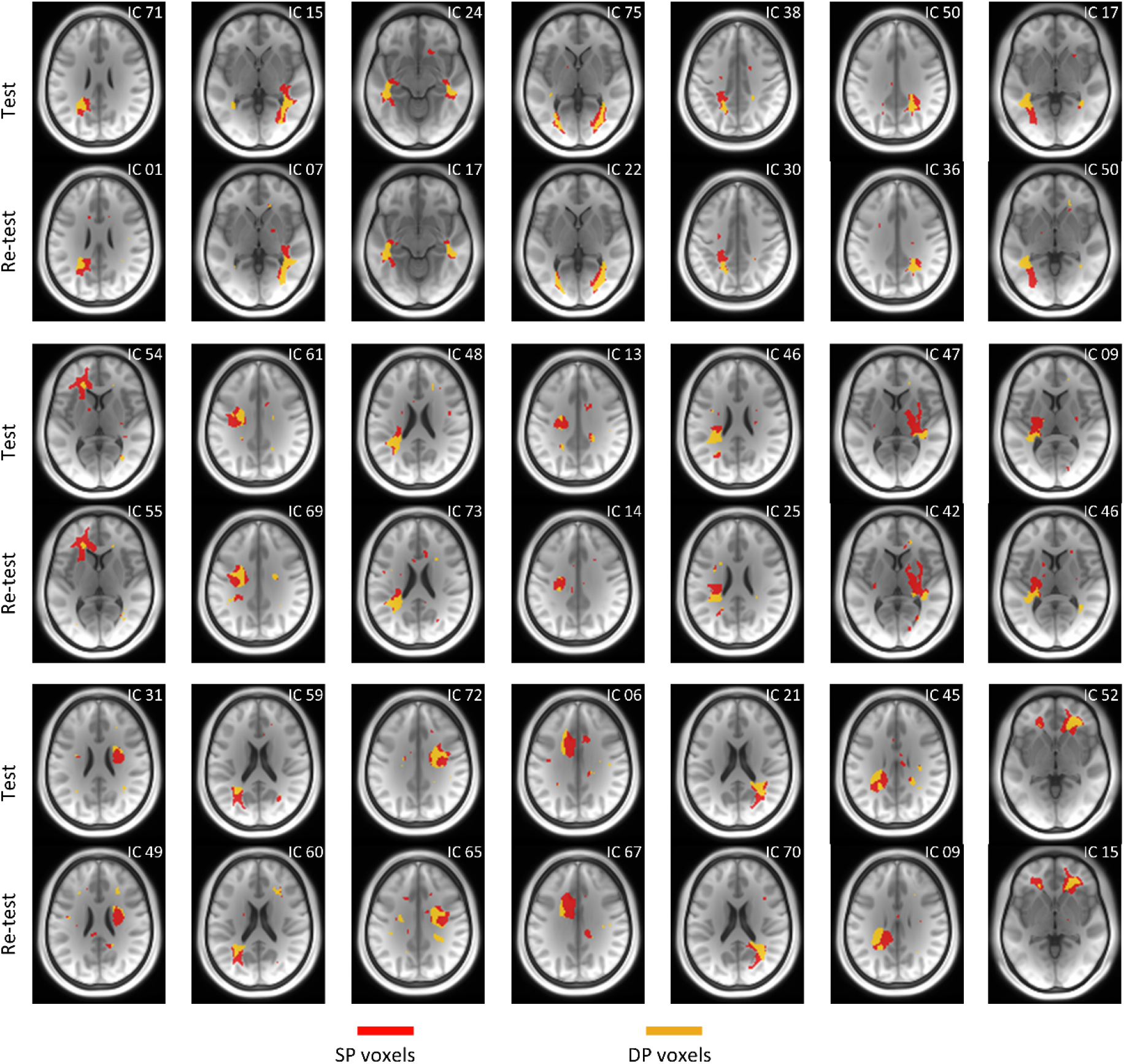
Test‐ retest reproducibility of the spatial distribution of SP and DP voxels in selected WM ICs. The test and retest spatial maps are calculated using the same workflow and based on two resting sessions acquired from same subjects over a period of two days.

**Figure 3.**
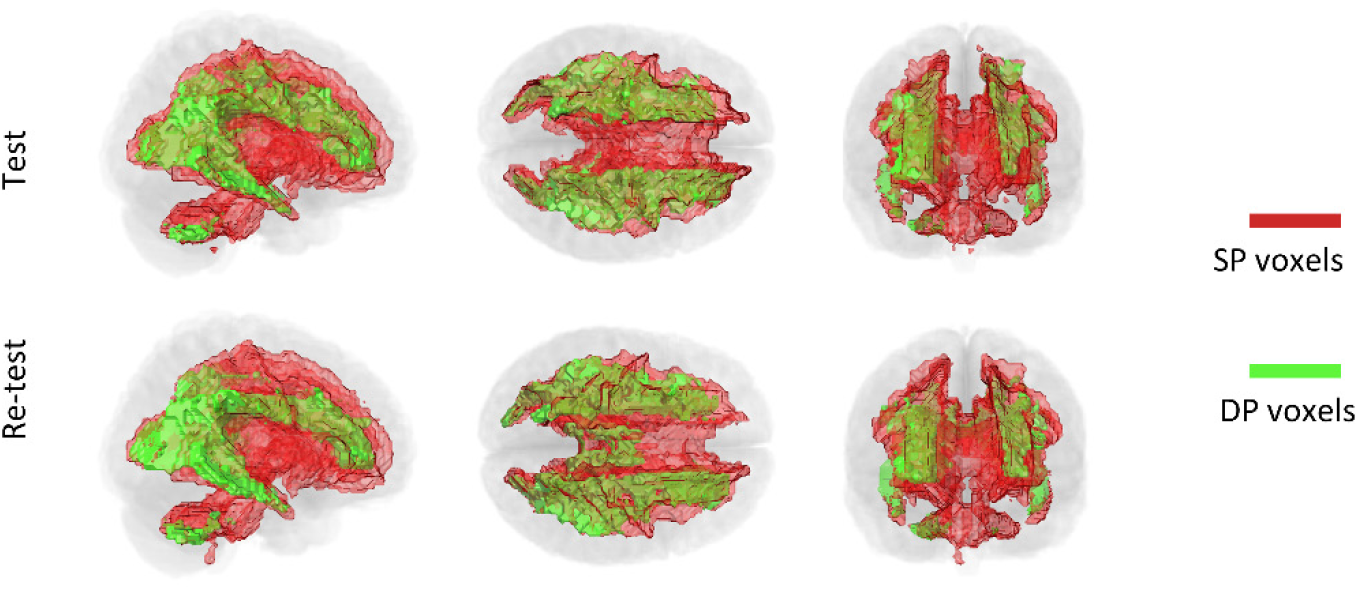
The global maps of SP and DP voxels. Maps derived from test and retest data are displayed in upper row and lower row, respectively.

### The relationship between HRFs and power spectra in WM ICs

HRFs were estimated using the RS‐HRF toolbox (see methods for details) for SP and DP voxels separately in each IC. For each panel in Figure 4, the left figure shows the distribution of SP and DP voxels in the IC. The middle figure displays the HRFs of SP and DP voxels in different colors, where each line represents the HRF averaged over 199 subjects at the same voxel. We observed that the DP voxels exhibit more prominent initial dips whose magnitudes are significantly lower (p<0.05, Bonferroni corrected) compared with SP voxels in 80 out of 80 ICs. To further confirm that this difference was not observed by chance, we randomly divided each IC (on a subject level) into two groups and observed that the HRFs fully overlapped between the two random groups, as shown in the right figure of each panel.

**Figure 4.**
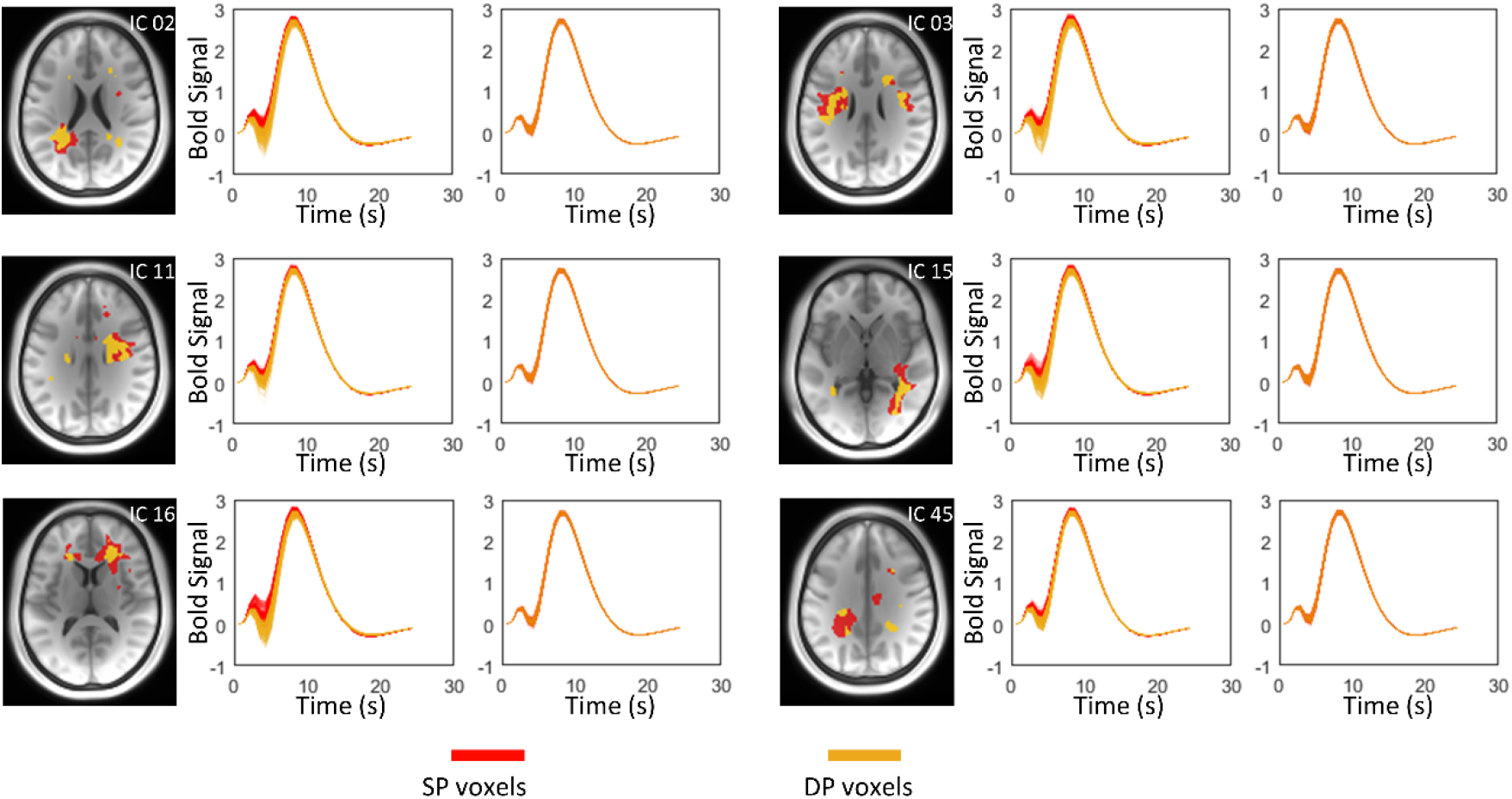
The estimated HRFs in SP and DP voxels in selected WM ICs. For each panel (IC): the left figure visualizes the spatial distribution of SP and DP voxels within the IC. The middle figure visualizes the estimated HRFs of SP and DP voxels respectively in two colors. The right figure represents the estimated HRFs from voxels that are randomly divided into two groups. Note that the random group 1 and 2 consists of same number of voxels as the SP and DP.

The magnitudes of initial dips, particularly in DP voxels, appear to vary with location. For example, the magnitudes of dips for IC 16 are much lower than those for IC 45 as shown in Figure 4. Likewise, the power spectra exhibit variable second peaks across ICs as shown in Figure 5b. To examine the relationship between HRFs and power spectra for DP voxels, we correlated the magnitudes of the initial dips in HRFs and four measurements that varied between power spectra across 80 ICs, including the magnitudes of the first and second peaks, along with their ratios, and the frequencies associated with the second peaks. We observed that the magnitudes of the initial dips of the HRFs significantly correlated with all four measurements as shown in Figure 5 c‐f. The highest correlation was found between the magnitudes of dips and the ratio of the two peaks in the power spectra, as shown in Figure 5c.

**Figure 5.**
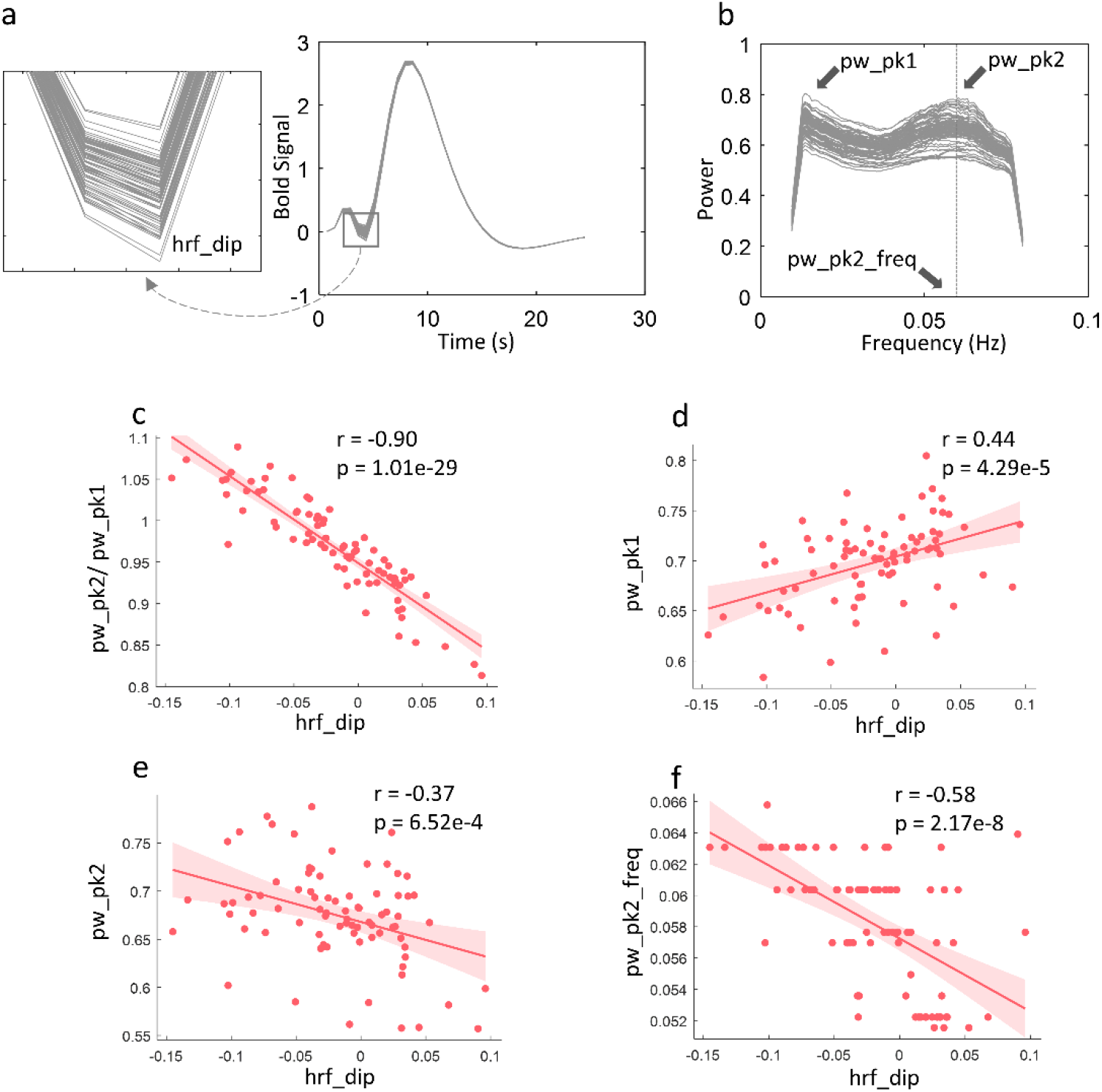
The relationship between HRFs and power spectral in DP voxels across 80 WM ICs. Estimated HRFs of 80 WM ICs. (b) Power spectra of 80 WM ICs. (c) Correlation between the dips of HRFs and the ratio the two peaks in power spectra across 80 WM ICs. (d) Correlation between the dips of HRFs and the magnitudes of first peaks in power spectra across 80 WM ICs. (e) Correlation between the dips of HRFs and the magnitudes of second peaks in power spectra across 80 WM ICs. (e) Correlation between the dips of HRFs and the coordinates of second peaks on frequency band across 80 WM ICs.

### The relationship between white matter tracts and power spectra patterns in DP voxels

The relationship between anatomical configurations and different power spectra in DP voxels was examined by first comparing the spatial distribution of the DP voxels and metrics of fiber complexities calculated from diffusion data acquired from the same 199 subjects (see methods for details) across the entire WM. As shown in Figure 6, these two maps are highly consistent with each other, showing overlapping areas in the corona radiations, posterior thalamic radiations, and sagittal stratum, where commissure tracts and major longitudinal tracts, such as the inferior longitudinal fasciculus, inferior fronto‐occipital fasciculus, and fronto‐occipital fasciculus, also pass through.

**Figure 6.**
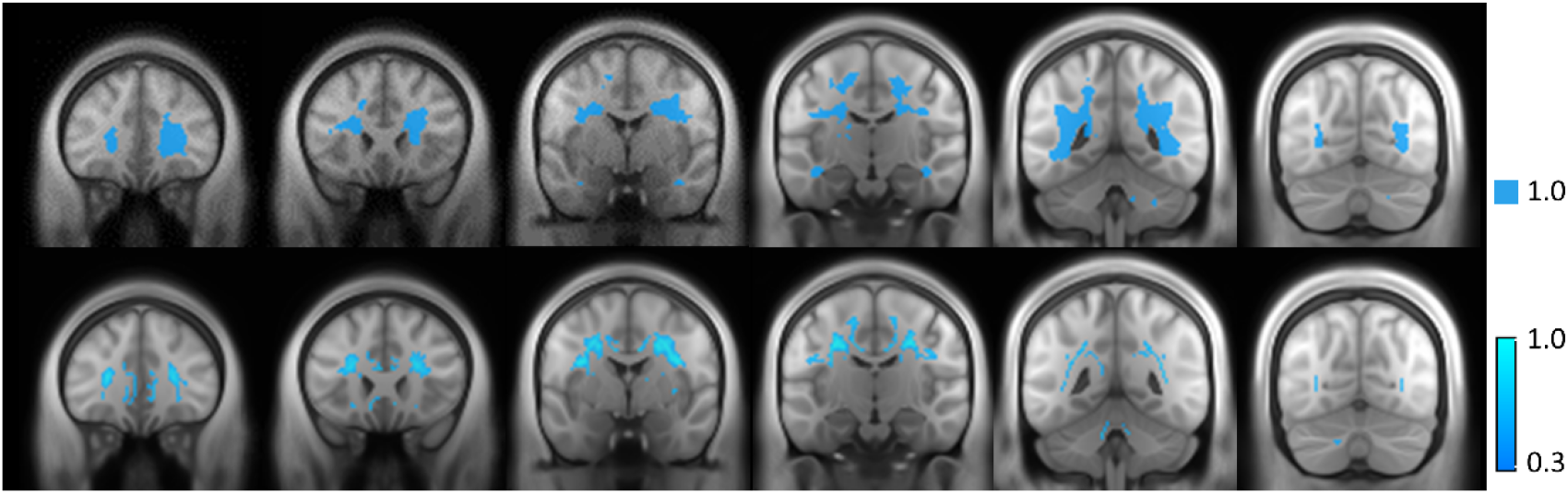
Spatial distribution of DP voxels (upper row) and the fiber complexity (lower row) over the brain WM.

To quantify this relationship further, we compared fiber complexity between SP and DP areas in each IC across 199 subjects. We observed, as shown in Figure 7, that 64 out of 80 (80%) of ICs show significantly higher complexity (p<0.05, Bonferroni corrected) in the DP area than that in the SP voxels, indicating that the second peak in the power spectra corresponds to voxels with more crossing fibers.

**Figure 7.**
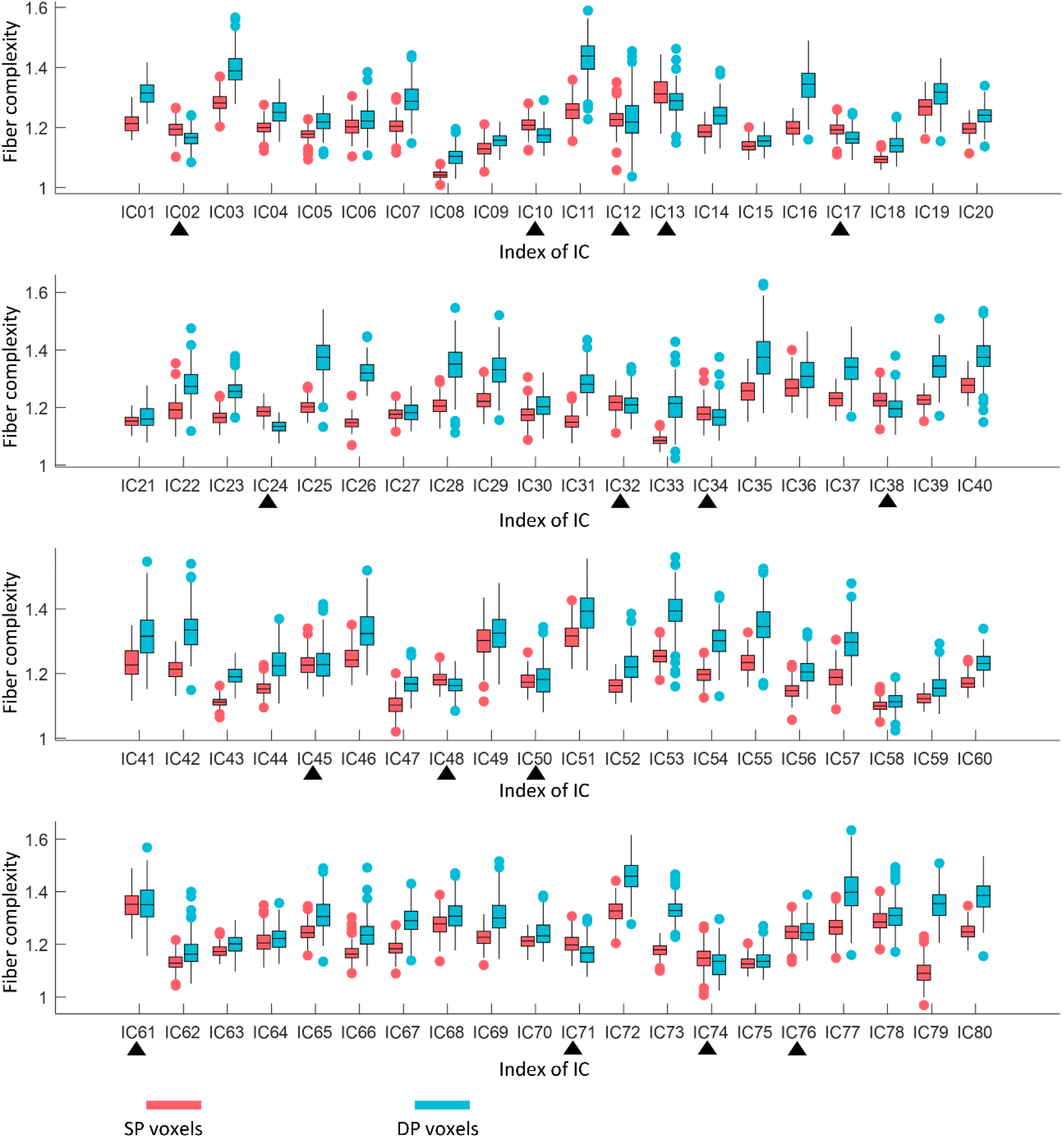
Comparison of fiber complexity between SP and DP areas in 80 WM ICs. 64 out of 80 ICs shows significantly higher fiber complexity in DP area compared to SP area. Black triangles indicate non‐significant differences.

### Functional correlations based on entire IC, SP voxels, and DP voxels

In a resting state, the BOLD signals from white matter regions show correlations with other white and grey matter areas in a manner similar to the correlations used to infer functional connectivity between cortical volumes ^21^. We, therefore, constructed the resting state matrices showing the correlations between each pair of the 80 ICs for each entire component as well as only the SP and DP portions respectively. Figure 8 a‐c shows the DP ‐based matrices exhibit the highest connectivities. To quantify the differences, we compared the number of links, i.e., the sum of the binarized matrix when applying different thresholds to the matrix, for 199 subjects. It can be seen in Figure 8d that the matrices based on DP areas show a higher number of links than those using SP or the entire IC across all thresholds we selected.

**Figure 8.**
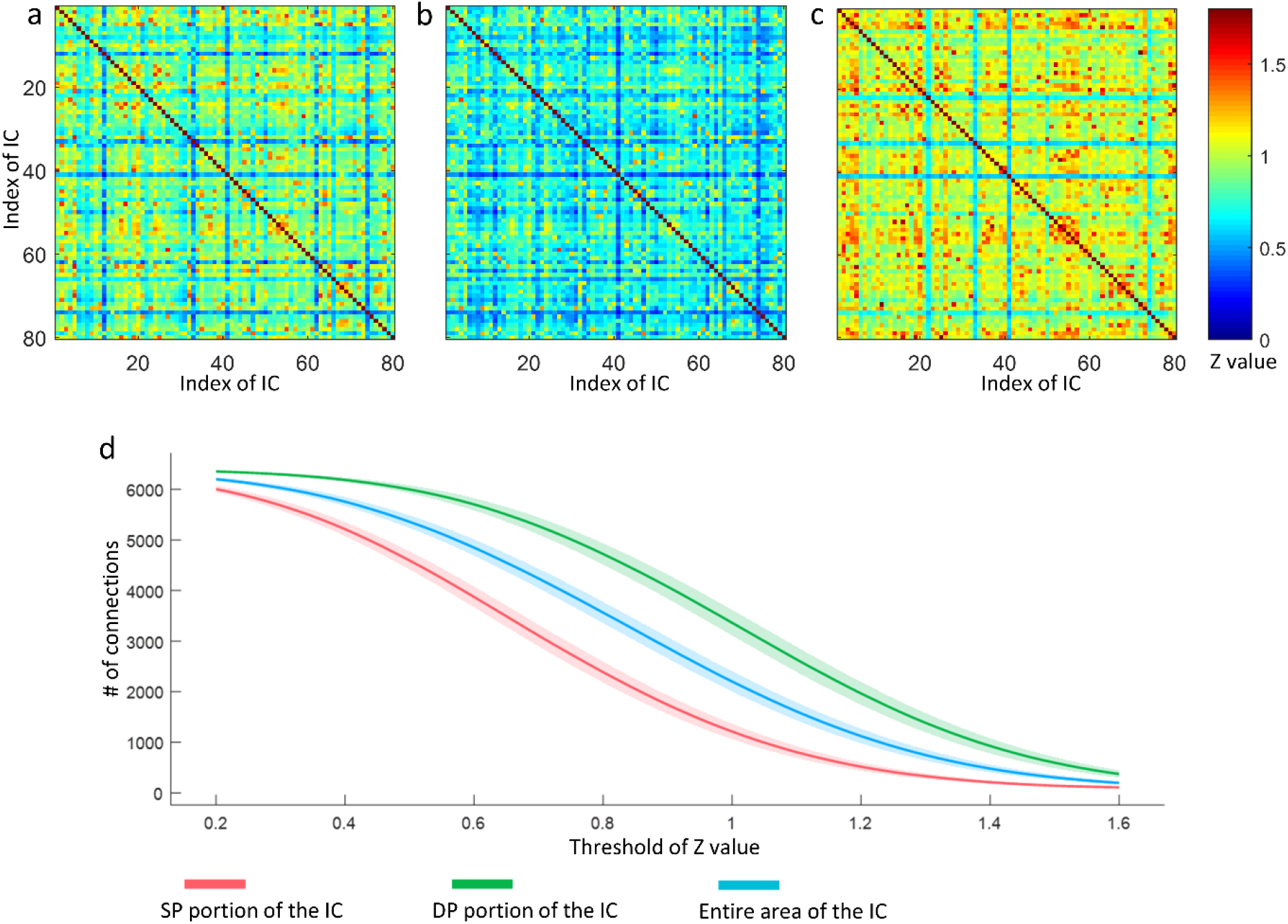
Comparison of FC matrix reconstructed by (a) entire areas of ICs, (b) SP portions of ICs and (c) DP portions of ICs. (d) Number of connections when applying different threshold to the FC matrix regarding 199 subjects.

### Relationship between power spectra measurements and behavioral scores

The database we used for image analysis also contained measures of cognitive performance and behavior. We evaluated the relationship between specific behavioral scores and the power spectral peaks for DP voxels in each IC across 199 subjects Figure 9 shows that episodic memory scores were significantly (p<0.05) correlated with the ratio between the two peaks for IC 39 (overlapped with anterior coronal radiation left, anterior limb of internal capsule left and superior corona radiation left). Reading decoding scores were significantly (p<0.05) correlated with the ratio between the two peaks for IC 54 (overlapped with anterior corona radiation right). Note that there are other ICs that showed significant (p<0.05) correlations with behavior scores. Here we report those with the highest correlation coefficient with behavior.

**Figure 9.**
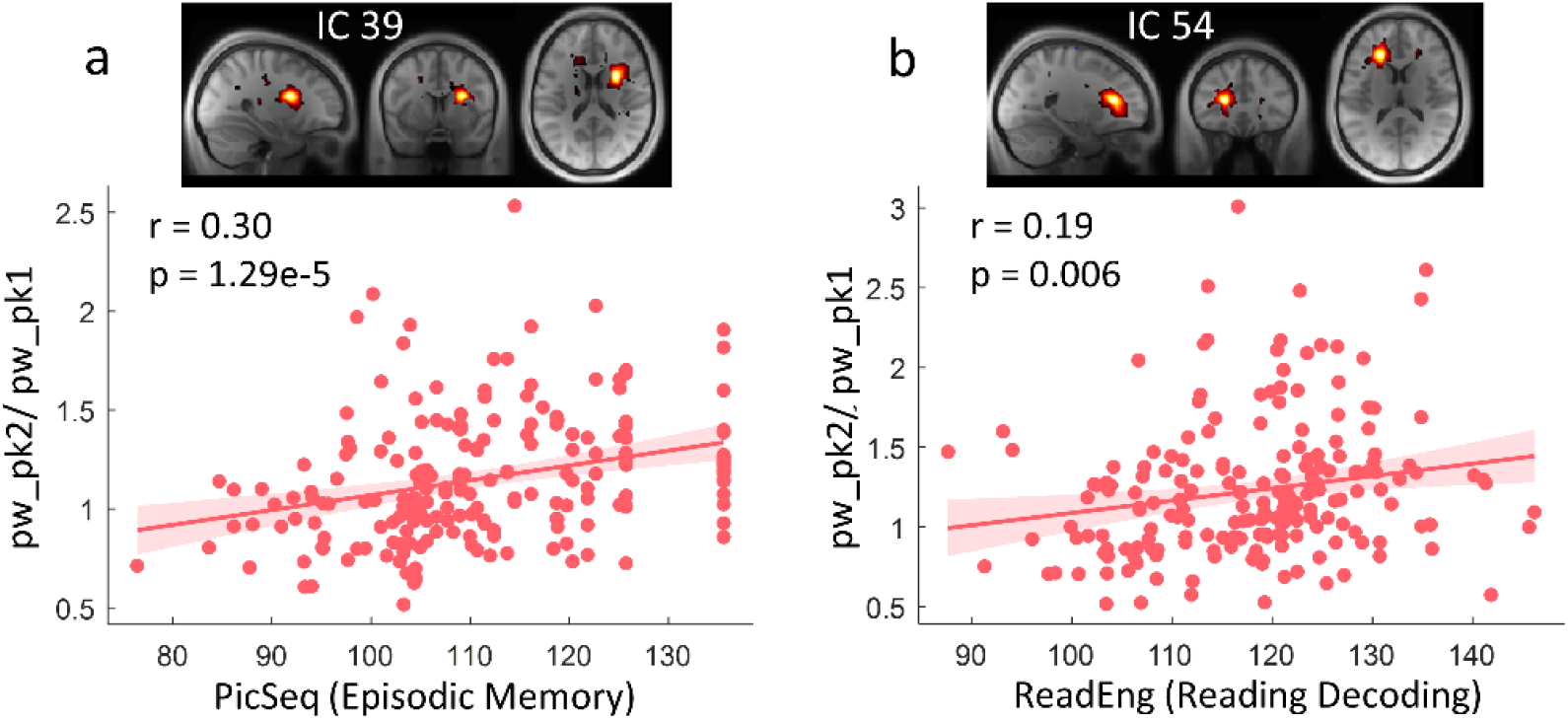
The relationship between behavior scores and power spectral peaks in DP voxels across 199 subjects. (a) Correlation between Episodic Memory Test Score and the ratio the two peaks in power spectra across 199 subjects. (b) Correlation between Reading Decoding Test Score and the ratio the two peaks in power spectra across 199 subjects.

## Discussions

We have measured the power spectra of resting‐state BOLD signals in areas identified as functionally coherent and examined in detail the signal power within a low‐frequency band. We identified two categories of voxels, namely, SP and DP voxels, that exhibited distinct spectral distributions. Specifically, SP voxels, analogous to GM, presented a single‐peaked spectrum, whereas DP voxels exhibit an additional peak at a higher frequency. SP and DP voxels were clustered in certain locations where distinct HRFs were also observed. DP voxels showed significantly lower initial dips compared to SP voxels, and the magnitudes of initial dips differed among ICs and are significantly correlated with the magnitudes of the ratio between two peaks in powers. In addition, the distributions of SP and DP voxels corresponded with the detailed structure of WM within the IC, as shown by the significant differences in fiber complexities in SP and DP voxels. Moreover, inter‐IC correlations evaluated using different components of the ICs suggest that the time courses of signals from DP voxels within an IC are synchronous with a higher number of other ICs compared to the entire IC or the SP portions, demonstrating their different engagement in brain connectivity. Last, the power spectral measurements in voxels with two peaks in specific components predict specific human behaviors.

Different frequency bands in power spectra may correspond to distinct physiological processes, Biswal et al. for the first time reported that low‐frequency spontaneous BOLD signals reflect neurophysiological activity, laying the foundation for extensive resting‐state fMRI studies ^2^. Since then, power spectra have been frequently used to characterize spontaneous activities in different functional units or among study groups. However, dual‐peak patterns within the low‐frequency range, as we reliably observed in DP voxels, have not been previously reported in GM. In addition, while recent studies have reported significant changes in the amplitude of low-frequency fluctuations (ALFF) in WM in patients with schizophrenia and autism spectrum disorder ^33,34^, these findings were restricted to the intensity of spontaneous neural activity. The patterns of power spectra over the frequency range of interest correspond to variations in underlying HRF characteristics and may add insights to the functional mechanism underlying the association between signal features, neurovascular coupling, and anatomical substrates.

Despite the fact that time courses were similar within the same IC according to the criteria of ICA, a clear separation of two categories of voxels can be observed within each IC based on the power spectra patterns. These voxels were reliably detected at locations within each IC, and DP voxels tended to distribute in deeper areas of WM, where the hemodynamic environment differ more from those closer to GM. Specifically, deep WM tends to be more distant from supply vessels so that it takes a longer time to compensate for any increase in oxygen consumption ^35^. As observed in this study, HRFs of DP voxels exhibited more prominent initial dips compared to SP voxels, which is consistent with our previous findings regarding depth‐dependent changes in WM HRFs based on an event‐related task ^17^. The lower dips of DP voxels may represent early focal increases in oxygen consumption, which induce longer‐lasting negative BOLD signals than in SP voxels before the arterial supply increases flow sufficiently to meet the tissue demand. Alternately, it may arise if larger boluses of deoxygenated venous blood drain from nearby active GM. These more drastic changes in BOLD signals over a certain period of time presumably correspond to the increased magnitudes in high‐frequency spectral components. Quantitatively, for DP voxels, the magnitudes of the second peaks in power are negatively correlated with the magnitudes of the initial dips in HRFs over 80 ICs, proving their linkage with the neurovascular conditions.

The distribution of DP and SP voxels appear to correspond to structural variations within WM as characterized by diffusion‐based tractography. Previous studies have suggested that structural alterations, such as a lesion or demyelination, can induce BOLD changes in WM ^20–23,36^ so changes in patterns of power spectra, as well as HRF, may indicate WM integrity or WM complexity. By visual inspection and according to prior knowledge, DP voxels are mainly distributed in areas where multiple WM tracts cross. Quantitative measures confirmed that WM complexity was strongly related to the distribution of SP and DP voxels, and a major proportion ICs showed significantly higher complexity in the DP areas than in SP areas. WM areas with higher complexity are interpreted as being responsible for signal transmission along multiple neural pathways, potentially influencing local BOLD effects and causing lower dips in HRFs and increased high‐frequency power in DP voxels compared to SP voxels.

The inter‐IC connectivities evaluated separately for SP and DP sub‐regions may reflect the extent to which these two categories of voxels are differentially engaged in functional integration in the brain. Our data suggest that the number of correlations between DP voxels was significantly greater than for SP voxels. This is consistent with the premise that DP voxels correspond to fiber‐crossing areas, in which the BOLD signal time courses represent the combination of contributions from multiple tracts, and which therefore are synchronous with (and dependent on) a greater number of ICs compared with SP voxels.

In summary, we characterized the power spectra of the resting‐state time courses in 80 ICs that distribute over the WM. A clear separation of two categories of voxels is evident in each IC based on their power spectral patterns. These voxels are location‐specific, and their distributions in each IC are related to underlying anatomical structures. Moreover, they differed in their involvement in apparent functional connectedness in the brain as judged from resting‐state correlations. Taken together, these findings add to the existing understanding of WM BOLD changes during resting state and reveal a strong structural‐vascular‐functional association in WM.

## Supporting information

support file 1

support file 2

support file 3

## Acknowledgments

This work was supported by the National Institutes of Health (NIH) grant R01 NS093669 (J.C.G) and R01 NS113832 (J.C.G), and Vanderbilt Discovery Grant FF600670 (Gao). Imaging data were provided by the Human Connectome Project, WU‐Minn Consortium (Principal Investigators: David Van Essen and Kamil Ugurbil; 1U54MH091657) funded by the 16 NIH Institutes and Centers that support the NIH Blueprint for Neuroscience Research; and by the McDonnell Center for Systems Neuroscience at Washington University.

We declare no conflict of interest.

## Methods

### HCP data

199 subjects were randomly selected from the HCP S1200 release ^37^. They were healthy young adults, 87 M, and 112 F, whose ages ranged between 22 and 35 years. The images used in the present study include two sessions of resting‐state fMRI acquired on two separate days, T1 weighted MRI and diffusion MRI. The imaging protocols are described in detail in previous work ^37^. Briefly, data were acquired using a 3T Siemens Skyra scanner (Siemens AG, Erlanger, Germany). The resting‐state data were acquired using multiband gradient‐echo echo‐planar imaging (EPI). Each session consists of two runs (left‐to‐right and right‐to‐left phase encoding) of 14 min and 33 s each, repetition time (TR) = 720 ms, echo time (TE) = 33.1 ms, voxel size = 2 mm isotropic, number of volumes = 1200. Physiological data, including cardiac and respiratory signals, were recorded during fMRI scanning. The diffusion MRI included 6 runs (9 min and 50 s each) that were acquired using a multiband spin‐echo EPI, representing 3 different gradient combinations, with each acquired once with right‐to‐left and left‐to‐right phase encoding polarities, TR=5520 ms, TE =89.5 s, voxel size =1.25 mm isotropic. Diffusion weighting consisted of 3 shells of b=1000, 2000, and 3000 s/mm^2^ interspersed with an approximately equal number of acquisitions on each shell within each run. T1 images were acquired using a 3D magnetization‐prepared rapid acquisition with gradient echo (MPRAGE), TR = 2400 ms, TE = 2.14 s, voxels size = 0.7 mm isotropic.

Assessments of cognitive ability in the HCP data include tasks from the University of Pennsylvania Computerized Neurocognitive Battery ^38^, and the Blueprint for Neuroscience Research ‐ funded NIH Toolbox for Assessment of Neurological and Behavioral function (http://www.nihtoolbox.org). We selected behavior scores corresponding to four tasks, including episodic memory, reading decoding, spatial orientation, and Inhibition, for evaluation.

### Preprocessing

The images drawn from the HCP repository were preprocessed through the minimal preprocessing pipelines (MPP) as detailed elsewhere ^39^. Briefly, T1 images were nonlinearly registered to MNI space using FNIRT ^40^ and underwent a Freesurfer pipeline which produced surface and volume parcellations as well as morphometric measurements ^41^. For fMRI, the pipeline included motion correction, distortion correction using reversed‐phase encoding directions, and nonlinear registration to MNI space. We performed additional processing of the images that were sourced from the HCP database, including regression of nuisance variables, including head movement parameters (using one of the outputs of motion correction in the MPP pipeline), and cardiac and respiratory noise modeled by the RETROICOR approach ^42^, and followed by a correction for linear trends and temporal filtering with a band‐pass filter (0.01 – 0.08 Hz). As the analyses were restricted to WM, a group‐wise WM mask was reconstructed by averaging the WM parcellations that were derived from Freesurfer across all subjects and thresholded at 0.9. Then the fMRI data were spatially smoothed within the WM mask with a 4 mm FWHM Gaussian kernel. For comparison, data were also smoothed within a GM mask that was reconstructed in a similar manner but using a lower threshold (0.6) due to higher individual variabilities in GM. For diffusion images, the MPP pipeline included a b0 intensity normalization, EPI distortion correction using reversed‐phase encoding directions, and coregistration to T1 native space.

### Group ICA and power spectra of the ICs

ICA is capable of decomposing data into ICs that are assumed to make up the data by an unknown, but linear, mixing process. In this study, the GIFT toolbox was used to analyze the data ^43^. To estimate 80 ICs, the temporal dimension of each subject was reduced from 1200 to 120 (1.5 times the target number of ICs) using spatial principal component analysis (PCA). Those PCs were then concatenated along their temporal dimensions across 199 subjects to evaluate signal changes over 199*120 dynamics. The group data again underwent a PCA, further having its dimension reduced from 120 to 80, producing PCs that accounted for maximal variances on the group level, from which 80 ICs were subsequently estimated using infomax ICA ^44^. The spatial map (group‐level) of each IC was reconstructed and converted to z‐scores and thresholded at z>1. Note that the z‐score here has no statistical interpretation but is used for descriptive purposes only ^45^. The ICs were overlaid back as masks on each subject’s data to extract time courses of interest, from which power spectra were estimated using a multi‐taper Fourier transform ^46^. The power spectra for each voxel in each IC were averaged over 199 subjects and assigned to SP (if a single peak was observed) or DP (if double peaks were observed) using a k‐means clustering algorithm (two clusters, 20 iterations) based on their patterns. Before clustering, the power spectra were normalized to unit variance. Otherwise, the clusters may mainly reflect the differences in magnitudes of the powers rather than patterns of power over frequencies.

### Estimation of HRFs

HRFs were estimated from resting‐state timecourses *b(t)* in each subject using a blind deconvolution approach ^47,48^. The method requires no prior hypothesis about the HRF and is based on the notion that relatively large amplitude BOLD signal peaks represent the occurrence of separable, major, spontaneous events. In our study, first, such events were detected as peaks beyond a specified threshold (here, greater than 1.5 standard deviations over the mean). For each event, a general linear model was fitted using a combination of *s*_*n*_*(t)*, the onset of the event, and *h(t)*, which represents a linear combination of two gamma functions and its temporal derivative. Here *n* characterizes the time from the onset to the peaks *s(t)*, where *s(t)* =1 only if *t* corresponds to the peaks (events) we detected. The double gamma functions together with temporal derivative are capable of modeling an initial dip and time delay in the response ^49,50^. By searching for an *n (n∈ 0‐ 12 s)* and minimizing the covariance of the residuals *cov[b(t) – conv(s*_n_*(t), h(t))]*, several parameters that model *h(t)* can be estimated, so the HRF *h*_*n*_*(t)* can be obtained. The relationship between the HRF and power spectra in DP voxels were evaluated in terms of Pearson’s correlation coefficients comparing the magnitudes of initial dips in HRFs and magnitudes of power of the two peaks.

### Estimation of fiber complexity

The analysis was conducted using DSI Studio (http://dsi-studio.labsolver.org). The diffusion data were modeled using generalized q‐sampling imaging, a model‐free reconstruction method that quantifies the density of diffusing water at different orientations ^51^. This derives the spin distribution function (SDF), an orientation distribution function which has been shown to have high sensitivity and specificity to white matter characteristics and pathology. In our study, for each voxel, three fibers were resolved. The SDF includes the orientations estimated for each of the three fibers, as well as their fractions, reflecting the likelihood that a fiber along that orientation can be identified. The fiber complexity was determined by the number of fractions that exceeded a threshold (0.1 in this study). Note that the preprocessed diffusion data sourced from the HCP database were aligned to T1 native space only. For group analysis, the complexity map for each subject was then spatially transformed to MNI space using the “warp information” recorded during the FNIRT registration of T1 images. The differences in fiber complexity between SP and DP voxels were evaluated using a two‐sample t‐test over 199 subjects. Bonferroni‐adjusted p<0.05 was considered significant.

### Connectivity matrix

The connectivity matrices were reconstructed by calculating Pearson’s correlation coefficients between time courses of pair‐wise ICs on the subject level. Each IC was split into two portions, consisting of SP voxels and DP voxels respectively. The matrices regarding the entire area of IC, the SP portion, and DP portion of IC were reconstructed separately, converted to z values using Fisher’s r to z transformation, and averaged across 199 subjects for further assessments. The number of connections equals the sum of the binarized matrix when applying different thresholds to the connectivity matrix.

