## Supplementary material for "Power spectra reveal distinct BOLD resting‐state time courses in white matter": support file 1

### Group ICA Parameters

```
..................................................

Number of Subjects : 199
Number of Sessions : 1
Number of Independent Components : 80
ICA Algorithm : Infomax
Number Of Scans/Timepoints : 1200
Mask File : ICA_mask2
Data Pre-processing Type : Remove Mean Per Timepoint
PCA Type : Standard
Group PCA Type : Subject Specific
Group ICA Type : Spatial
Back Reconstruction Type : GICA
Scaling Components : Z-scores
Stability analysis type : none
Group analysis mode: Serial
Anatomical file: h:\ICA_HCP_20200909\wm_ICs\T1_avg200.nii,1
Slice Plane: Axial
Image values: Positive
Convert to Z-scores: yes
Threshold: 1

..................................................
```

### ICASSO Plots

### Mean Components

Mean across all subjects and sessions is computed for each component

- **a) Timecourse** - Mean timecourse is converted to z-scores.
- **b) Spectra** - Timecourses spectra is computed for each data-set and averaged across sessions. Mean and standard error of mean is shown in the figure.

- **c) Montage** - Axial slices are shown.
- **d) Ortho slices** - Ortho plot is shown for the peak voxel and coordinates are reported.

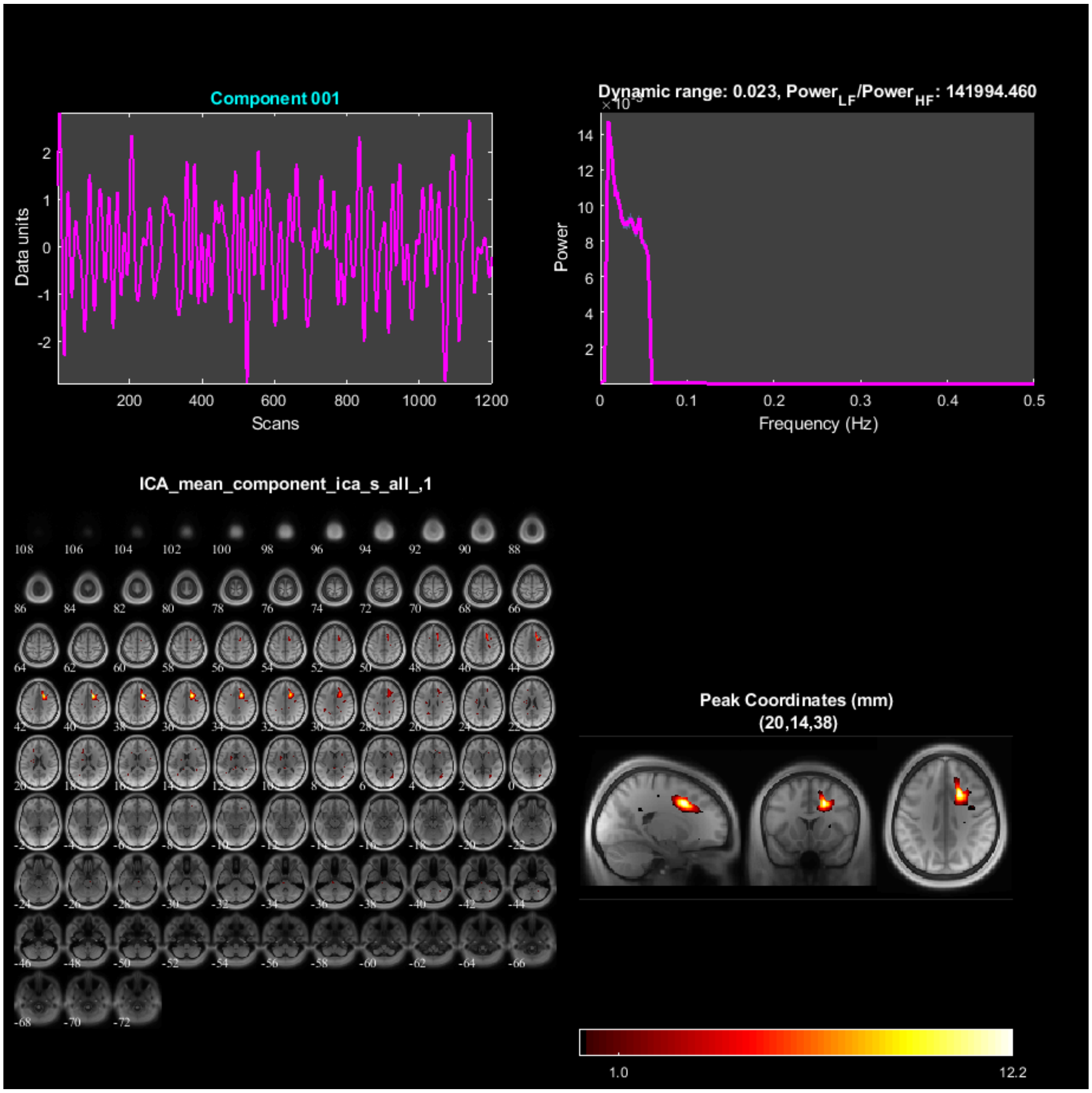

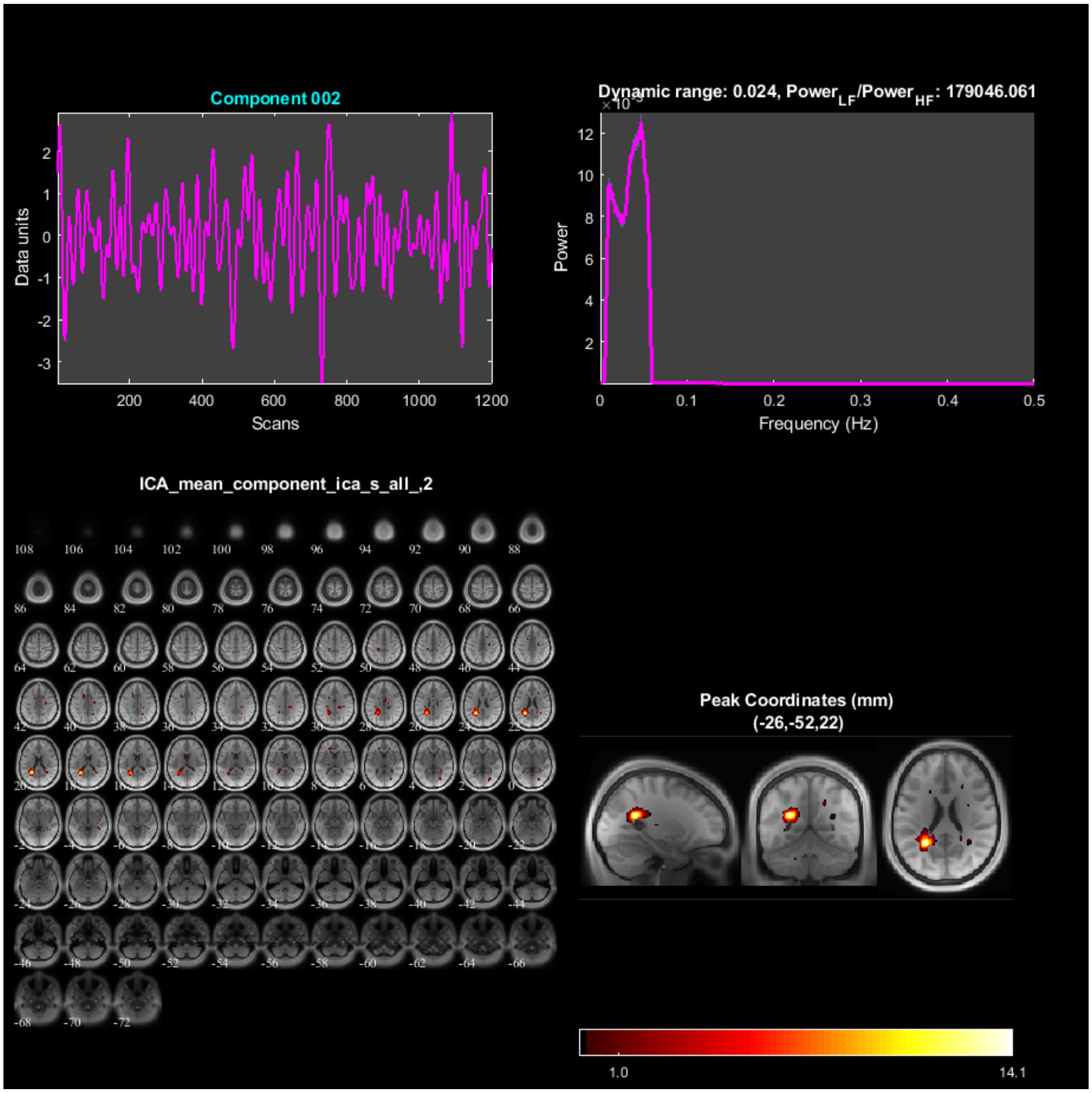
Component 002
Data units
Scans
Dynamic range: 0.024, Power_LF/Power_HF: 179046.061
Power
Frequency (Hz)
ICA_mean_component_ica_s_all_,2
Peak Coordinates (mm)
(-26,-52,22)
1.0
14.1

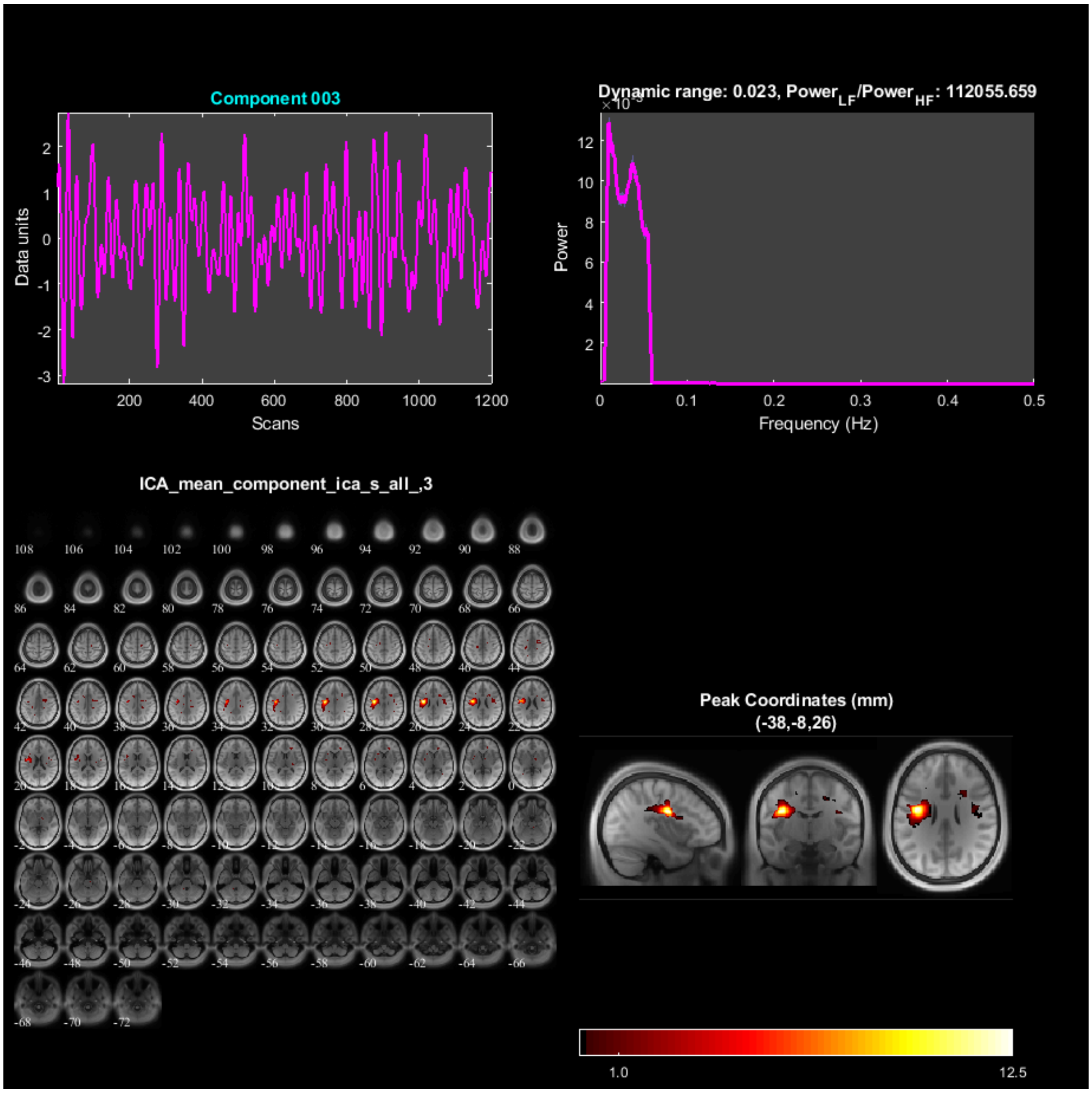

Component 003
Data units
Scans
Dynamic range: 0.023, Power$_{LF}$/Power$_{HF}$: 112055.659
Power
Frequency (Hz)
ICA_mean_component_ica_s_all_,3
Peak Coordinates (mm)
(-38,-8,26)
1.0
12.5

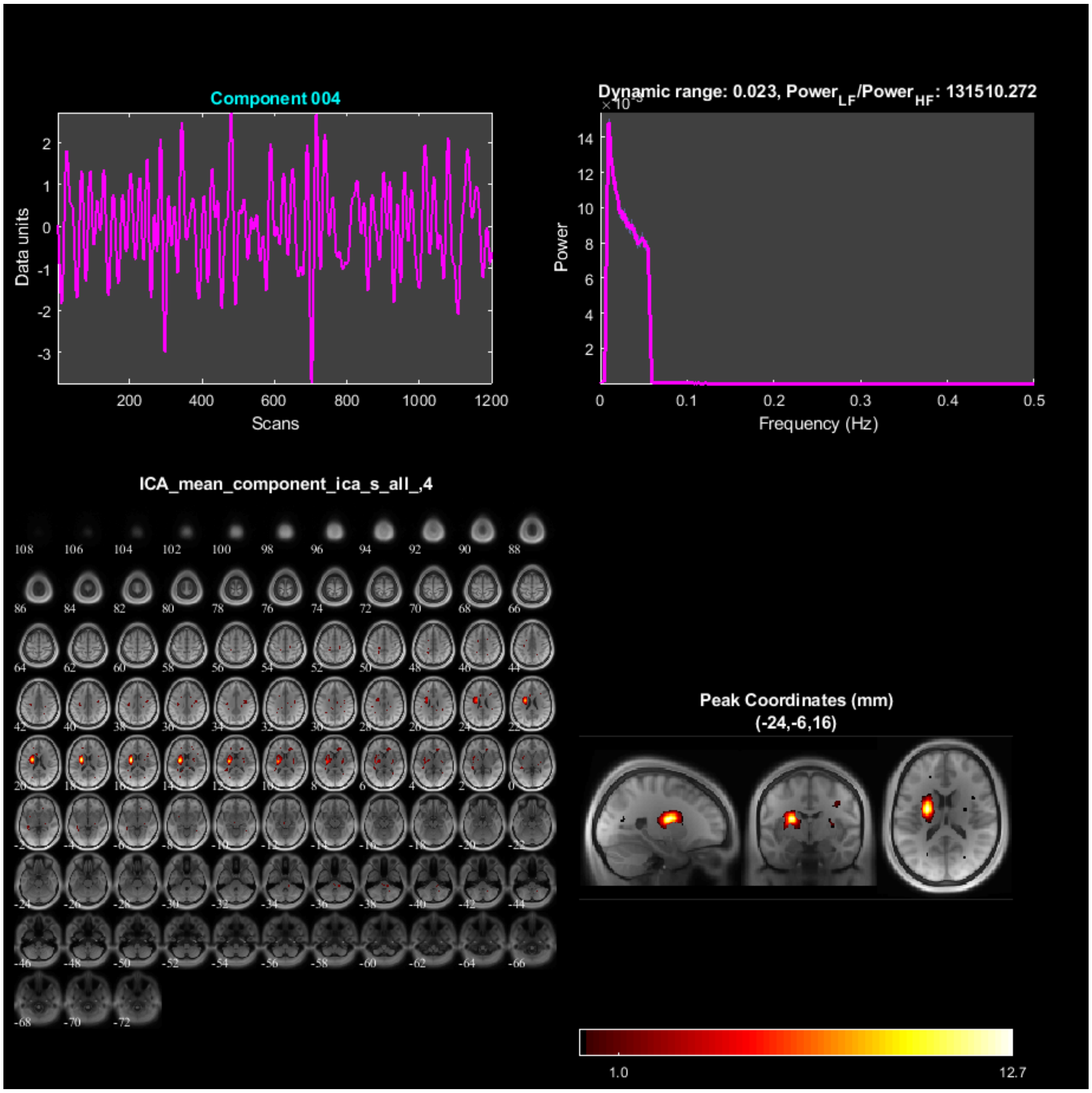
Component 004
Data units
Scans
Dynamic range: 0.023, Power_LF/Power_HF: 131510.272
Power
Frequency (Hz)
ICA_mean_component_ica_s_all_,4
Peak Coordinates (mm)
(-24,-6,16)
1.0
12.7

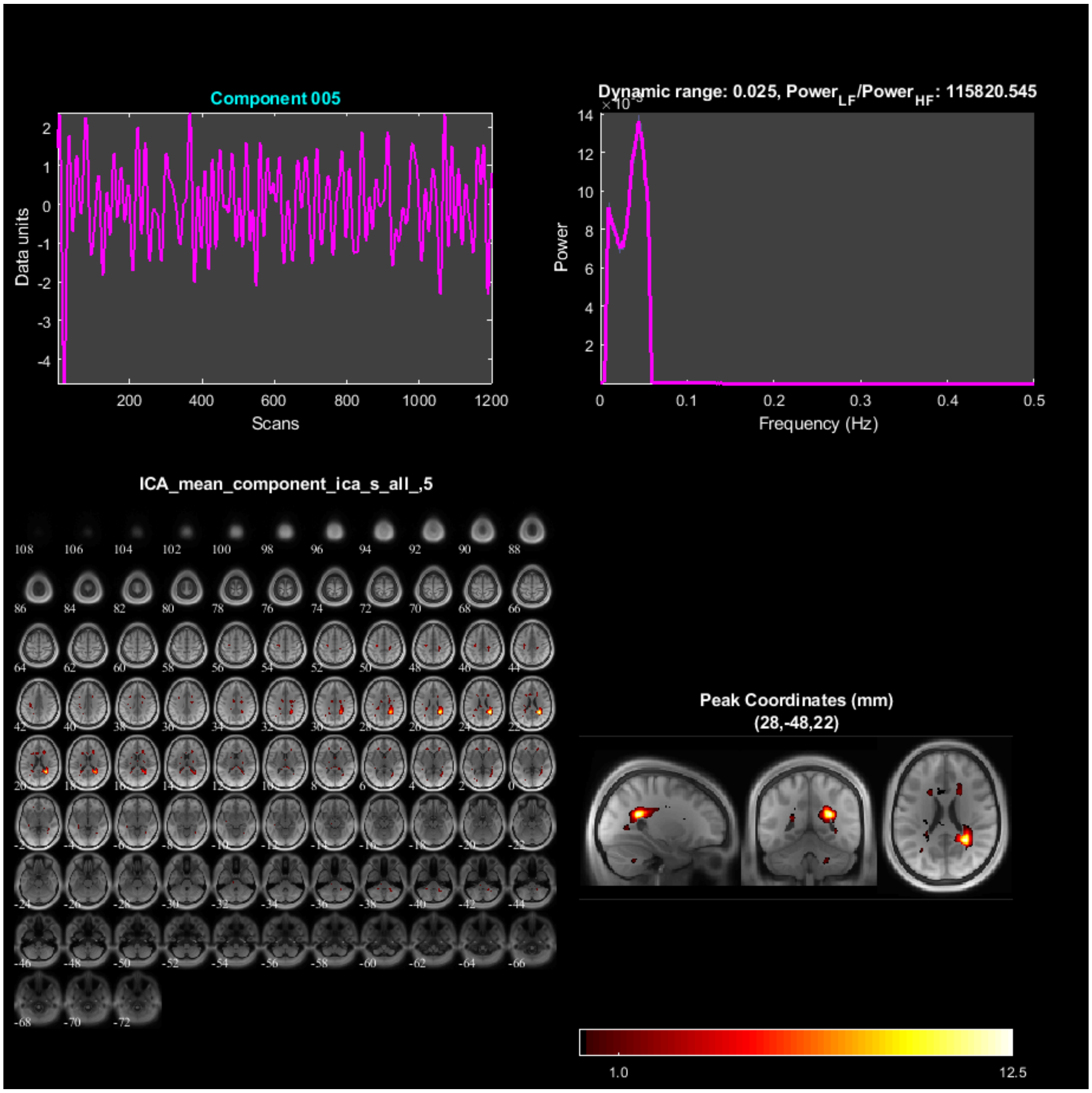
Component 005
Data units
Scans
Dynamic range: 0.025, Power_LF/Power_HF: 115820.545
Power
Frequency (Hz)
ICA_mean_component_ica_s_all_,5
Peak Coordinates (mm)
(28,-48,22)
1.0
12.5

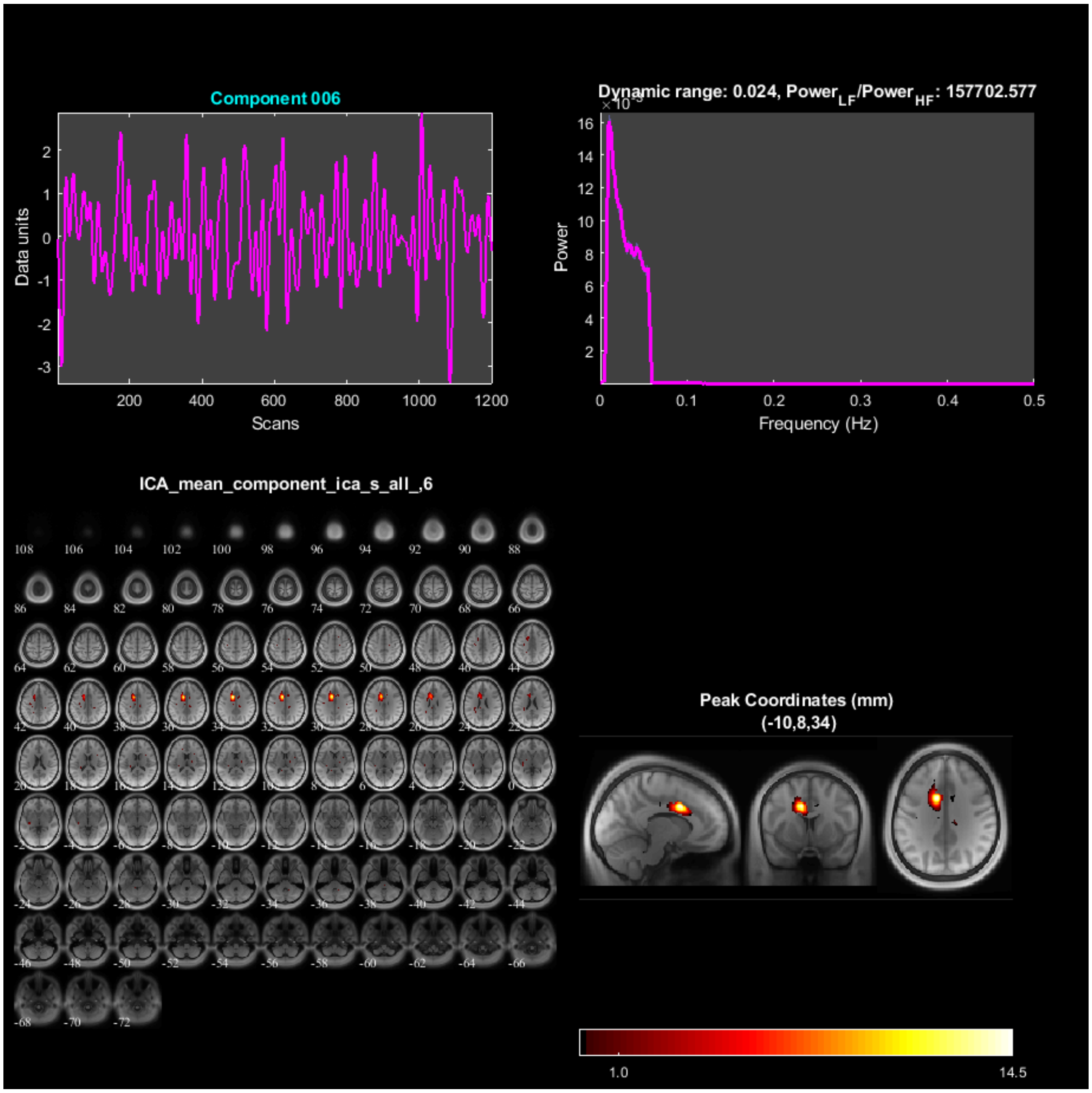

Component 006
Data units
Scans
Dynamic range: 0.024, Power$_{LF}$/Power$_{HF}$: 157702.577
Power
Frequency (Hz)
ICA_mean_component_ica_s_all_,6
Peak Coordinates (mm)
(-10,8,34)
1.0
14.5

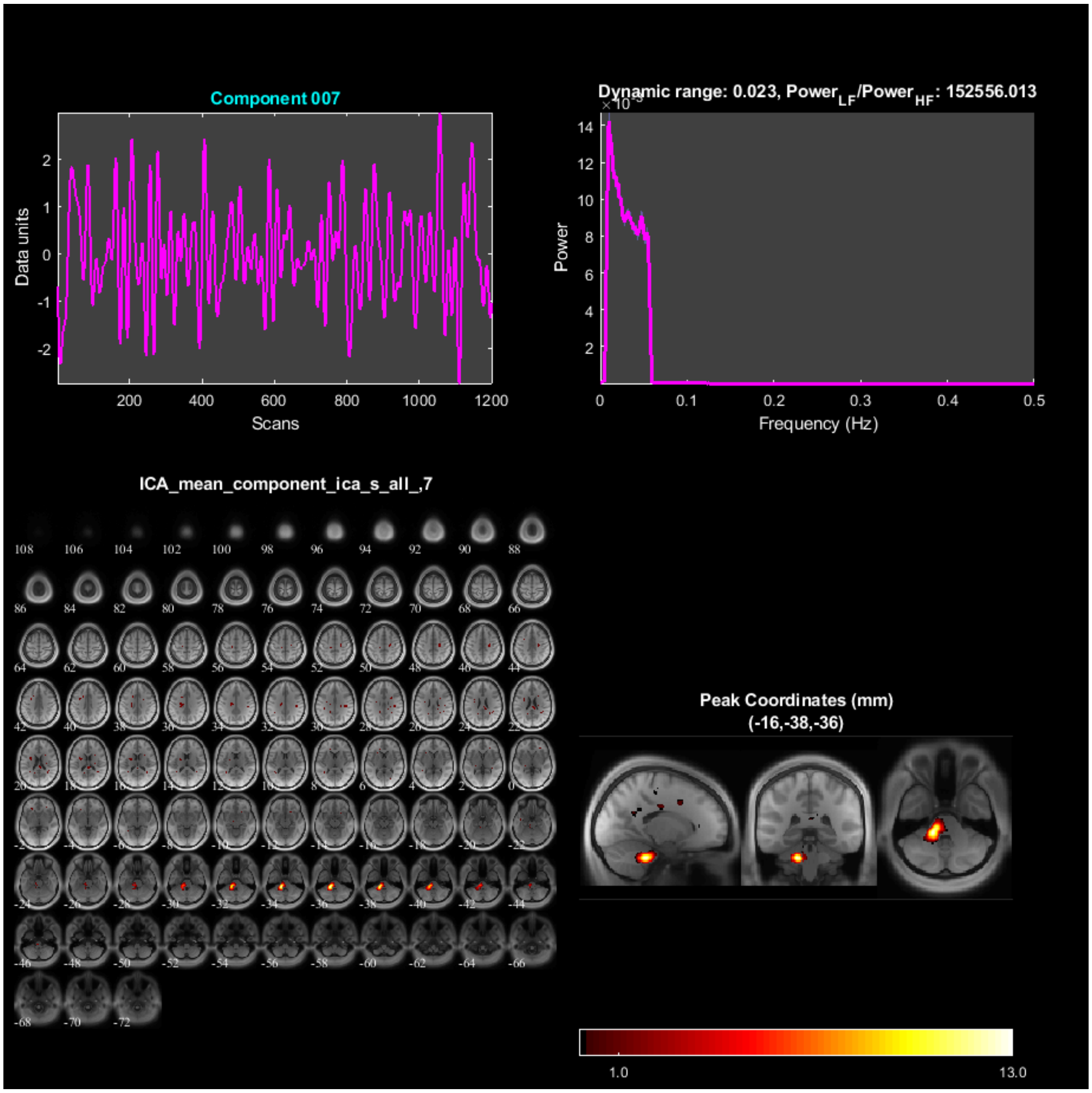

Component 007
Data units
Scans
Dynamic range: 0.023, Power_LF/Power_HF: 152556.013
Power
Frequency (Hz)
ICA_mean_component_ica_s_all_,7
Peak Coordinates (mm)
(-16,-38,-36)
1.0
13.0

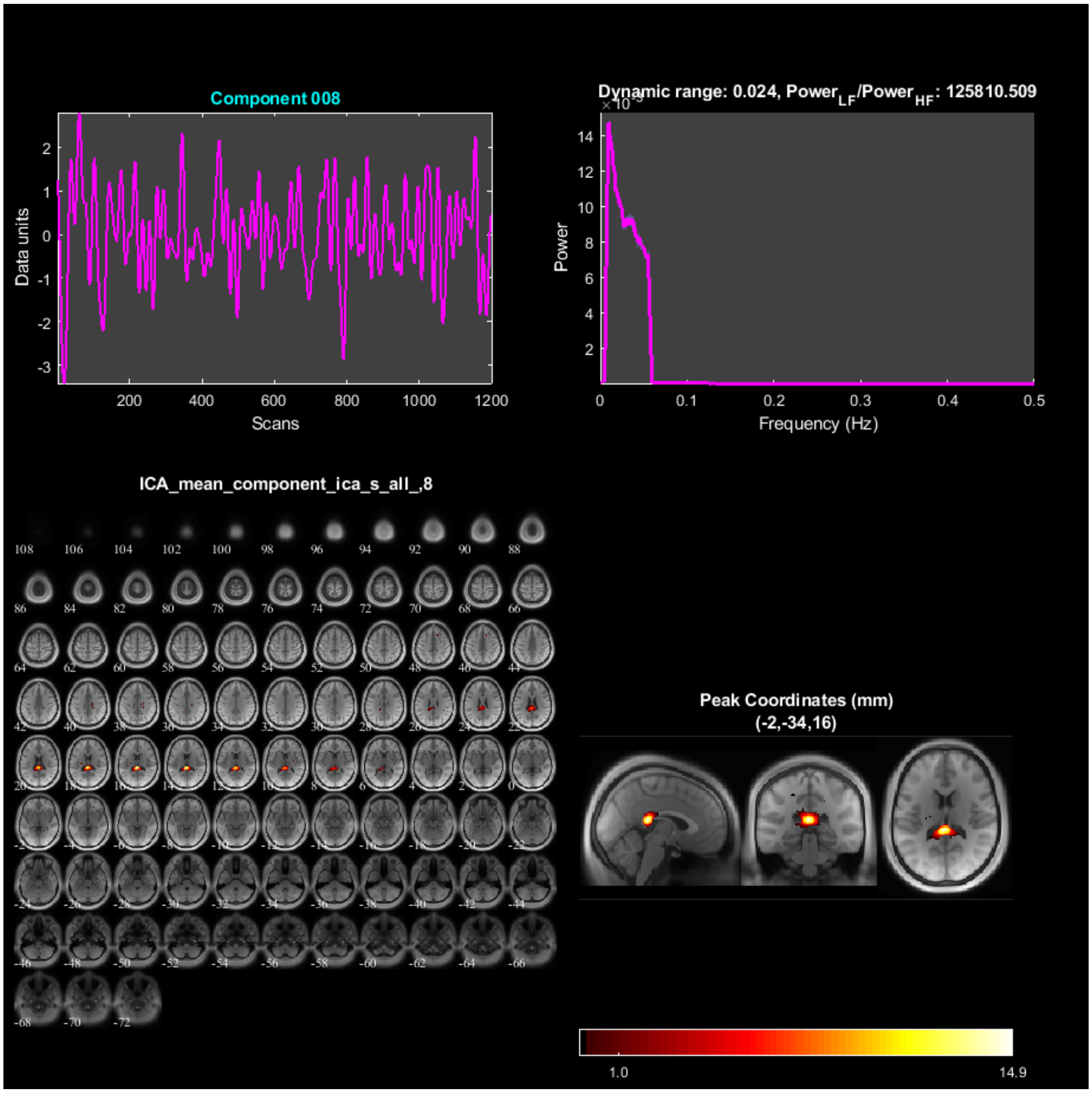
Component 008
Data units
Scans
Dynamic range: 0.024, Power$_{LF}$/Power$_{HF}$: 125810.509
Power
Frequency (Hz)
ICA_mean_component_ica_s_all_,8
Peak Coordinates (mm)
(-2,-34,16)
1.0
14.9

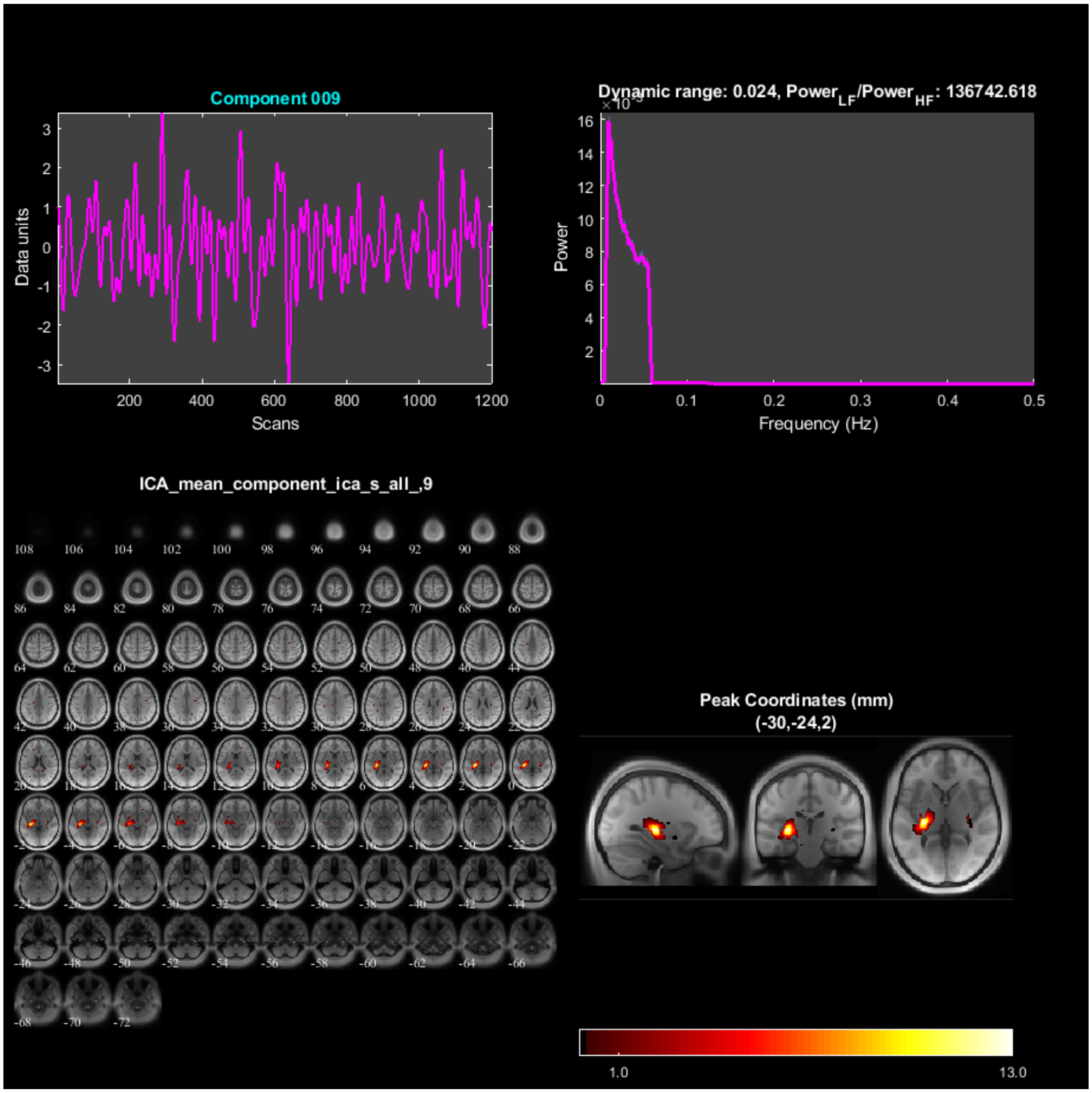

Component 009
Data units
Scans
Dynamic range: 0.024, Power$_{LF}$/Power$_{HF}$: 136742.618
Power
Frequency (Hz)
ICA_mean_component_ica_s_all_,9
Peak Coordinates (mm)
(-30,-24,2)
1.0
13.0

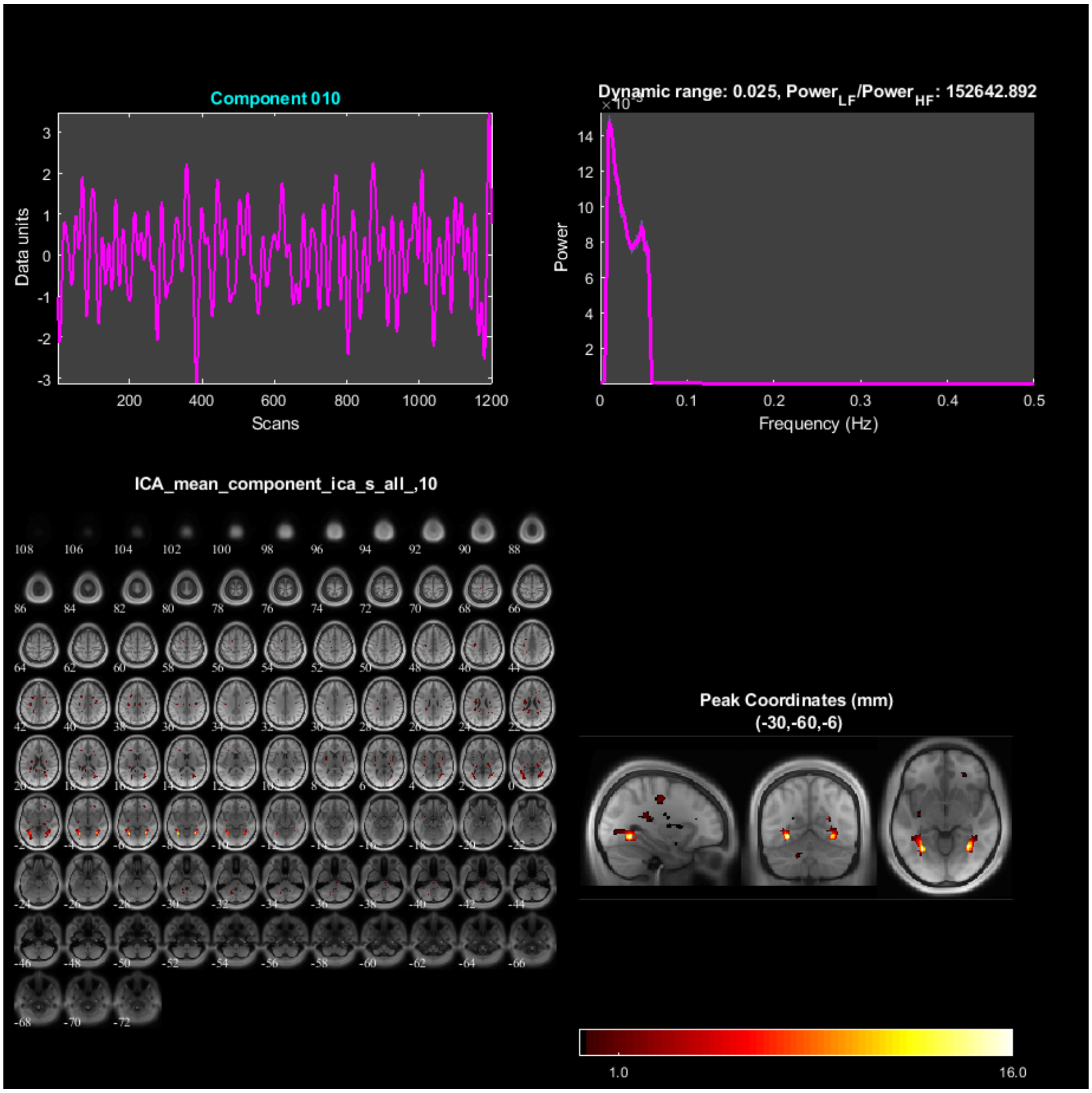

Component 010
Data units
Scans
Dynamic range: 0.025, Power$_{LF}$/Power$_{HF}$: 152642.892
Power
Frequency (Hz)
ICA_mean_component_ica_s_all_,10
Peak Coordinates (mm)
(-30,-60,-6)
1.0
16.0

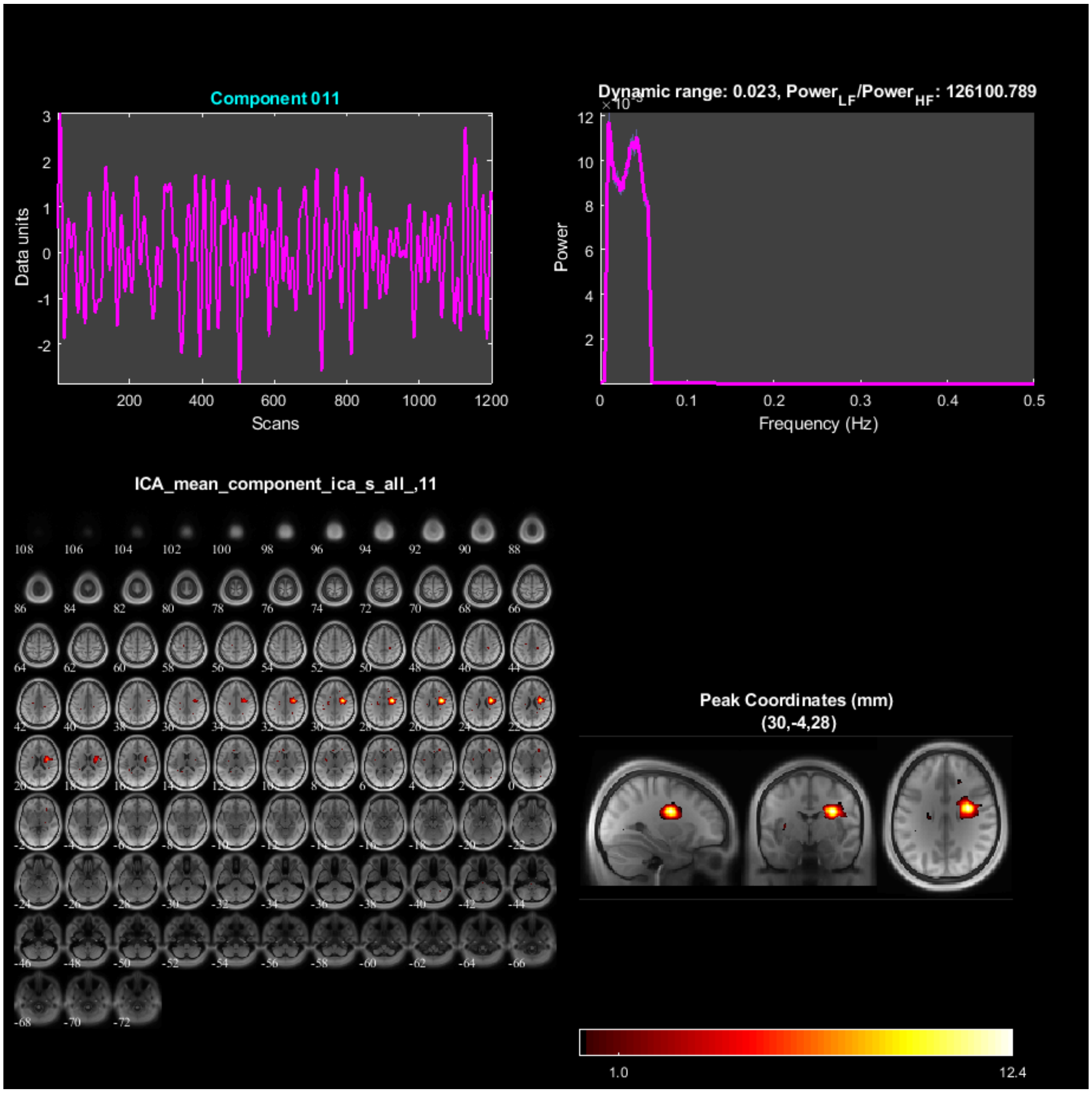
Component 011
Data units
Scans
Dynamic range: 0.023, Power$_{LF}$/Power$_{HF}$: 126100.789
Power
Frequency (Hz)
ICA_mean_component_ica_s_all_,11
Peak Coordinates (mm)
(30,-4,28)
1.0
12.4

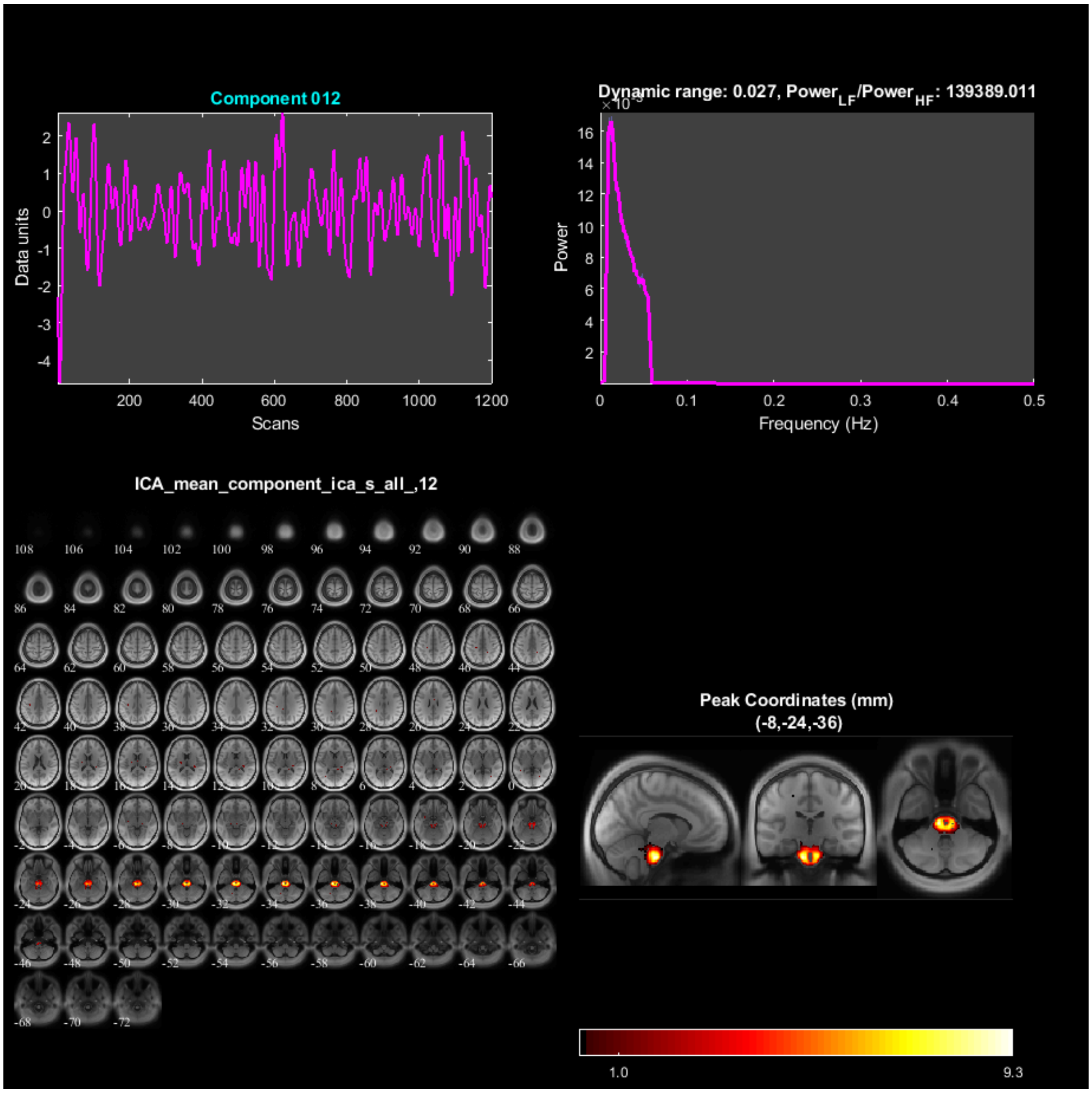

Component 012
Data units
Scans
Dynamic range: 0.027, Power$_{LF}$/Power$_{HF}$: 139389.011
Power
Frequency (Hz)
ICA_mean_component_ica_s_all_,12
Peak Coordinates (mm)
(-8,-24,-36)
1.0
9.3

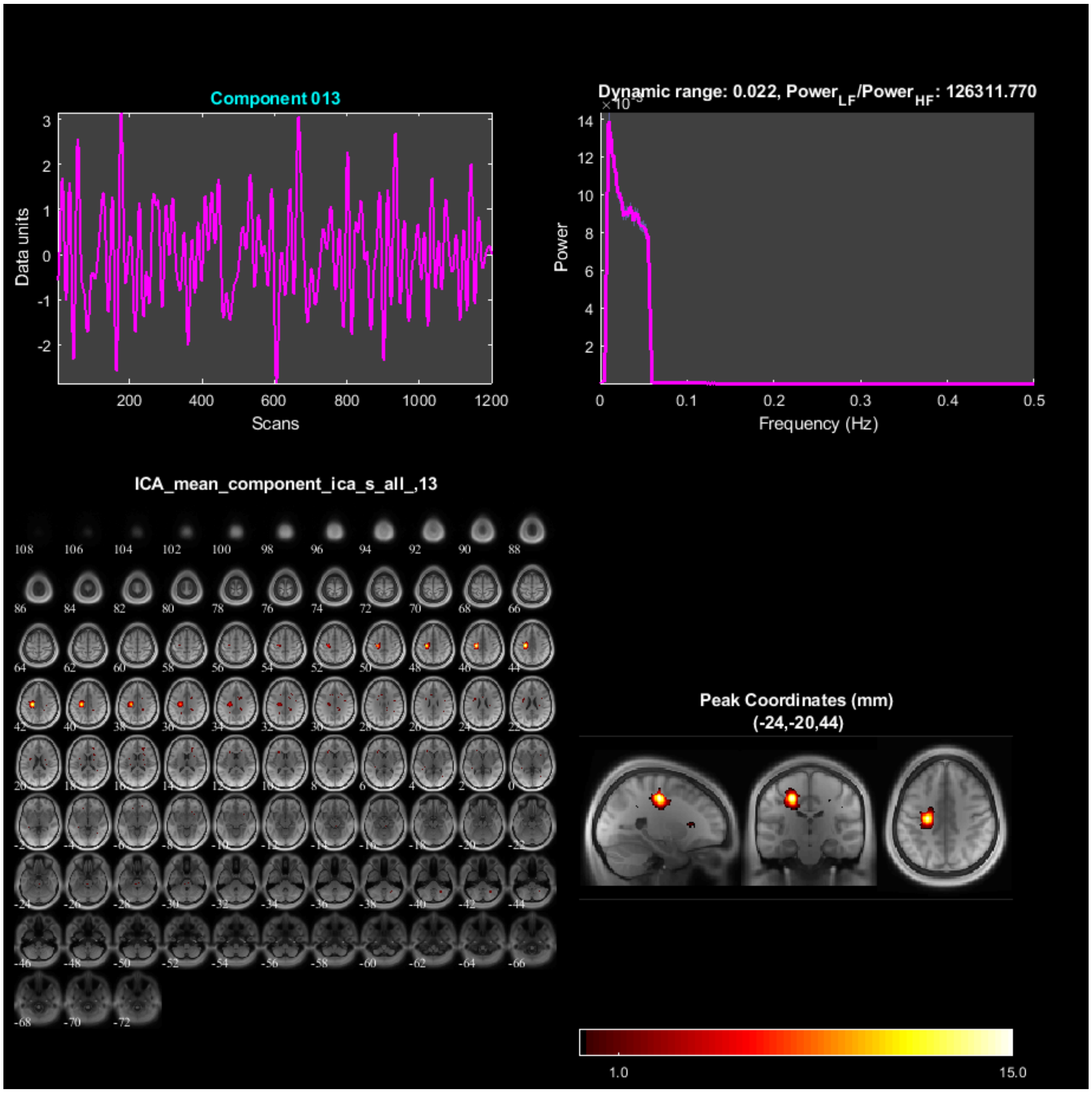
Component 013
Data units
Scans
Dynamic range: 0.022, Power$_{LF}$/Power$_{HF}$: 126311.770
Power
Frequency (Hz)
ICA_mean_component_ica_s_all_,13
Peak Coordinates (mm)
(-24,-20,44)
1.0
15.0

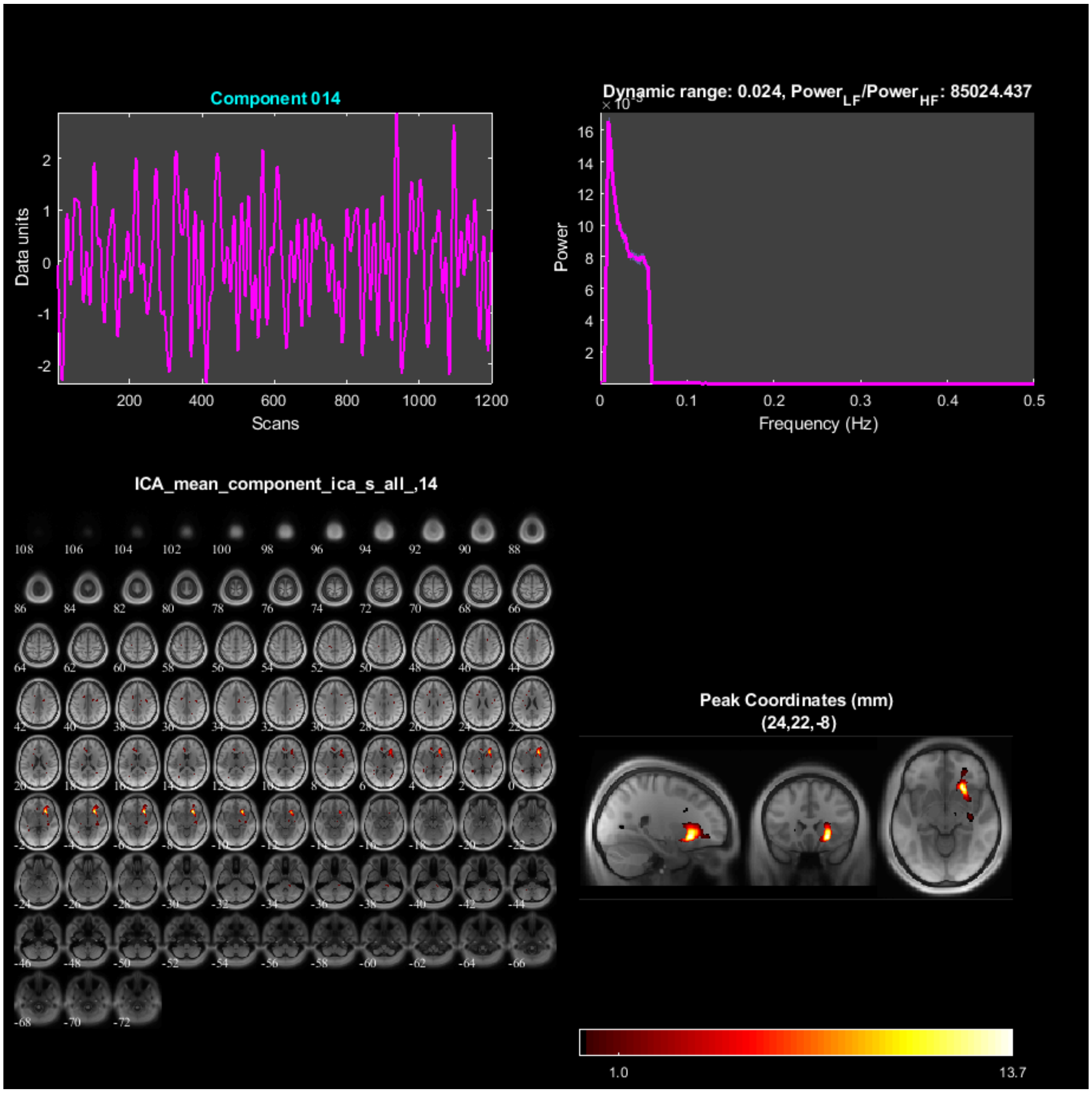

Component 014
Data units
Scans
Dynamic range: 0.024, Power_LF/Power_HF: 85024.437
Power
Frequency (Hz)
ICA_mean_component_ica_s_all_,14
Peak Coordinates (mm)
(24,22,-8)
1.0
13.7

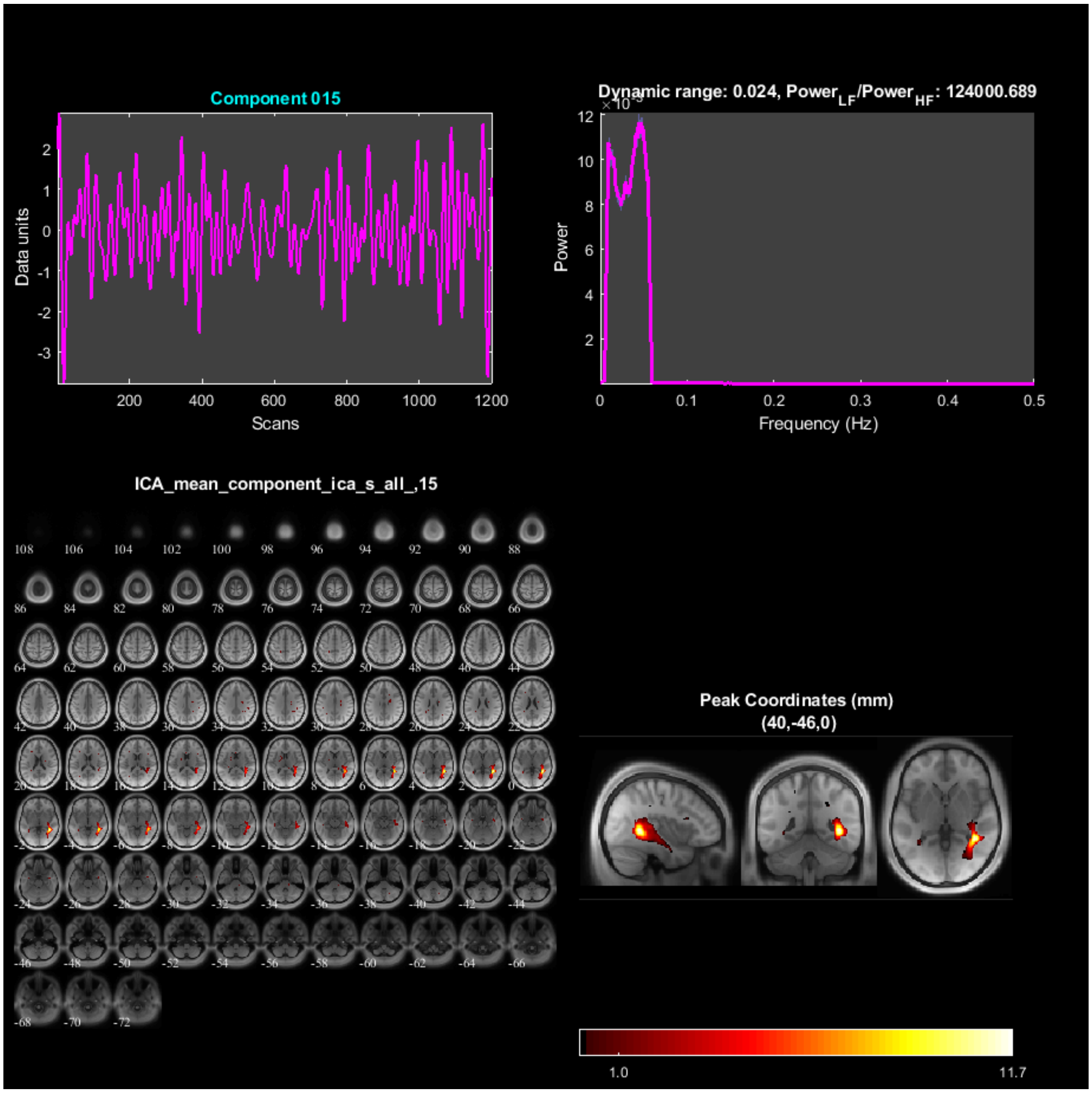
Component 015
Data units
Scans
Dynamic range: 0.024, Power$_{LF}$/Power$_{HF}$: 124000.689
Power
Frequency (Hz)
ICA_mean_component_ica_s_all_,15
Peak Coordinates (mm)
(40,-46,0)
1.0
11.7

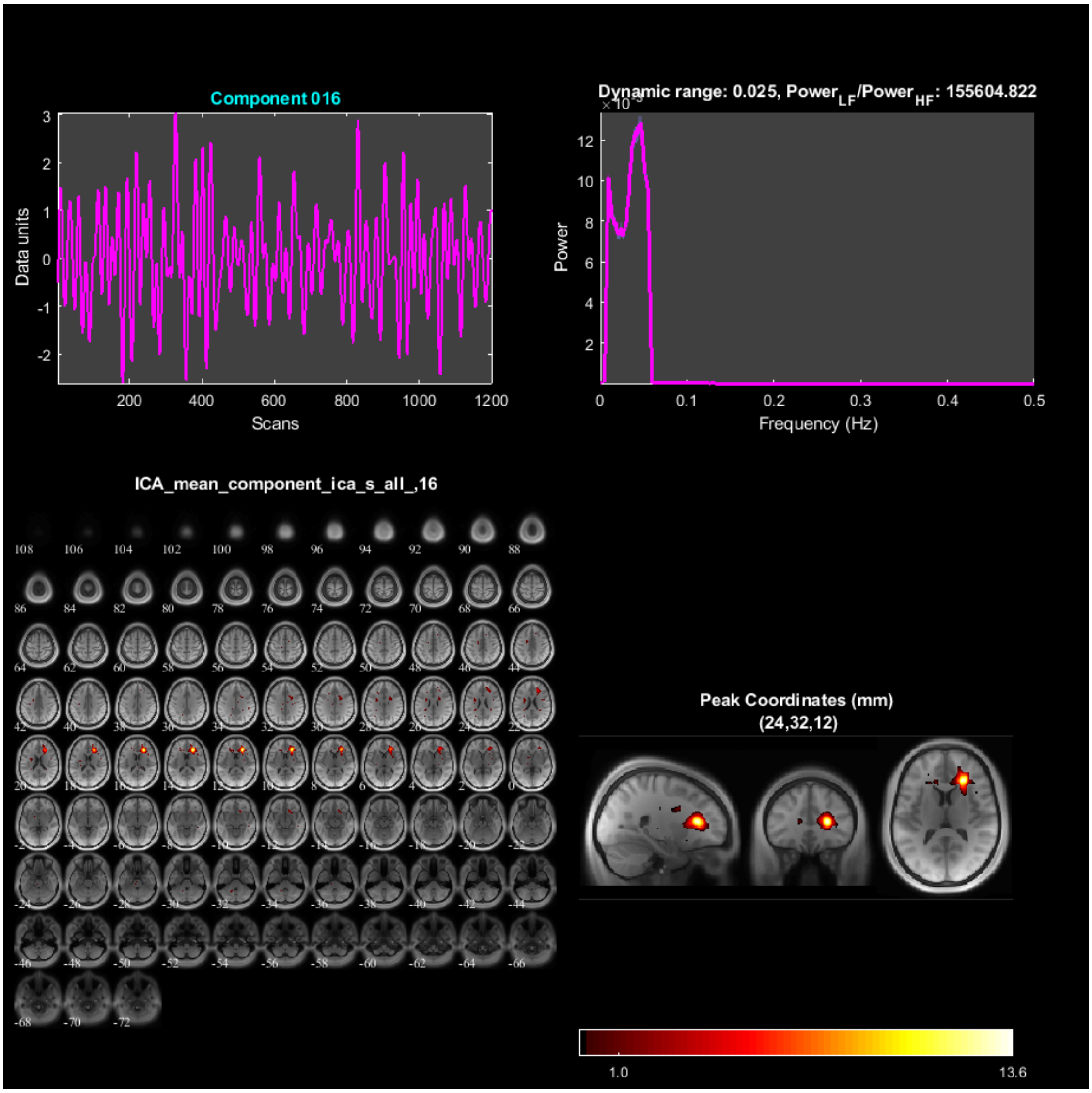
Component 016
Data units
Scans
Dynamic range: 0.025, Power$_{LF}$/Power$_{HF}$: 155604.822
Power
Frequency (Hz)
ICA_mean_component_ica_s_all_,16
Peak Coordinates (mm)
(24,32,12)
1.0
13.6

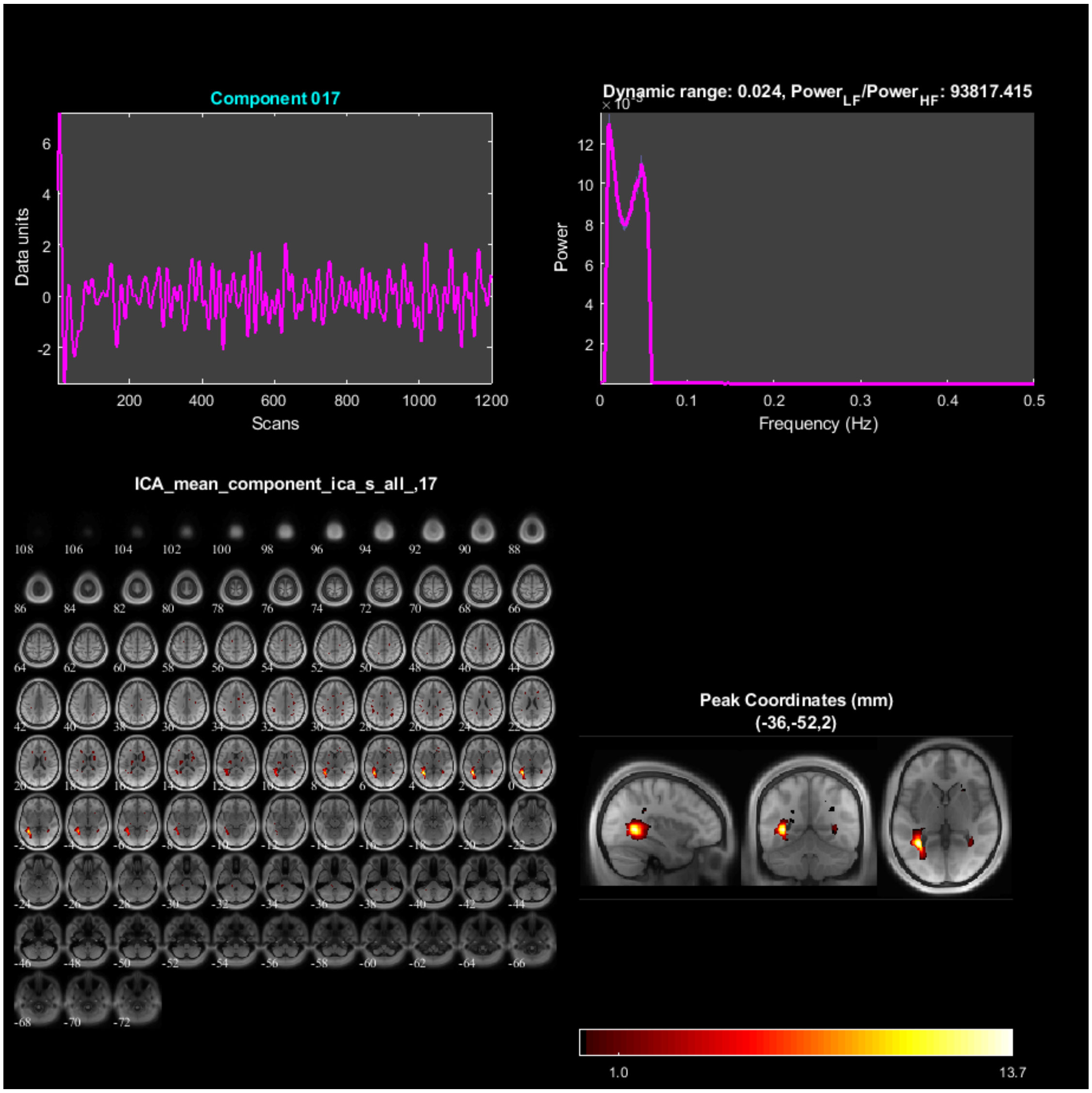

Component 017
Data units
Scans
Dynamic range: 0.024, Power$_{LF}$/Power$_{HF}$: 93817.415
Power
Frequency (Hz)
ICA_mean_component_ica_s_all_,17
Peak Coordinates (mm)
(-36,-52,2)
1.0
13.7

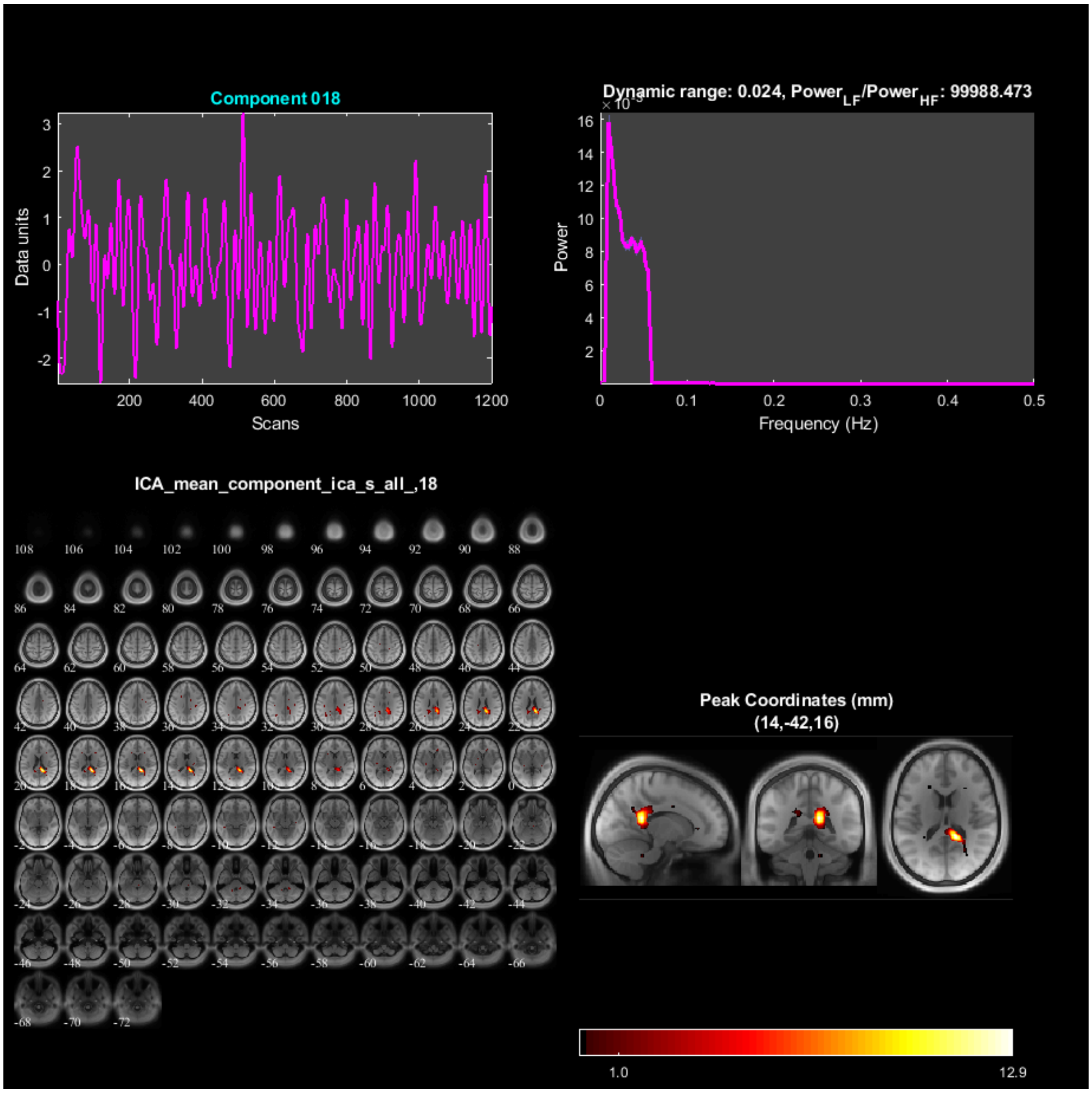
Component 018
Data units
Scans
Dynamic range: 0.024, Power_LF/Power_HF: 99988.473
Power
Frequency (Hz)
ICA_mean_component_ica_s_all_,18
Peak Coordinates (mm)
(14,-42,16)
1.0
12.9

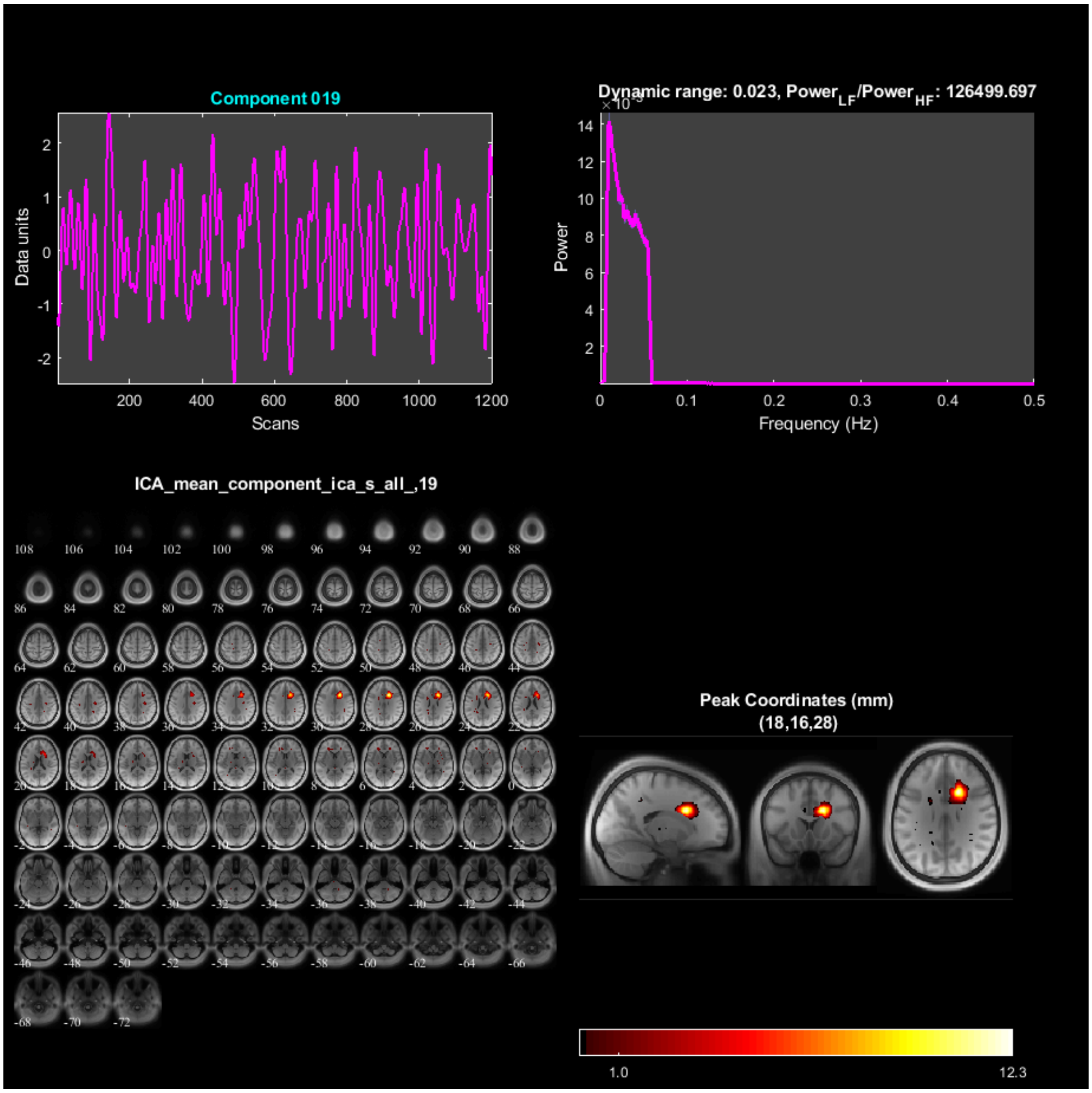

Component 019
Data units
Scans
Dynamic range: 0.023, Power$_{LF}$/Power$_{HF}$: 126499.697
Power
Frequency (Hz)
ICA_mean_component_ica_s_all_,19
Peak Coordinates (mm)
(18,16,28)
1.0
12.3

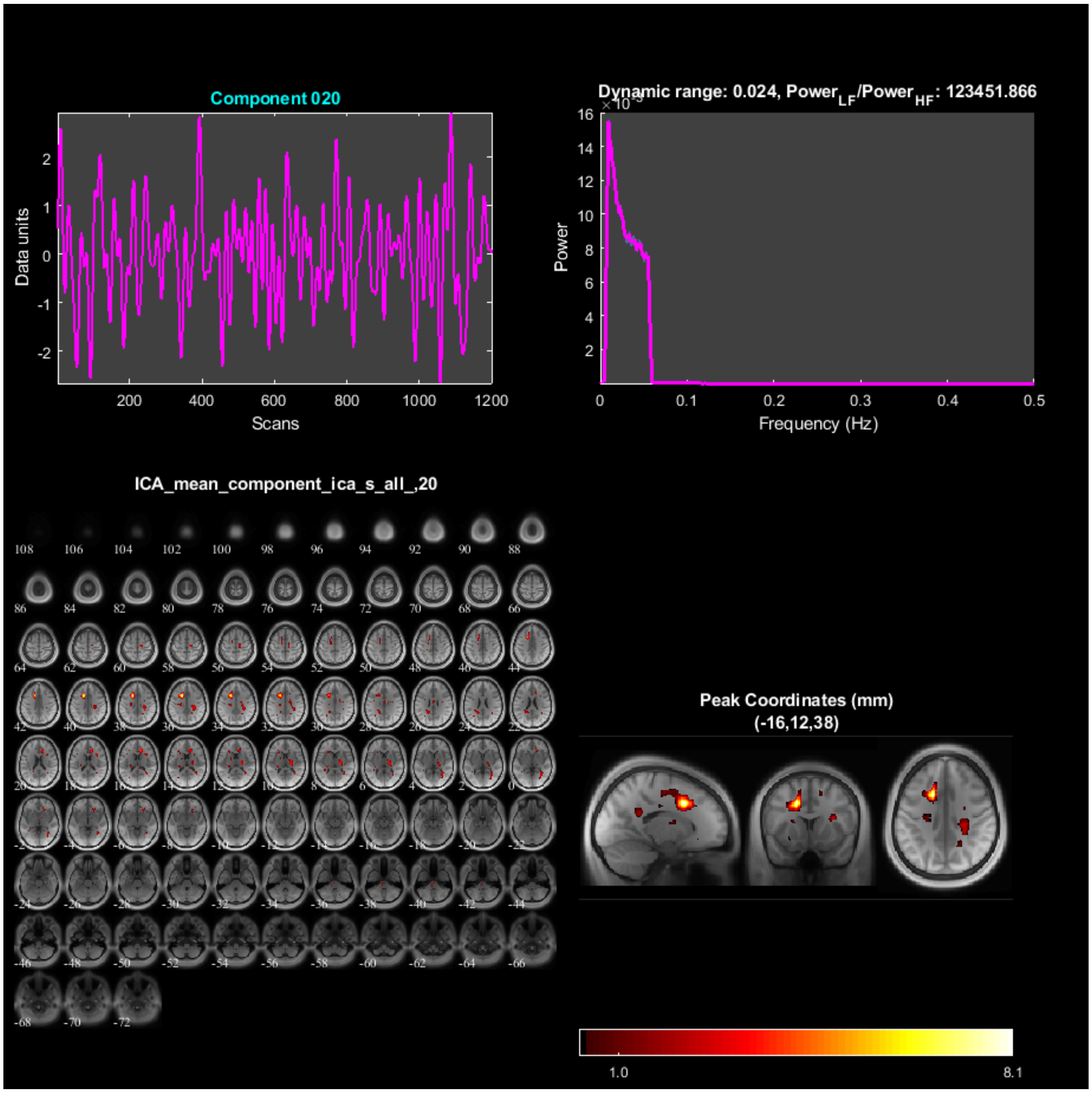

Component 020
Data units
Scans
Dynamic range: 0.024, Power$_{LF}$/Power$_{HF}$: 123451.866
Power
Frequency (Hz)
ICA_mean_component_ica_s_all_,20
Peak Coordinates (mm)
(-16,12,38)
1.0
8.1

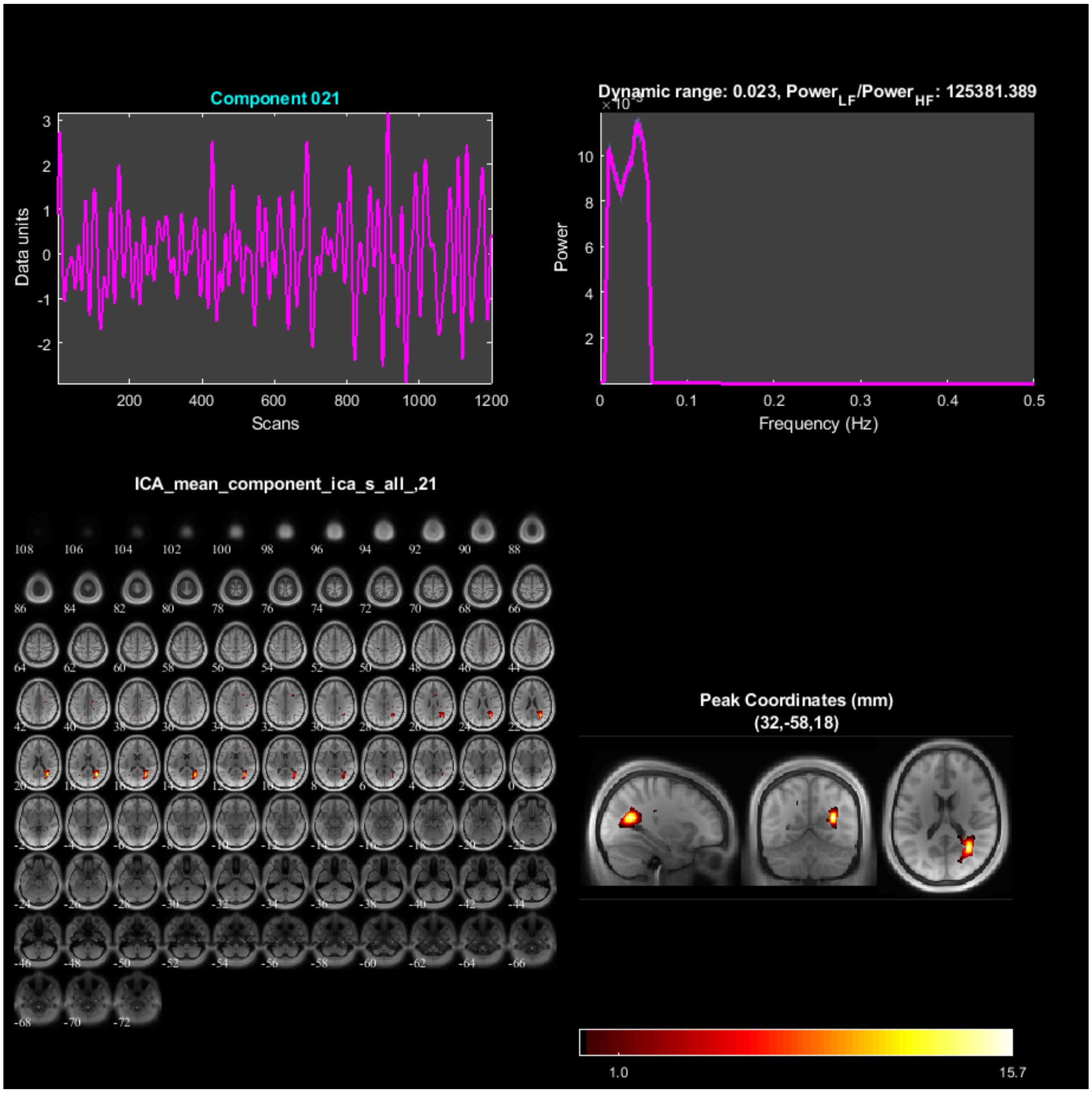

Component 021
Data units
Scans
Dynamic range: 0.023, Power$_{LF}$/Power$_{HF}$: 125381.389
Power
Frequency (Hz)
ICA_mean_component_ica_s_all_,21
Peak Coordinates (mm)
(32,-58,18)
1.0
15.7

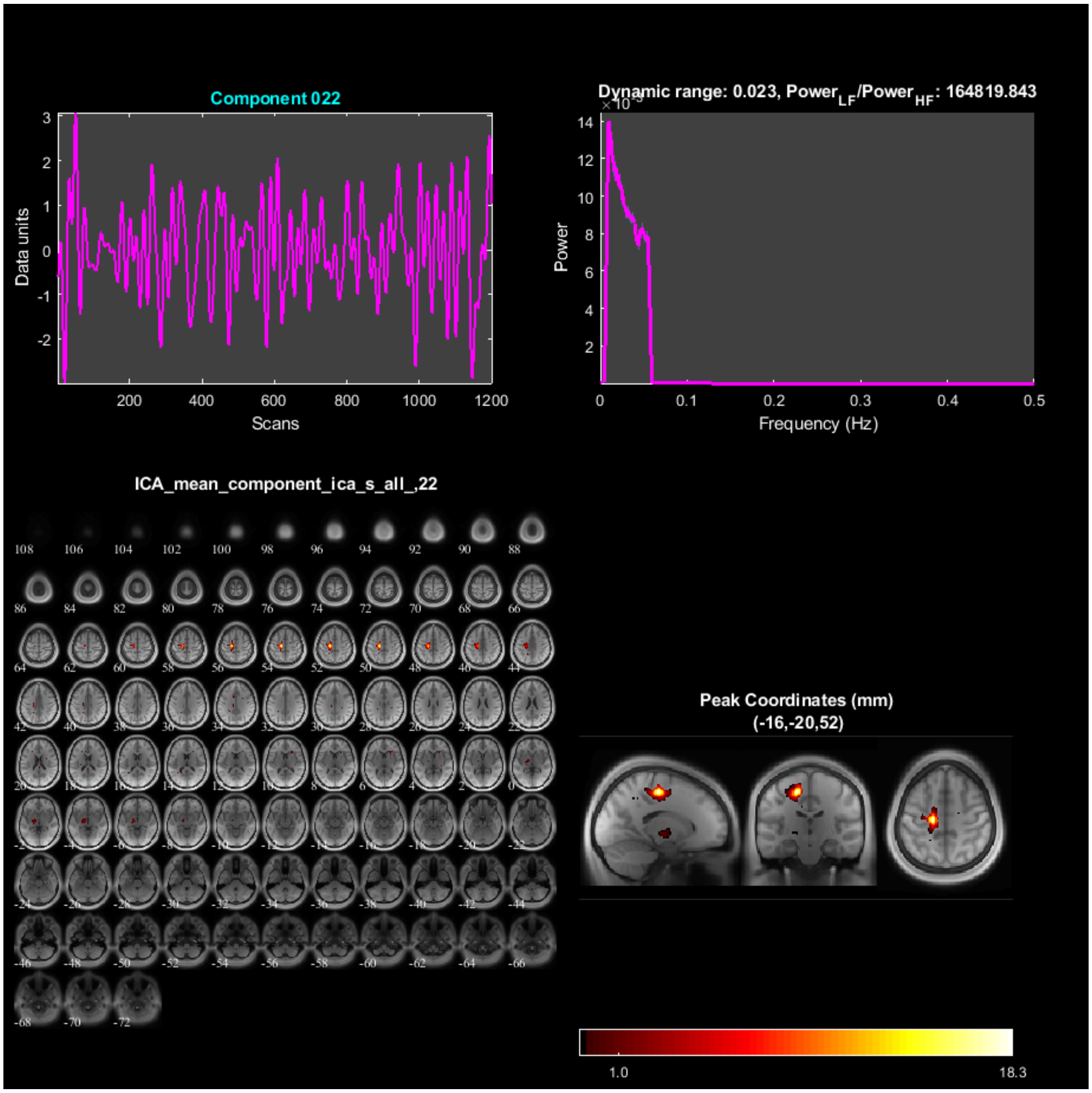
Component 022
Data units
Scans
Dynamic range: 0.023, Power$_{LF}$/Power$_{HF}$: 164819.843
Power
Frequency (Hz)
ICA_mean_component_ica_s_all_,22
Peak Coordinates (mm)
(-16,-20,52)
1.0
18.3

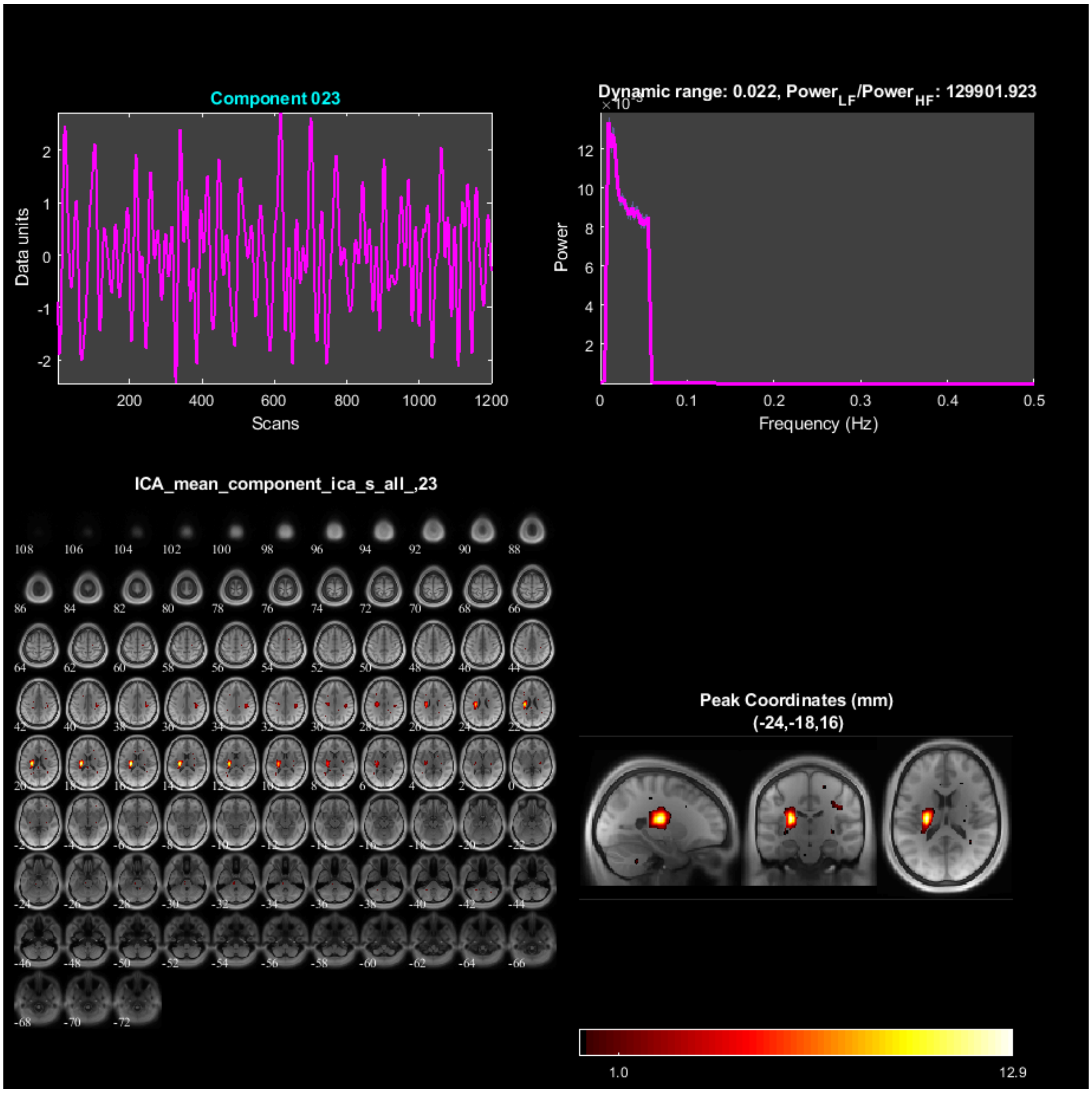

Component 023
Data units
Scans
Dynamic range: 0.022, Power$_{LF}$/Power$_{HF}$: 129901.923
Power
Frequency (Hz)
ICA_mean_component_ica_s_all_,23
Peak Coordinates (mm)
(-24,-18,16)
1.0
12.9

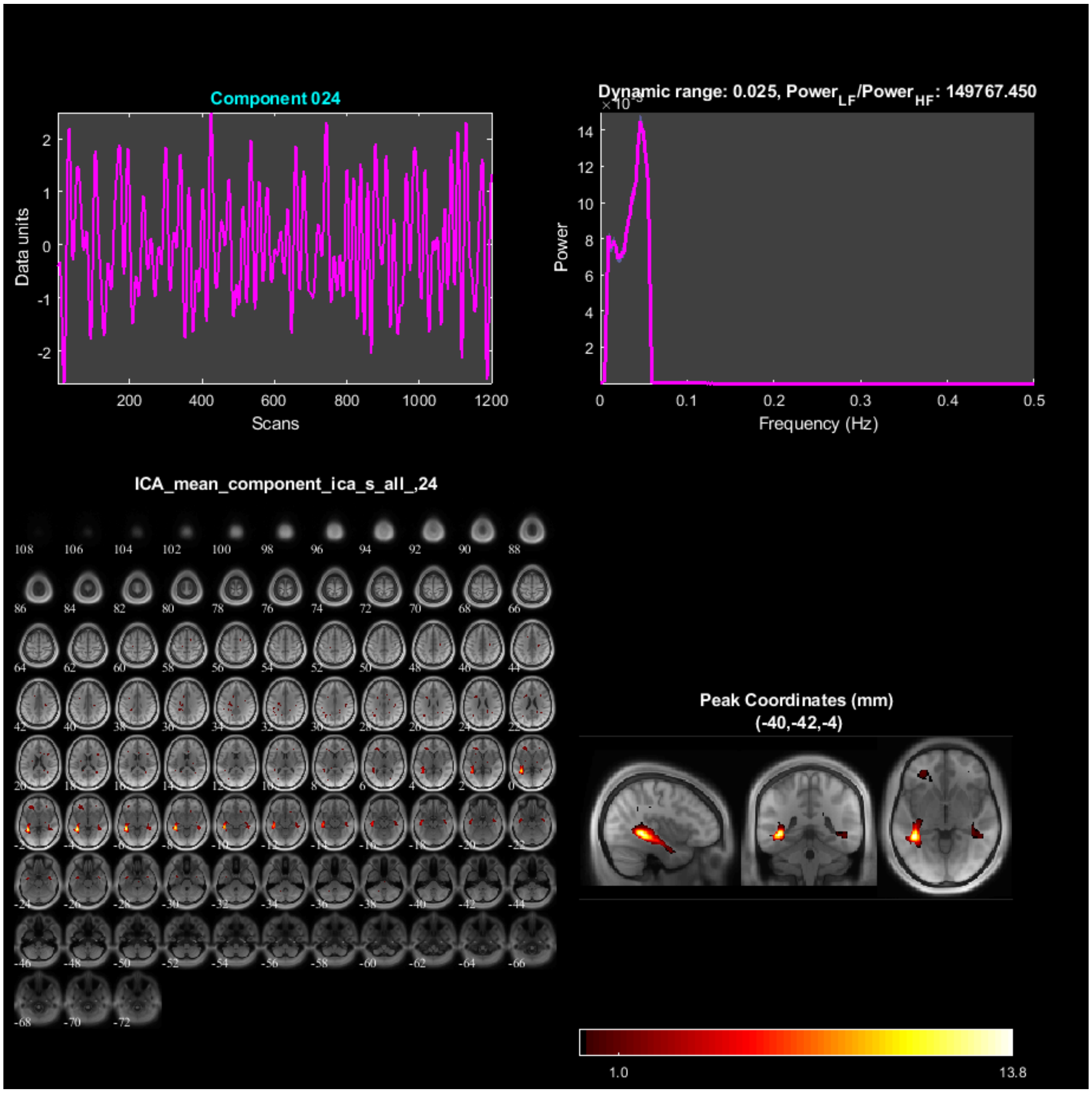

Component 024
Data units
Scans
Dynamic range: 0.025, Power$_{LF}$/Power$_{HF}$: 149767.450
Power
Frequency (Hz)
ICA_mean_component_ica_s_all_,24
Peak Coordinates (mm)
(-40,-42,-4)
1.0
13.8

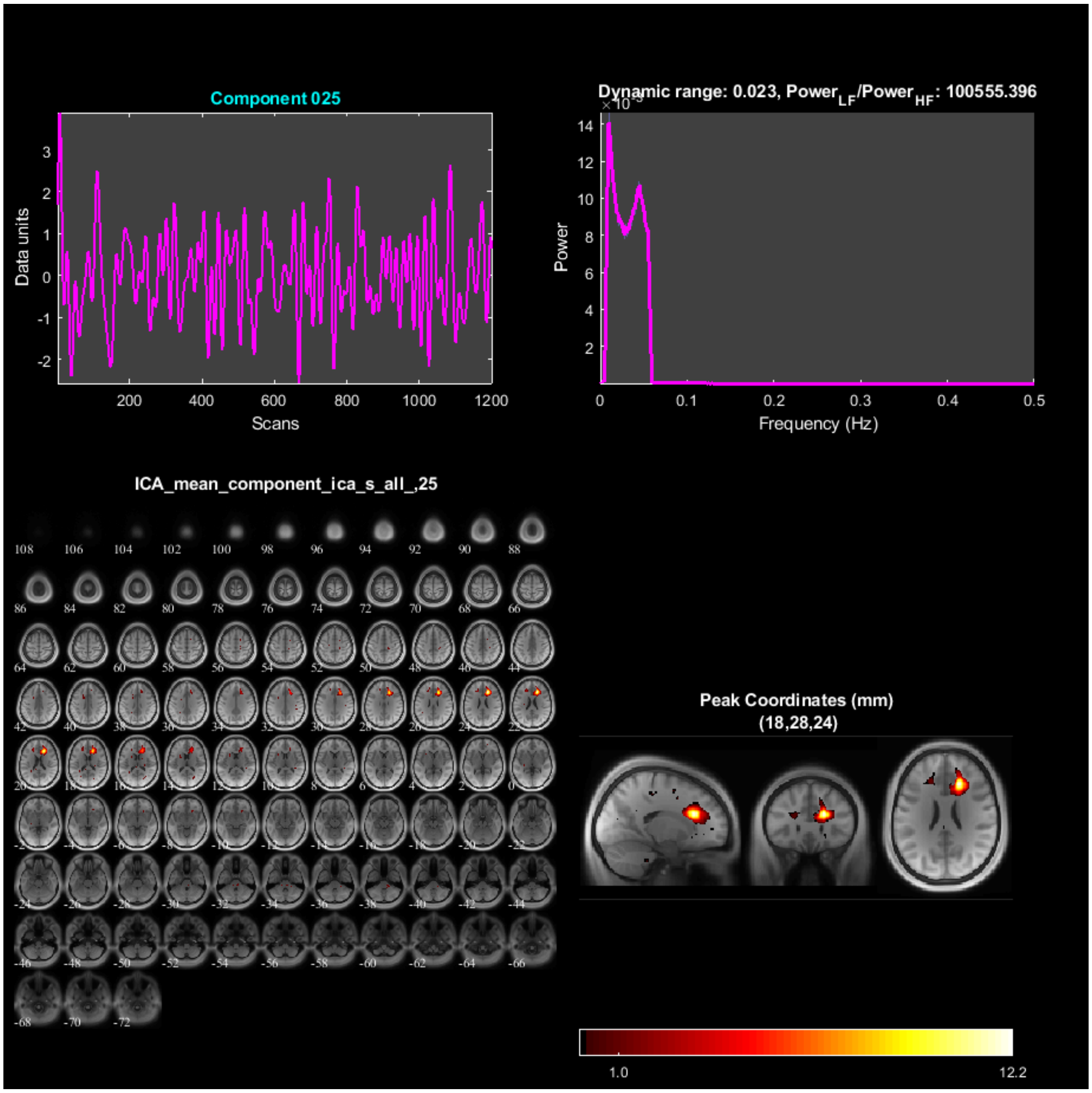
Component 025
Data units
Scans
Dynamic range: 0.023, Power$_{LF}$/Power$_{HF}$: 100555.396
Power
Frequency (Hz)
ICA_mean_component_ica_s_all_,25
Peak Coordinates (mm)
(18,28,24)
1.0
12.2

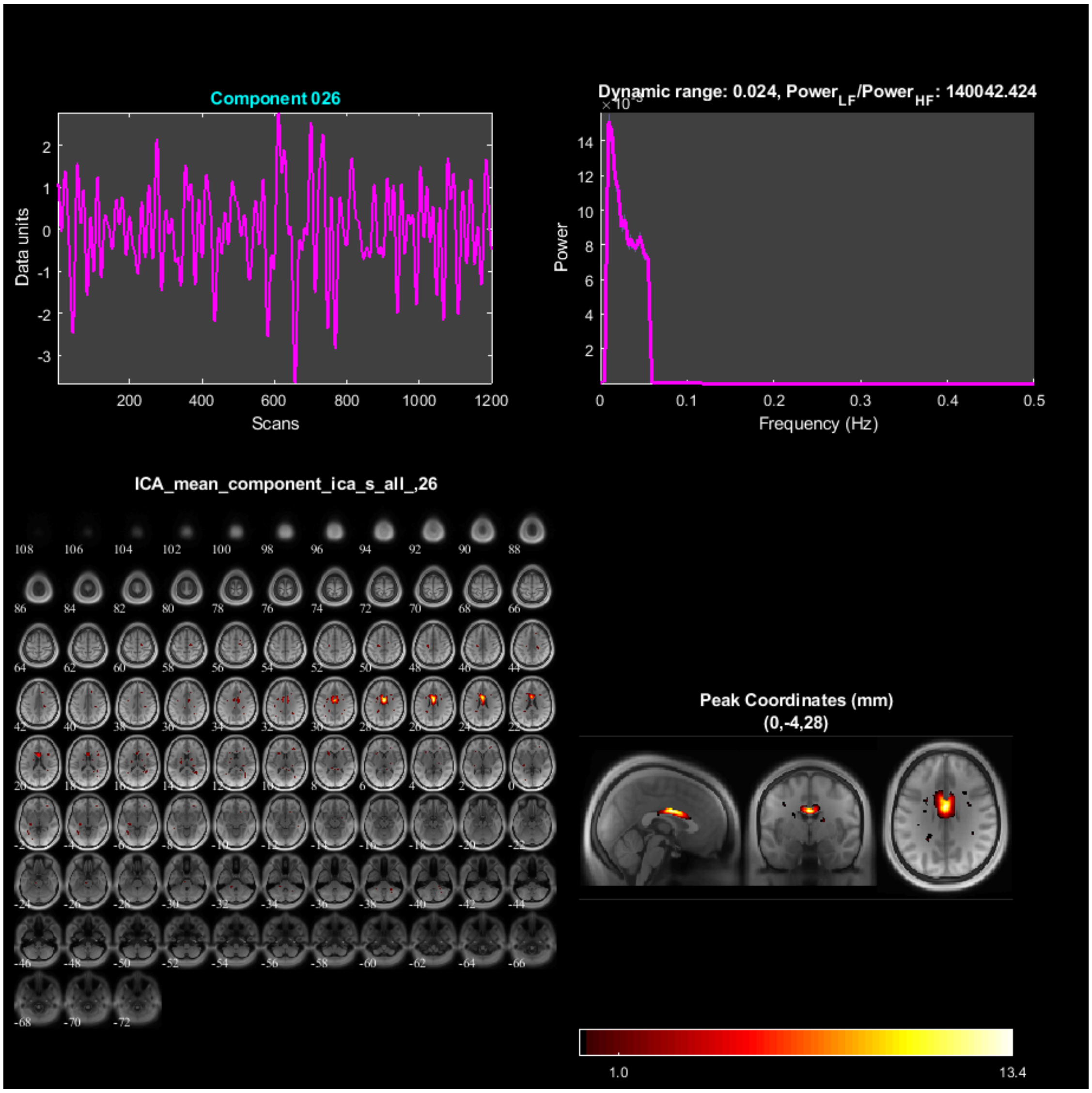

Component 026
Data units
Scans
Dynamic range: 0.024, Power$_{LF}$/Power$_{HF}$: 140042.424
Power
Frequency (Hz)
ICA_mean_component_ica_s_all_,26
Peak Coordinates (mm)
(0,-4,28)
1.0
13.4

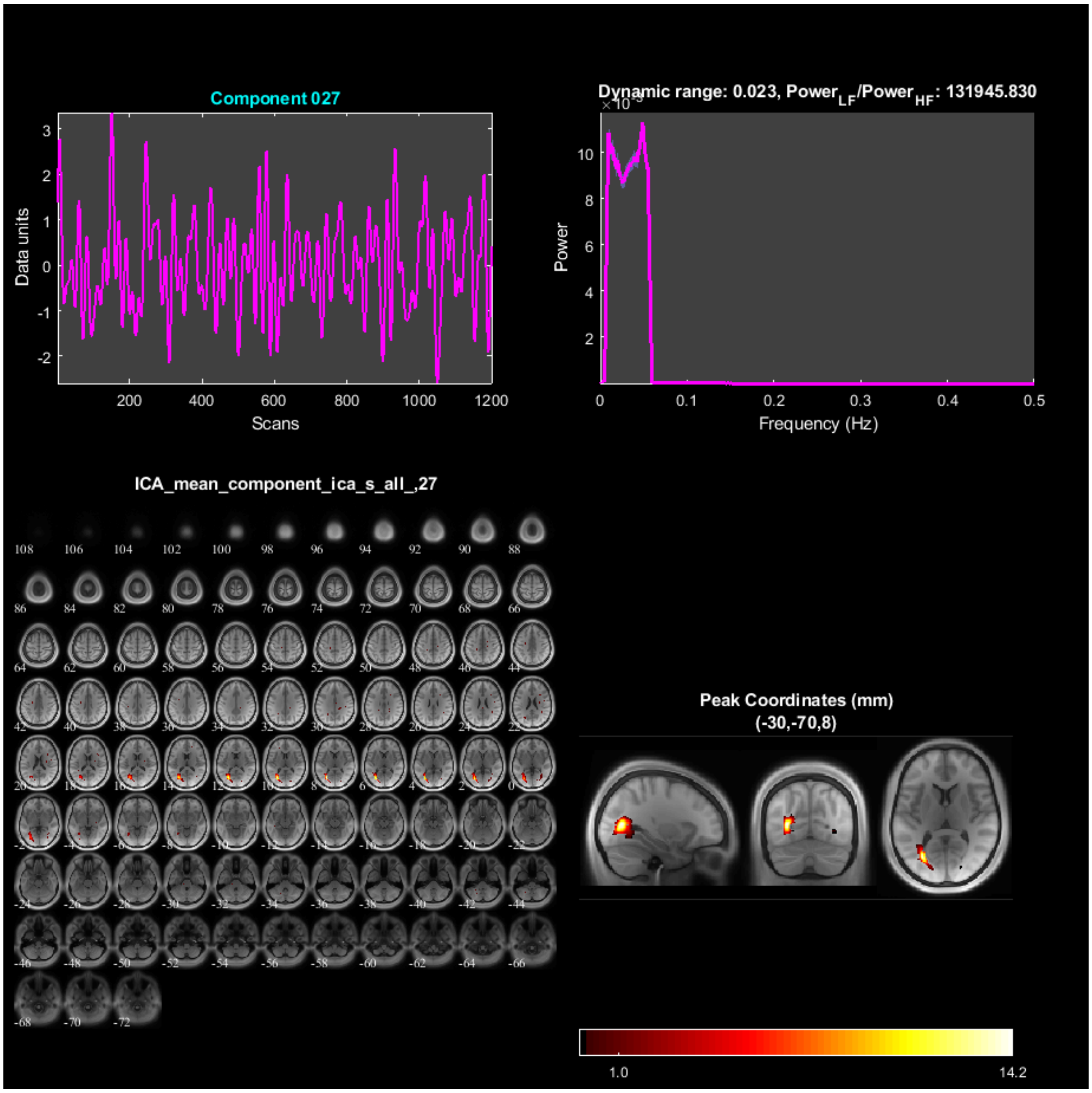

Component 027
Data units
Scans
Dynamic range: 0.023, Power$_{LF}$/Power$_{HF}$: 131945.830
Power
Frequency (Hz)
ICA_mean_component_ica_s_all_,27
Peak Coordinates (mm)
(-30,-70,8)
1.0
14.2

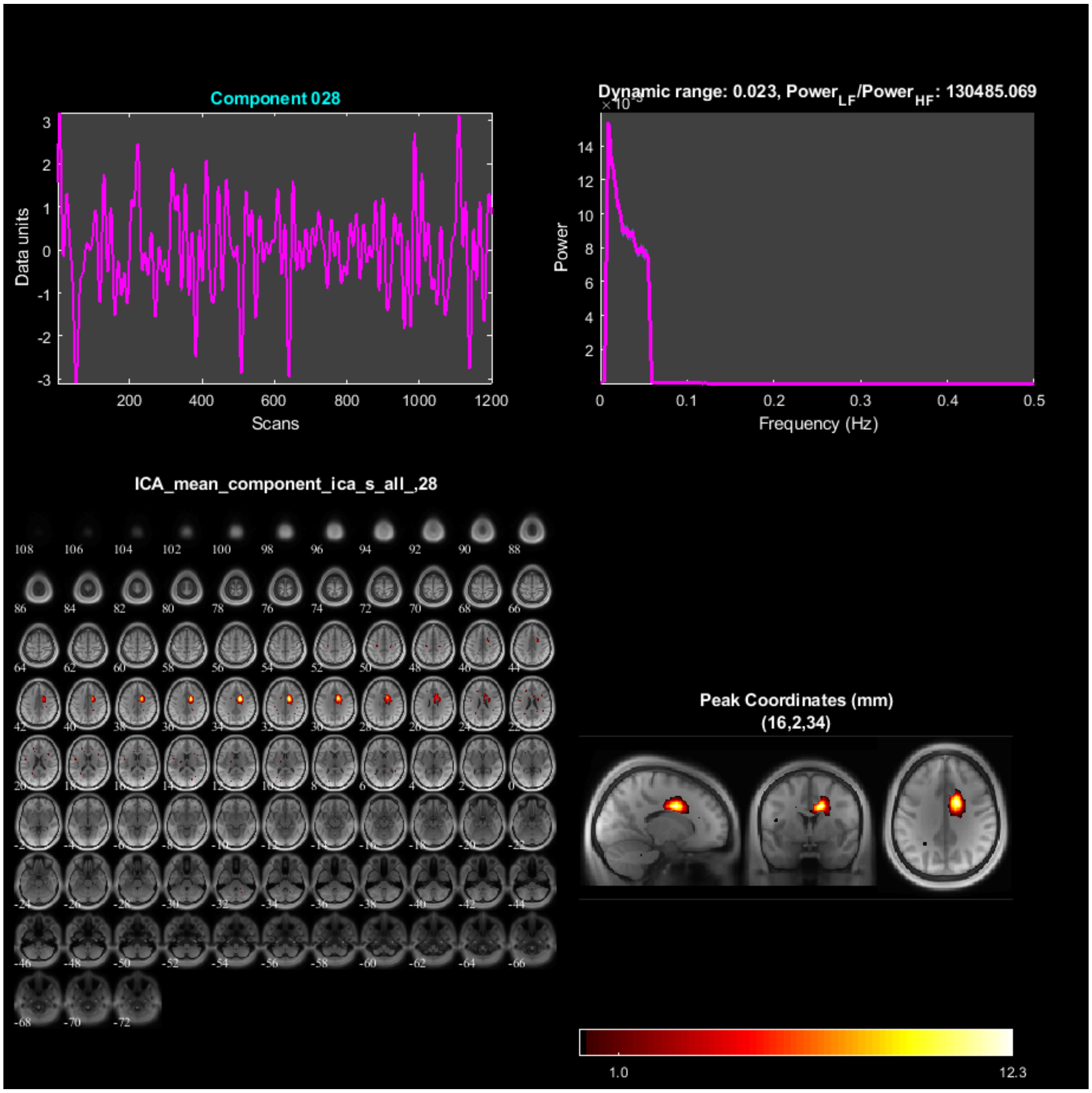

Component 028
Data units
Scans
Dynamic range: 0.023, Power$_{LF}$/Power$_{HF}$: 130485.069
Power
Frequency (Hz)
ICA_mean_component_ica_s_all_,28
Peak Coordinates (mm)
(16,2,34)
1.0
12.3

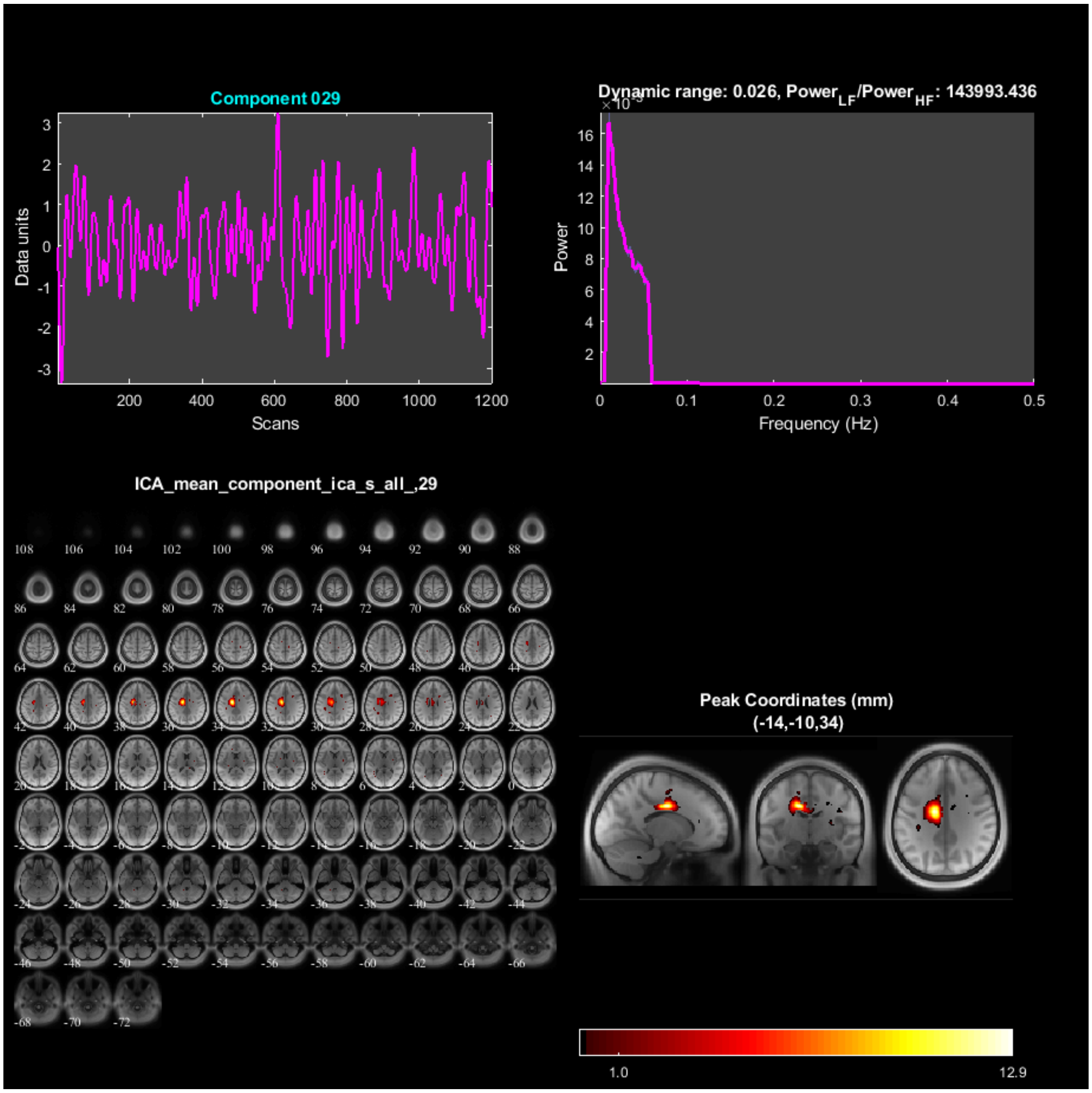

Component 029
Data units
Scans
Dynamic range: 0.026, Power$_{LF}$/Power$_{HF}$: 143993.436
Power
Frequency (Hz)
ICA_mean_component_ica_s_all_,29
Peak Coordinates (mm)
(-14,-10,34)
1.0
12.9

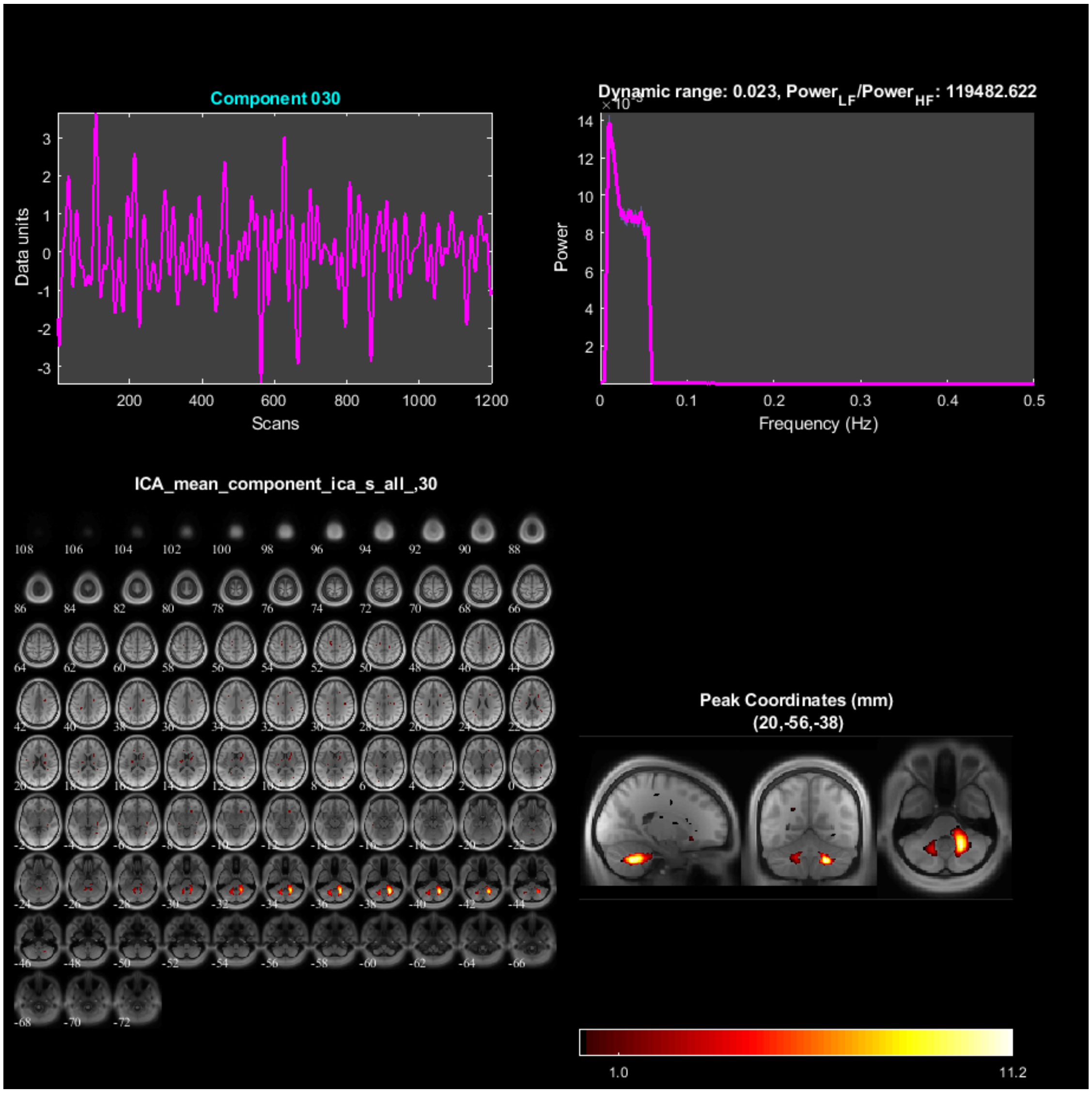
Component 030
Data units
Scans
Dynamic range: 0.023, Power$_{LF}$/Power$_{HF}$: 119482.622
Power
Frequency (Hz)
ICA_mean_component_ica_s_all_,30
Peak Coordinates (mm)
(20,-56,-38)
1.0
11.2

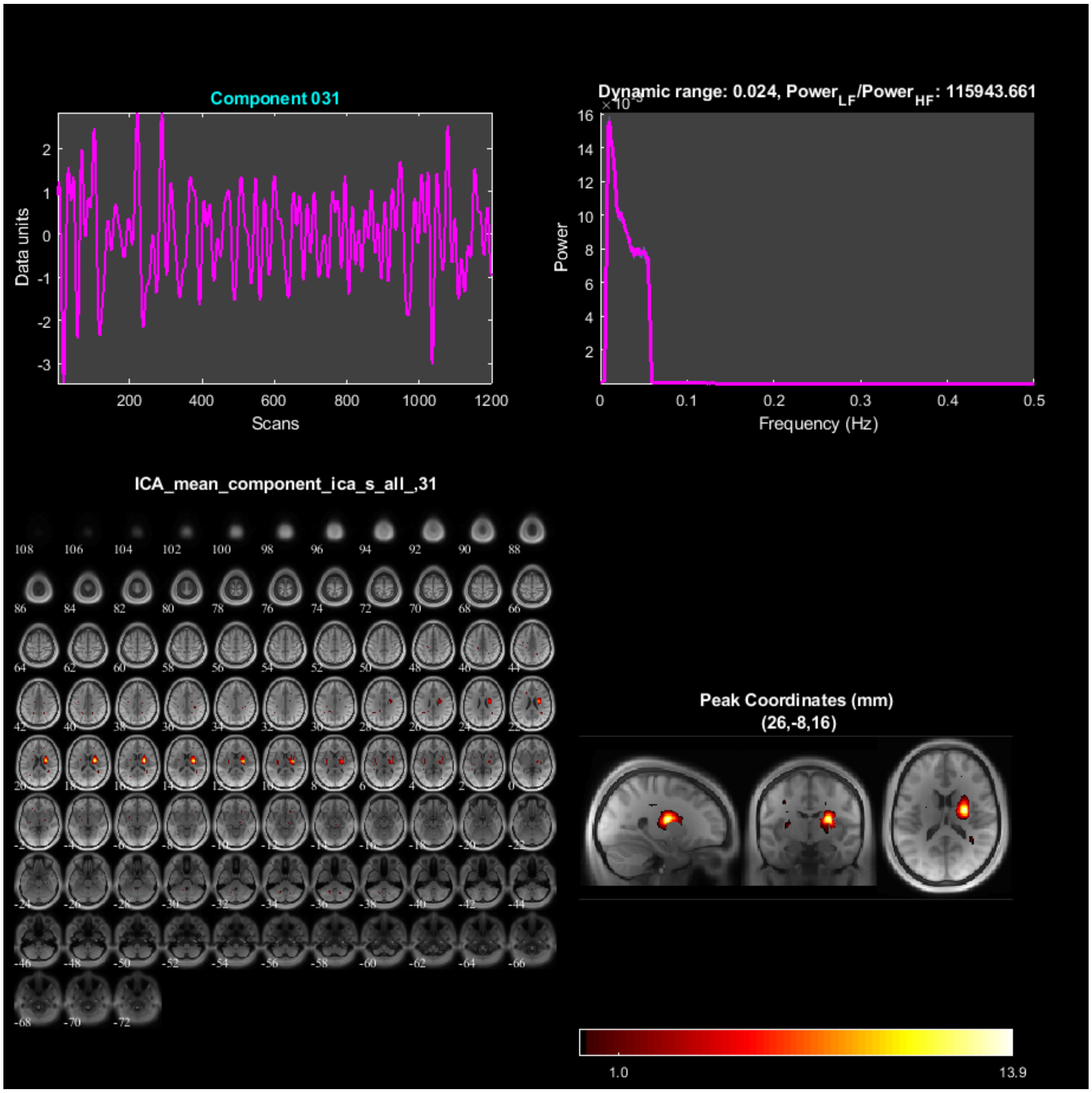

Component 031
Data units
Scans
Dynamic range: 0.024, Power$_{LF}$/Power$_{HF}$: 115943.661
Power
Frequency (Hz)
ICA_mean_component_ica_s_all_,31
Peak Coordinates (mm)
(26,-8,16)
1.0
13.9

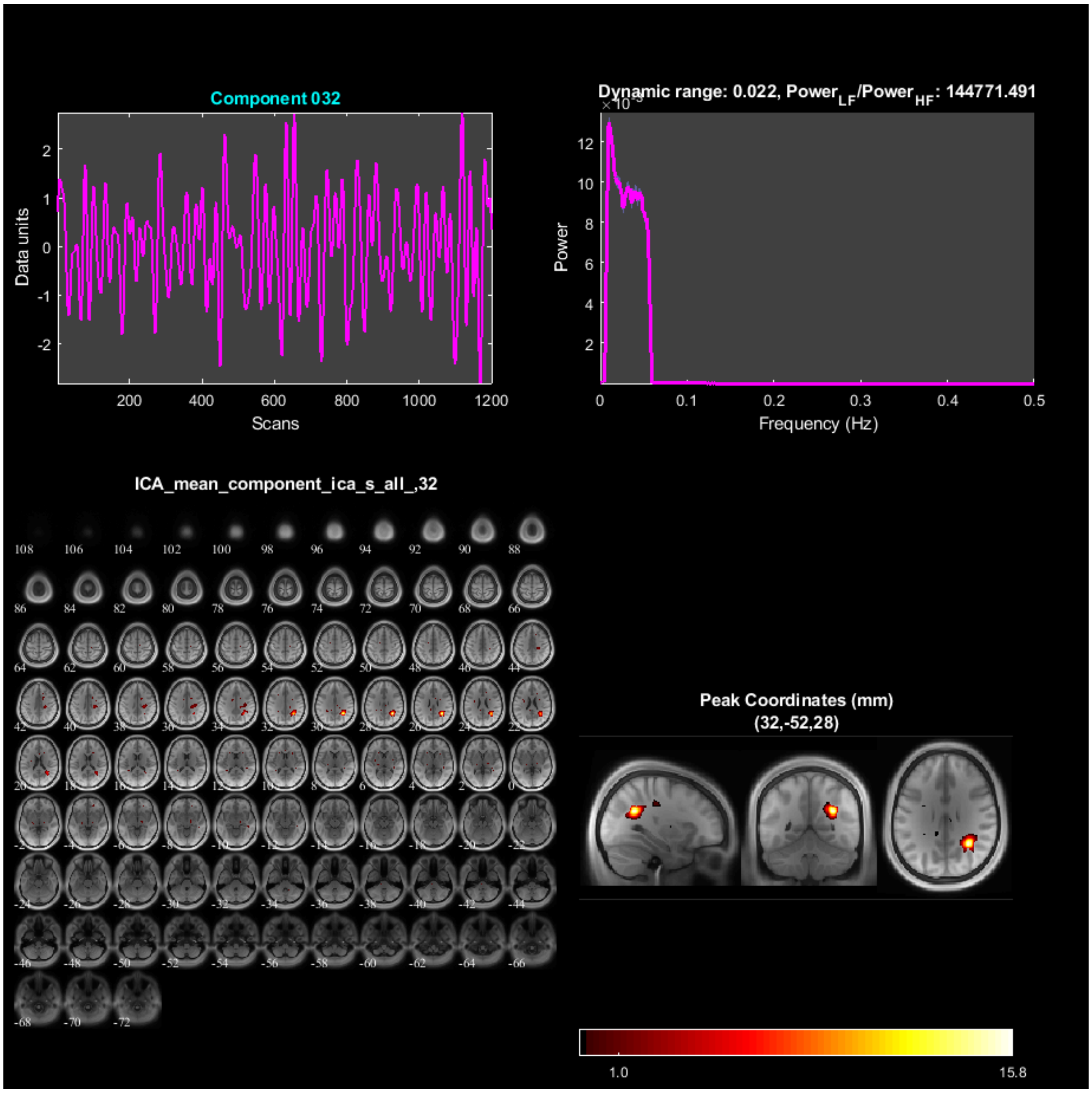

Component 032
Data units
Scans
Dynamic range: 0.022, Power_LF/Power_HF: 144771.491
Power
Frequency (Hz)
ICA_mean_component_ica_s_all_,32
Peak Coordinates (mm)
(32,-52,28)
1.0
15.8

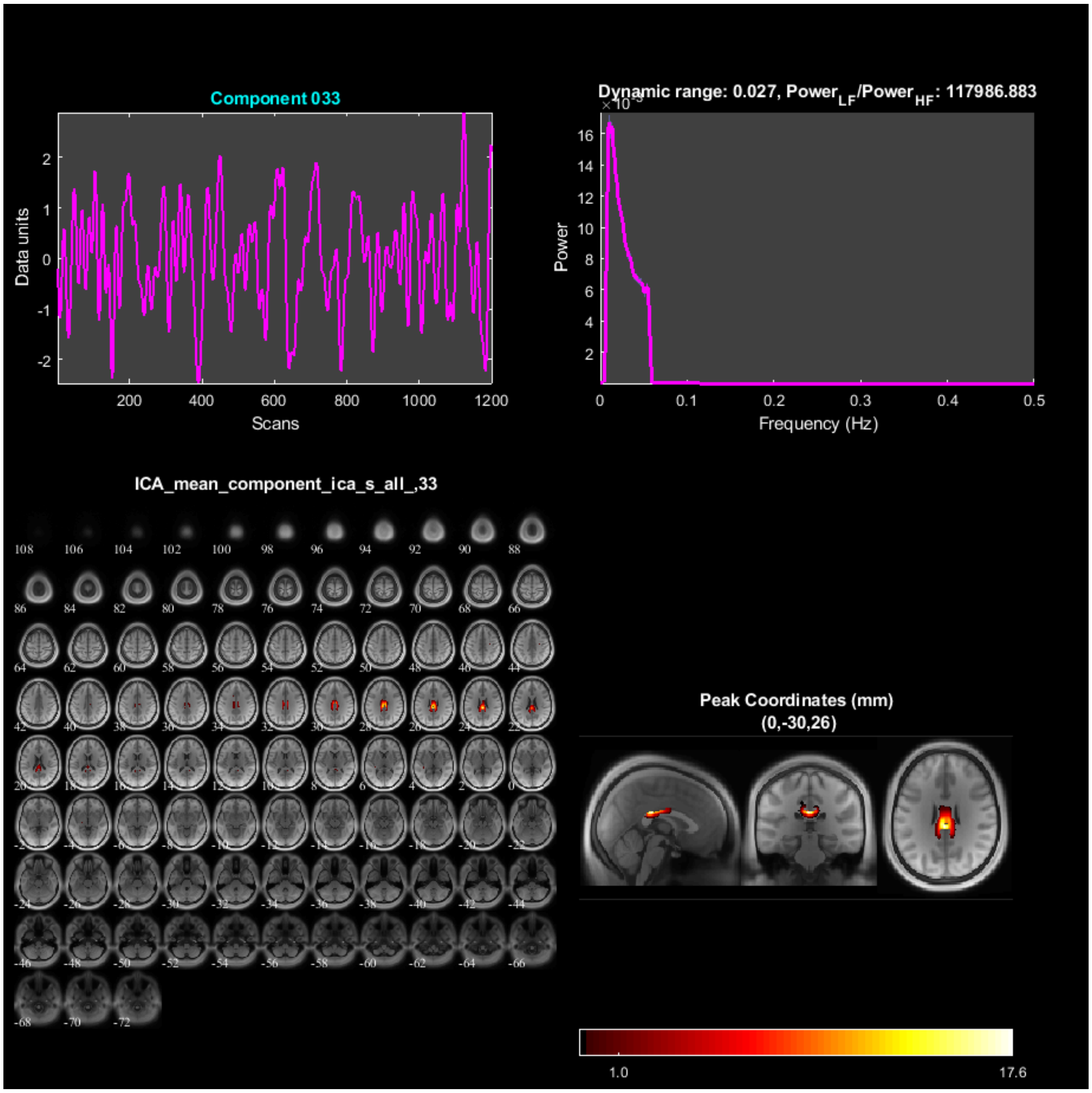
Component 033
Data units
Scans
Dynamic range: 0.027, Power_LF/Power_HF: 117986.883
Power
Frequency (Hz)
ICA_mean_component_ica_s_all_,33
Peak Coordinates (mm)
(0,-30,26)
1.0
17.6

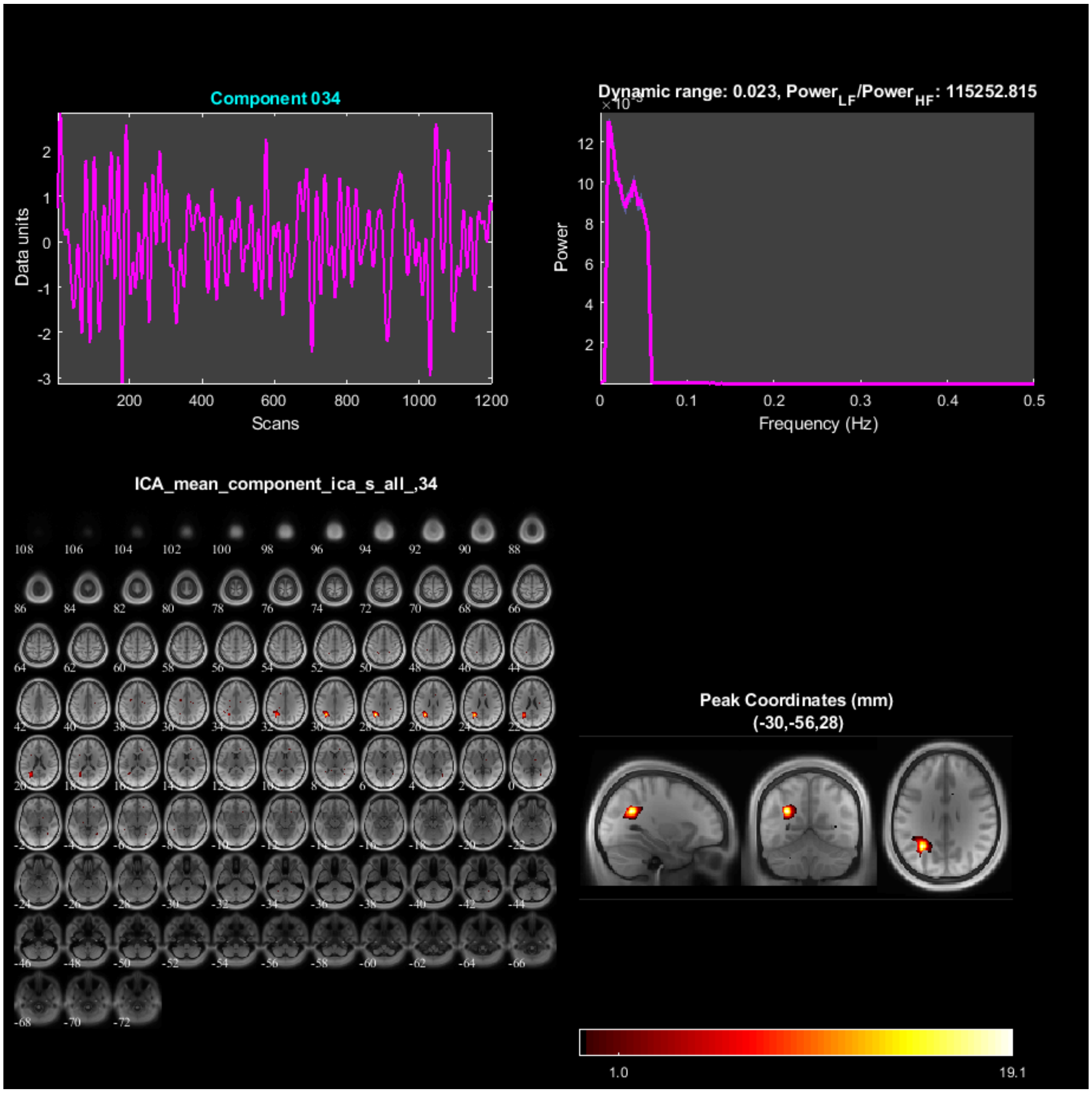
Component 034
Data units
Scans
Dynamic range: 0.023, Power_LF/Power_HF: 115252.815
Power
Frequency (Hz)
ICA_mean_component_ica_s_all_,34
Peak Coordinates (mm)
(-30,-56,28)
1.0
19.1

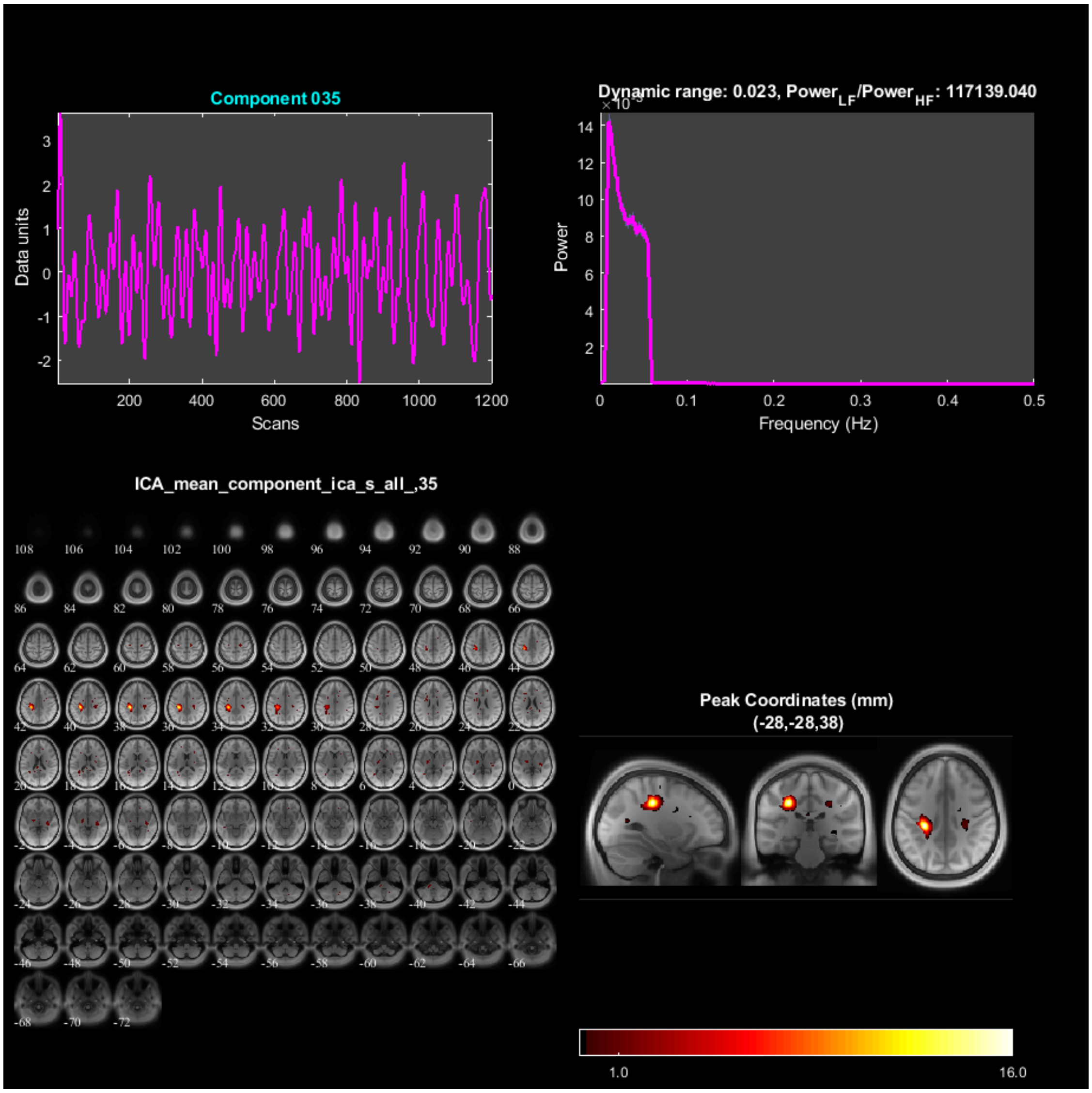
Component 035
Data units
Scans
Dynamic range: 0.023, Power_LF/Power_HF: 117139.040
Power
Frequency (Hz)
ICA_mean_component_ica_s_all_,35
Peak Coordinates (mm)
(-28,-28,38)
1.0
16.0

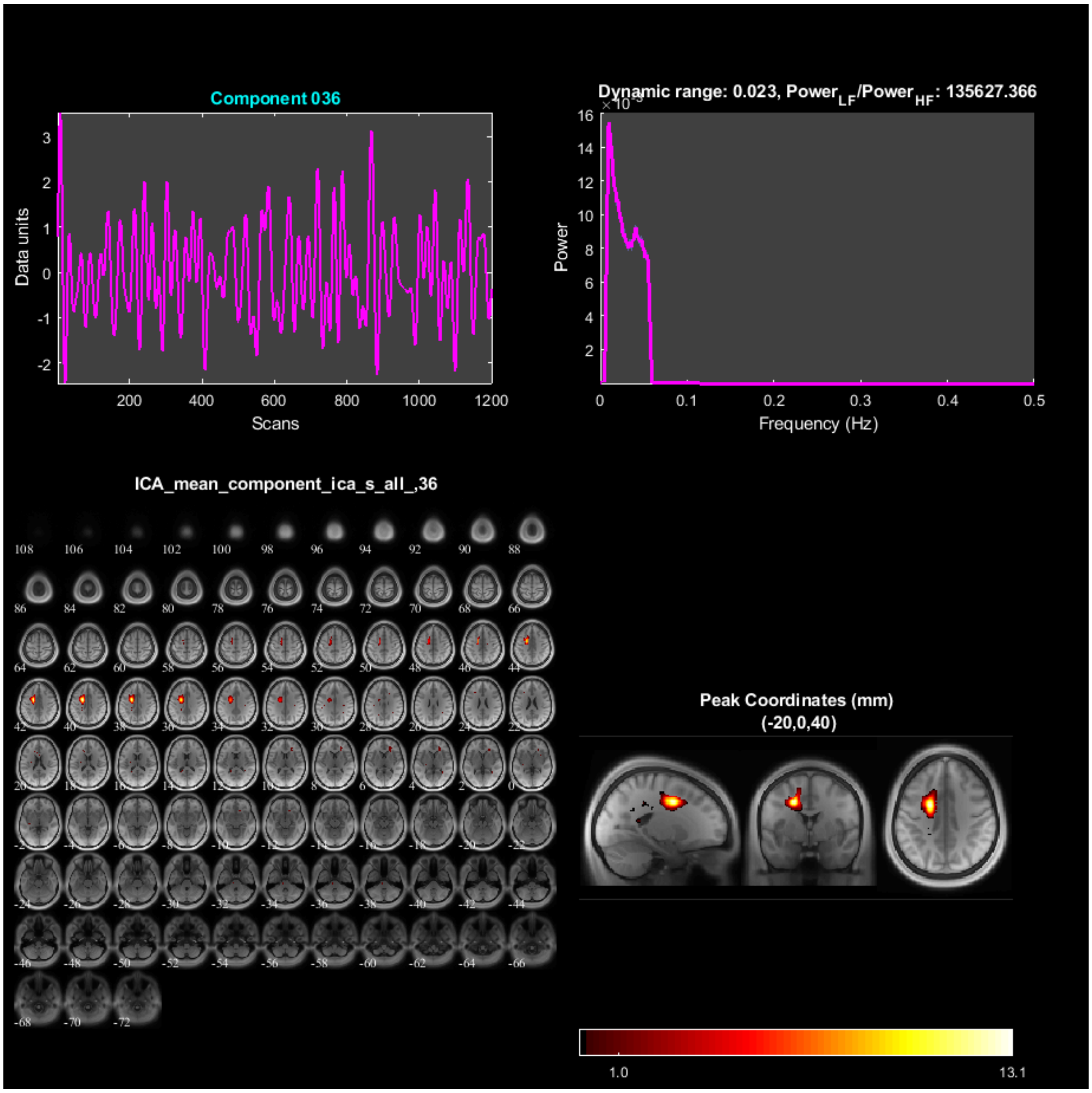

Component 036
Data units
Scans
Dynamic range: 0.023, Power$_{LF}$/Power$_{HF}$: 135627.366
Power
Frequency (Hz)
ICA_mean_component_ica_s_all_,36
Peak Coordinates (mm)
(-20,0,40)
1.0
13.1

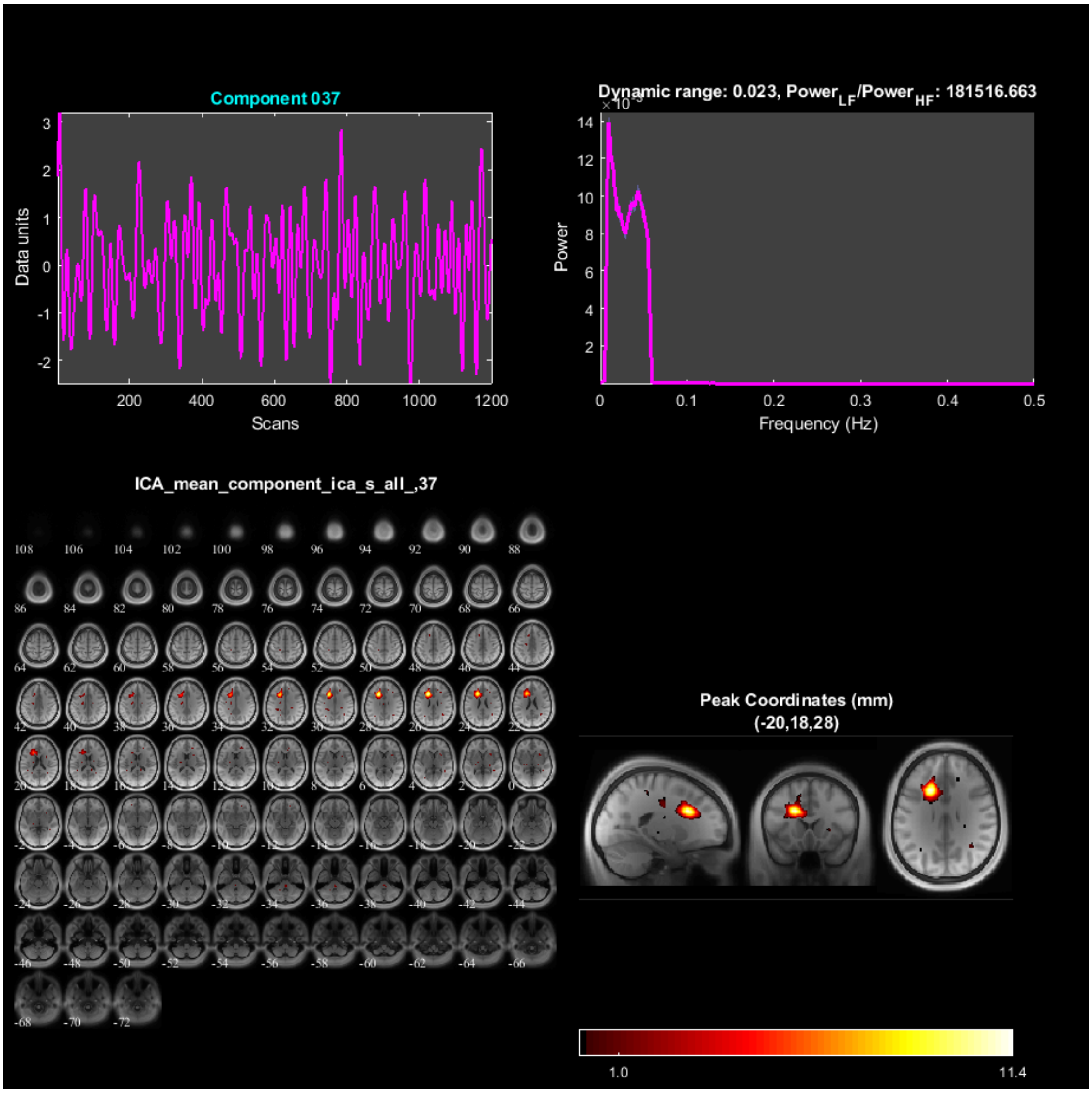

Component 037
Data units
Scans
Dynamic range: 0.023, Power$_{LF}$/Power$_{HF}$: 181516.663
Power
Frequency (Hz)
ICA_mean_component_ica_s_all_,37
Peak Coordinates (mm)
(-20,18,28)
1.0
11.4

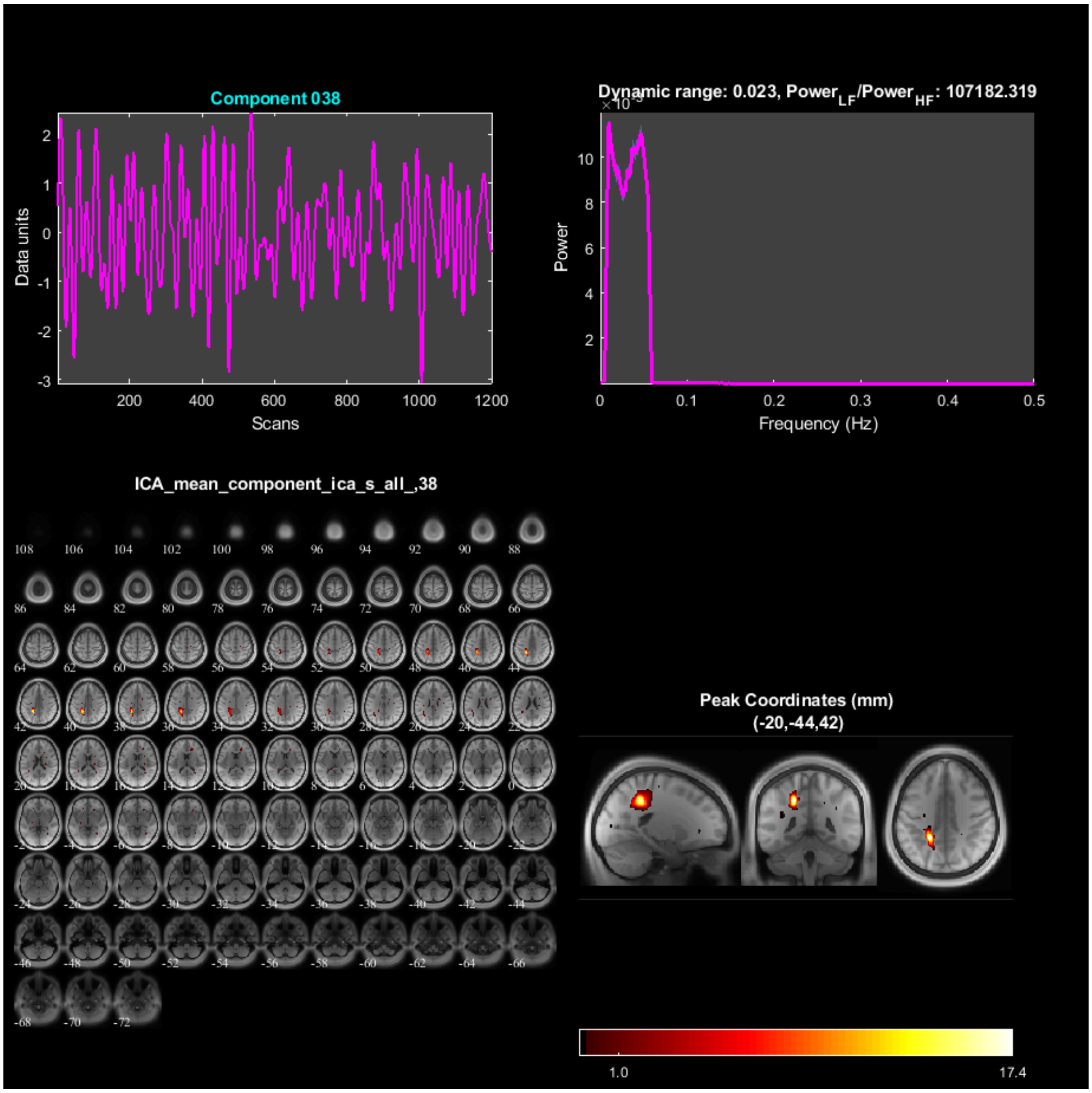
Component 038
Data units
Scans
Dynamic range: 0.023, Power LF/Power HF: 107182.319
Power
Frequency (Hz)
ICA_mean_component_ica_s_all_,38
Peak Coordinates (mm)
(-20,-44,42)
1.0
17.4

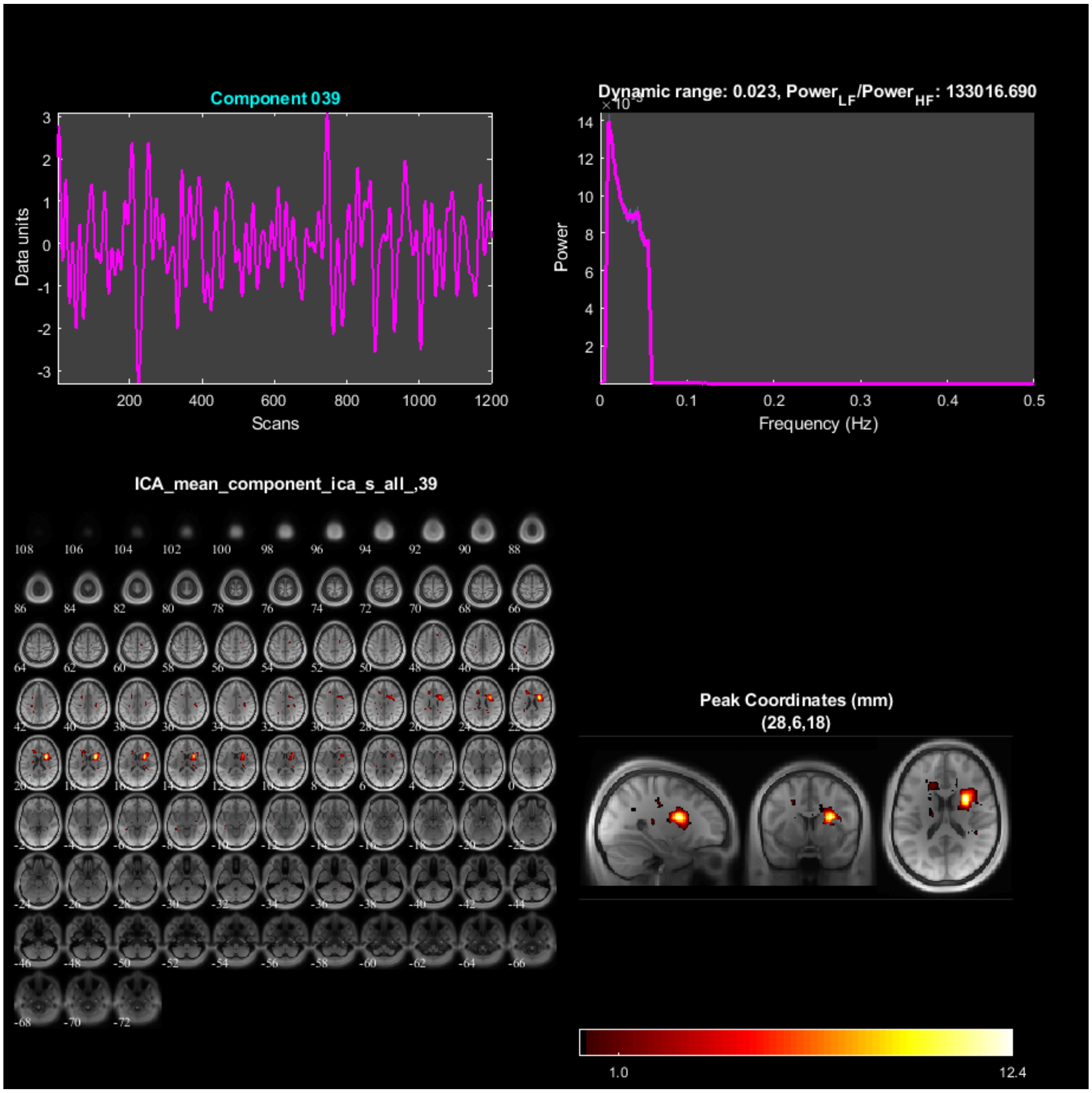

Component 039
Data units
Scans
Dynamic range: 0.023, Power$_{LF}$/Power$_{HF}$: 133016.690
Power
Frequency (Hz)
ICA_mean_component_ica_s_all_,39
Peak Coordinates (mm)
(28,6,18)
1.0
12.4

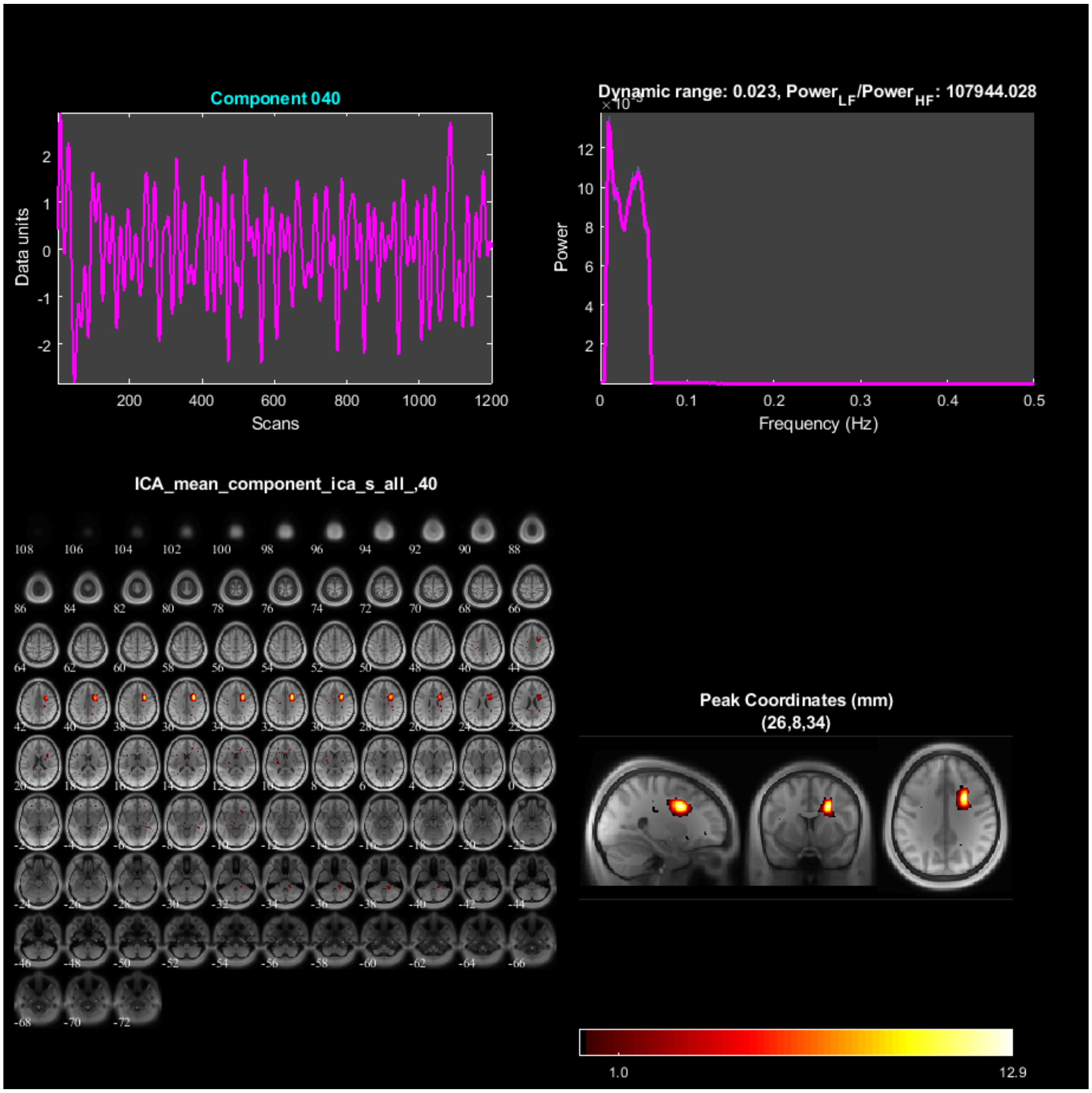

Component 040
Data units
Scans
Dynamic range: 0.023, Power_LF/Power_HF: 107944.028
Power
Frequency (Hz)
ICA_mean_component_ica_s_all_,40
Peak Coordinates (mm)
(26,8,34)
1.0
12.9

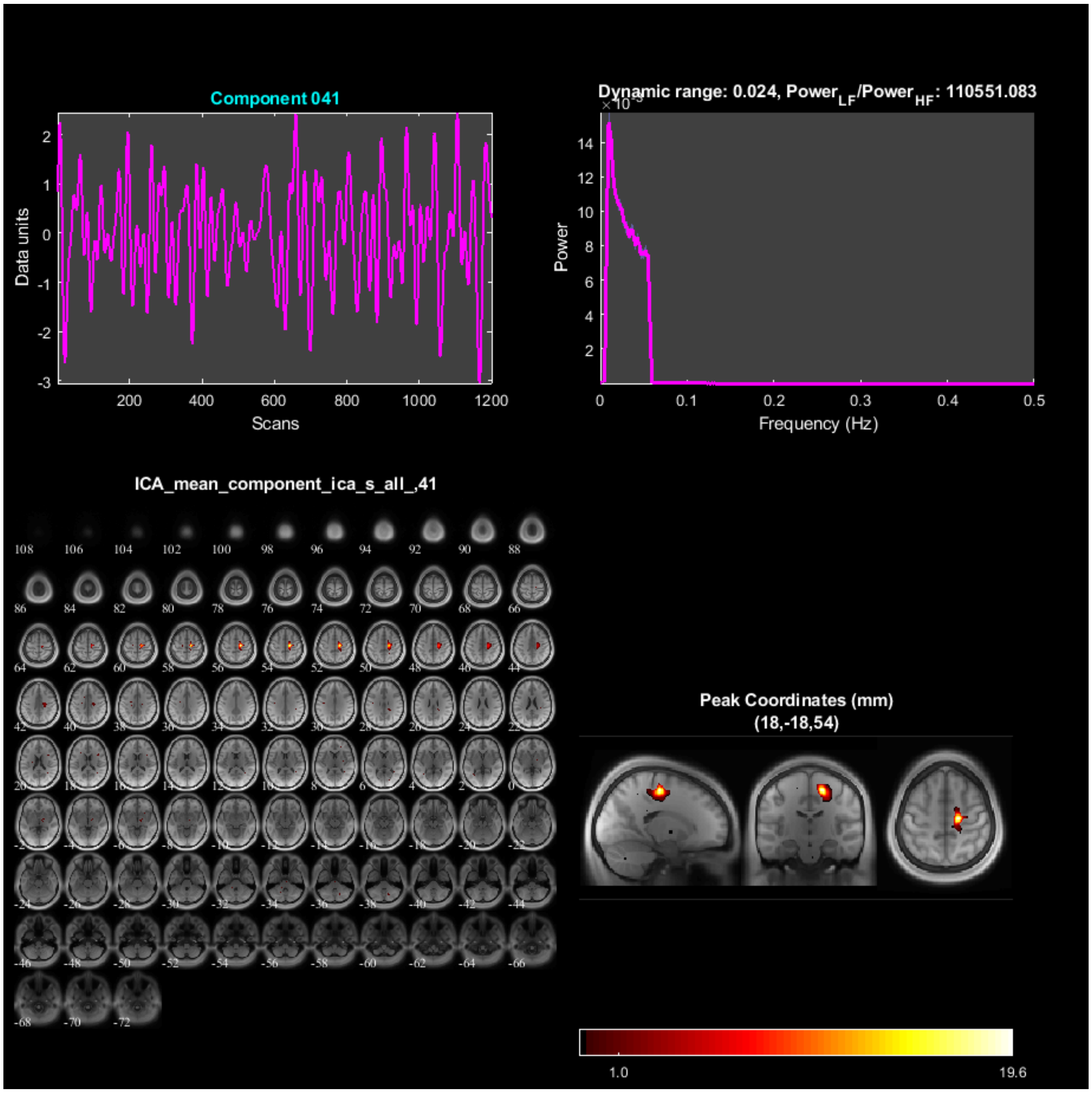
Component 041
Data units
Scans
Dynamic range: 0.024, Power_LF/Power_HF: 110551.083
Power
Frequency (Hz)
ICA_mean_component_ica_s_all_,41
Peak Coordinates (mm)
(18,-18,54)
1.0
19.6

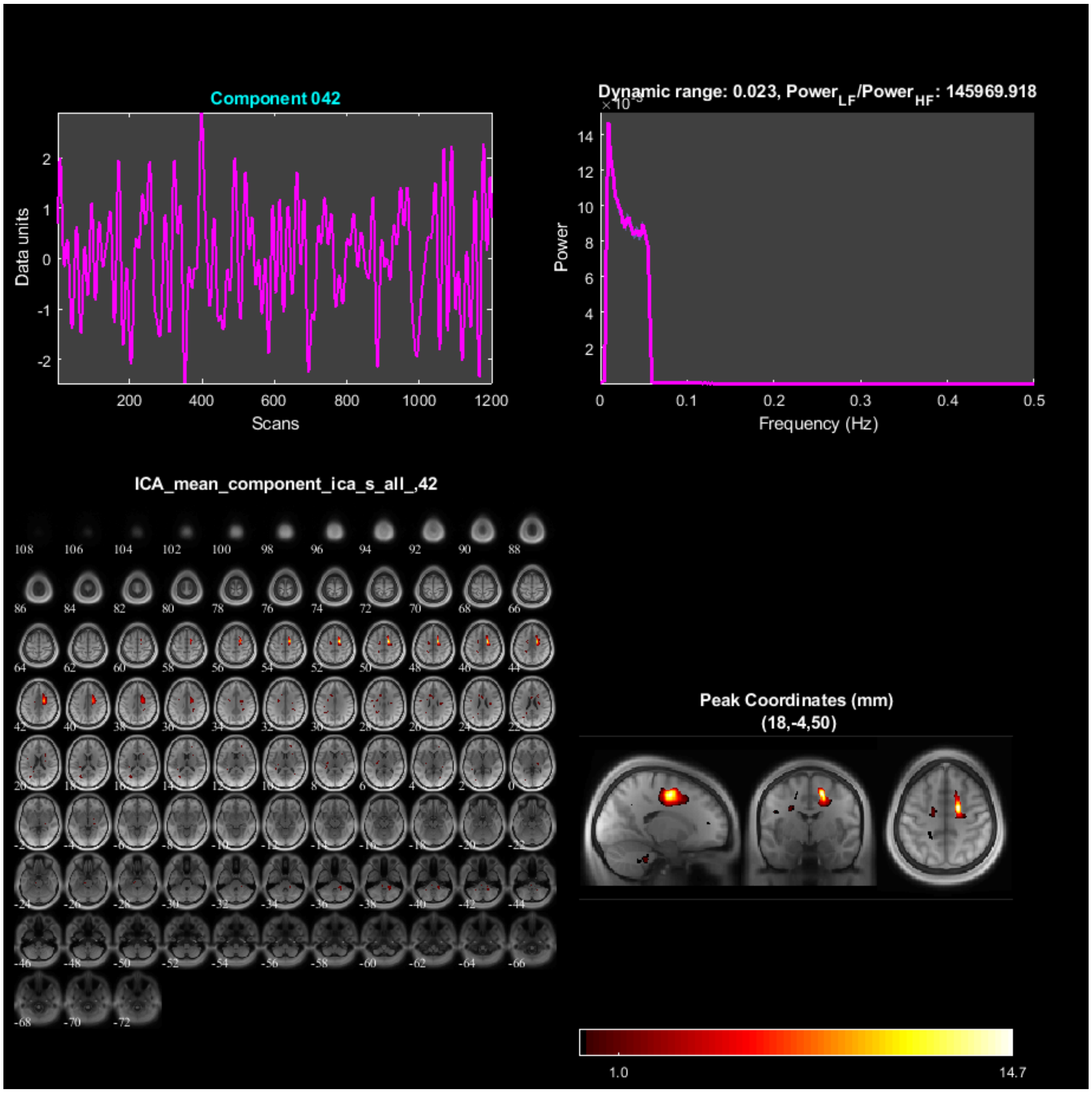

Component 042
Data units
Scans
Dynamic range: 0.023, Power$_{LF}$/Power$_{HF}$: 145969.918
Power
Frequency (Hz)
ICA_mean_component_ica_s_all_,42
Peak Coordinates (mm)
(18,-4,50)
1.0
14.7

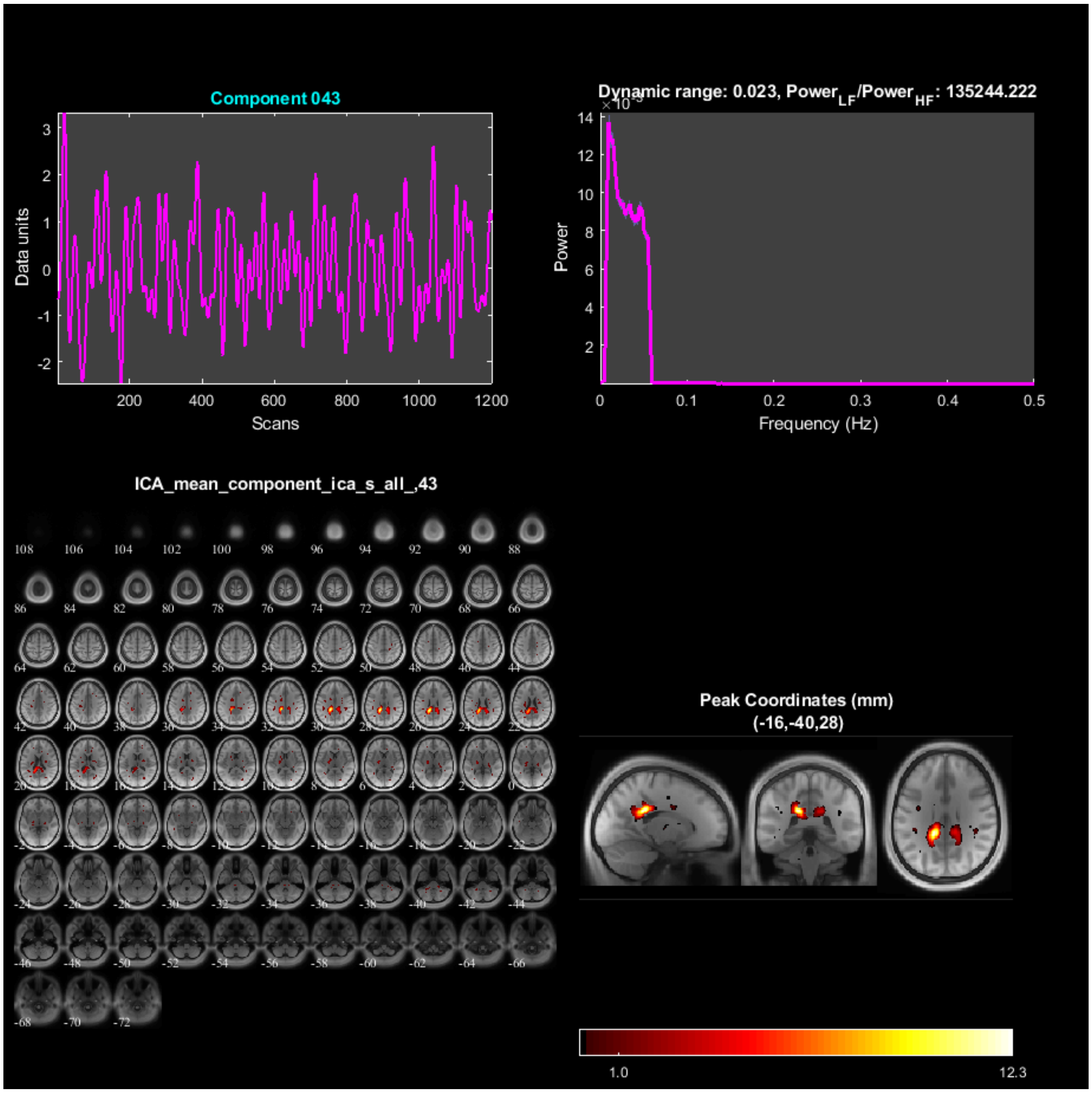
Component 043
Data units
Scans
Dynamic range: 0.023, Power$_{LF}$/Power$_{HF}$: 135244.222
Power
Frequency (Hz)
ICA_mean_component_ica_s_all_,43
Peak Coordinates (mm)
(-16,-40,28)
1.0
12.3

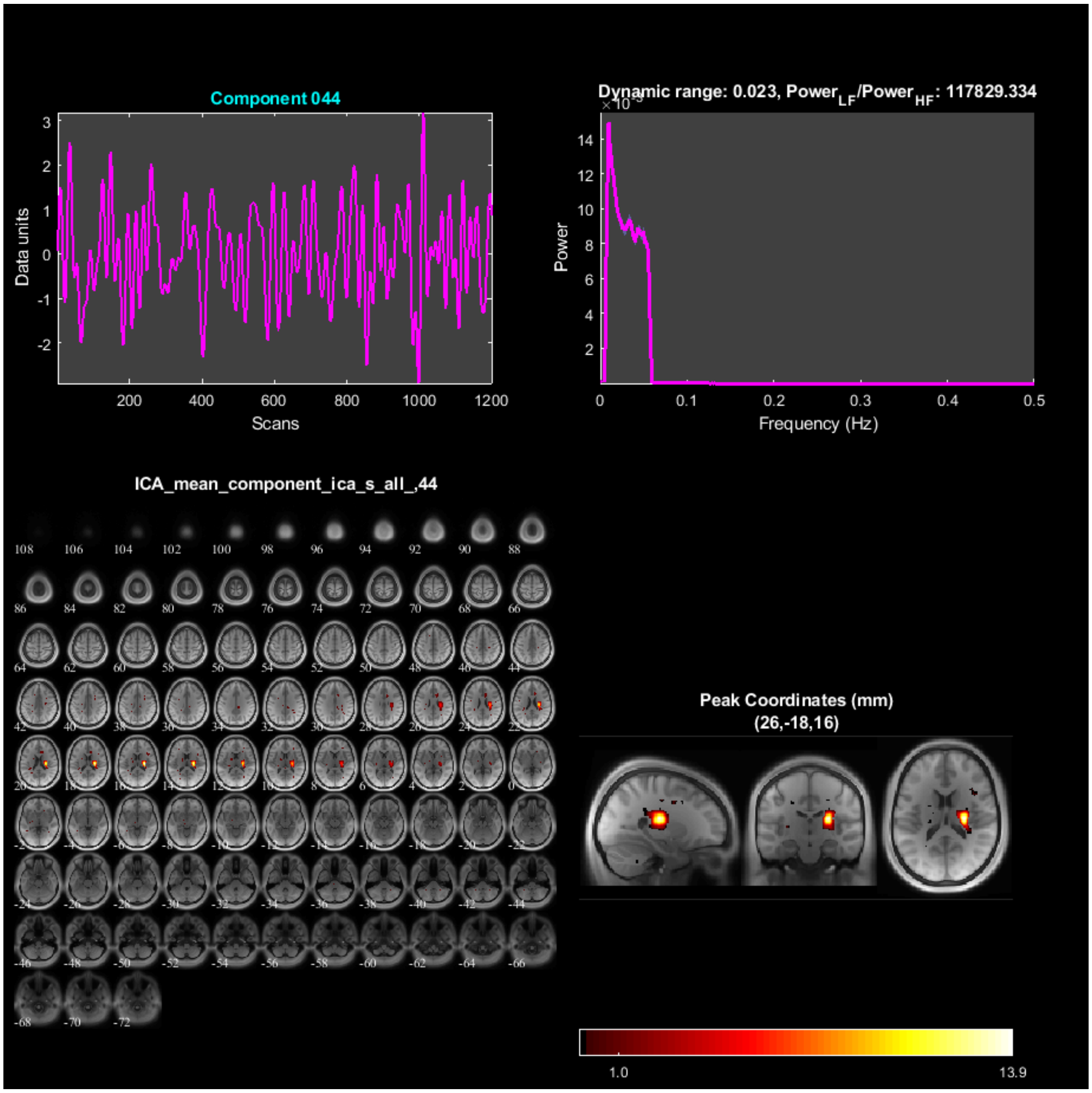

Component 044
Data units
Scans
Dynamic range: 0.023, Power$_{LF}$/Power$_{HF}$: 117829.334
Power
Frequency (Hz)
ICA_mean_component_ica_s_all_,44
Peak Coordinates (mm)
(26,-18,16)
1.0
13.9

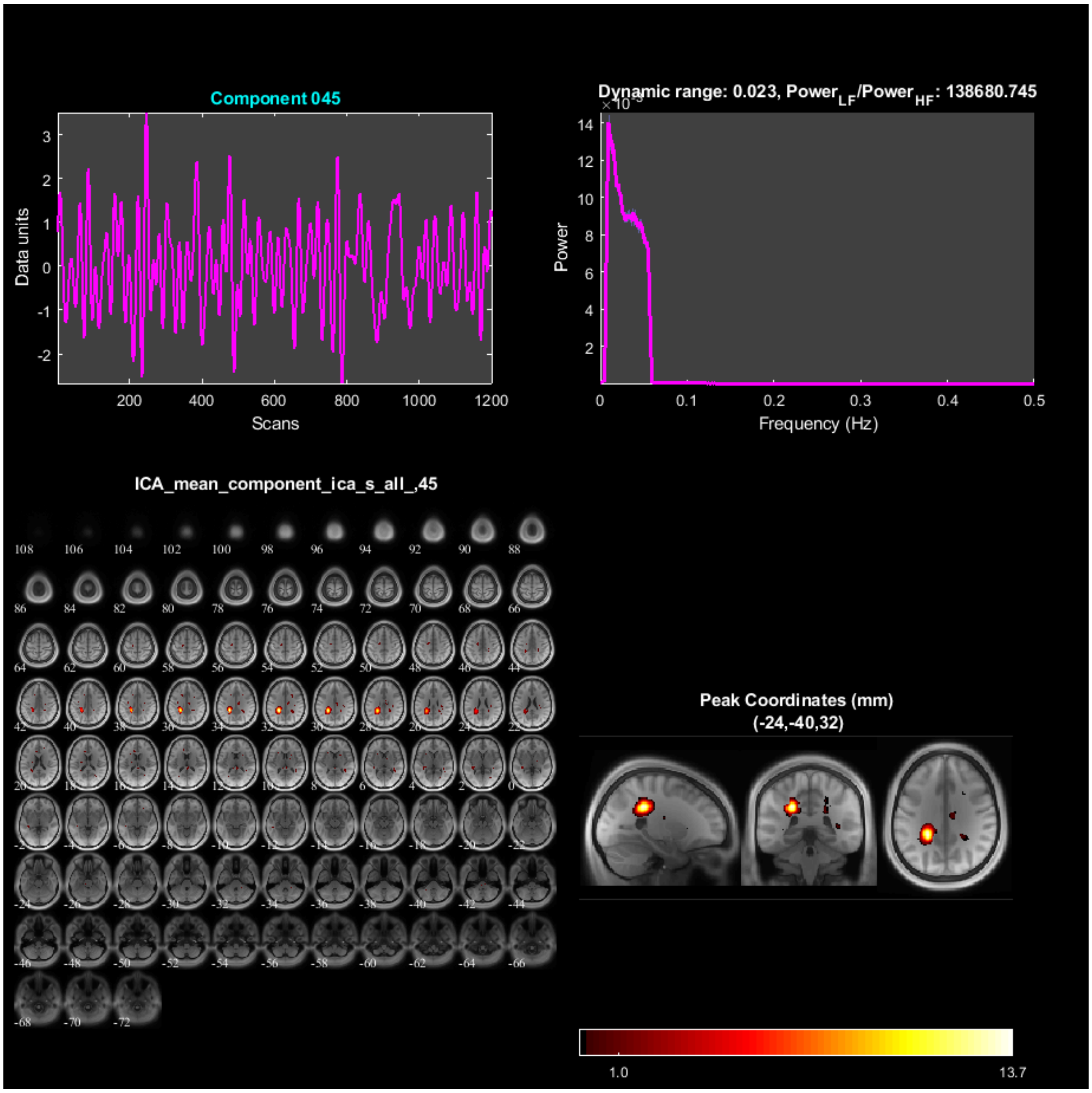
Component 045
Data units
Scans
Dynamic range: 0.023, Power_LF/Power_HF: 138680.745
Power
Frequency (Hz)
ICA_mean_component_ica_s_all_,45
Peak Coordinates (mm)
(-24,-40,32)
1.0
13.7

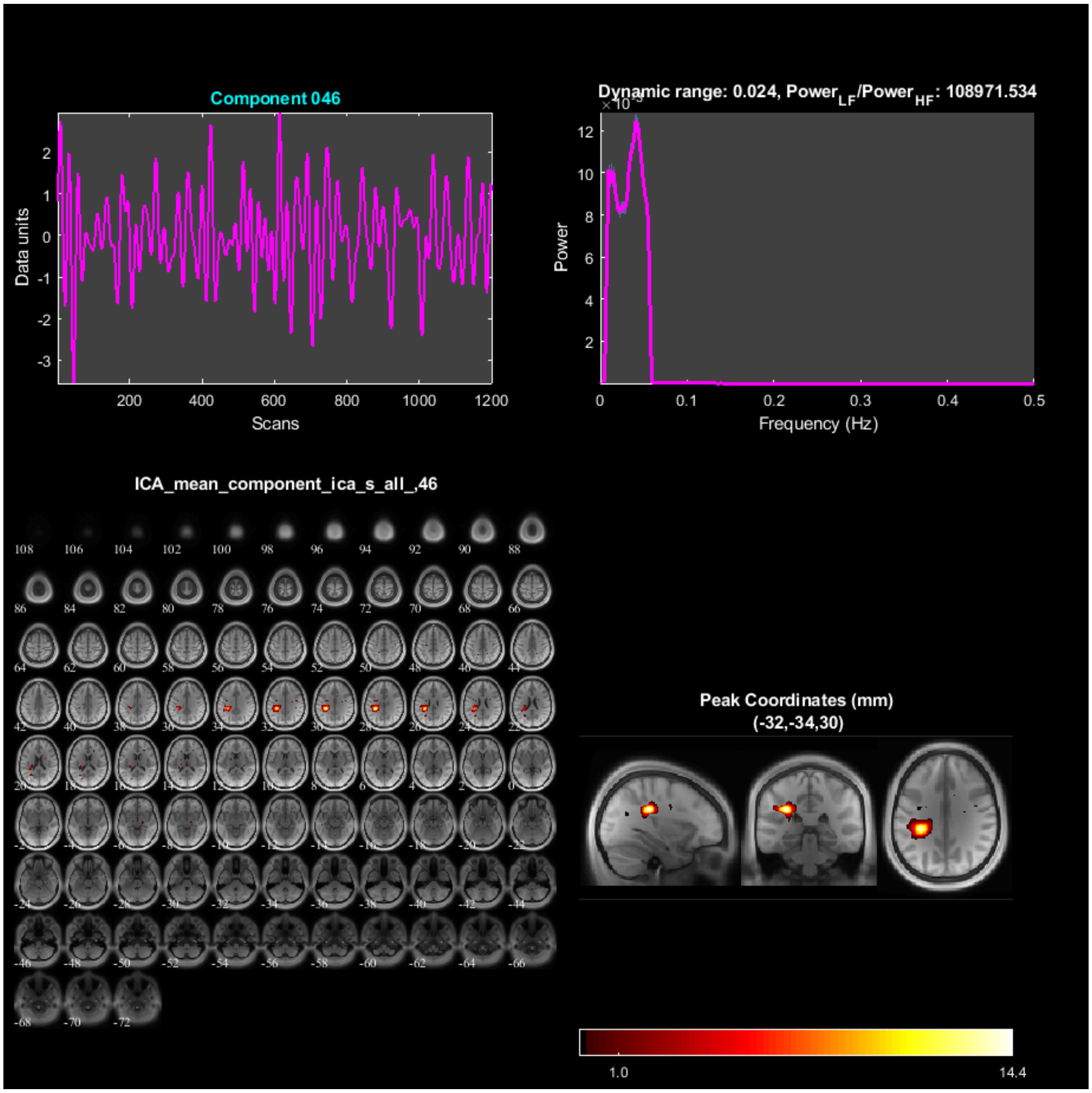
Component 046
Data units
Scans
Dynamic range: 0.024, Power$_{LF}$/Power$_{HF}$: 108971.534
Power
Frequency (Hz)
ICA_mean_component_ica_s_all_,46
Peak Coordinates (mm)
(-32,-34,30)
1.0
14.4

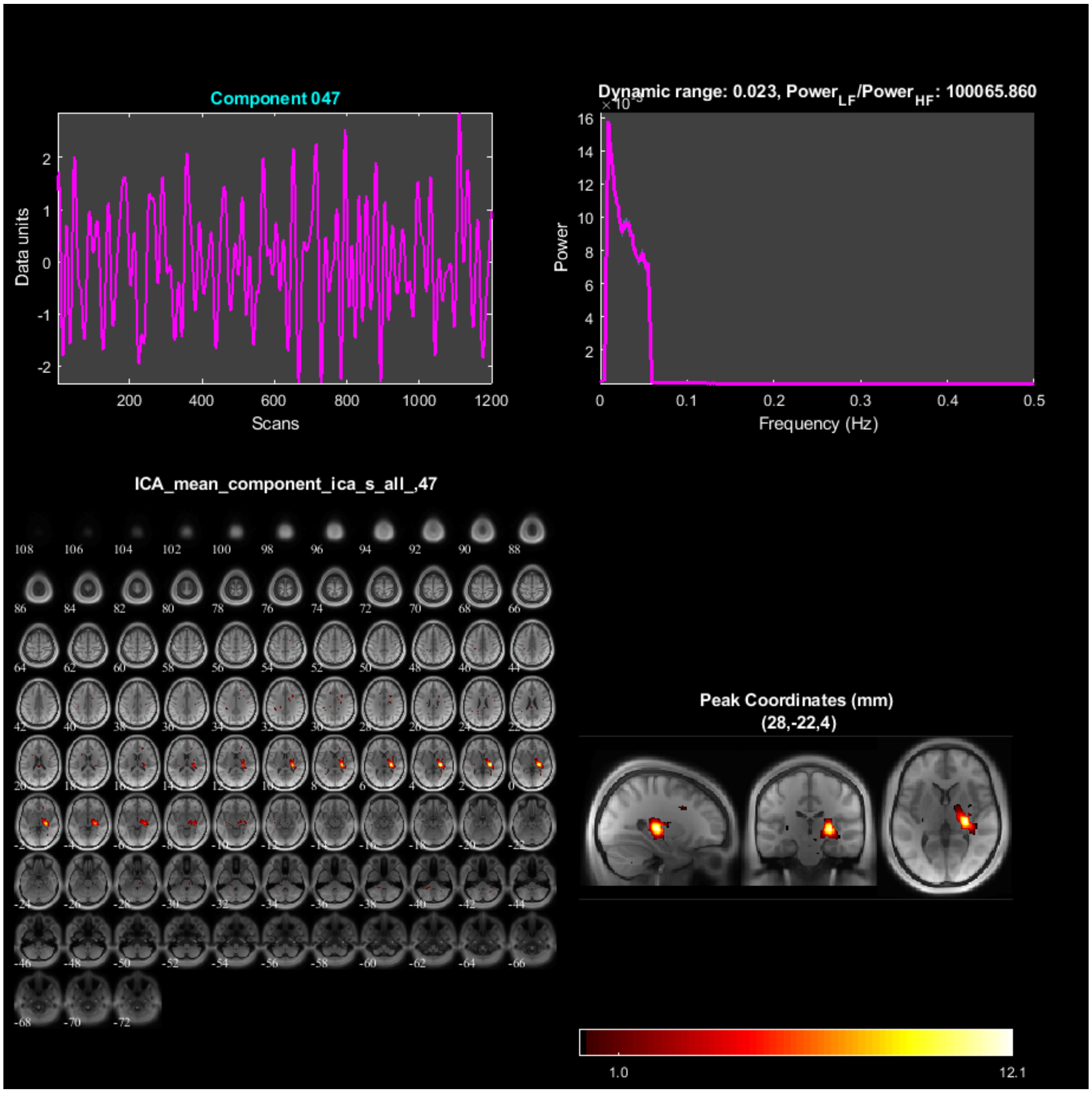
Component 047
Data units
Scans
Dynamic range: 0.023, Power_LF/Power_HF: 100065.860
Power
Frequency (Hz)
ICA_mean_component_ica_s_all_,47
Peak Coordinates (mm)
(28,-22,4)
1.0
12.1

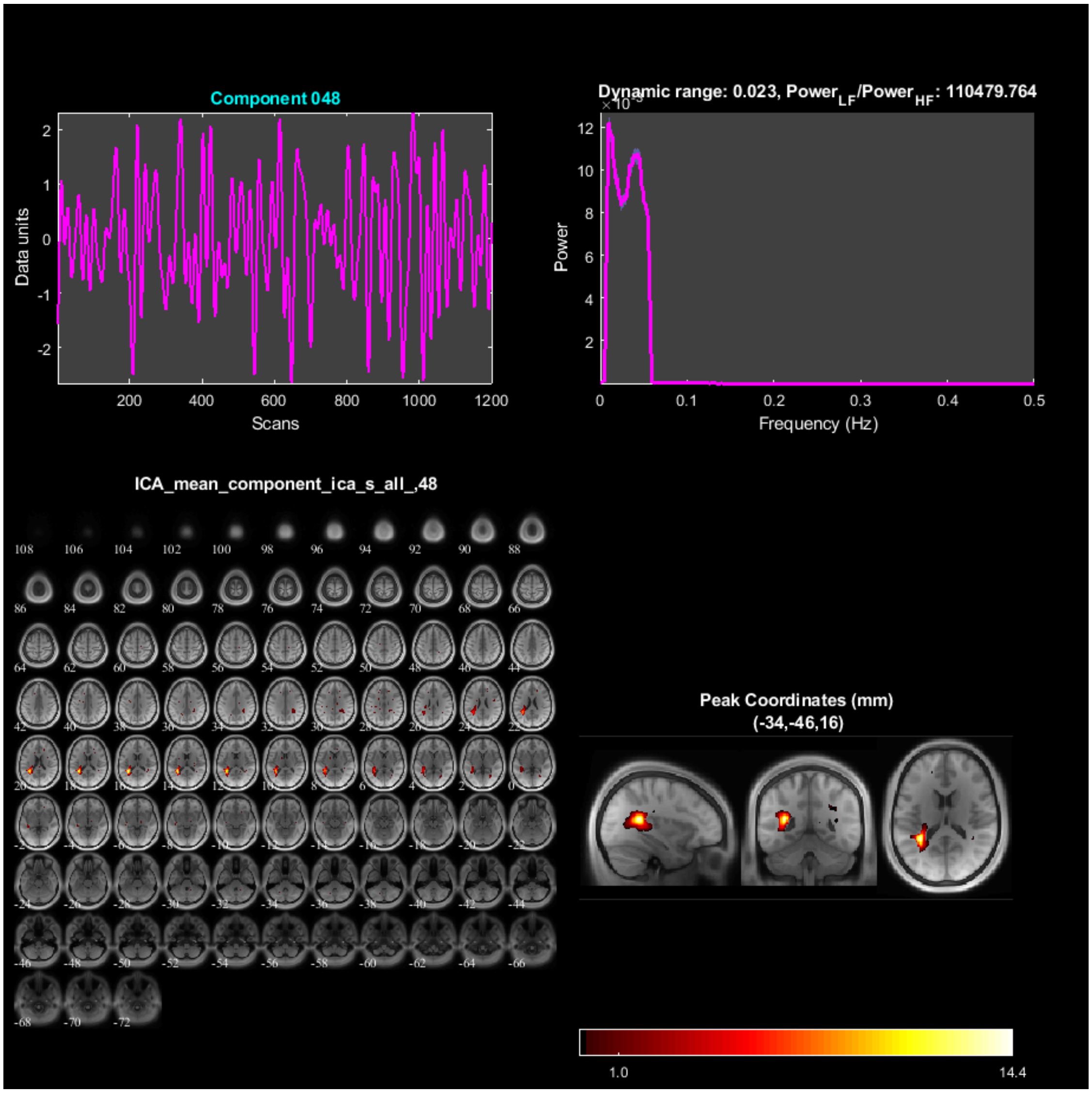

Component 048
Data units
Scans
Dynamic range: 0.023, Power$_{LF}$/Power$_{HF}$: 110479.764
Power
Frequency (Hz)
ICA_mean_component_ica_s_all_,48
Peak Coordinates (mm)
(-34,-46,16)
1.0
14.4

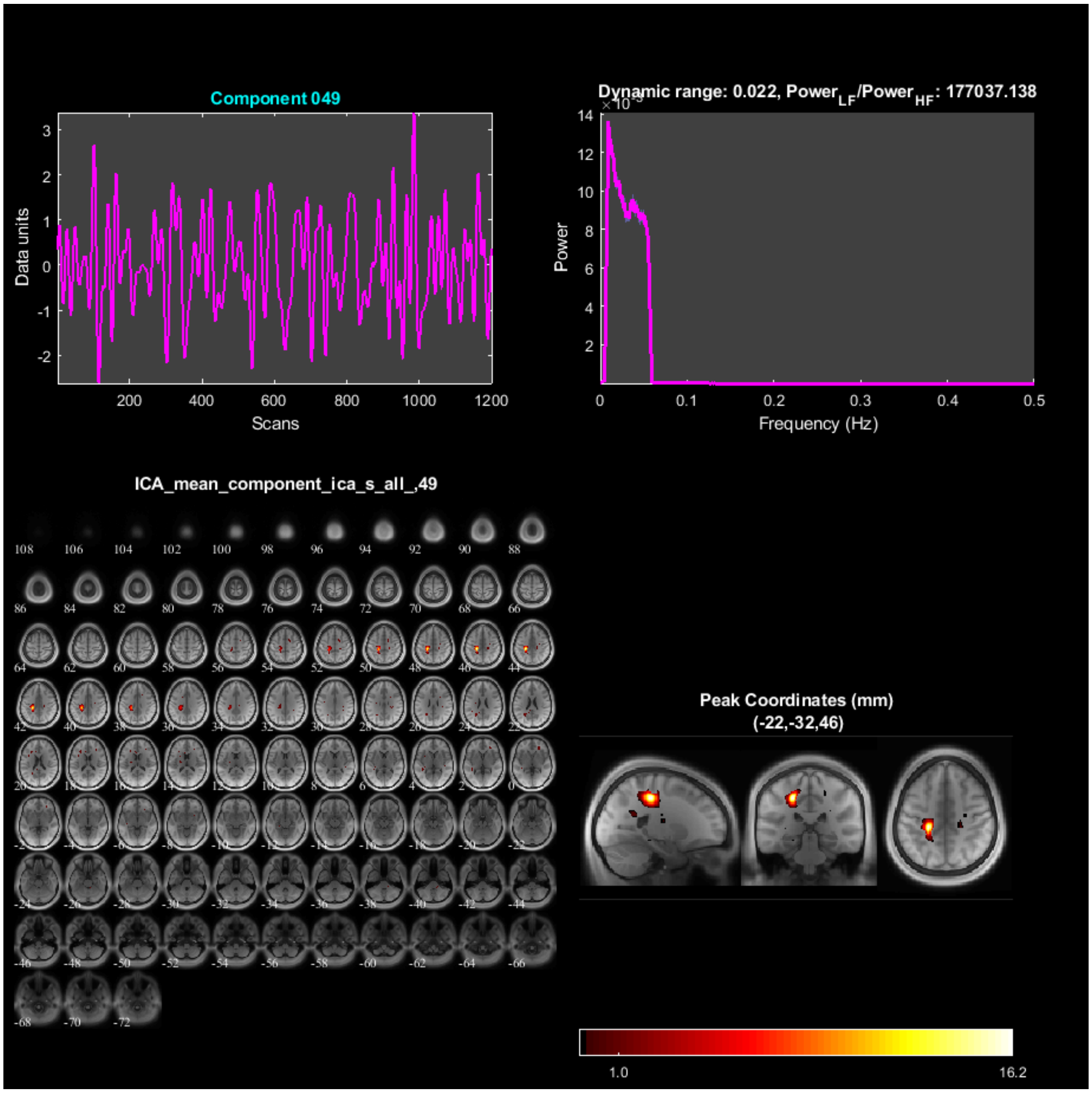

Component 049
Data units
Scans
Dynamic range: 0.022, Power$_{LF}$/Power$_{HF}$: 177037.138
Power
Frequency (Hz)
ICA_mean_component_ica_s_all_,49
Peak Coordinates (mm)
(-22,-32,46)
1.0
16.2

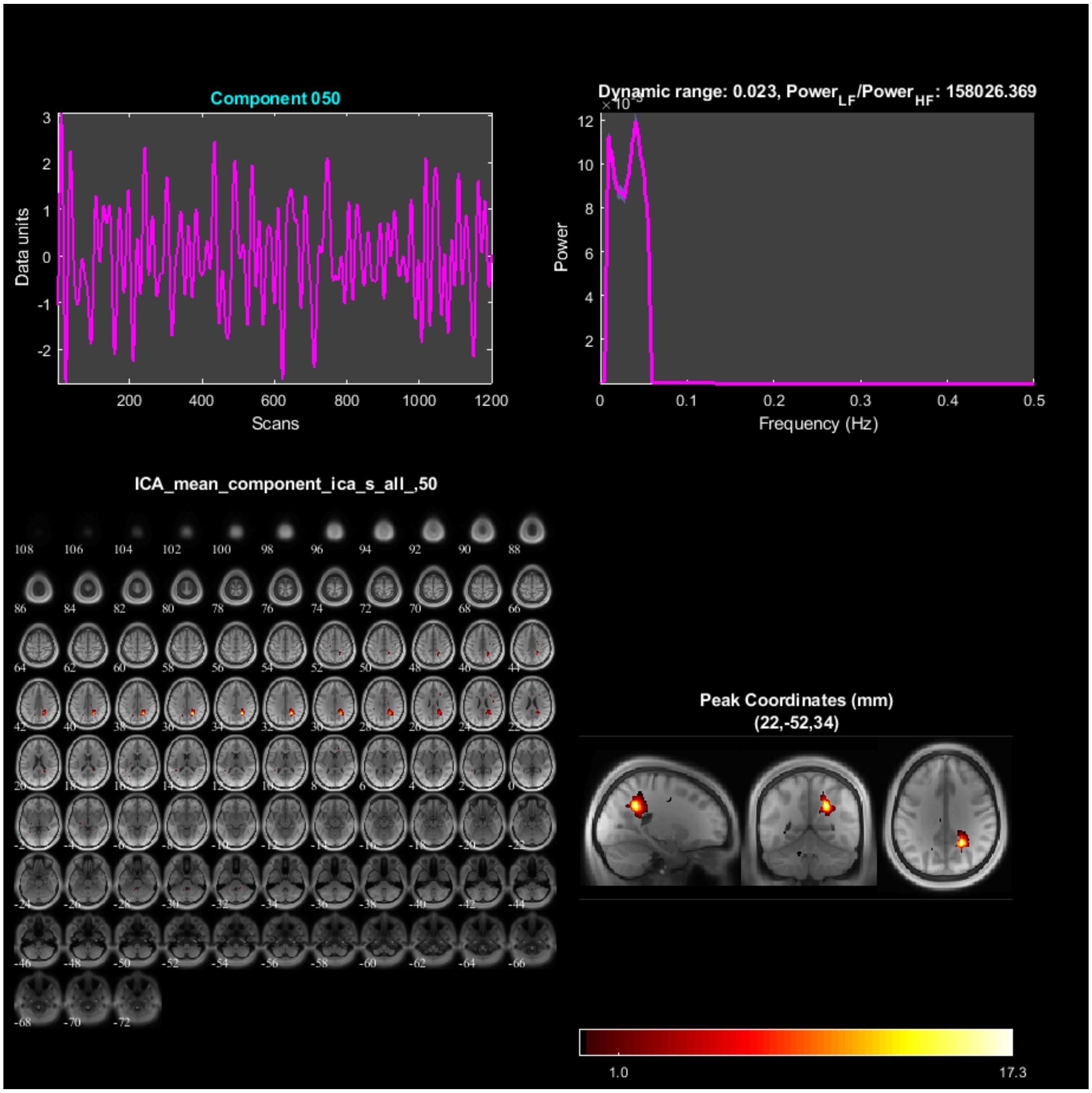
Component 050
Data units
Scans
Dynamic range: 0.023, Power$_{LF}$/Power$_{HF}$: 158026.369
Power
Frequency (Hz)
ICA_mean_component_ica_s_all_,50
Peak Coordinates (mm)
(22,-52,34)
1.0
17.3

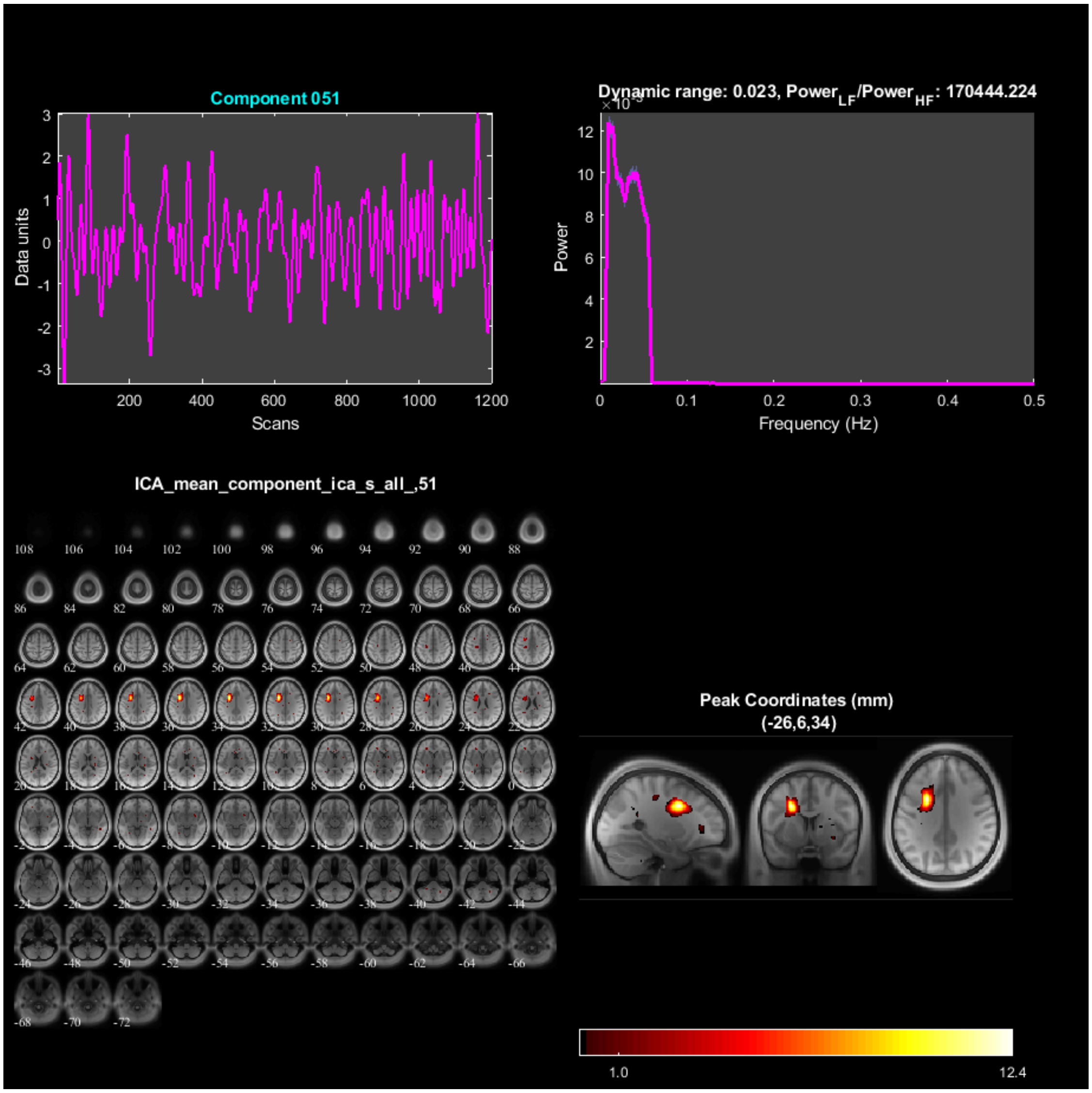

Component 051
Data units
Scans
Dynamic range: 0.023, Power$_{LF}$/Power$_{HF}$: 170444.224
Power
Frequency (Hz)
ICA_mean_component_ica_s_all_,51
Peak Coordinates (mm)
(-26,6,34)
1.0
12.4

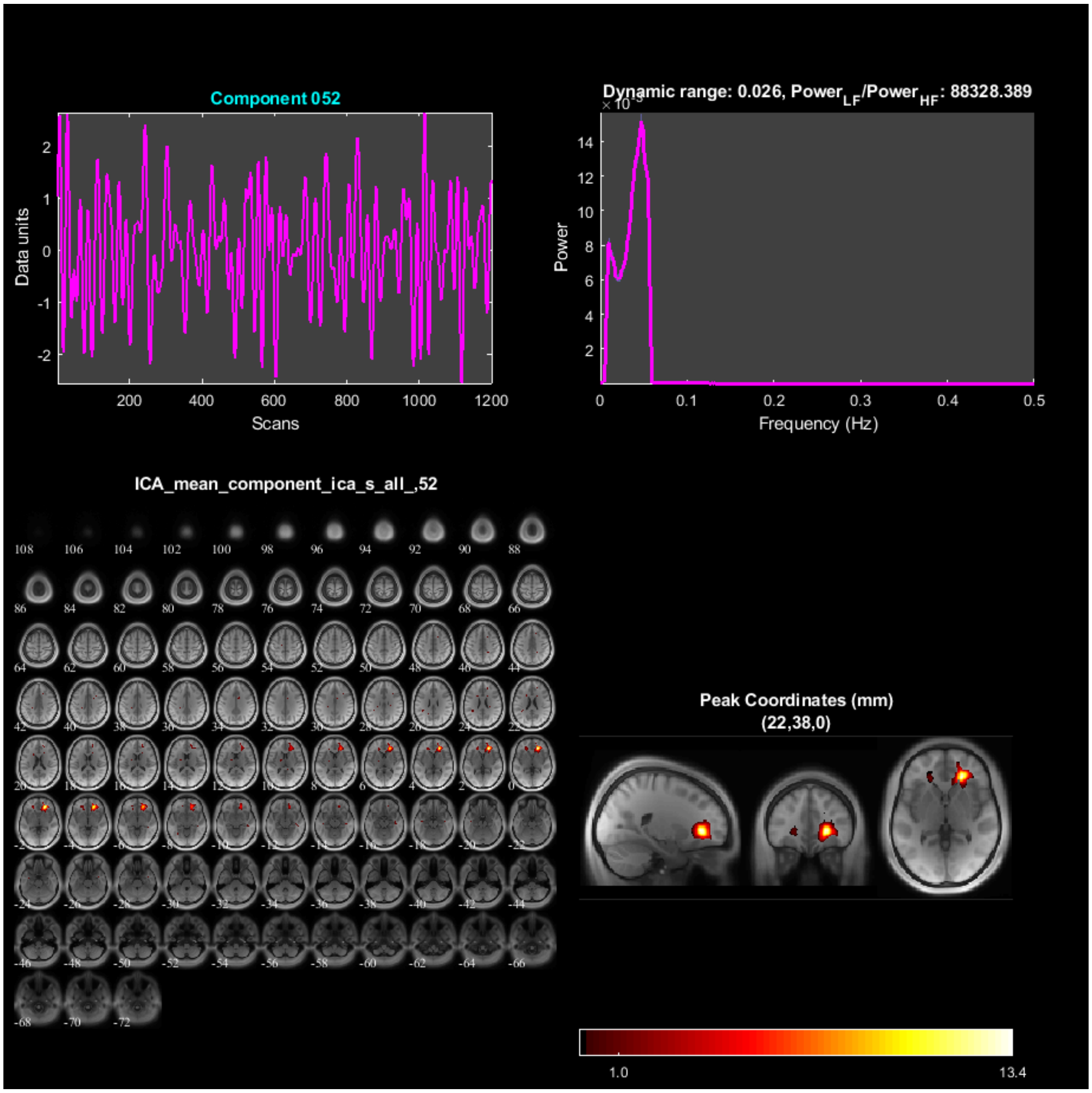
Component 052
Data units
Scans
Dynamic range: 0.026, Power$_{LF}$/Power$_{HF}$: 88328.389
Power
Frequency (Hz)
ICA_mean_component_ica_s_all_,52
Peak Coordinates (mm)
(22,38,0)
1.0
13.4

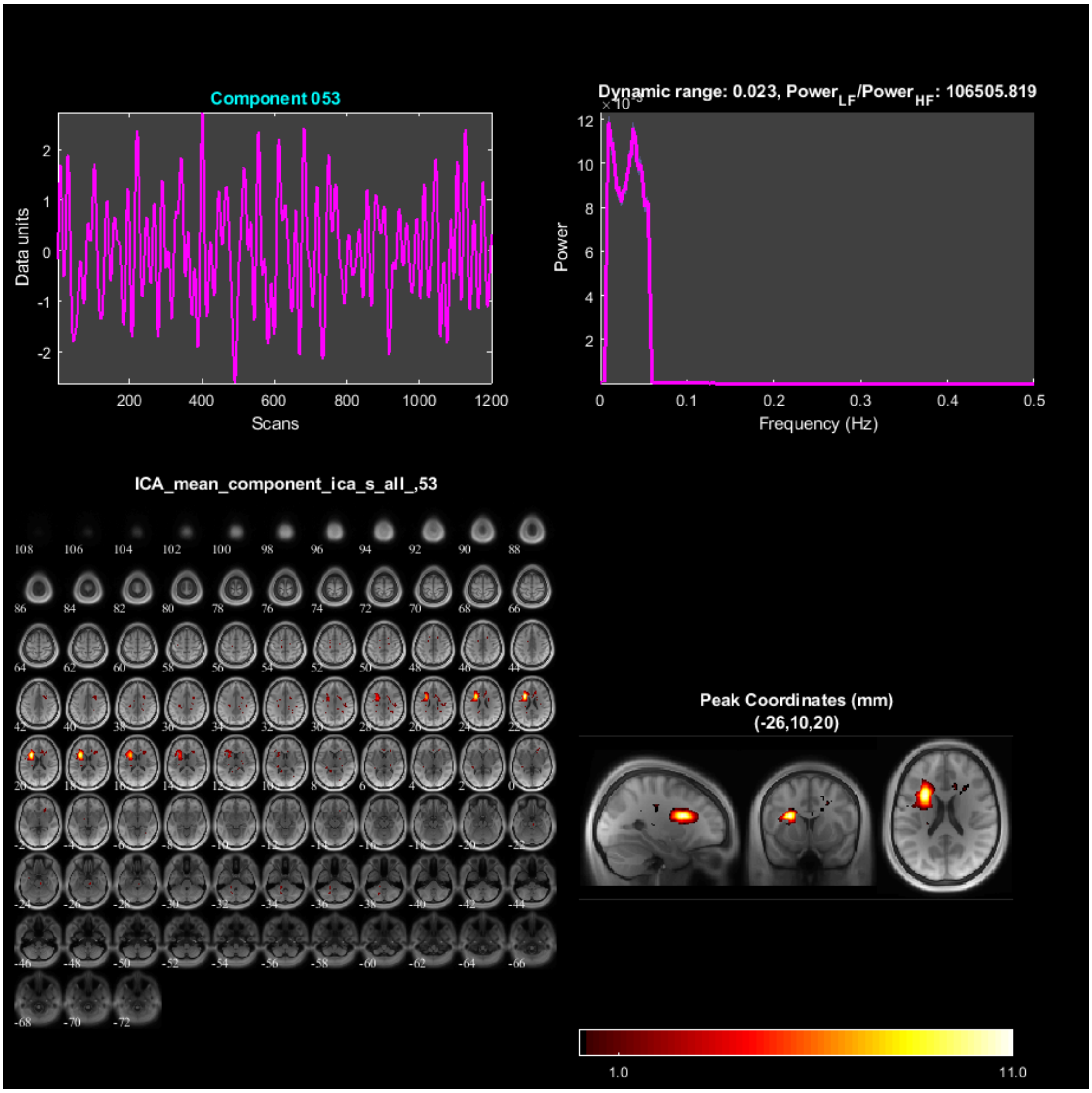

Component 053
Data units
Scans
Dynamic range: 0.023, Power$_{LF}$/Power$_{HF}$: 106505.819
Power
Frequency (Hz)
ICA_mean_component_ica_s_all_,53
Peak Coordinates (mm)
(-26,10,20)
1.0
11.0

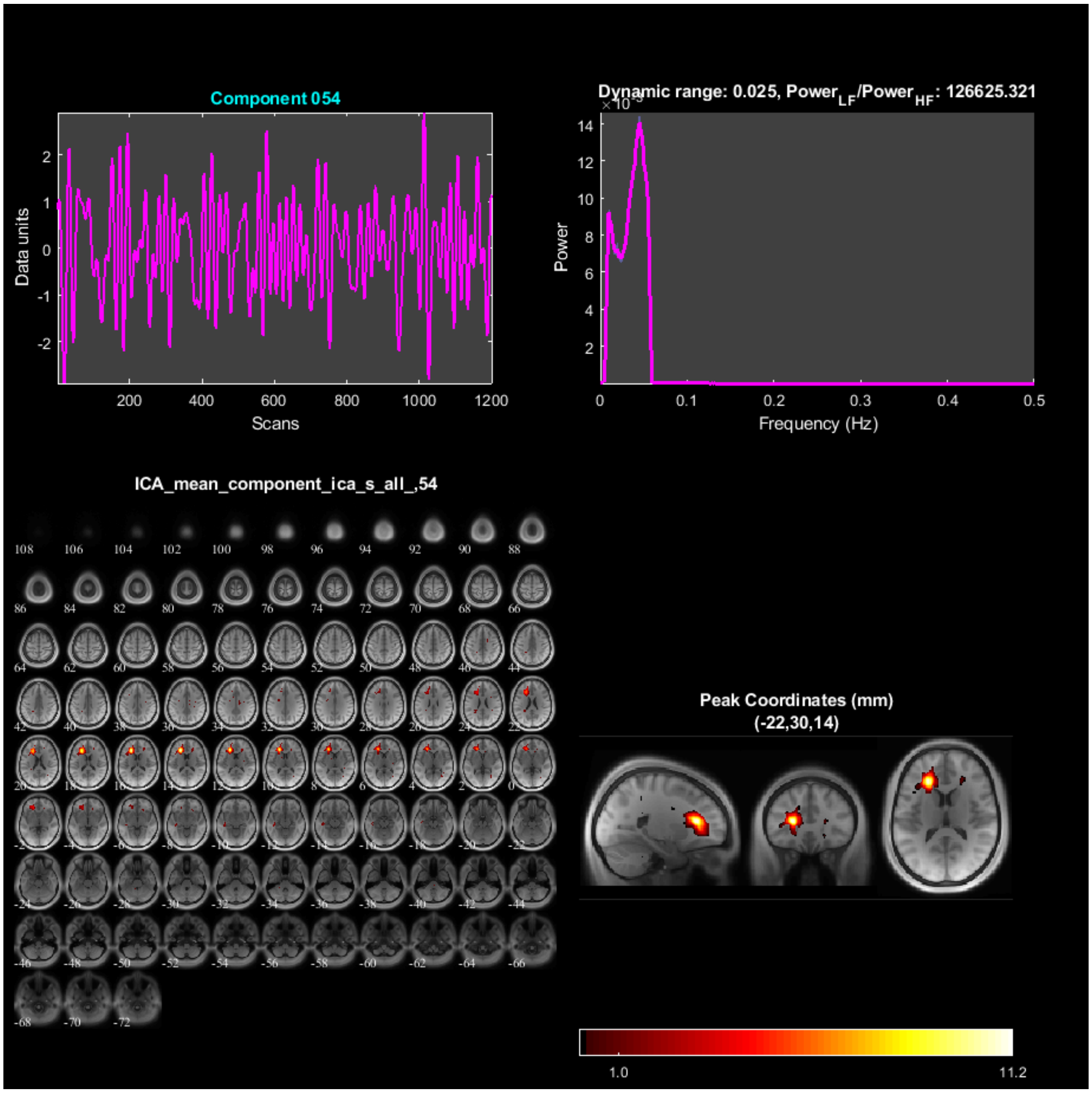
Component 054
Data units
Scans
Dynamic range: 0.025, Power$_{LF}$/Power$_{HF}$: 126625.321
Power
Frequency (Hz)
ICA_mean_component_ica_s_all_,54
Peak Coordinates (mm)
(-22,30,14)
1.0
11.2

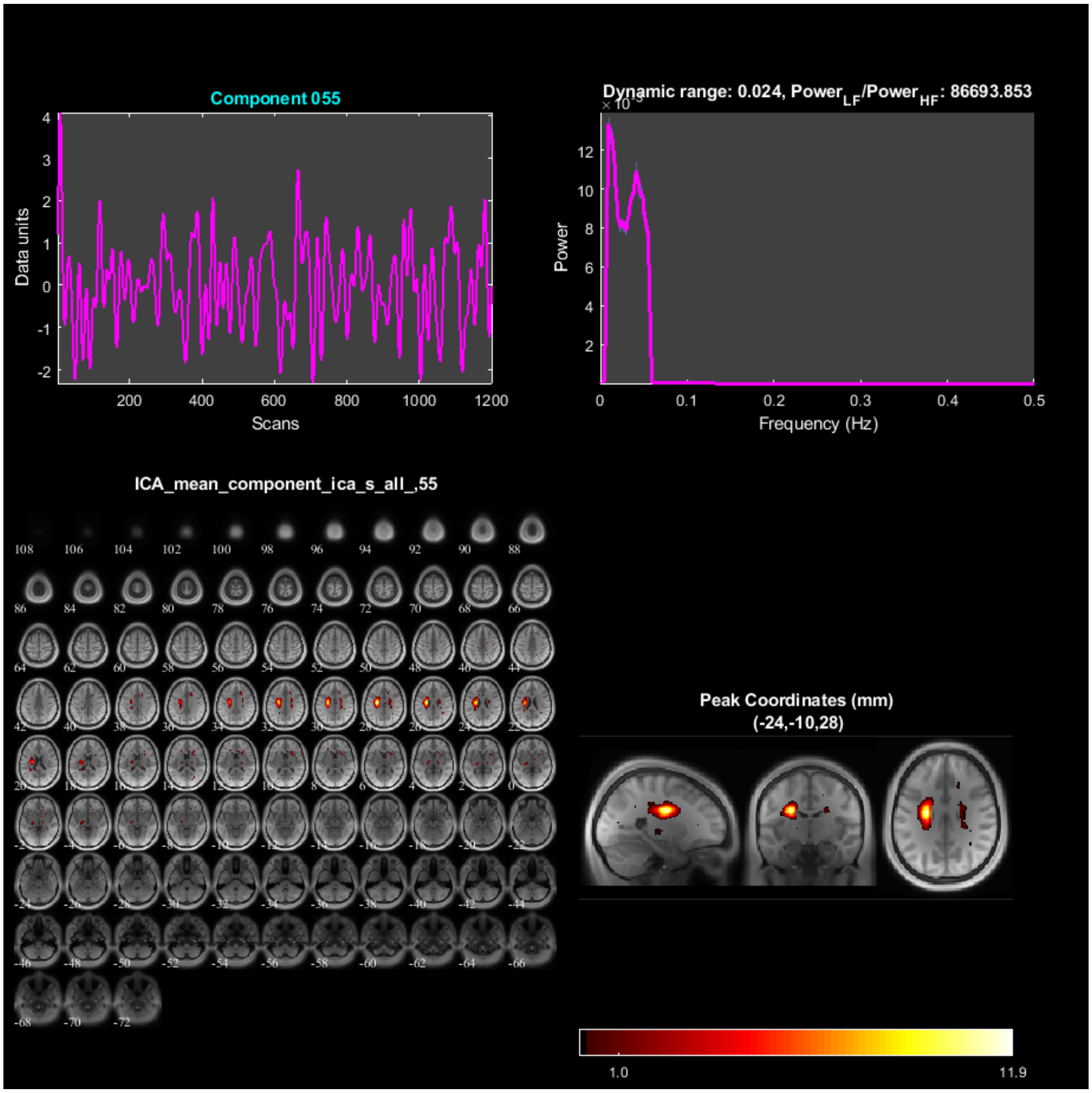
Component 055
Data units
Scans
Dynamic range: 0.024, Power$_{LF}$/Power$_{HF}$: 86693.853
Power
Frequency (Hz)
ICA_mean_component_ica_s_all_,55
Peak Coordinates (mm)
(-24,-10,28)
1.0
11.9

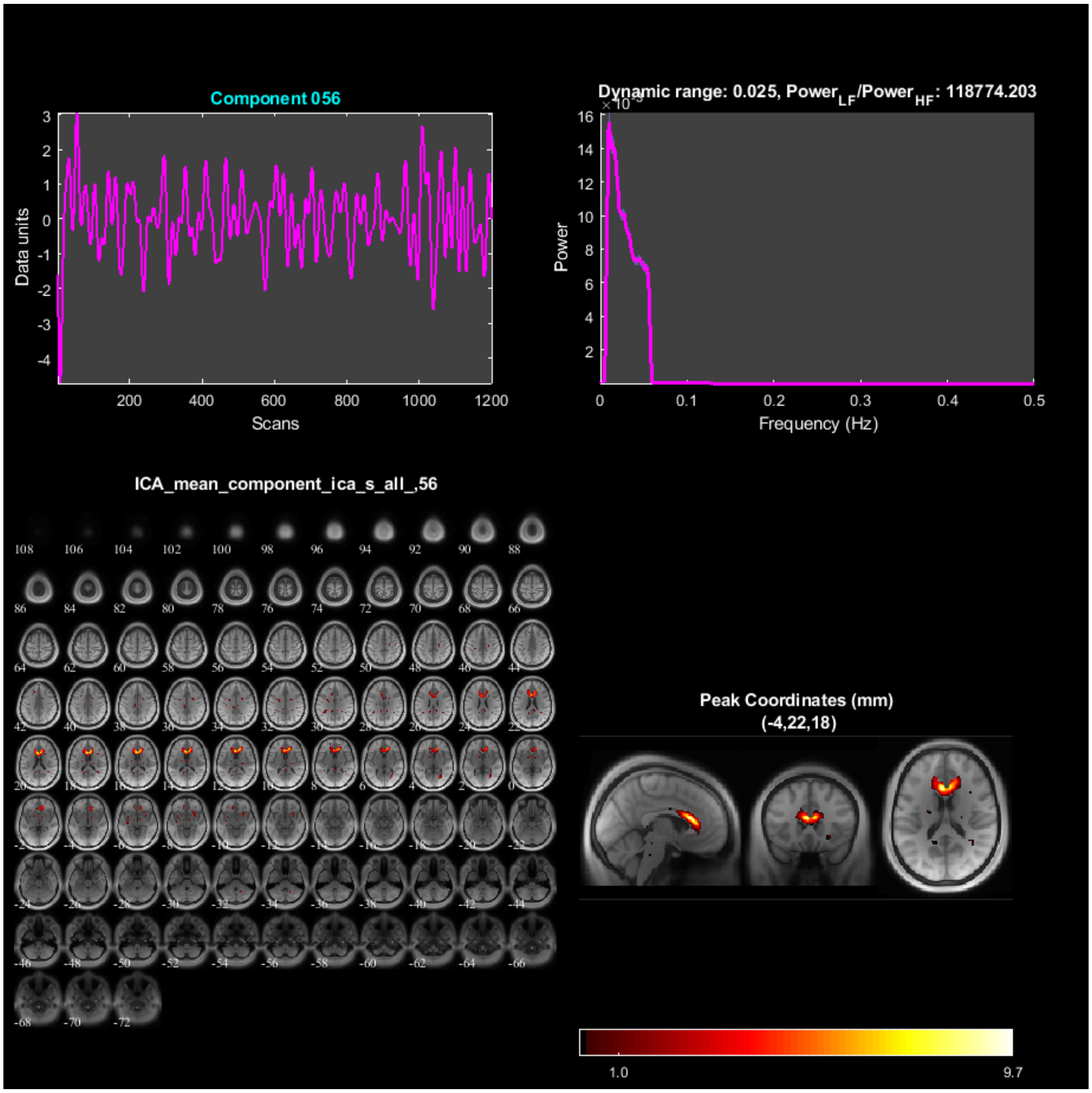

Component 056
Data units
Scans
Dynamic range: 0.025, Power$_{LF}$/Power$_{HF}$: 118774.203
Power
Frequency (Hz)
ICA_mean_component_ica_s_all_,56
Peak Coordinates (mm)
(-4,22,18)
1.0
9.7

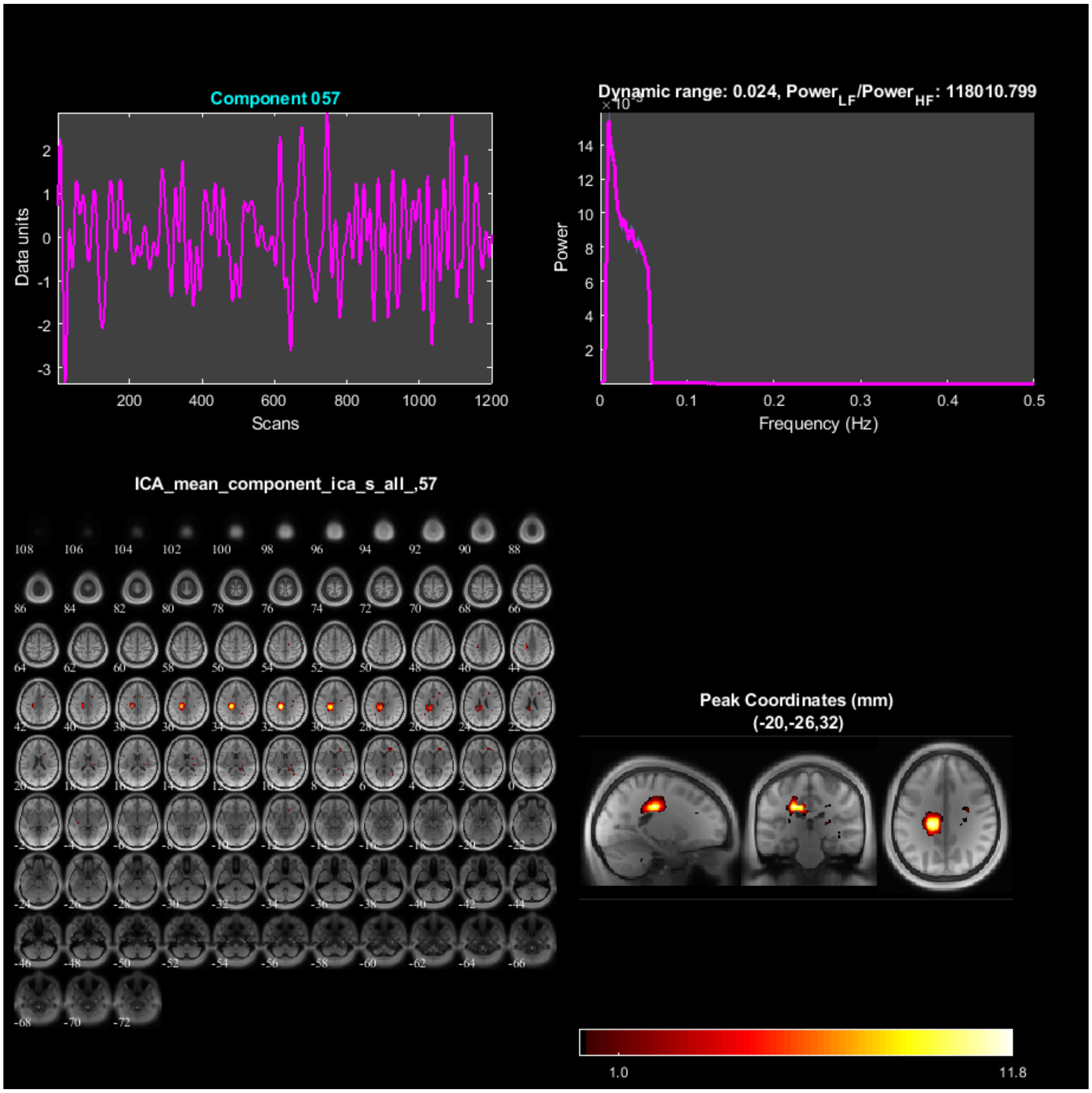
Component 057
Data units
Scans
Dynamic range: 0.024, Power$_{LF}$/Power$_{HF}$: 118010.799
Power
Frequency (Hz)
ICA_mean_component_ica_s_all_,57
Peak Coordinates (mm)
(-20,-26,32)
1.0
11.8

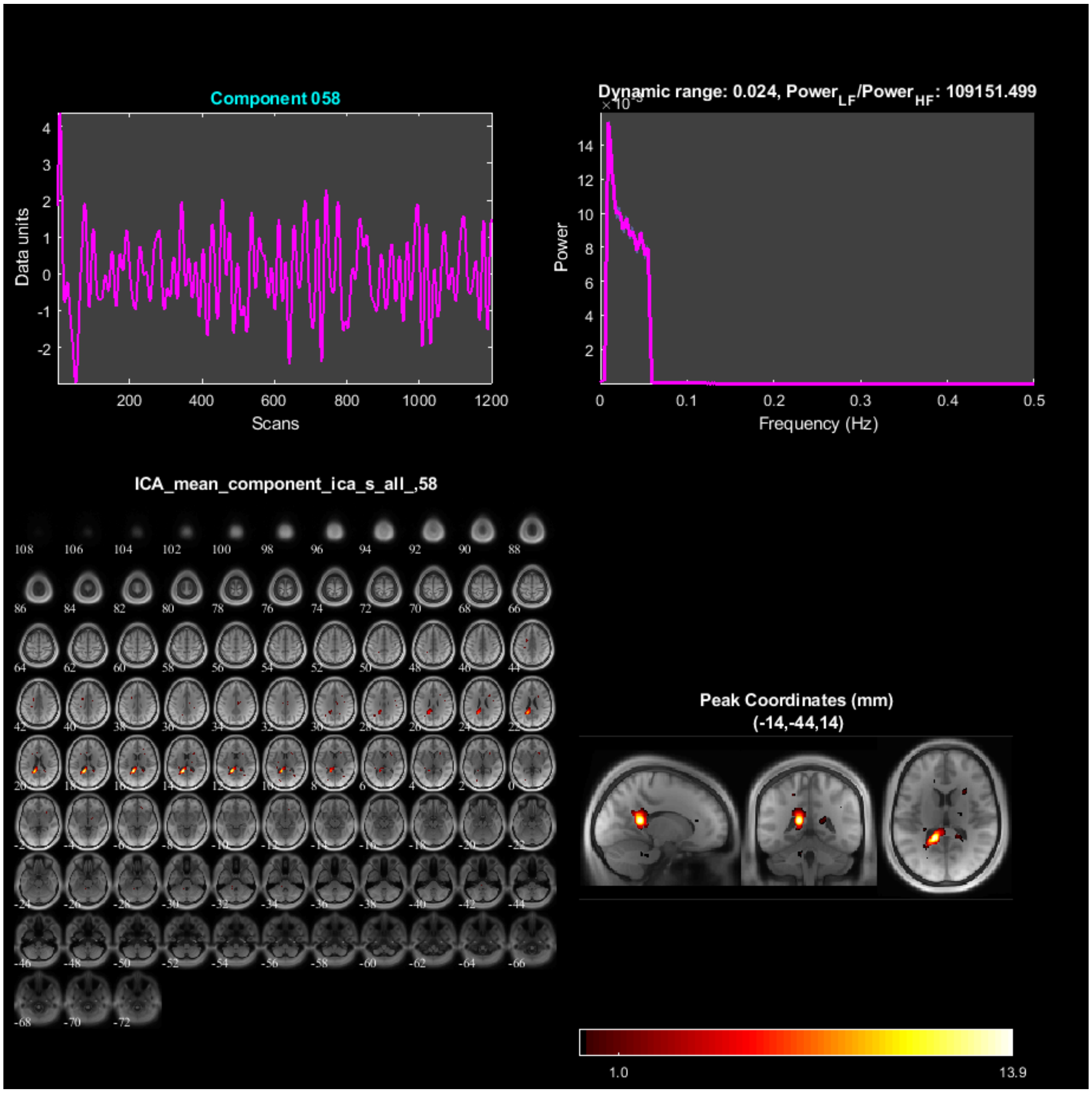
Component 058
Data units
Scans
Dynamic range: 0.024, Power_LF/Power_HF: 109151.499
Power
Frequency (Hz)
ICA_mean_component_ica_s_all_,58
Peak Coordinates (mm)
(-14,-44,14)
1.0
13.9

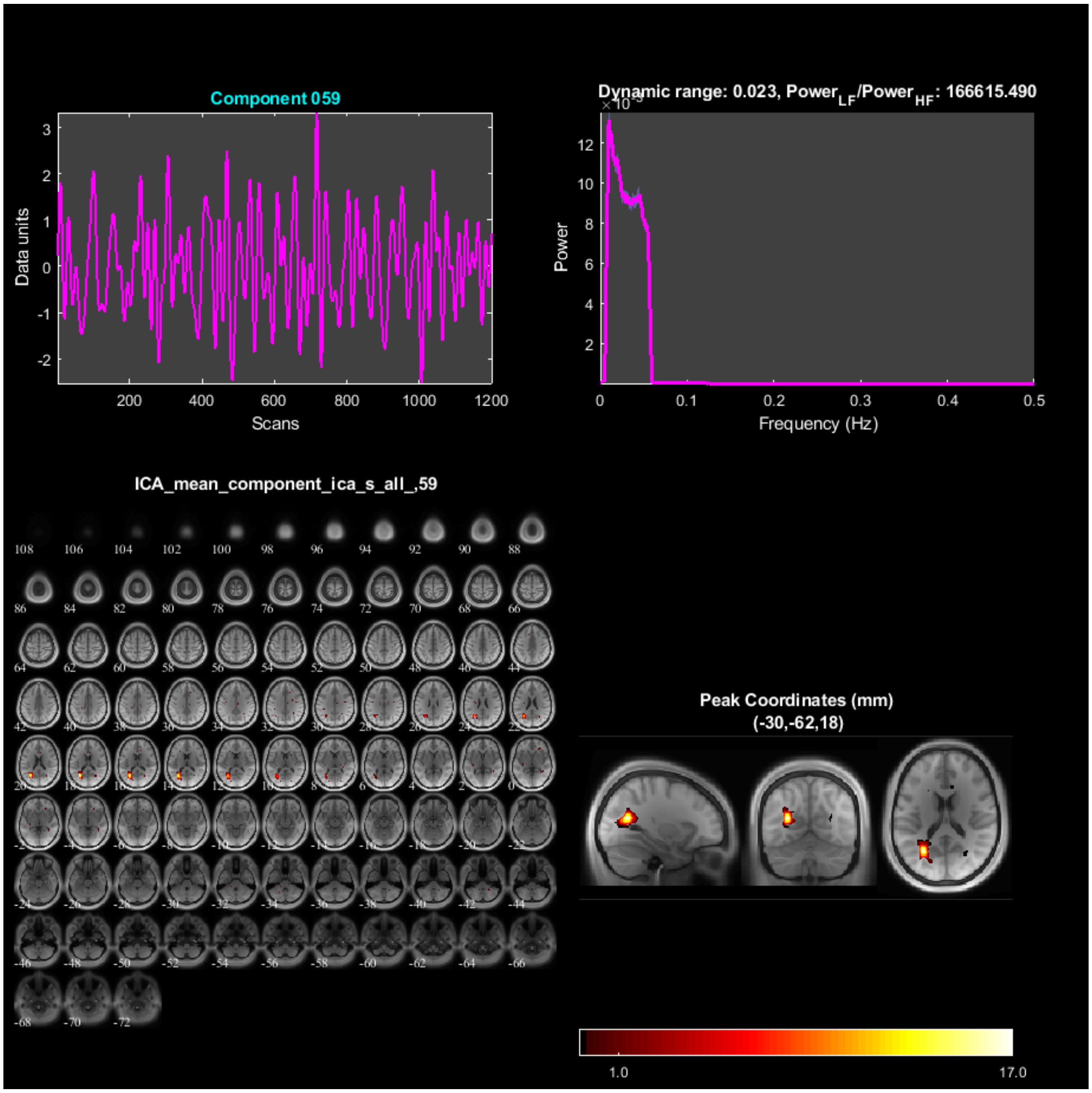

Component 059
Data units
Scans
Dynamic range: 0.023, Power$_{LF}$/Power$_{HF}$: 166615.490
Power
Frequency (Hz)
ICA_mean_component_ica_s_all_,59
Peak Coordinates (mm)
(-30,-62,18)
1.0
17.0

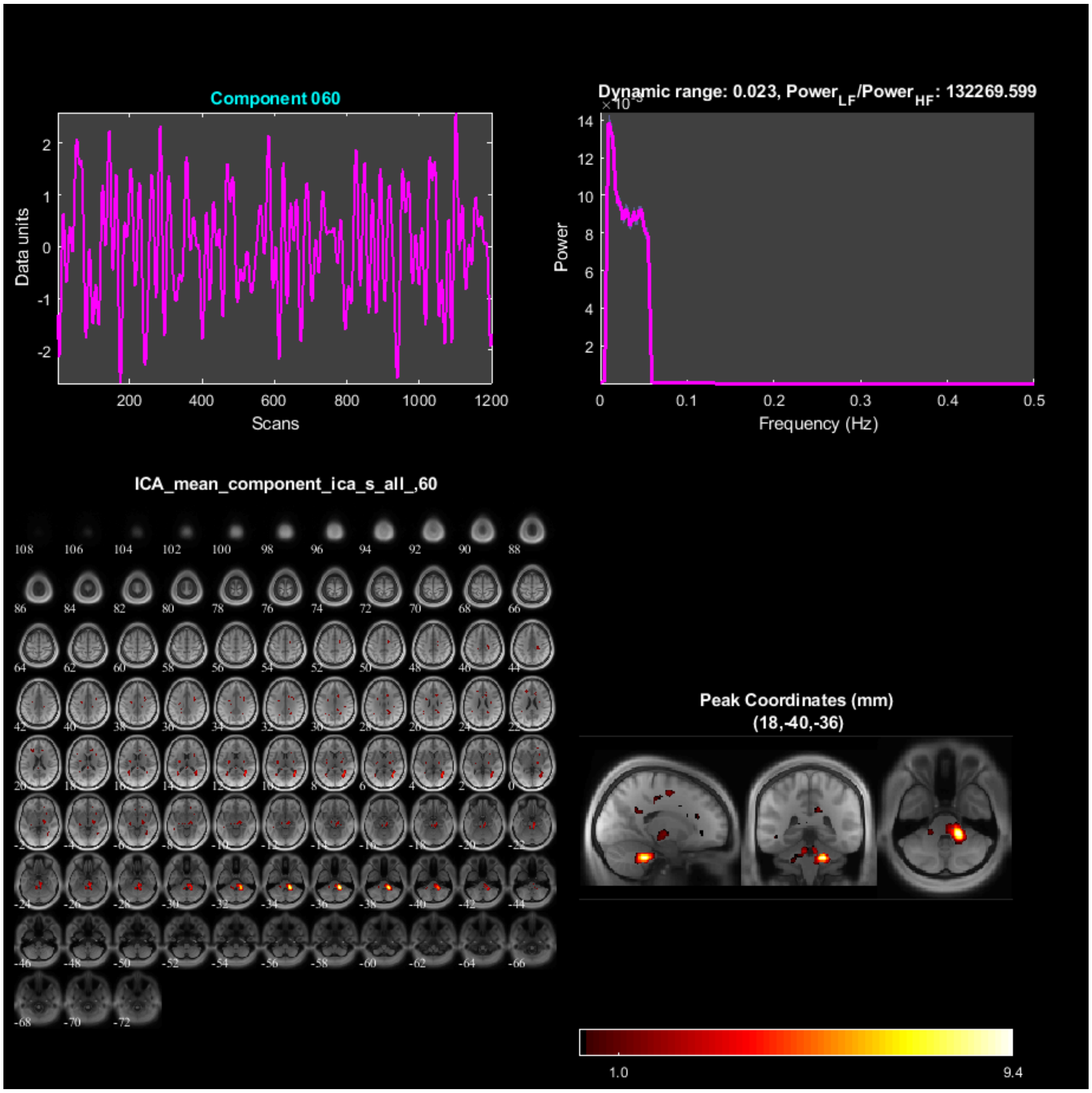
Component 060
Data units
Scans
Dynamic range: 0.023, Power_LF/Power_HF: 132269.599
Power
Frequency (Hz)
ICA_mean_component_ica_s_all_,60
Peak Coordinates (mm)
(18,-40,-36)
1.0
9.4

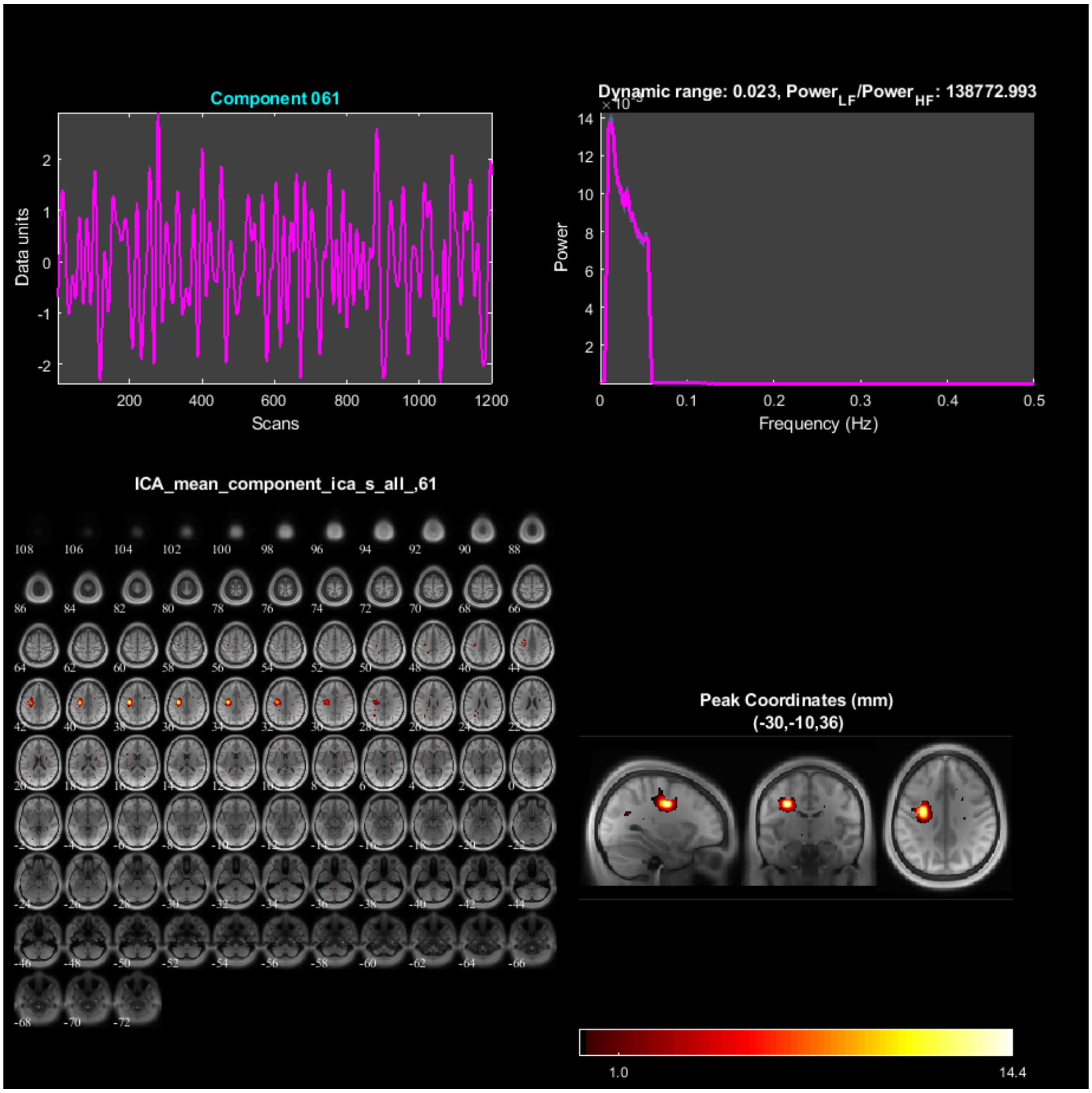
Component 061
Data units
Scans
Dynamic range: 0.023, Power$_{LF}$/Power$_{HF}$: 138772.993
Power
Frequency (Hz)
ICA_mean_component_ica_s_all_,61
Peak Coordinates (mm)
(-30,-10,36)
1.0
14.4

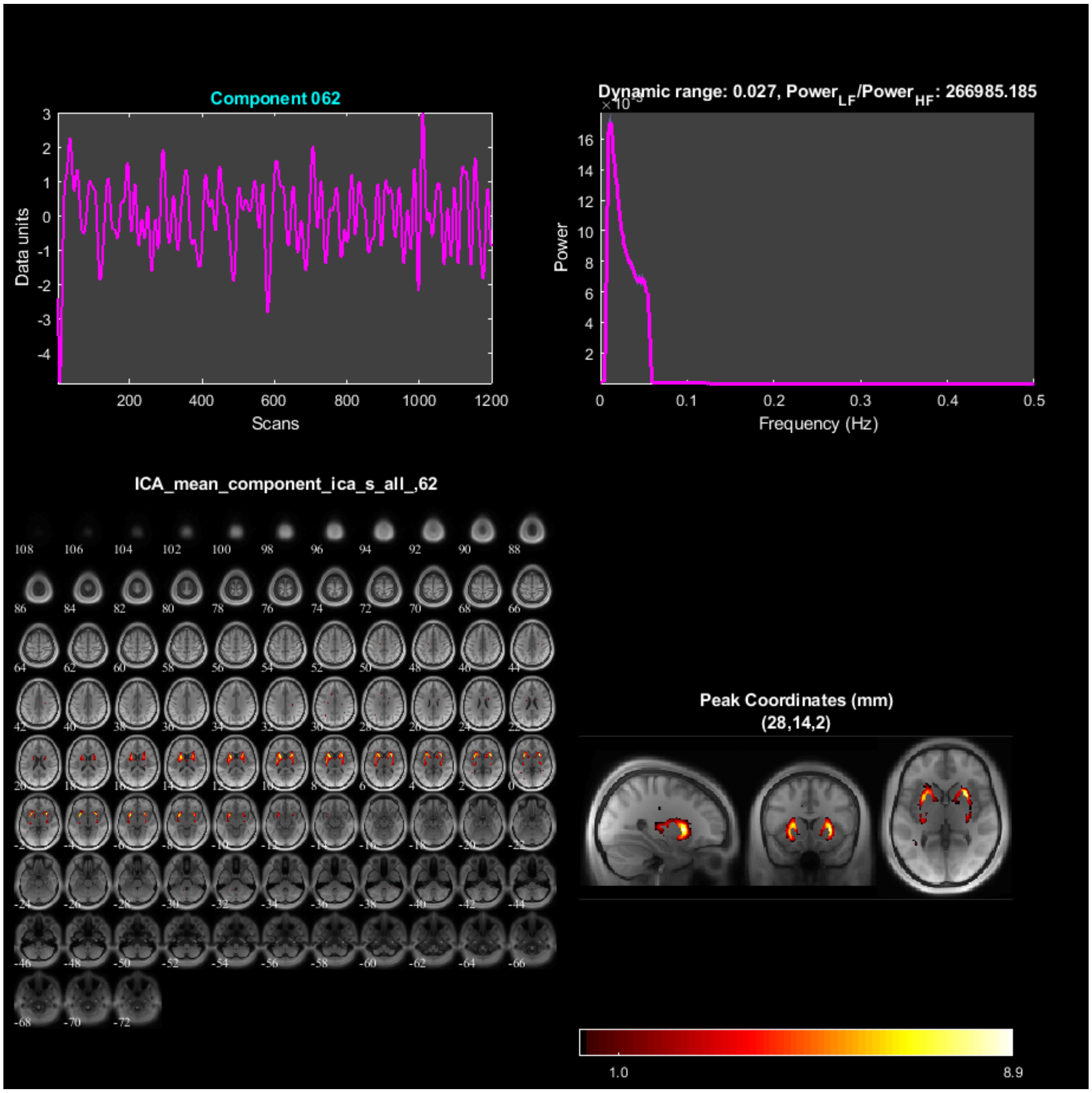
Component 062
Data units
Scans
Dynamic range: 0.027, Power$_{LF}$/Power$_{HF}$: 266985.185
Power
Frequency (Hz)
ICA_mean_component_ica_s_all_,62
Peak Coordinates (mm)
(28,14,2)
1.0
8.9

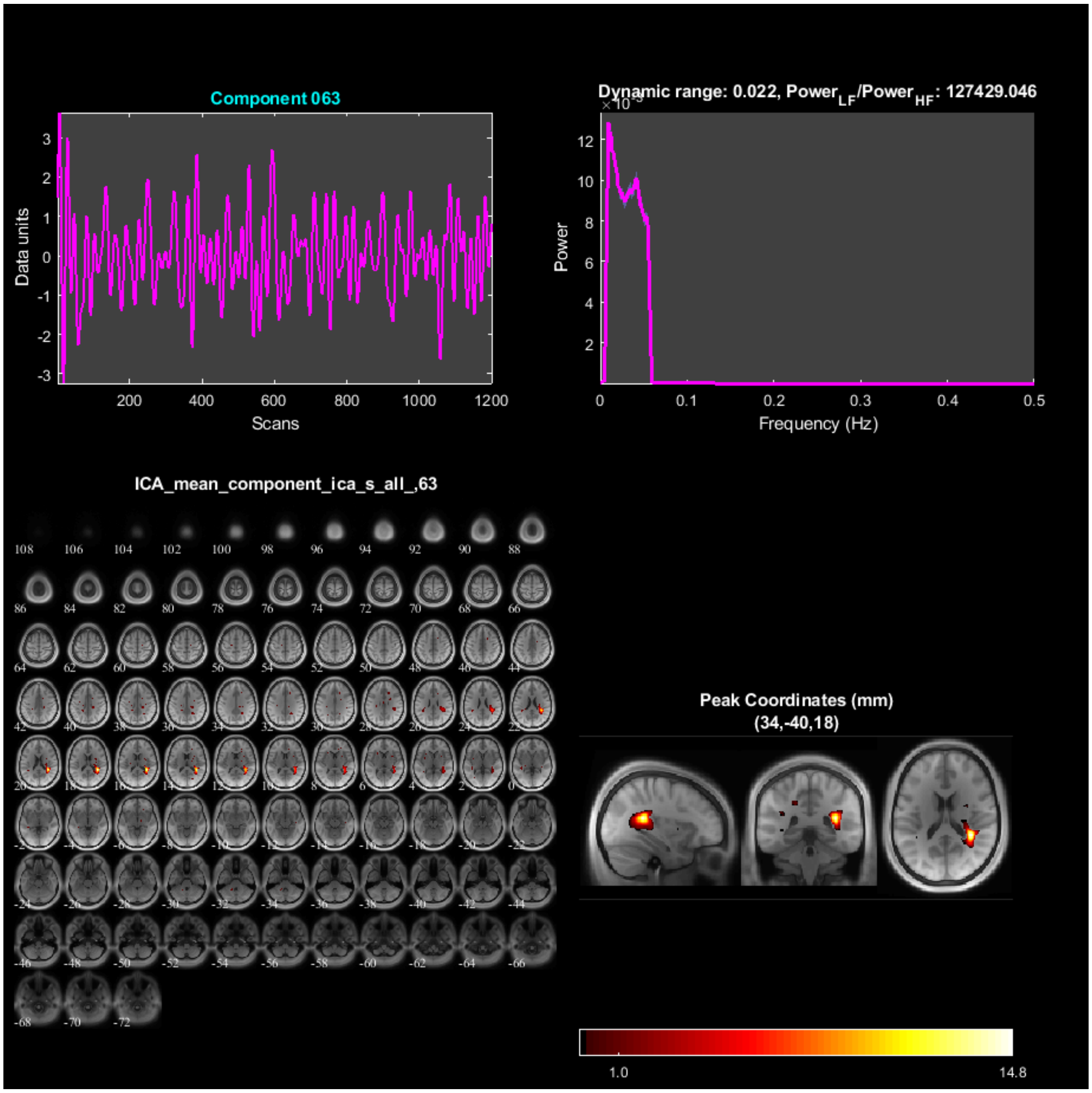

Component 063
Data units
Scans
Dynamic range: 0.022, Power_LF/Power_HF: 127429.046
Power
Frequency (Hz)
ICA_mean_component_ica_s_all_,63
Peak Coordinates (mm)
(34,-40,18)
1.0
14.8

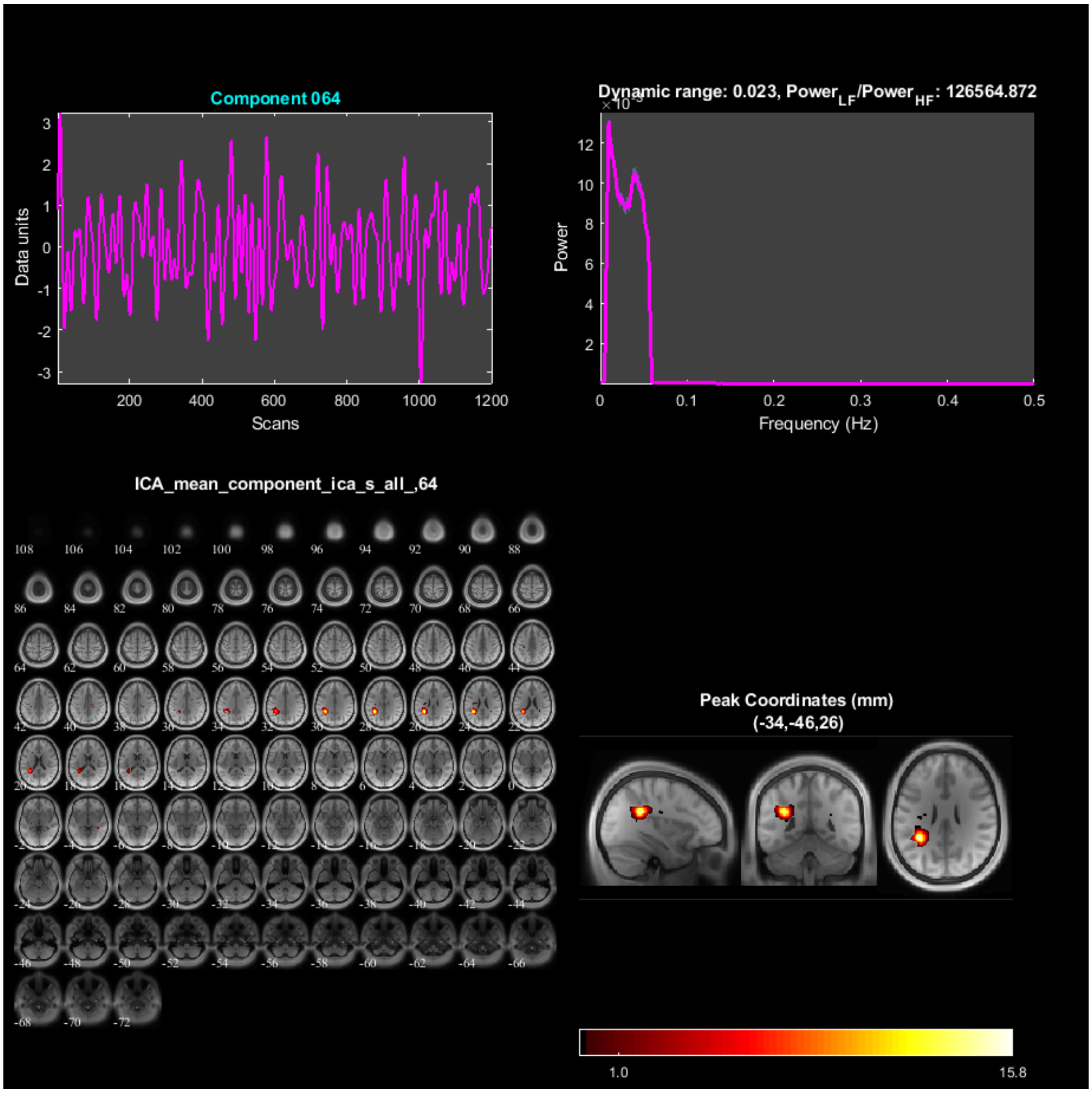
Component 064
Data units
Scans
Dynamic range: 0.023, Power_LF/Power_HF: 126564.872
Power
Frequency (Hz)
ICA_mean_component_ica_s_all_,64
Peak Coordinates (mm)
(-34,-46,26)
1.0
15.8

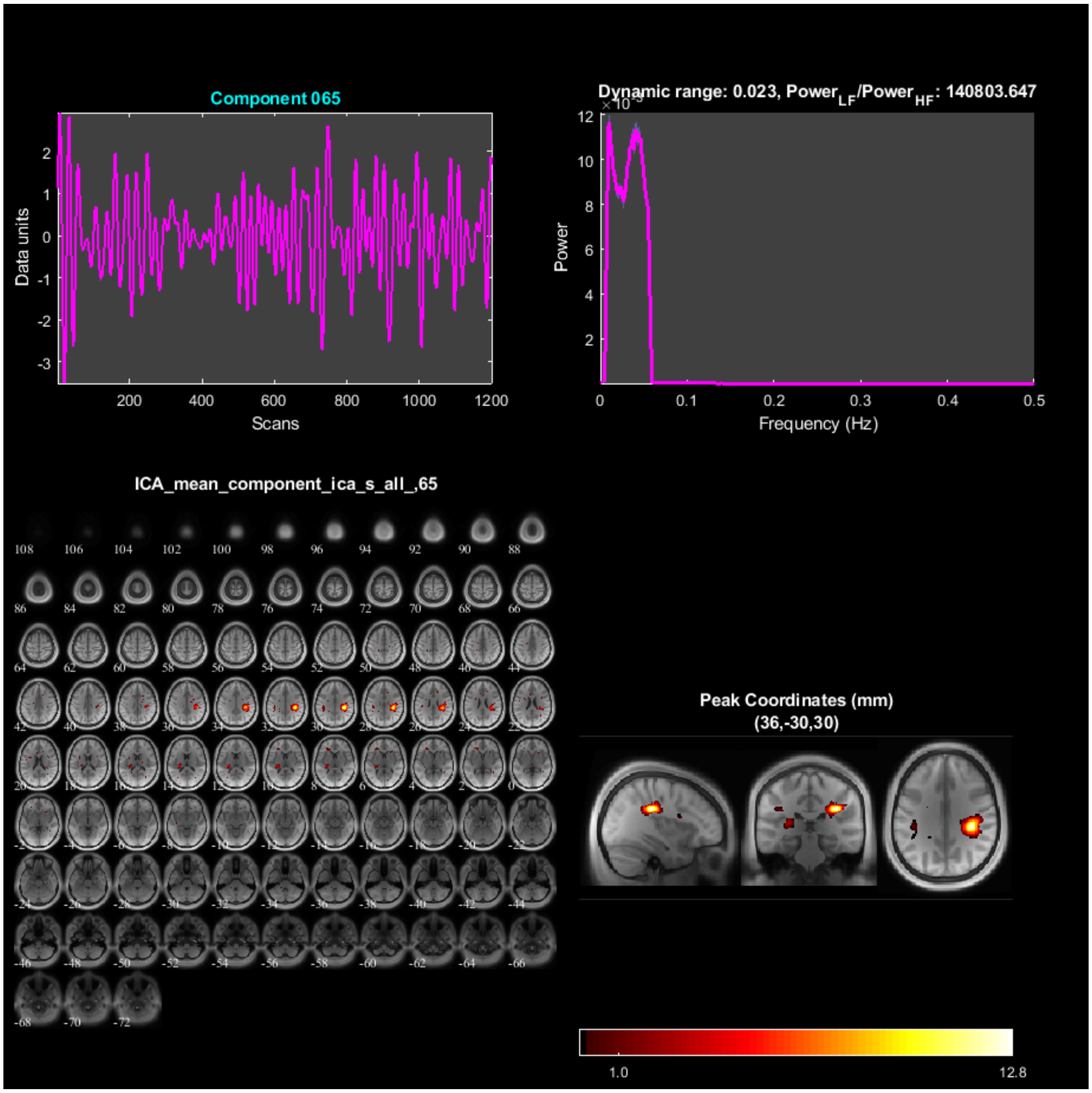

Component 065
Data units
Scans
Dynamic range: 0.023, Power$_{LF}$/Power$_{HF}$: 140803.647
Power
Frequency (Hz)
ICA_mean_component_ica_s_all_,65
Peak Coordinates (mm)
(36,-30,30)
1.0
12.8

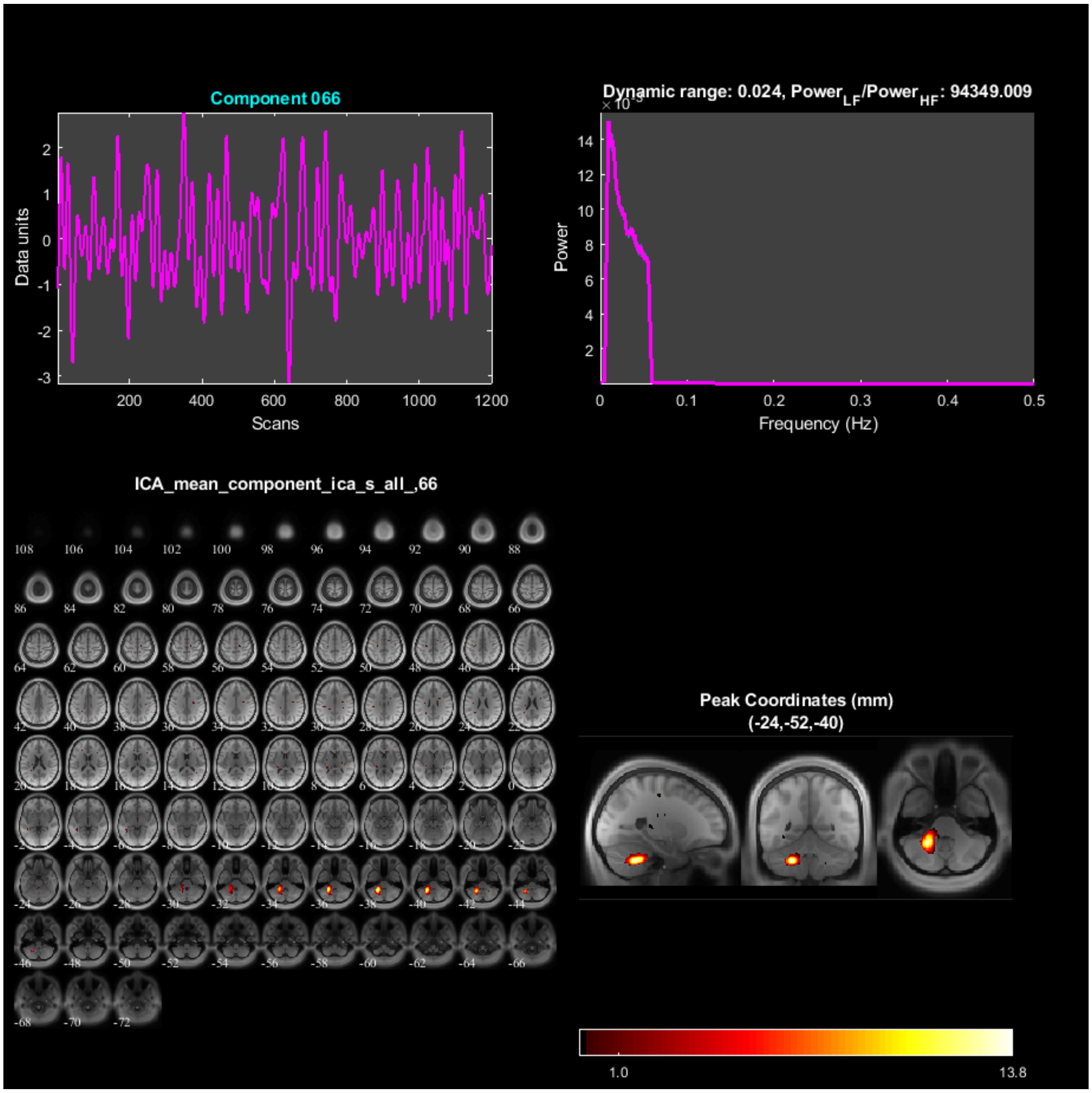

Component 066
Data units
Scans
Dynamic range: 0.024, Power$_{LF}$/Power$_{HF}$: 94349.009
Power
Frequency (Hz)
ICA_mean_component_ica_s_all_,66
Peak Coordinates (mm)
(-24,-52,-40)
1.0
13.8

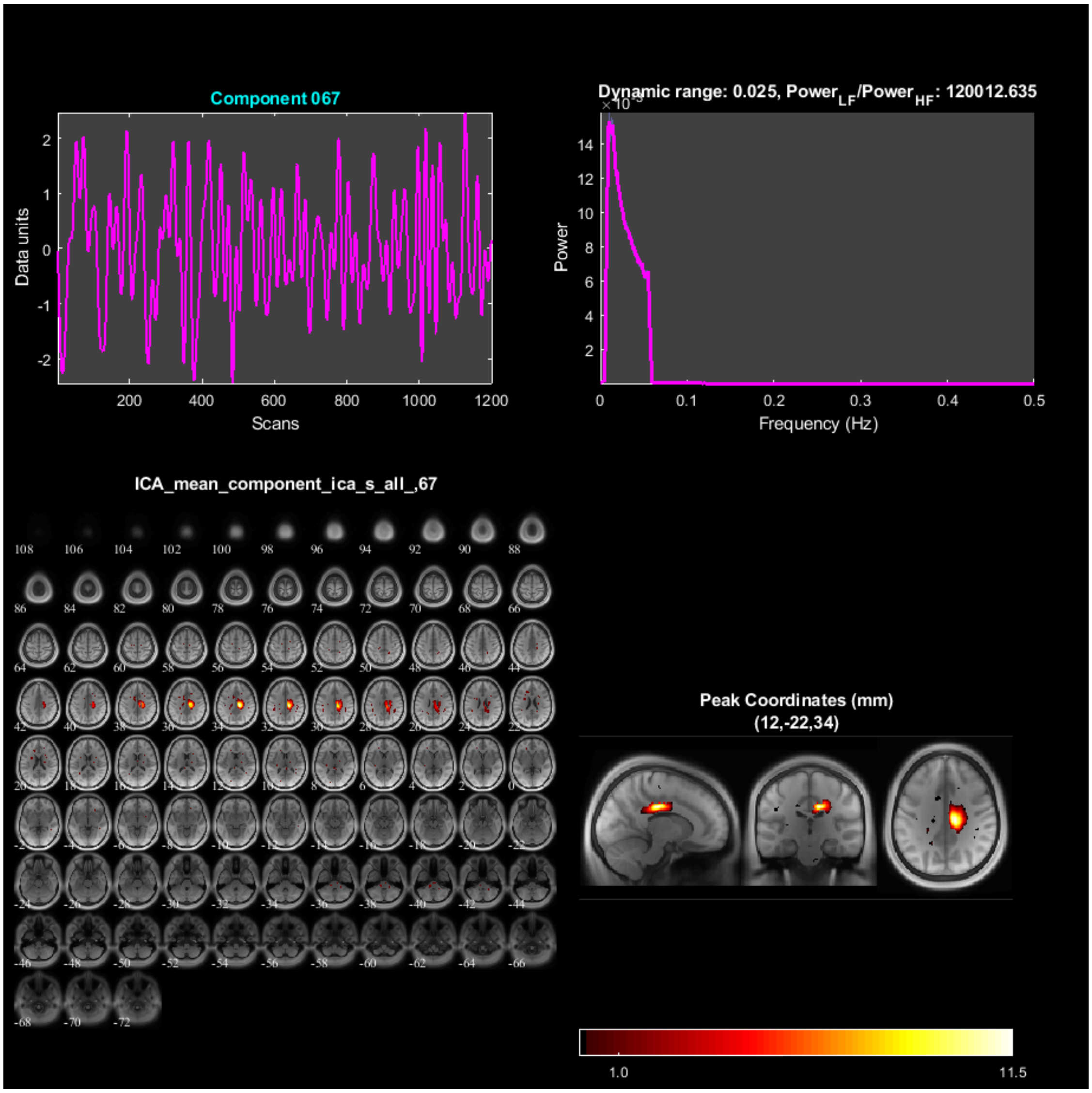

Component 067
Data units
Scans
Dynamic range: 0.025, Power$_{LF}$/Power$_{HF}$: 120012.635
Power
Frequency (Hz)
ICA_mean_component_ica_s_all_,67
Peak Coordinates (mm)
(12,-22,34)
1.0
11.5

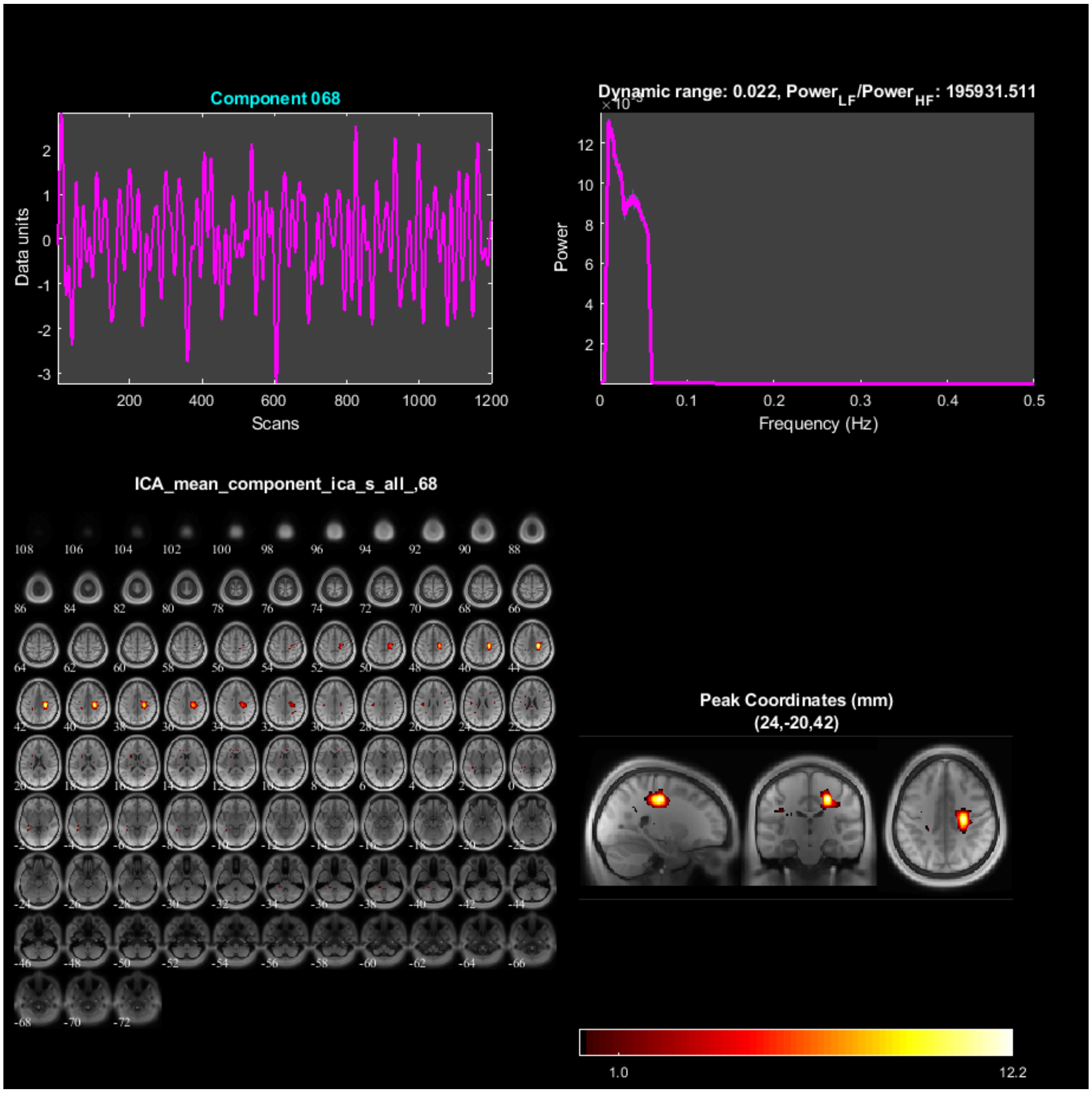

Component 068
Data units
Scans
Dynamic range: 0.022, Power$_{LF}$/Power$_{HF}$: 195931.511
Power
Frequency (Hz)
ICA_mean_component_ica_s_all_,68
Peak Coordinates (mm)
(24,-20,42)
1.0
12.2

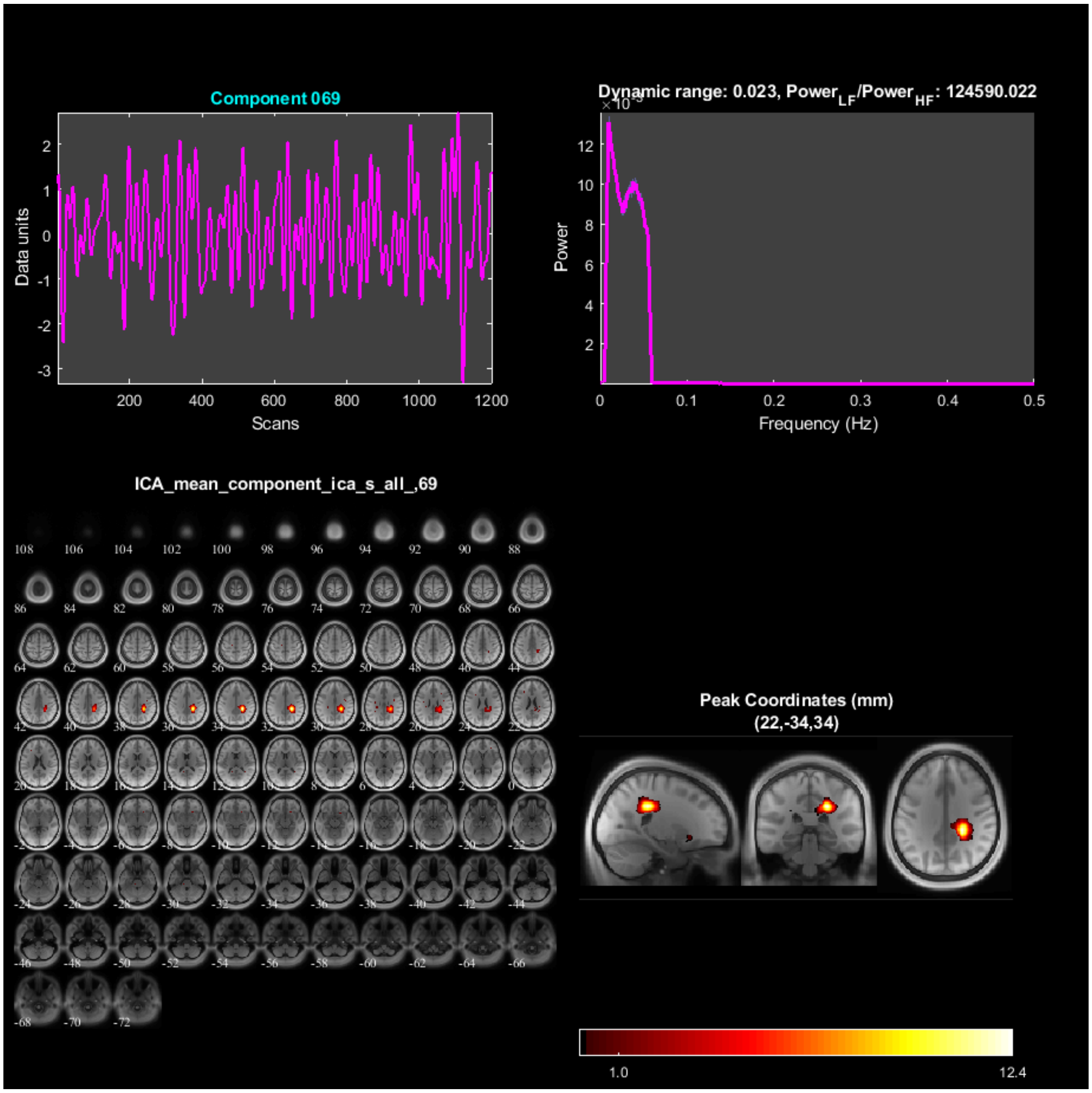
Component 069
Data units
Scans
Dynamic range: 0.023, Power_LF/Power_HF: 124590.022
Power
Frequency (Hz)
ICA_mean_component_ica_s_all_,69
Peak Coordinates (mm)
(22,-34,34)
1.0
12.4

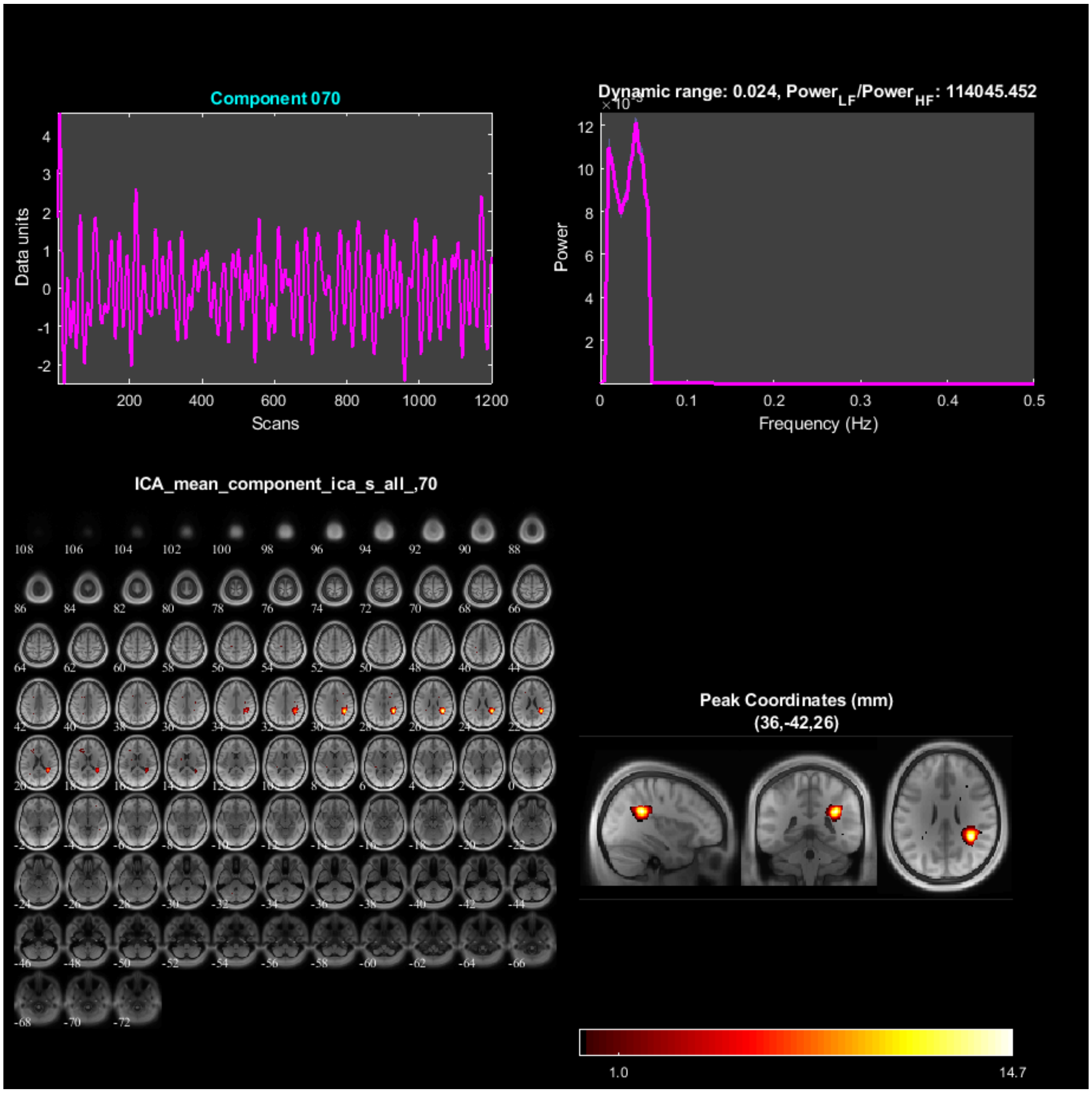
Component 070
Data units
Scans
Dynamic range: 0.024, Power$_{LF}$/Power$_{HF}$: 114045.452
Power
Frequency (Hz)
ICA_mean_component_ica_s_all_,70
Peak Coordinates (mm)
(36,-42,26)
1.0
14.7

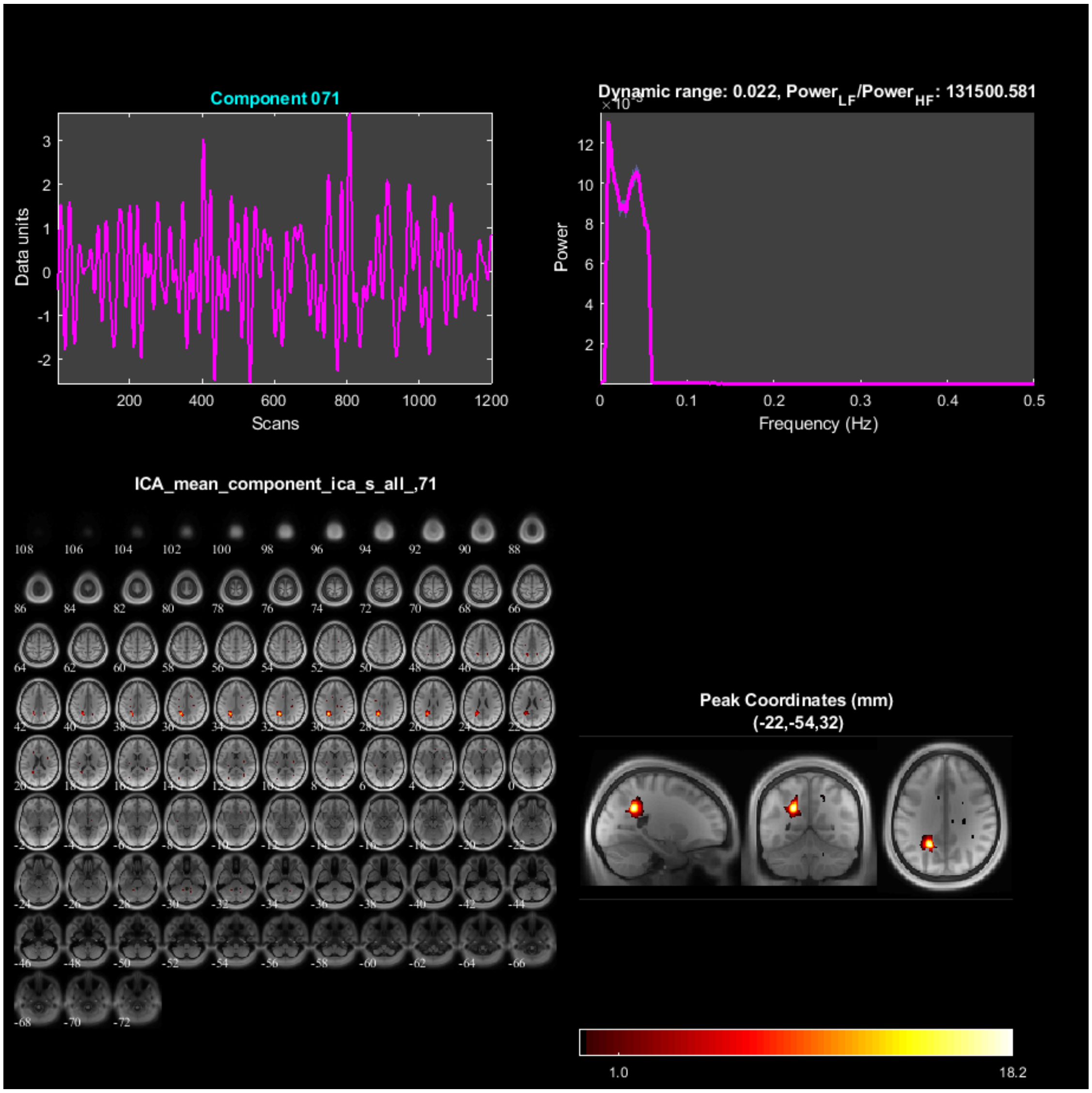

Component 071
Data units
Scans
Dynamic range: 0.022, Power$_{LF}$/Power$_{HF}$: 131500.581
Power
Frequency (Hz)
ICA_mean_component_ica_s_all_,71
Peak Coordinates (mm)
(-22,-54,32)
1.0
18.2

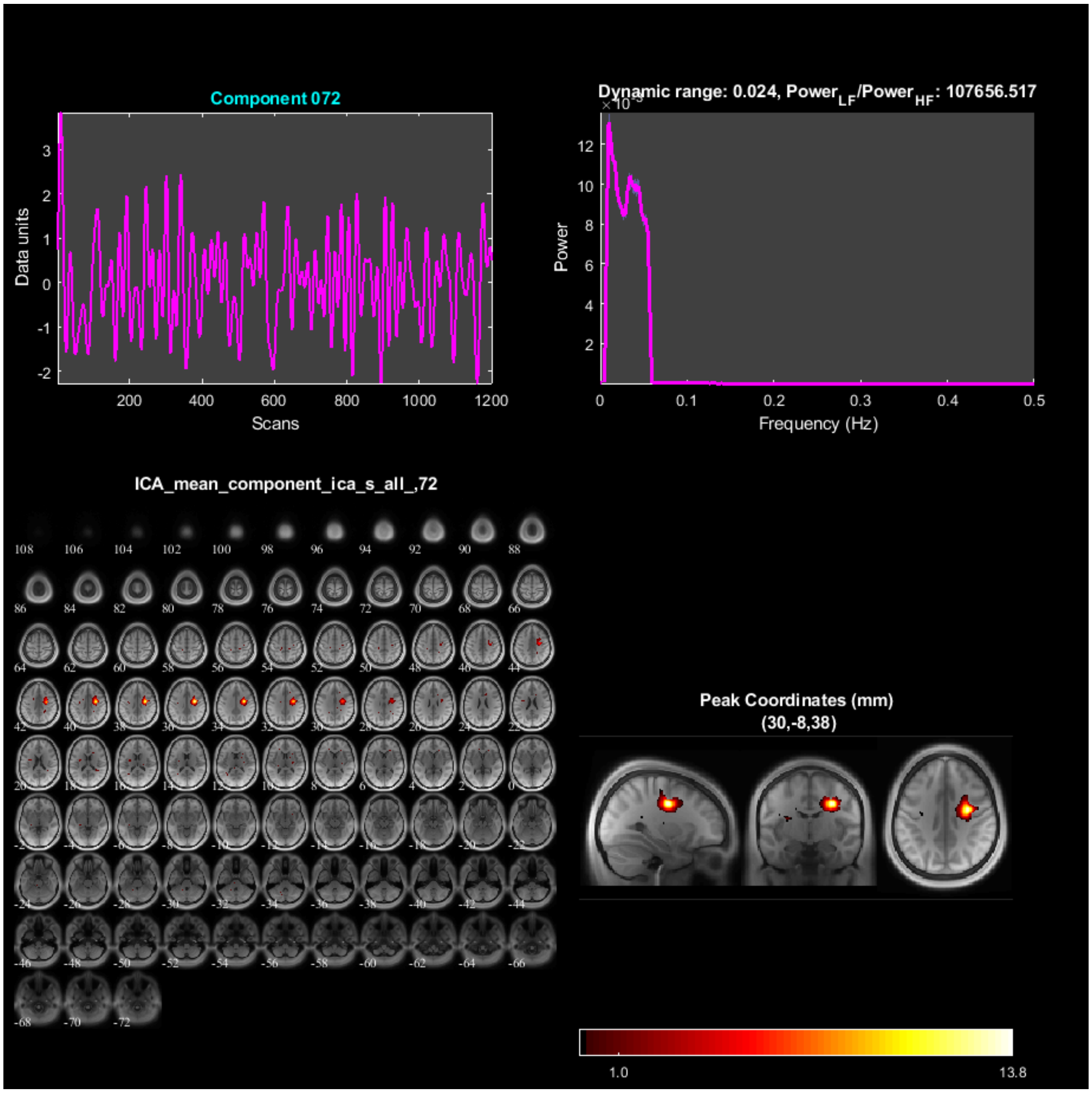

Component 072
Data units
Scans
Dynamic range: 0.024, Power$_{LF}$/Power$_{HF}$: 107656.517
Power
Frequency (Hz)
ICA_mean_component_ica_s_all_,72
Peak Coordinates (mm)
(30,-8,38)
1.0
13.8

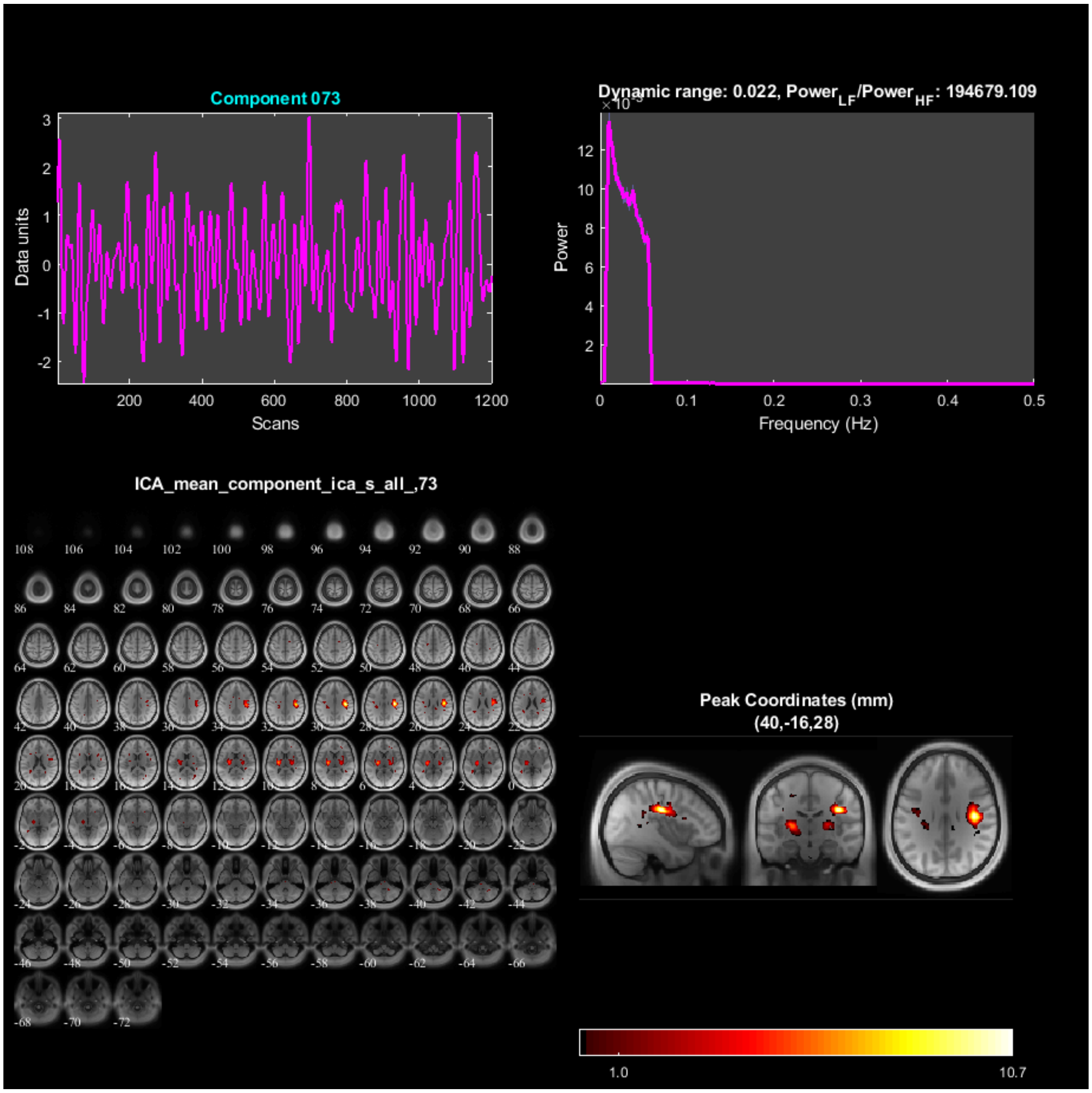

Component 073
Data units
Scans
Dynamic range: 0.022, Power$_{LF}$/Power$_{HF}$: 194679.109
Power
Frequency (Hz)
ICA_mean_component_ica_s_all_,73
Peak Coordinates (mm)
(40,-16,28)
1.0
10.7

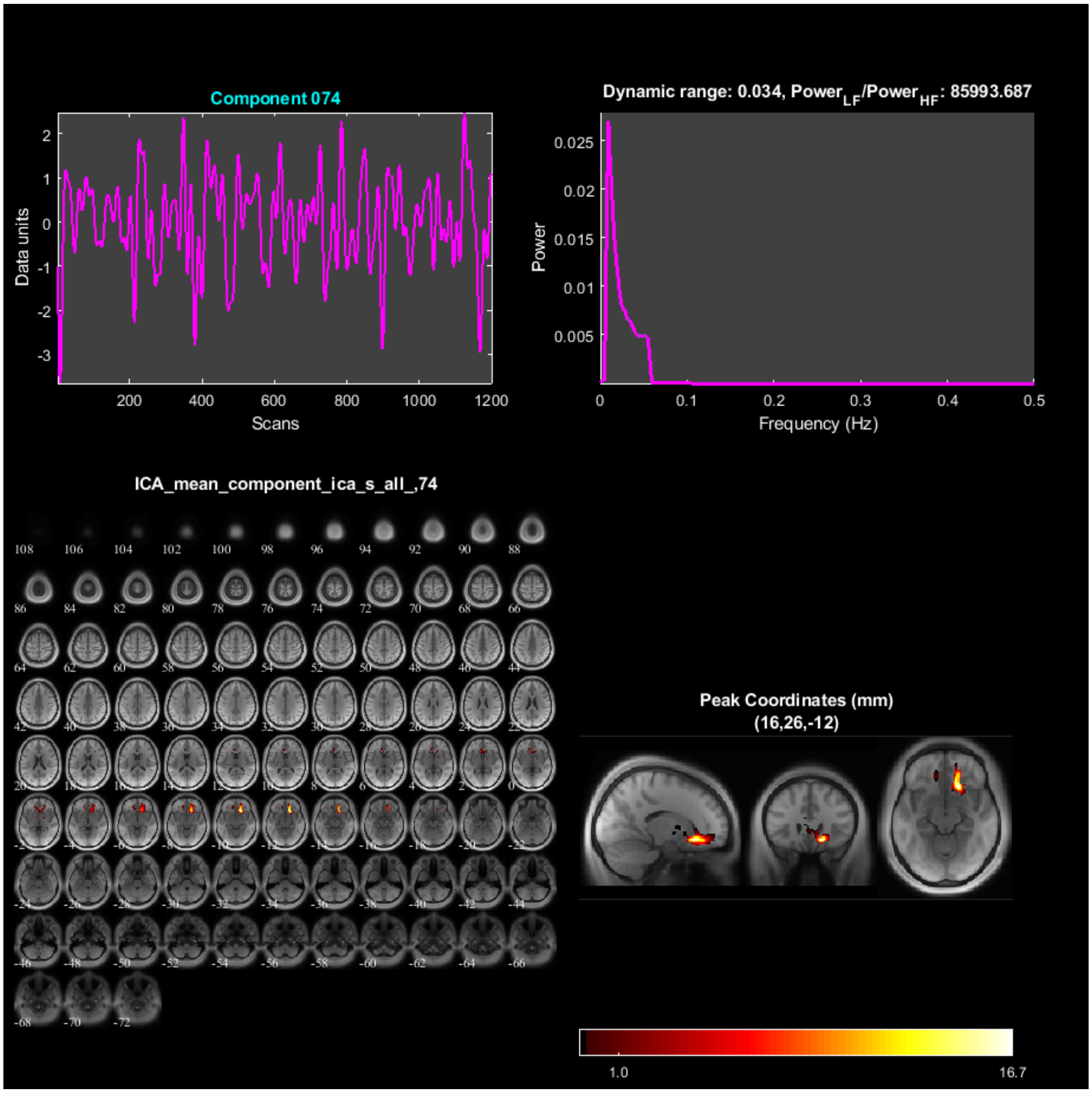

Component 074
Data units
Scans
Dynamic range: 0.034, Power$_{LF}$/Power$_{HF}$: 85993.687
Power
Frequency (Hz)
ICA_mean_component_ica_s_all_,74
Peak Coordinates (mm)
(16,26,-12)
1.0
16.7

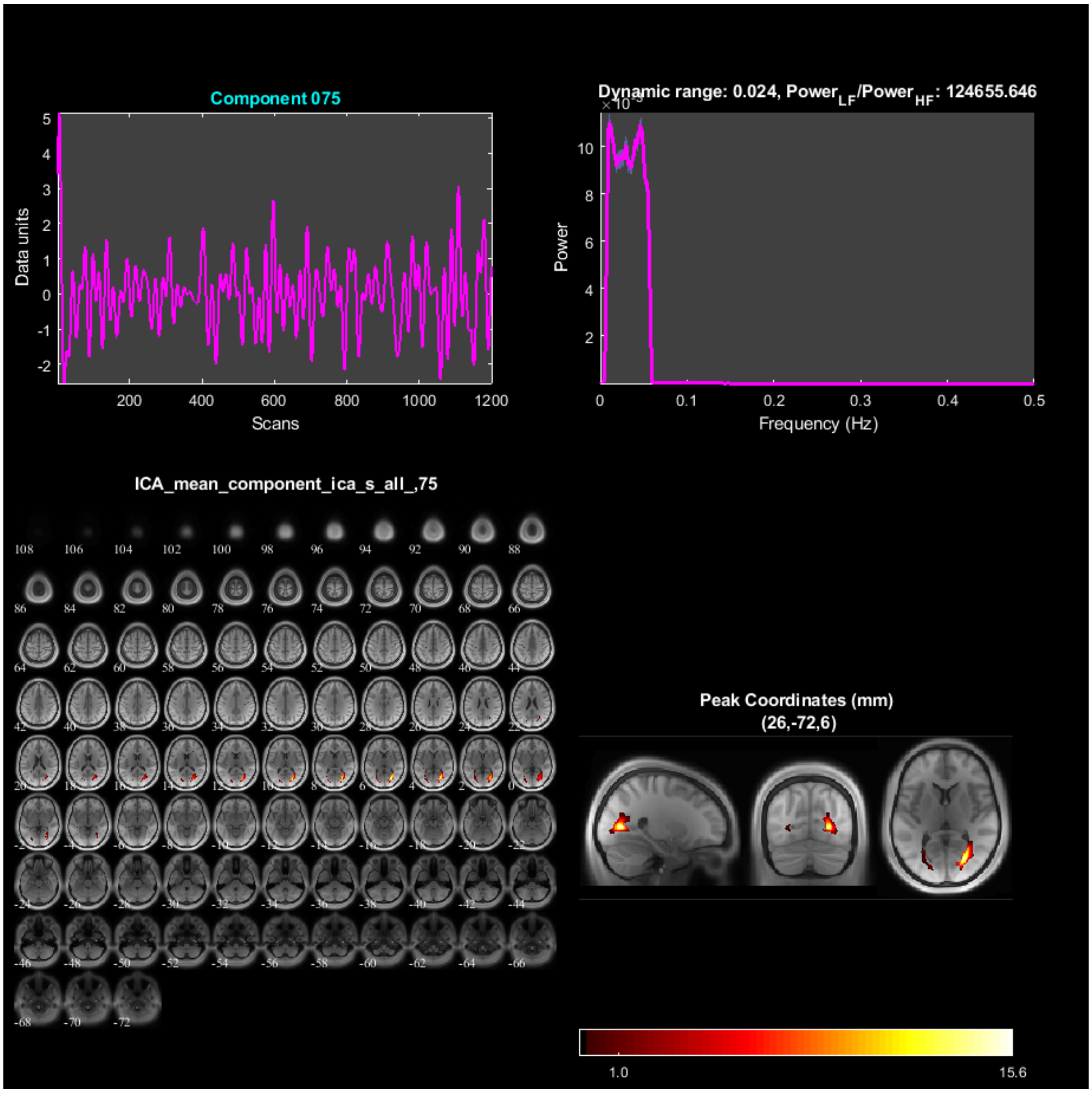
Component 075
Data units
Scans
Dynamic range: 0.024, Power_LF/Power_HF: 124655.646
Power
Frequency (Hz)
ICA_mean_component_ica_s_all_,75
Peak Coordinates (mm)
(26,-72,6)
1.0
15.6

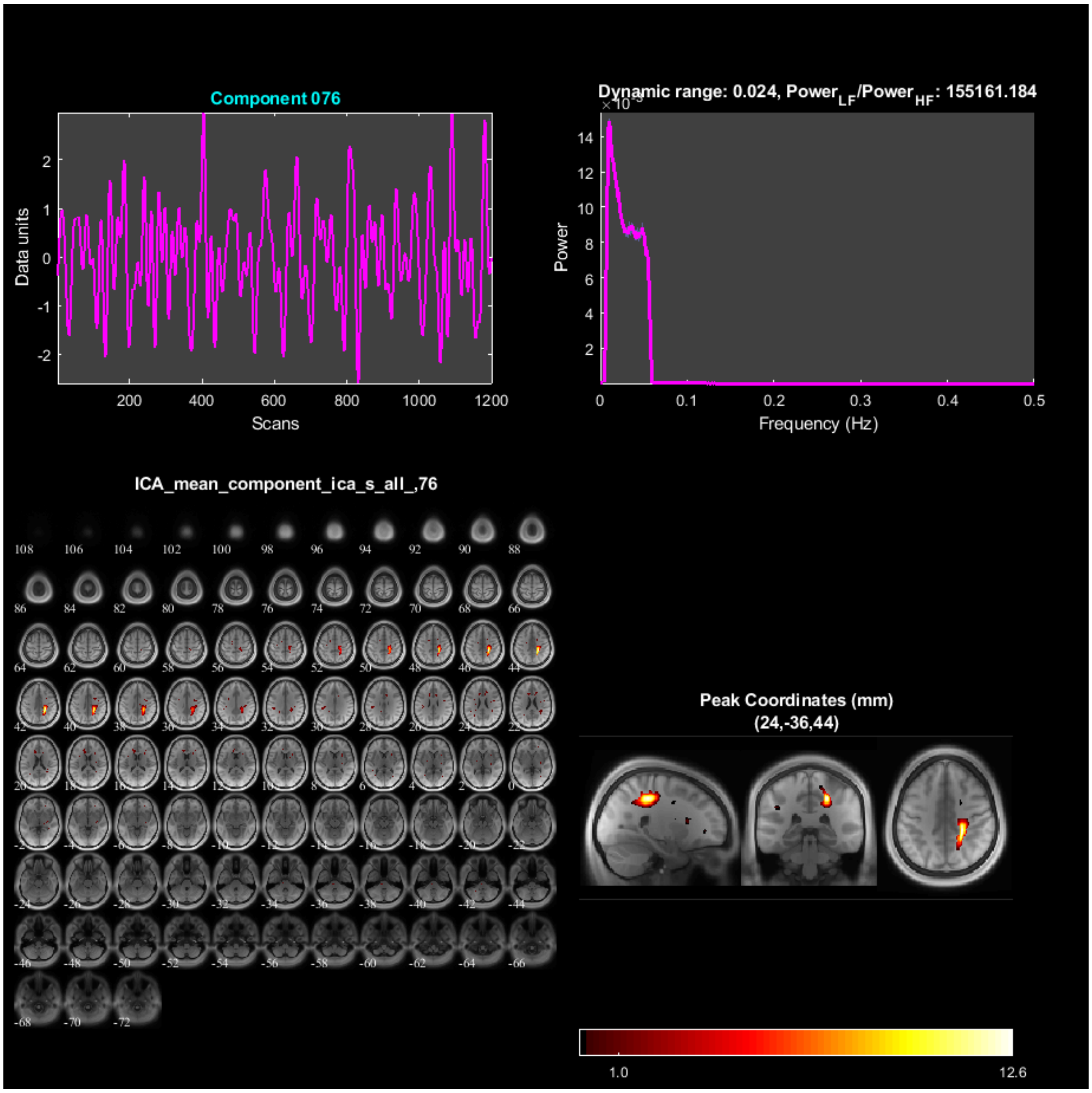
Component 076
Data units
Scans
Dynamic range: 0.024, Power_LF/Power_HF: 155161.184
Power
Frequency (Hz)
ICA_mean_component_ica_s_all_,76
Peak Coordinates (mm)
(24,-36,44)
1.0
12.6

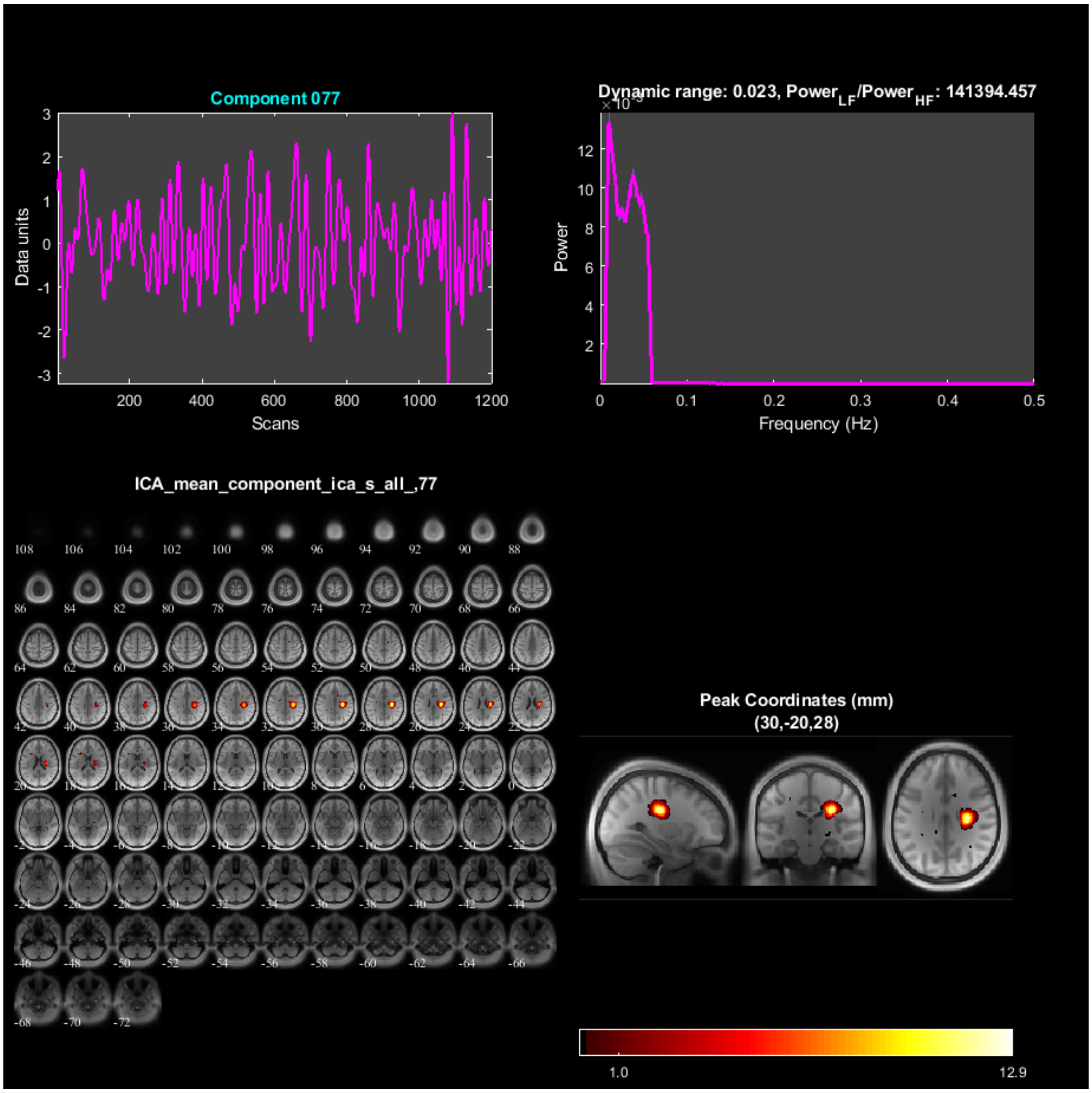
Component 077
Data units
Scans
Dynamic range: 0.023, Power$_{LF}$/Power$_{HF}$: 141394.457
Power
Frequency (Hz)
ICA_mean_component_ica_s_all_,77
Peak Coordinates (mm)
(30,-20,28)
1.0
12.9

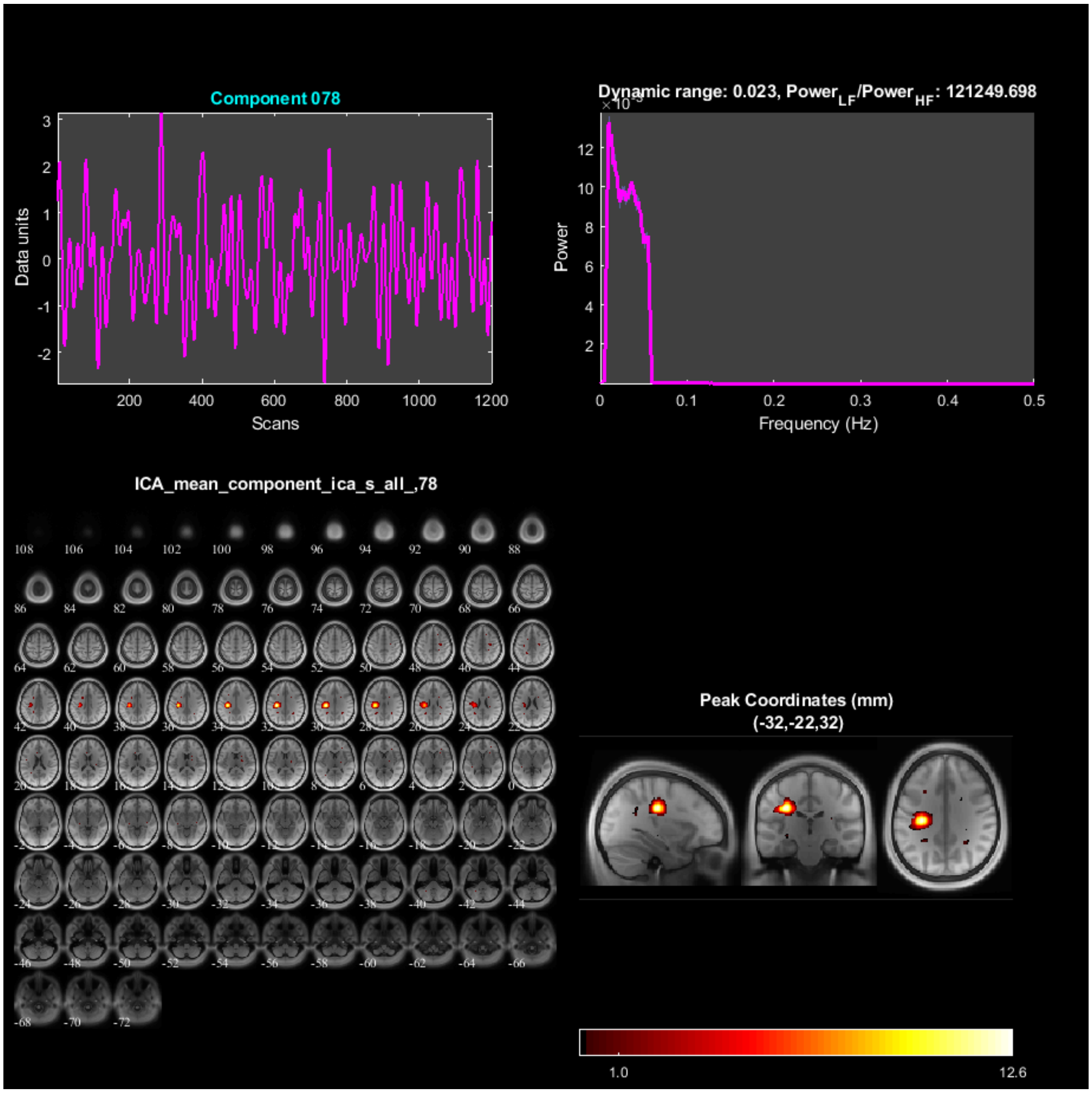
Component 078
Data units
Scans
Dynamic range: 0.023, Power$_{LF}$/Power$_{HF}$: 121249.698
Power
Frequency (Hz)
ICA_mean_component_ica_s_all_,78
Peak Coordinates (mm)
(-32,-22,32)
1.0
12.6

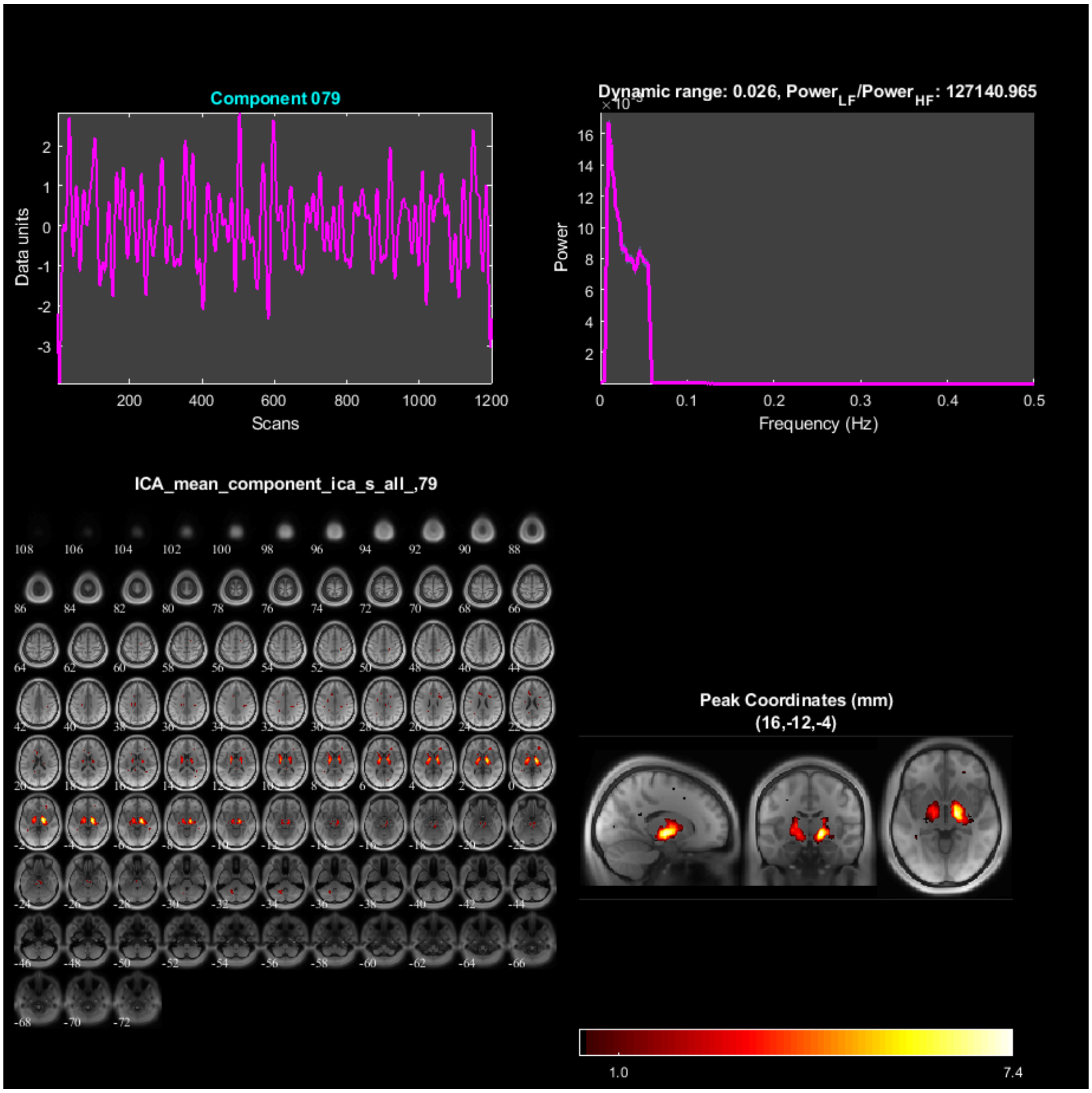
Component 079
Data units
Scans
Dynamic range: 0.026, Power$_{LF}$/Power$_{HF}$: 127140.965
Power
Frequency (Hz)
ICA_mean_component_ica_s_all_,79
Peak Coordinates (mm)
(16,-12,-4)
1.0
7.4

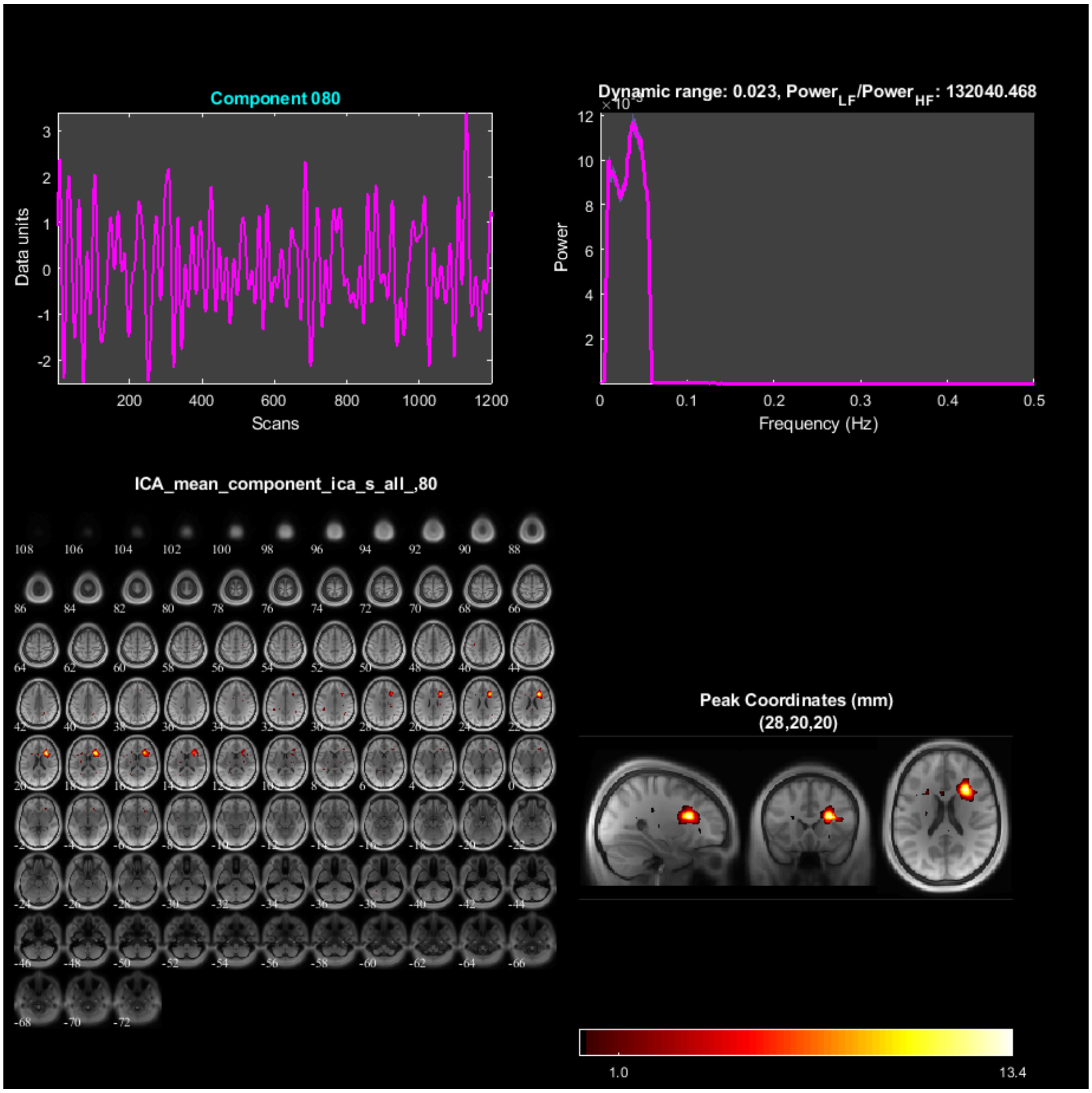

### Spectral Summary

- **a) dynamic_range** - Difference between the peak power and minimum power at frequencies to the right of the peak.
- **b) fALFF** - Low frequency to high frequency power ratio.

| *ComponentNumber* | *DynamicRange* | *fALFF* |
|---|---|---|
| *1* | *0.022669* | *1.4199e+05* |
| *2* | *0.024073* | *1.7905e+05* |
| *3* | *0.023438* | *1.1206e+05* |
| *4* | *0.023048* | *1.3151e+05* |
| *5* | *0.025145* | *1.1582e+05* |
| *6* | *0.024142* | *1.577e+05* |
| *7* | *0.022825* | *1.5256e+05* |

| | | |
|---|---|---|
| 8 | 0.023648 | 1.2581e+05 |
| 9 | 0.024073 | 1.3674e+05 |
| 10 | 0.025153 | 1.5264e+05 |
| 11 | 0.023199 | 1.261e+05 |
| 12 | 0.026635 | 1.3939e+05 |
| 13 | 0.022334 | 1.2631e+05 |
| 14 | 0.023867 | 85024 |
| 15 | 0.023664 | 1.24e+05 |
| 16 | 0.024845 | 1.556e+05 |
| 17 | 0.023849 | 93817 |
| 18 | 0.023909 | 99988 |
| 19 | 0.022929 | 1.265e+05 |
| 20 | 0.023553 | 1.2345e+05 |
| 21 | 0.02337 | 1.2538e+05 |
| 22 | 0.022981 | 1.6482e+05 |
| 23 | 0.022498 | 1.299e+05 |
| 24 | 0.025019 | 1.4977e+05 |
| 25 | 0.023481 | 1.0056e+05 |
| 26 | 0.023803 | 1.4004e+05 |
| 27 | 0.023019 | 1.3195e+05 |
| 28 | 0.023268 | 1.3049e+05 |
| 29 | 0.025571 | 1.4399e+05 |
| 30 | 0.023064 | 1.1948e+05 |
| 31 | 0.024062 | 1.1594e+05 |
| 32 | 0.022007 | 1.4477e+05 |
| 33 | 0.026787 | 1.1799e+05 |
| 34 | 0.022638 | 1.1525e+05 |
| 35 | 0.022942 | 1.1714e+05 |
| 36 | 0.023278 | 1.3563e+05 |
| 37 | 0.023381 | 1.8152e+05 |
| 38 | 0.022754 | 1.0718e+05 |
| 39 | 0.022506 | 1.3302e+05 |
| 40 | 0.02266 | 1.0794e+05 |
| 41 | 0.02368 | 1.1055e+05 |
| 42 | 0.022637 | 1.4597e+05 |
| 43 | 0.023216 | 1.3524e+05 |
| 44 | 0.02344 | 1.1783e+05 |
| 45 | 0.022853 | 1.3868e+05 |
| 46 | 0.02376 | 1.0897e+05 |
| 47 | 0.023402 | 1.0007e+05 |
| 48 | 0.022798 | 1.1048e+05 |
| 49 | 0.022074 | 1.7704e+05 |
| 50 | 0.023332 | 1.5803e+05 |
| 51 | 0.02289 | 1.7044e+05 |
| 52 | 0.025932 | 88328 |
| 53 | 0.023167 | 1.0651e+05 |
| 54 | 0.025325 | 1.2663e+05 |
| 55 | 0.024065 | 86694 |
| 56 | 0.025038 | 1.1877e+05 |
| 57 | 0.02402 | 1.1801e+05 |
| 58 | 0.023678 | 1.0915e+05 |
| 59 | 0.022817 | 1.6662e+05 |
| 60 | 0.022846 | 1.3227e+05 |
| 61 | 0.023363 | 1.3877e+05 |

| | | |
|---|---|---|
| 62 | 0.027258 | 2.6699e+05 |
| 63 | 0.022249 | 1.2743e+05 |
| 64 | 0.022959 | 1.2656e+05 |
| 65 | 0.023344 | 1.408e+05 |
| 66 | 0.023658 | 94349 |
| 67 | 0.02458 | 1.2001e+05 |
| 68 | 0.022109 | 1.9593e+05 |
| 69 | 0.022521 | 1.2459e+05 |
| 70 | 0.02363 | 1.1405e+05 |
| 71 | 0.022485 | 1.315e+05 |
| 72 | 0.023535 | 1.0766e+05 |
| 73 | 0.022119 | 1.9468e+05 |
| 74 | 0.034265 | 85994 |
| 75 | 0.023594 | 1.2466e+05 |
| 76 | 0.02374 | 1.5516e+05 |
| 77 | 0.023396 | 1.4139e+05 |
| 78 | 0.022966 | 1.2125e+05 |
| 79 | 0.025539 | 1.2714e+05 |
| 80 | 0.022712 | 1.3204e+05 |

### Temporal Stats On Beta Weights

Multiple regression is done using the timecourses from SPM design matrix as model and ICA timecourses as observations. R^2 values for each component are shown in bar plot. For each component, one sample t-test results of each session and condition are shown in the bar plots.

### Kurtosis of timecourses and spatial maps

Mean across subjects is reported in table. Figure shows mean+/- SEM across subjects

| ComponentNumber | Timecourses | SpatialMaps |
|---|---|---|
| 1 | 3.663 | 3.4307 |
| 2 | 3.1569 | 3.5067 |
| 3 | 3.5292 | 3.4349 |
| 4 | 3.4946 | 3.4424 |
| 5 | 3.1789 | 3.377 |
| 6 | 3.554 | 3.497 |
| 7 | 3.6048 | 3.4144 |
| 8 | 3.605 | 3.5763 |
| 9 | 3.6483 | 3.4499 |
| 10 | 3.3984 | 3.4542 |
| 11 | 3.3707 | 3.4675 |
| 12 | 3.8523 | 3.4137 |
| 13 | 3.43 | 3.5541 |
| 14 | 3.2882 | 3.4772 |
| 15 | 3.4997 | 3.3797 |
| 16 | 3.0602 | 3.4935 |
| 17 | 3.3783 | 3.4441 |
| 18 | 3.9579 | 3.4768 |

| | | |
|---|---|---|
| 19 | 3.4022 | 3.3986 |
| 20 | 3.3364 | 3.2722 |
| 21 | 3.3393 | 3.626 |
| 22 | 3.6117 | 3.6944 |
| 23 | 3.4666 | 3.4502 |
| 24 | 3.2488 | 3.4752 |
| 25 | 3.1839 | 3.4319 |
| 26 | 3.4484 | 3.3978 |
| 27 | 3.4302 | 3.5544 |
| 28 | 3.729 | 3.4449 |
| 29 | 3.5453 | 3.3889 |
| 30 | 3.7752 | 3.3763 |
| 31 | 3.6714 | 3.4575 |
| 32 | 3.2658 | 3.5714 |
| 33 | 3.2711 | 4.5667 |
| 34 | 3.2602 | 3.912 |
| 35 | 3.4833 | 3.5402 |
| 36 | 3.5844 | 3.441 |
| 37 | 3.2824 | 3.3999 |
| 38 | 3.3158 | 3.6316 |
| 39 | 3.2623 | 3.3957 |
| 40 | 3.3334 | 3.488 |
| 41 | 3.8948 | 3.7955 |
| 42 | 3.4288 | 3.5031 |
| 43 | 3.48 | 3.361 |
| 44 | 3.6697 | 3.4749 |
| 45 | 3.3675 | 3.525 |
| 46 | 3.1929 | 3.6086 |
| 47 | 3.9813 | 3.4406 |
| 48 | 3.2998 | 3.5091 |
| 49 | 3.3402 | 3.583 |
| 50 | 3.3914 | 3.7244 |
| 51 | 3.3805 | 3.4345 |
| 52 | 3.1738 | 3.5485 |
| 53 | 3.5525 | 3.3782 |
| 54 | 3.2283 | 3.3943 |
| 55 | 3.3798 | 3.3981 |
| 56 | 3.362 | 3.3059 |
| 57 | 3.3965 | 3.4149 |
| 58 | 3.5036 | 3.4724 |
| 59 | 3.2331 | 3.679 |
| 60 | 3.7541 | 3.2883 |
| 61 | 3.3134 | 3.5134 |
| 62 | 3.5937 | 3.3812 |
| 63 | 3.4876 | 3.4763 |
| 64 | 3.2151 | 3.6864 |
| 65 | 3.3004 | 3.4641 |
| 66 | 3.9347 | 3.5607 |
| 67 | 3.4212 | 3.3988 |
| 68 | 3.7793 | 3.4598 |
| 69 | 3.5396 | 3.4869 |
| 70 | 3.3612 | 3.6538 |
| 71 | 3.3521 | 3.773 |
| 72 | 3.729 | 3.4882 |

| | | |
|---|---|---|
| *73* | *3.6407* | *3.3116* |
| *74* | *3.5644* | *4.0216* |
| *75* | *3.3152* | *3.8072* |
| *76* | *3.6313* | *3.4859* |
| *77* | *3.6449* | *3.4719* |
| *78* | *3.2705* | *3.4915* |
| *79* | *3.826* | *3.2579* |
| *80* | *3.1483* | *3.4851* |

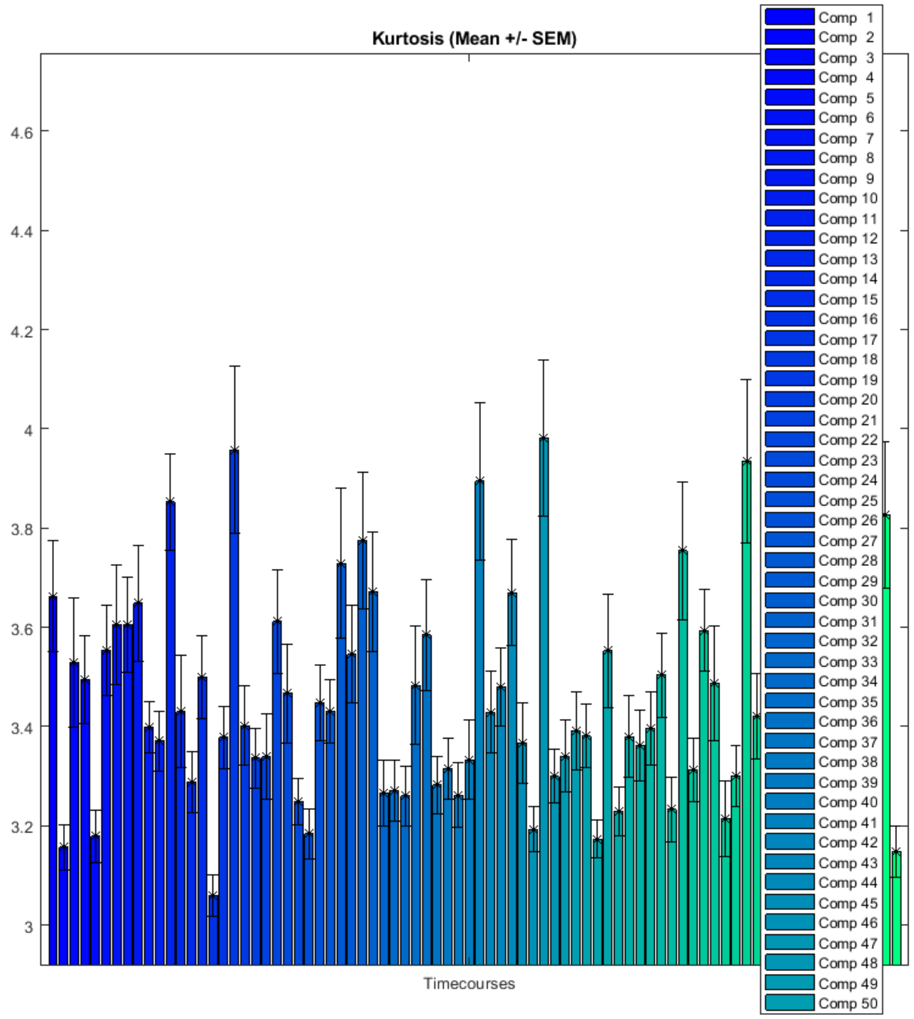

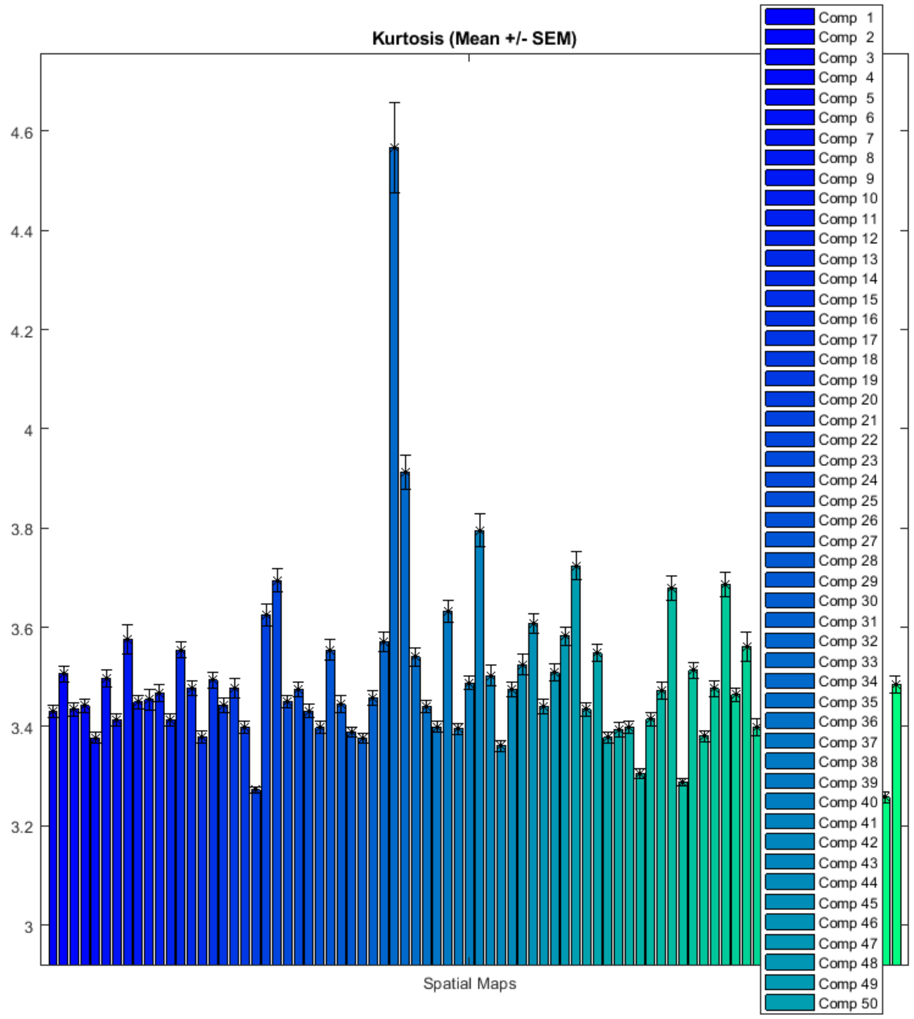

### FNC correlations

Functional network connectivity correlations are computed for each data-set and averaged across sessions.

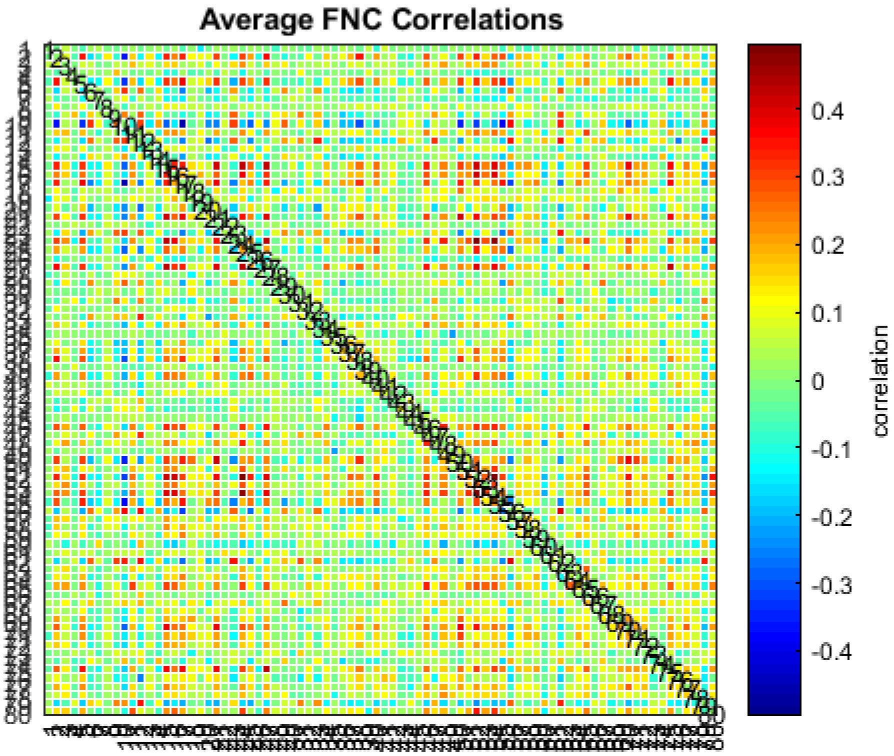

### FNC metrics of component spatial maps

Mutual information is computed between components spatially and averaged across data-sets.

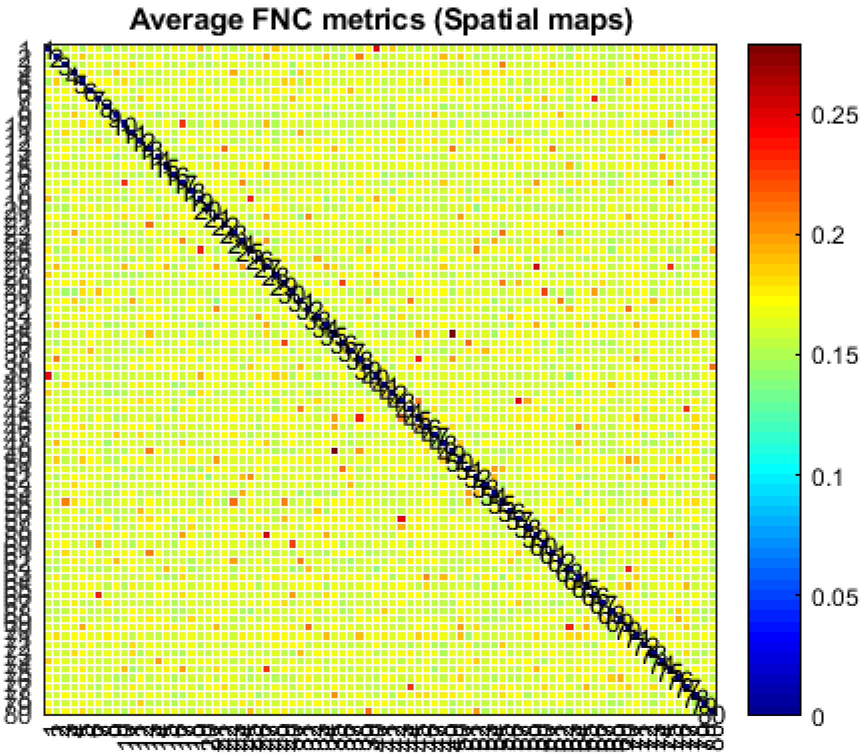

*Published with MATLAB® R2017b*
