## Supplementary material for "Power spectra reveal distinct BOLD resting‐state time courses in white matter": support file 2

Supporting file 2, page 1/4. Each figure shown below is the power spectra of the voxels within one WM IC. Each line represent the mean power spectra over 199 subjects at the same voxel. Each power spectral has been normalized to unit variance (z-score).

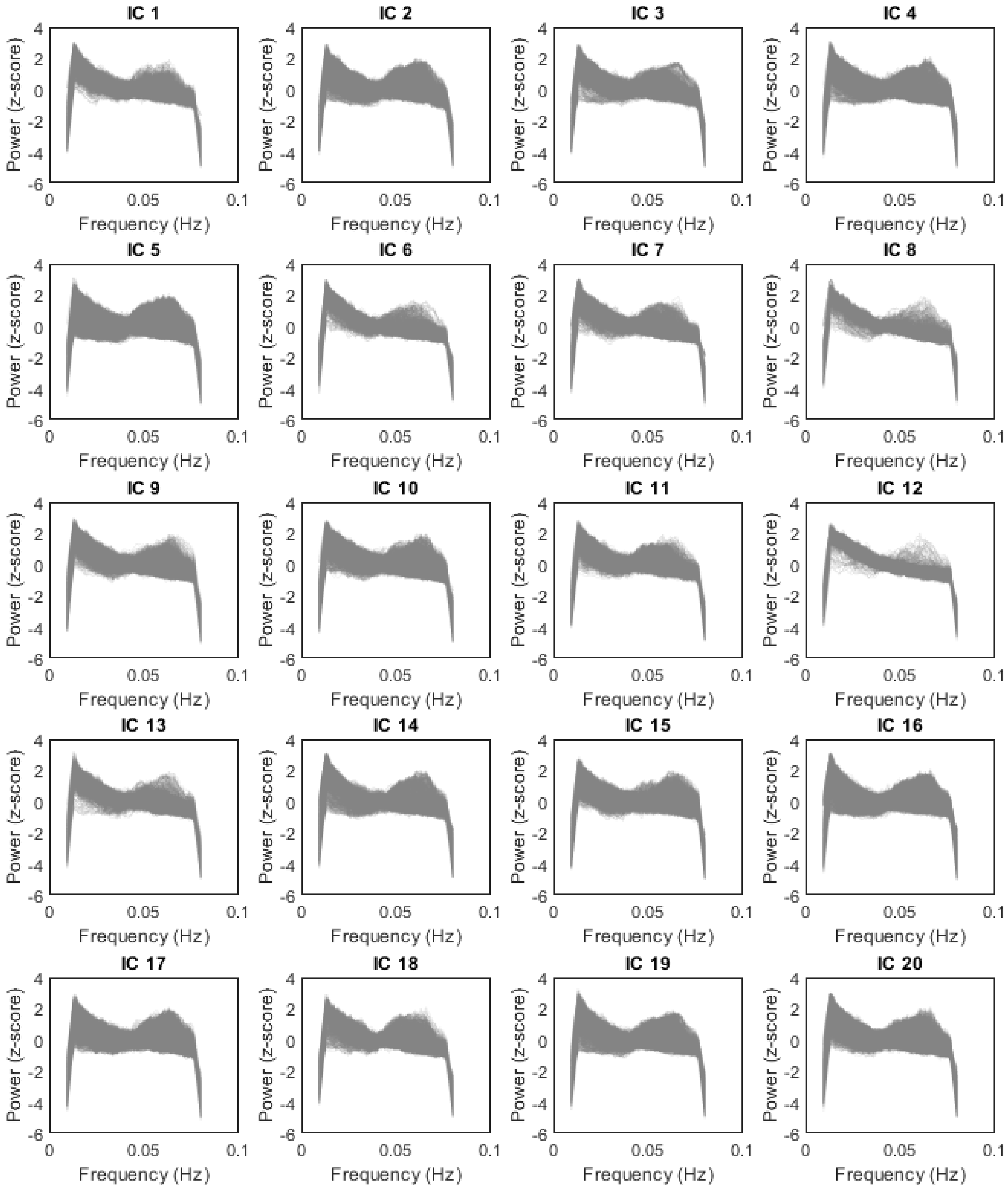

Supporting file 2, page 2/4. Each figure shown below is the power spectra of the voxels within one WM IC. Each line represent the mean power spectra over 199 subjects at the same voxel. Each power spectral has been normalized to unit variance (z-score).

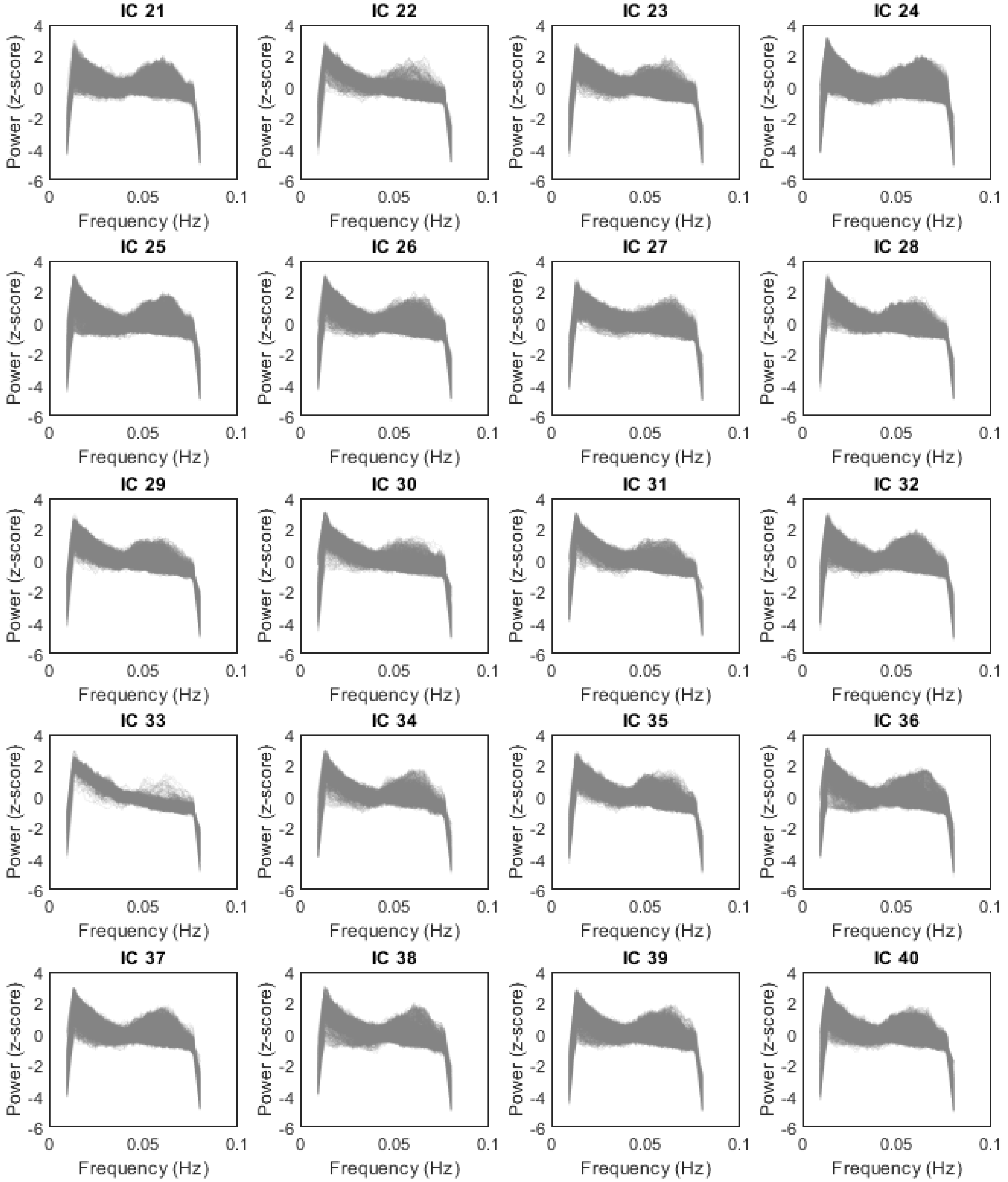

Supporting file 2, page 3/4. Each figure shown below is the power spectra of the voxels within one WM IC. Each line represent the mean power spectra over 199 subjects at the same voxel. Each power spectral has been normalized to unit variance (z-score).

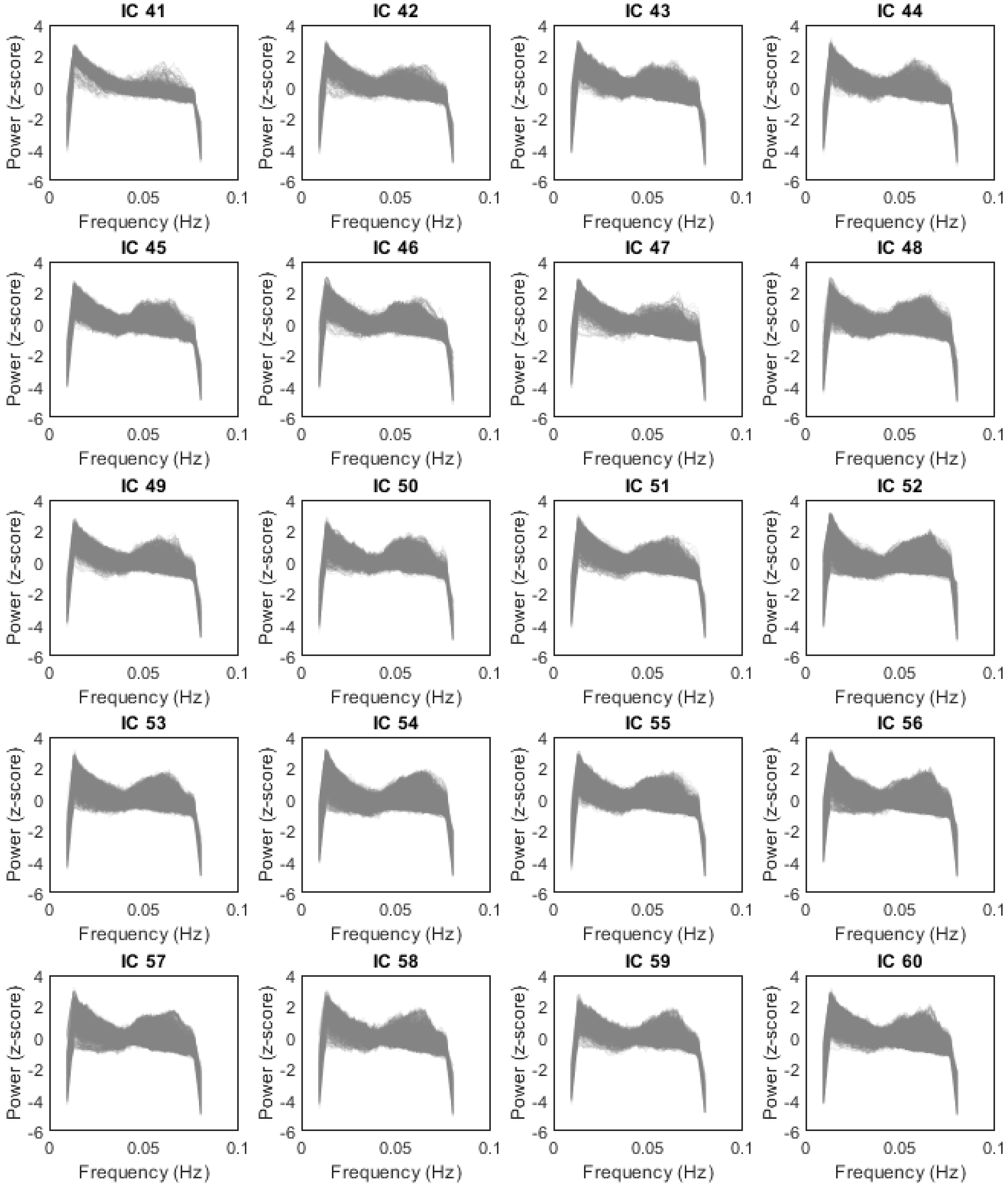

Supporting file 2, page 4/4. Each figure shown below is the power spectra of the voxels within one WM IC. Each line represent the mean power spectra over 199 subjects at the same voxel. Each power spectral has been normalized to unit variance (z-score).

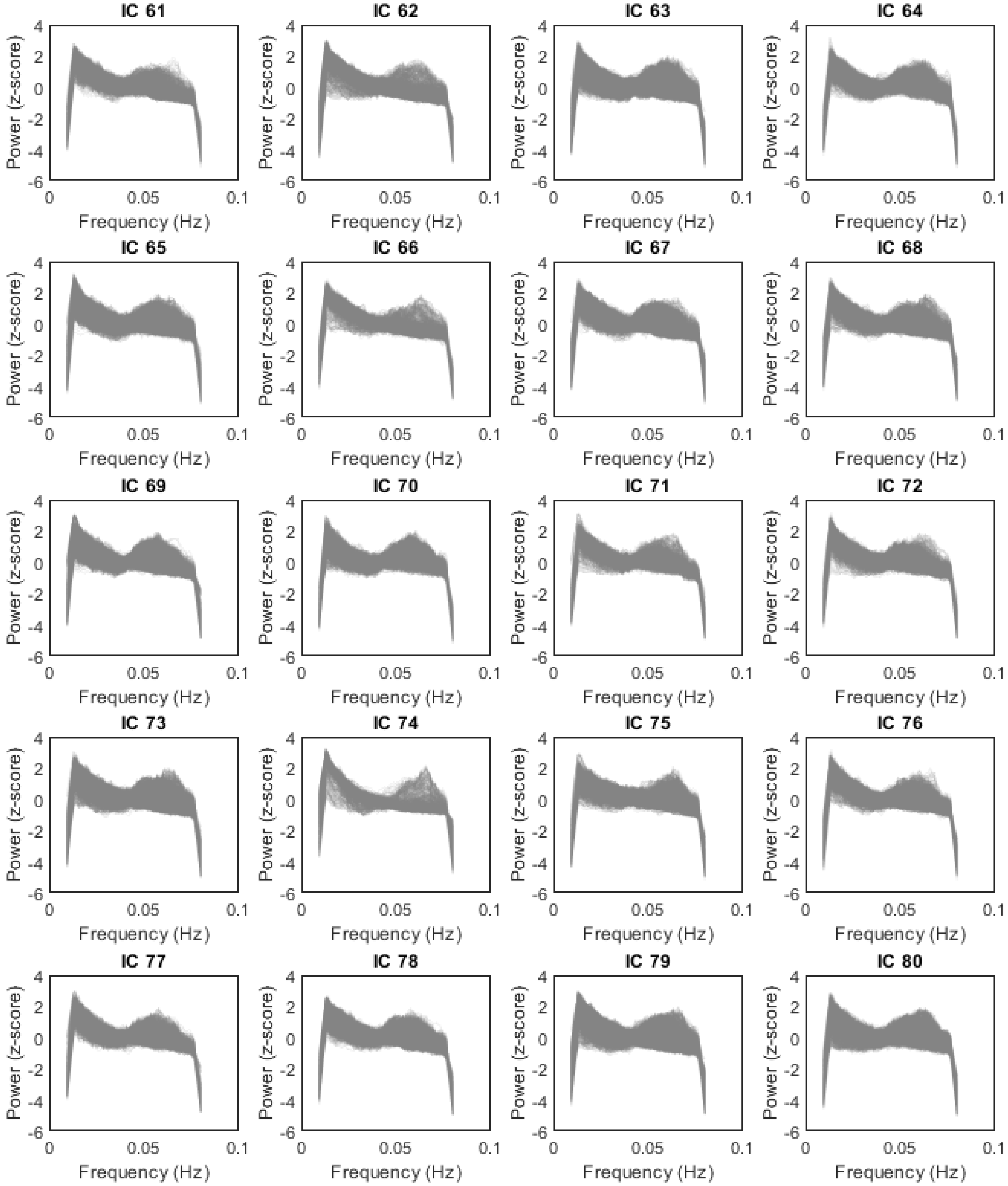
