## Supplementary material for "Power spectra reveal distinct BOLD resting‐state time courses in white matter": support file 3

### Group ICA Parameters

```
...................................................

Number of Subjects : 199
Number of Sessions : 1
Number of Independent Components : 80
ICA Algorithm : Infomax
Number Of Scans/Timepoints : 1200
Mask File : ICA_mask2_gm
Data Pre-processing Type : Remove Mean Per Timepoint
PCA Type : Standard
Group PCA Type : Subject Specific
Group ICA Type : Spatial
Back Reconstruction Type : GICA
Scaling Components : Z-scores
Stability analysis type : none
Group analysis mode: Serial
Anatomical file: H:\ICA_HCP_20200909\wm_ICs\T1_avg200.nii,1
Slice Plane: Axial
Image values: Positive
Convert to Z-scores: yes
Threshold: 1

...................................................
```

### ICASSO Plots

### Mean Components

Mean across all subjects and sessions is computed for each component

- **a) Timecourse** - Mean timecourse is converted to z-scores.
- **b) Spectra** - Timecourses spectra is computed for each data-set and averaged across sessions. Mean and standard error of mean is shown in the figure.

- **c) Montage** - Axial slices are shown.
- **d) Ortho slices** - Ortho plot is shown for the peak voxel and coordinates are reported.

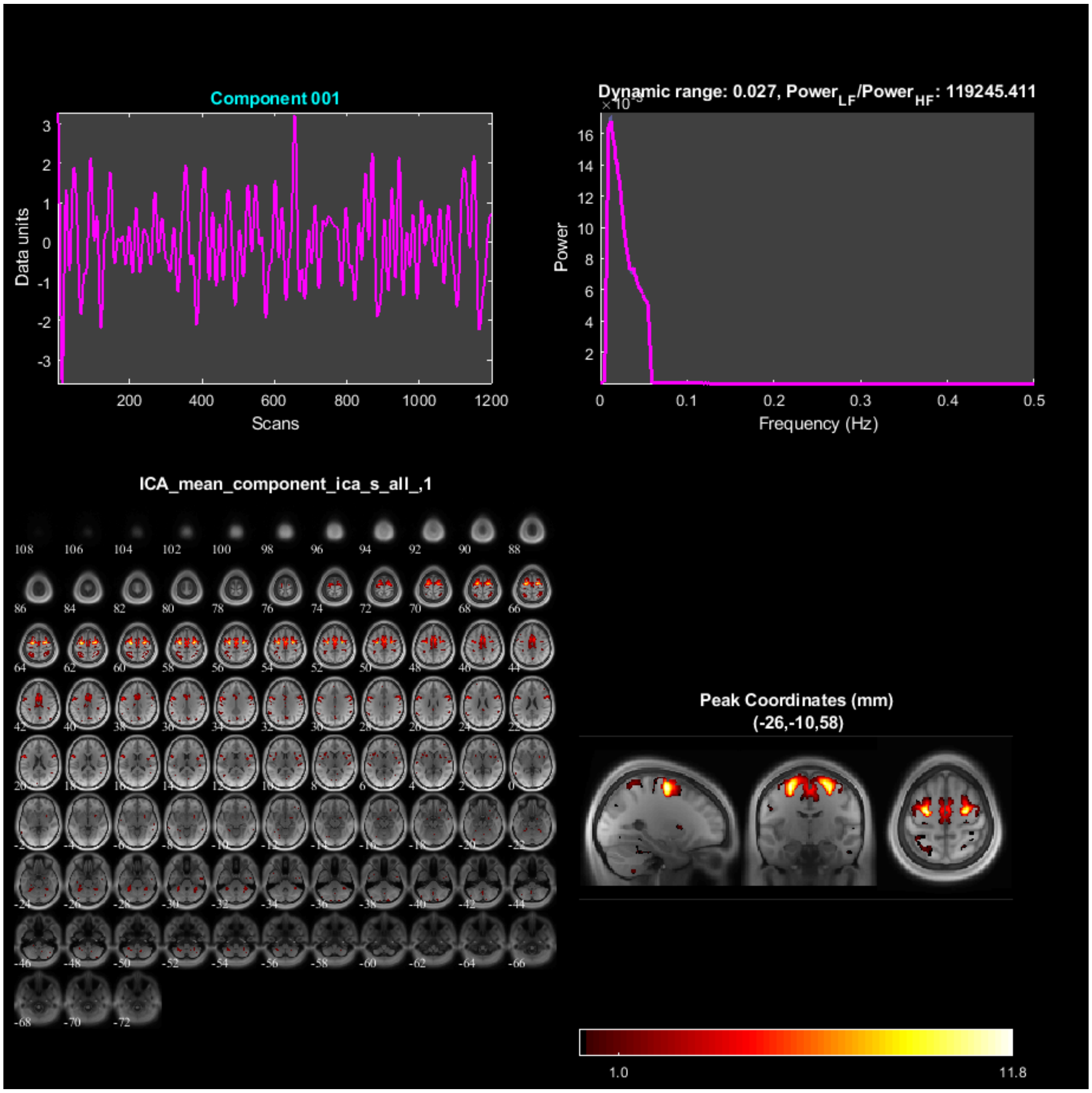

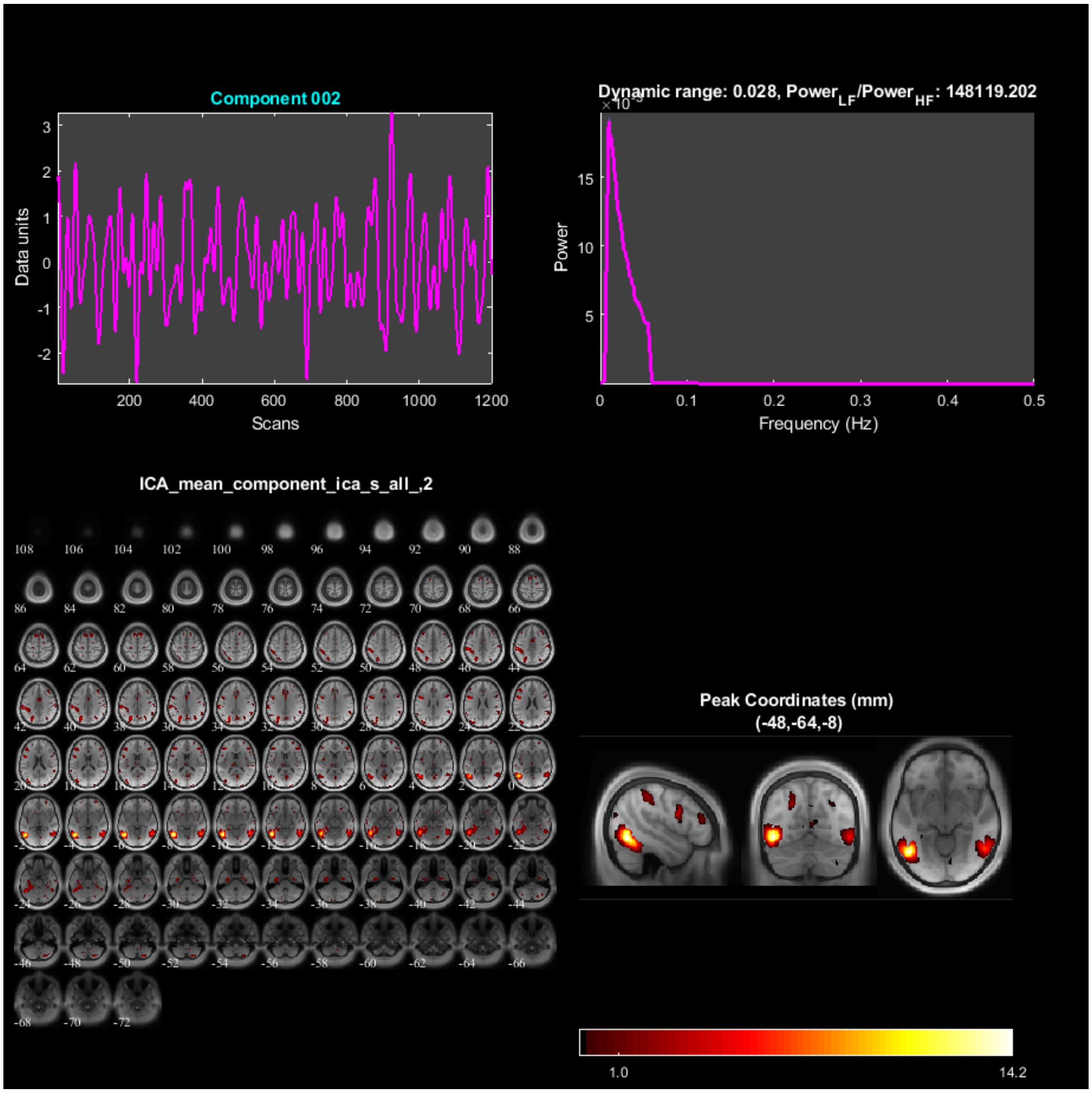
Component 002
Data units
Scans
Dynamic range: 0.028, Power$_{LF}$/Power$_{HF}$: 148119.202
Power
Frequency (Hz)
ICA_mean_component_ica_s_all_,2
Peak Coordinates (mm)
(-48,-64,-8)
1.0
14.2

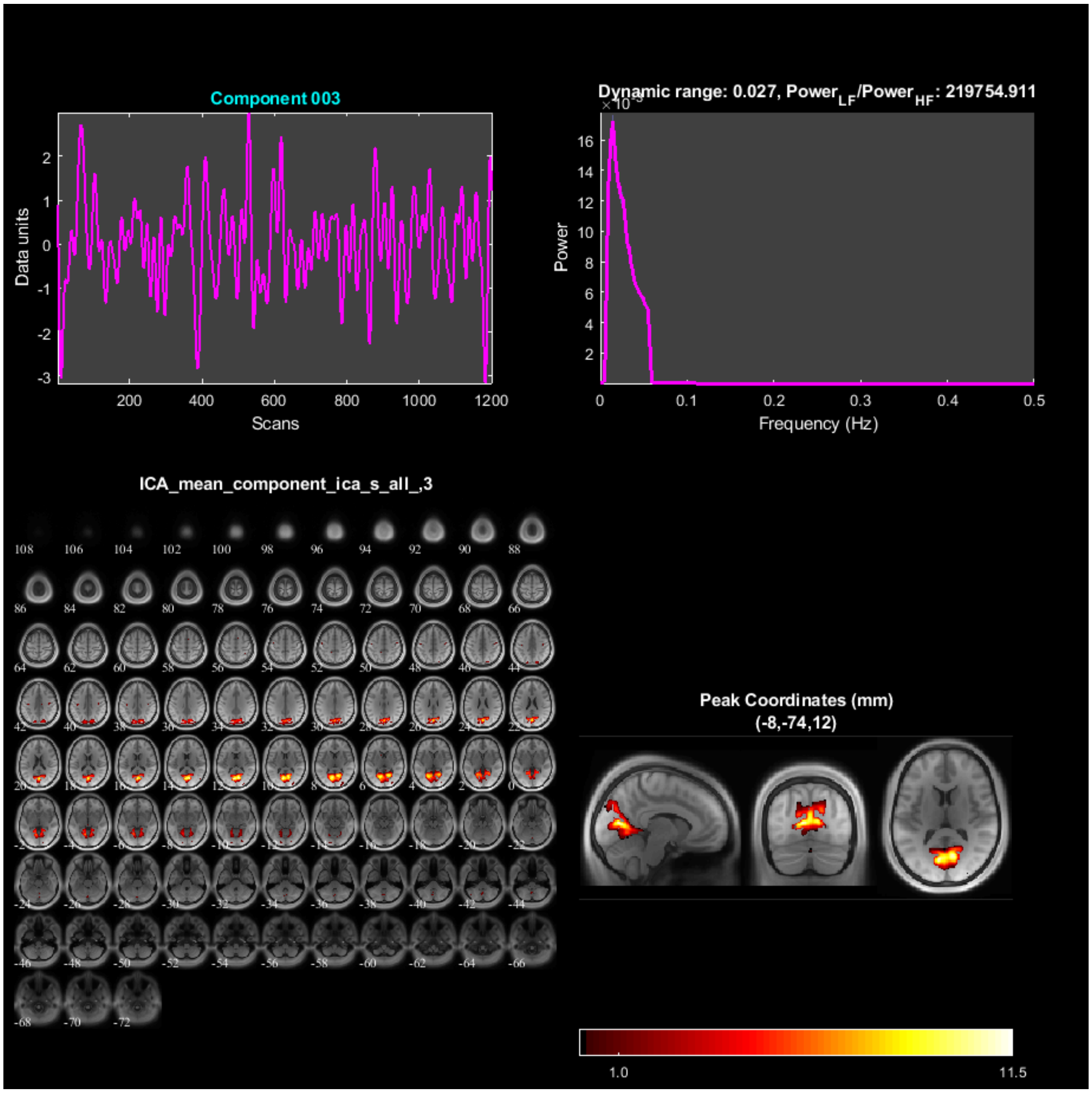
Component 003
Data units
Scans
Dynamic range: 0.027, Power$_{LF}$/Power$_{HF}$: 219754.911
Power
Frequency (Hz)
ICA_mean_component_ica_s_all_,3
Peak Coordinates (mm)
(-8,-74,12)
1.0
11.5

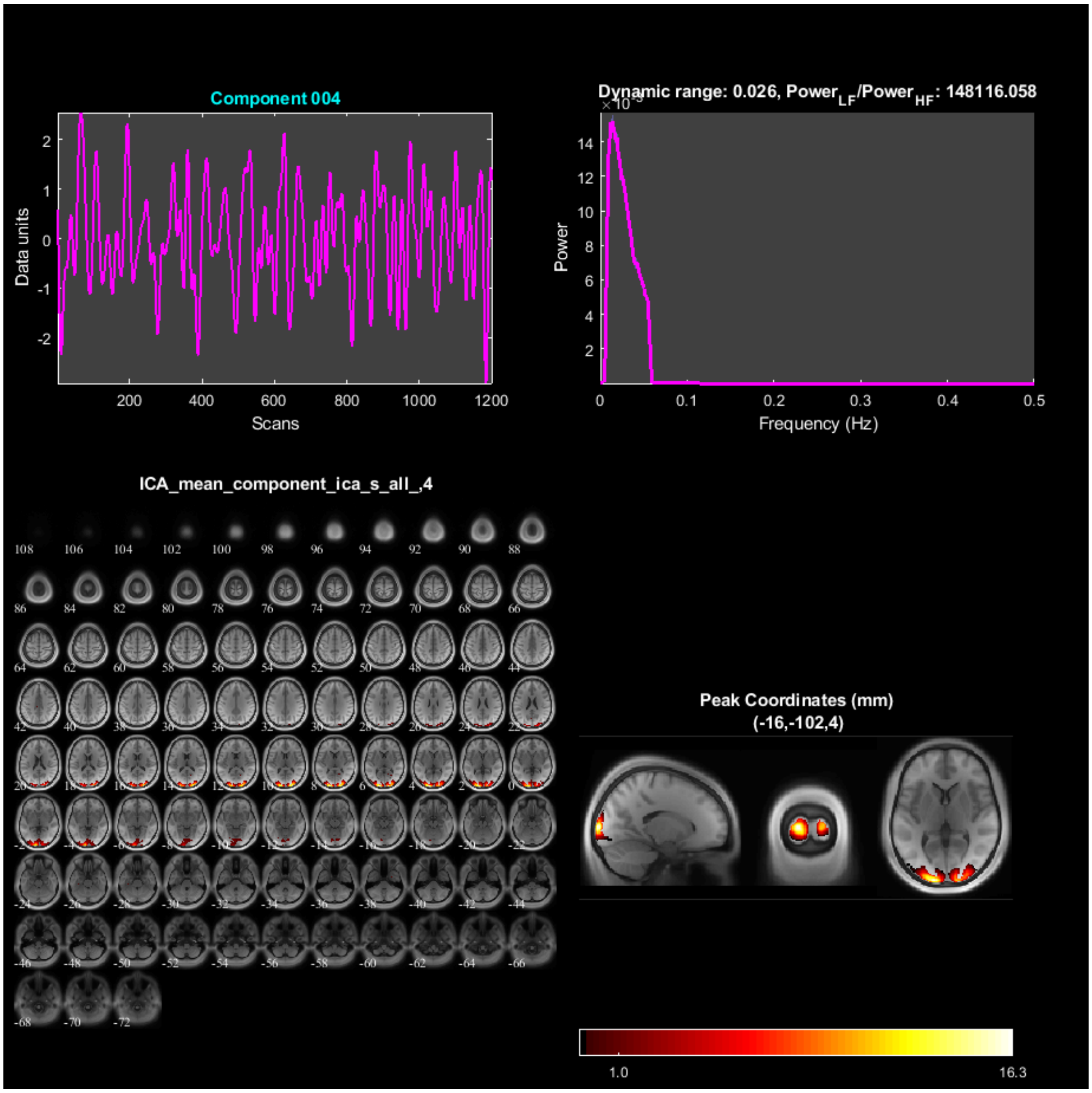
Component 004
Data units
Scans
Dynamic range: 0.026, Power$_{LF}$/Power$_{HF}$: 148116.058
Power
Frequency (Hz)
ICA_mean_component_ica_s_all_,4
Peak Coordinates (mm)
(-16,-102,4)
1.0
16.3

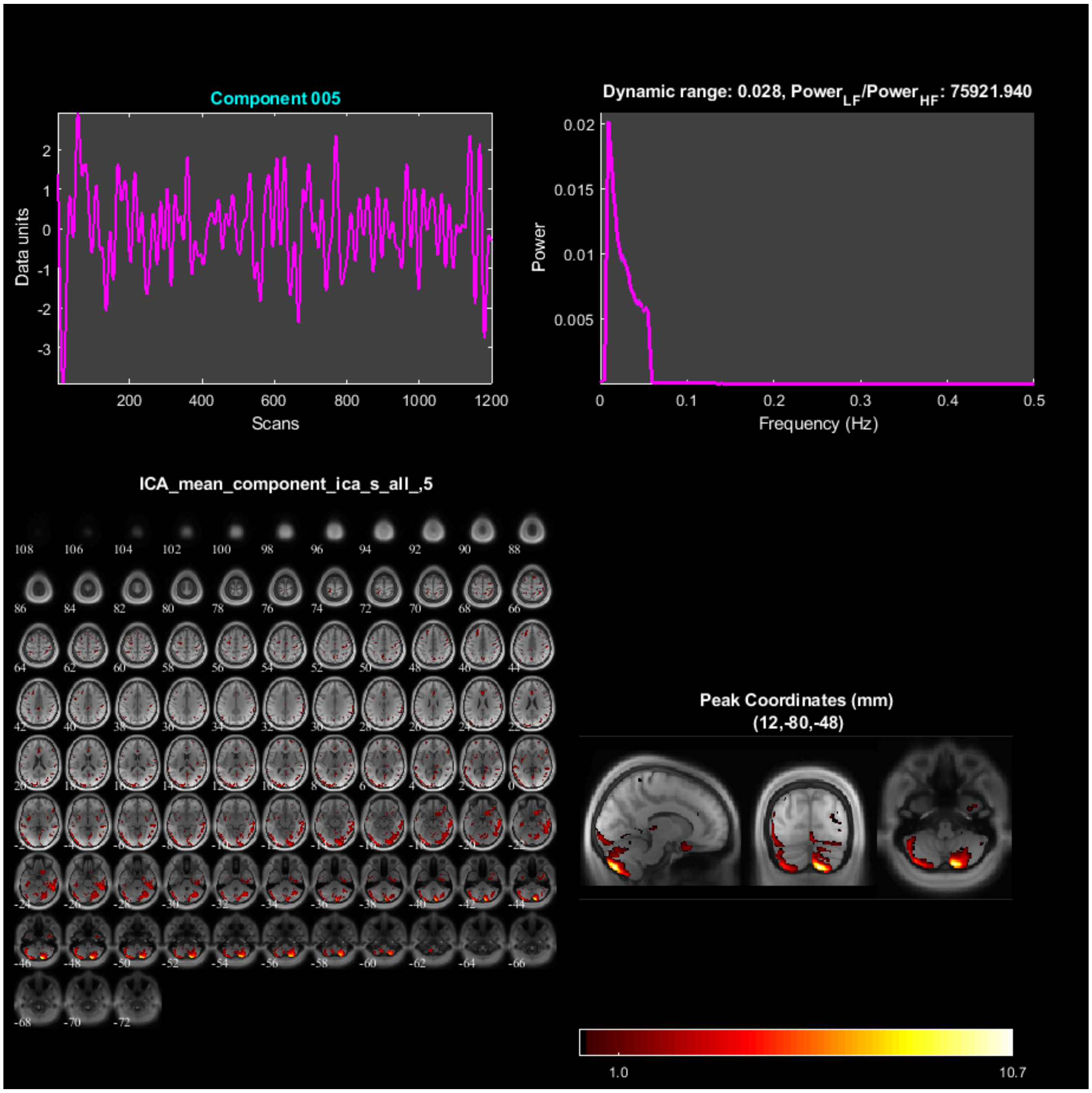

Component 005
Data units
Scans
Dynamic range: 0.028, Power$_{LF}$/Power$_{HF}$: 75921.940
Power
Frequency (Hz)
ICA_mean_component_ica_s_all_,5
Peak Coordinates (mm)
(12,-80,-48)
1.0
10.7

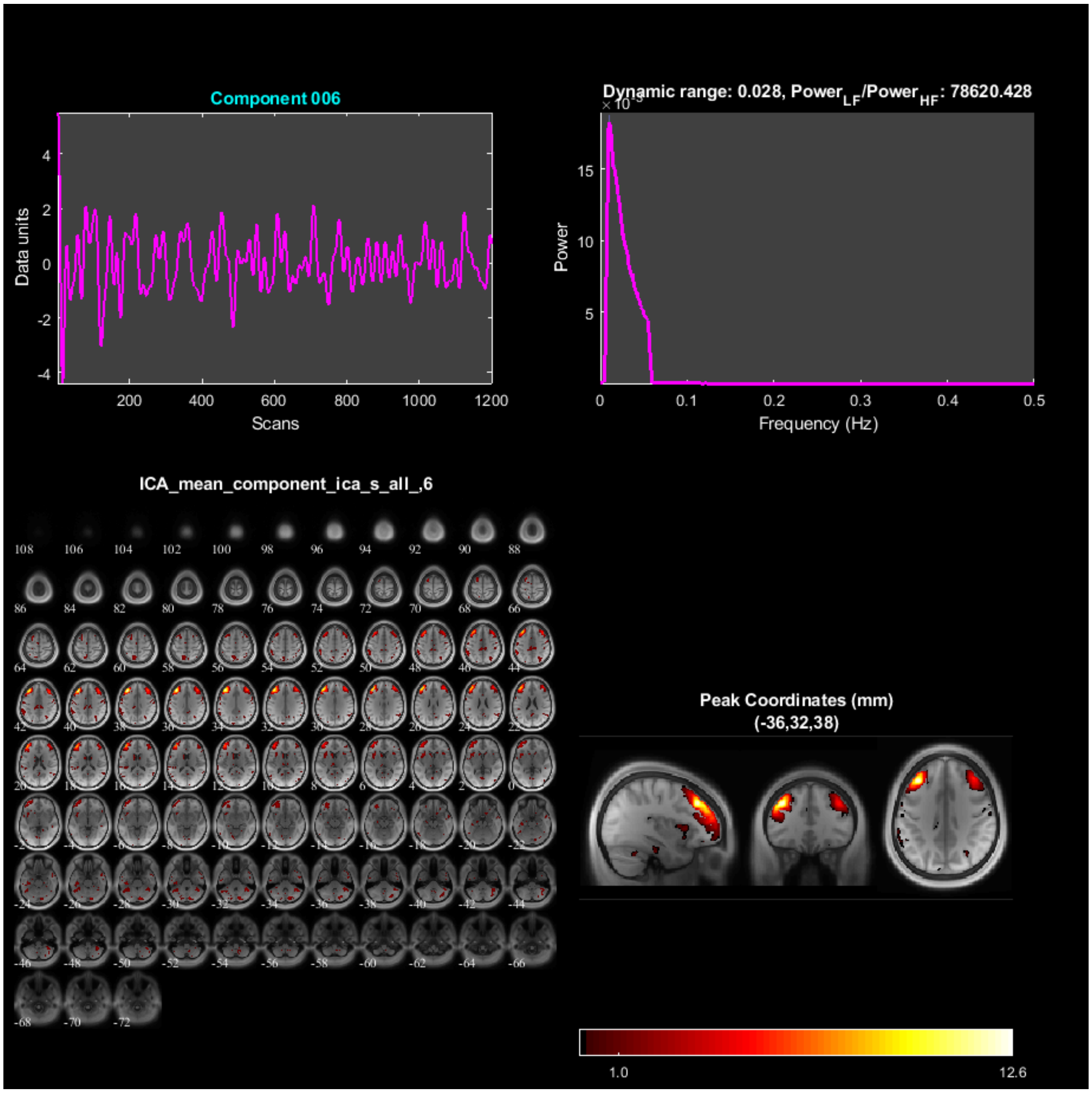

Component 006
Data units
Scans
Dynamic range: 0.028, Power$_{LF}$/Power$_{HF}$: 78620.428
Power
Frequency (Hz)
ICA_mean_component_ica_s_all_,6
Peak Coordinates (mm)
(-36,32,38)
1.0
12.6

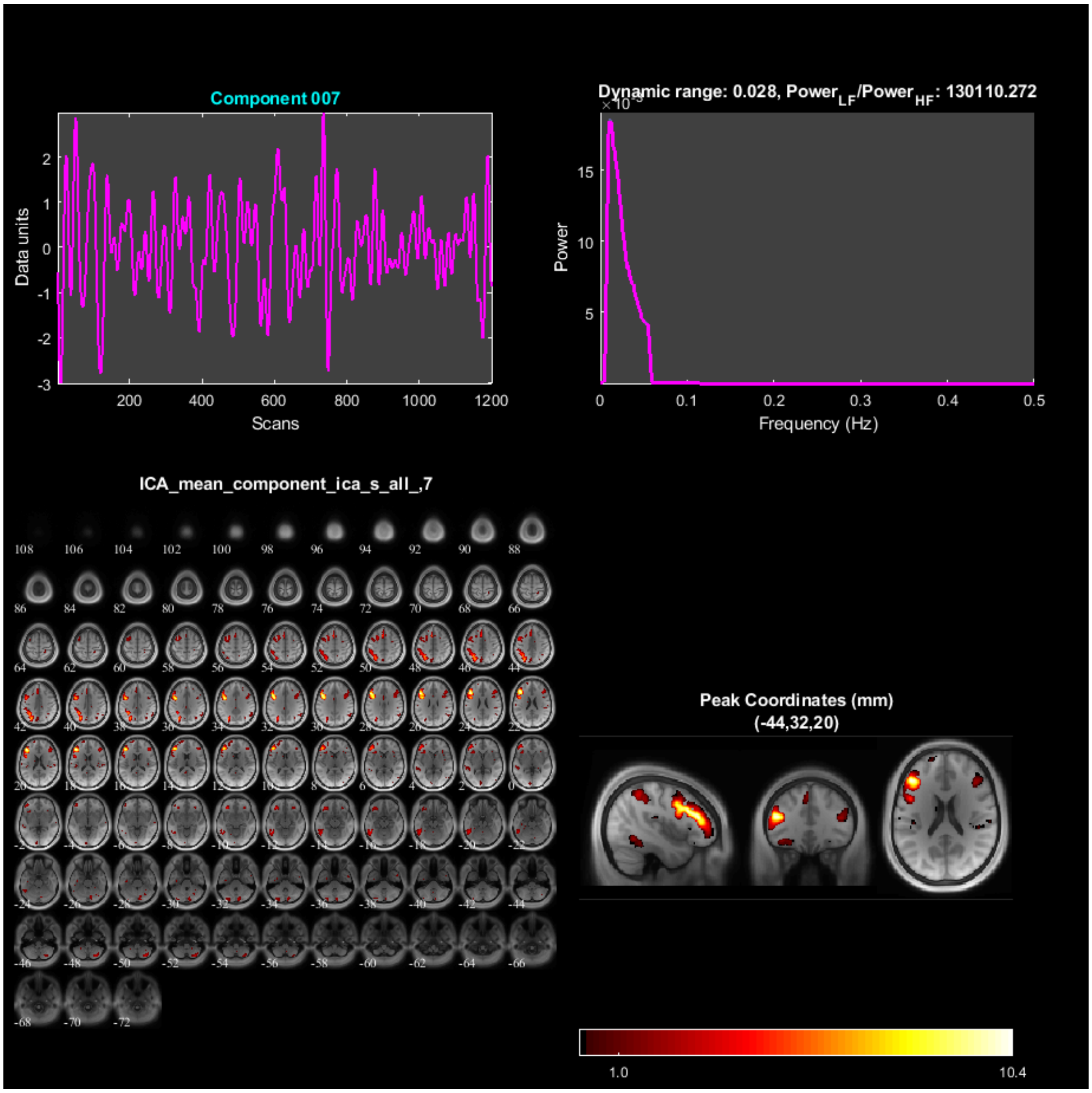

Component 007
Data units
Scans
Dynamic range: 0.028, Power$_{LF}$/Power$_{HF}$: 130110.272
Power
Frequency (Hz)
ICA_mean_component_ica_s_all_,7
Peak Coordinates (mm)
(-44,32,20)
1.0
10.4

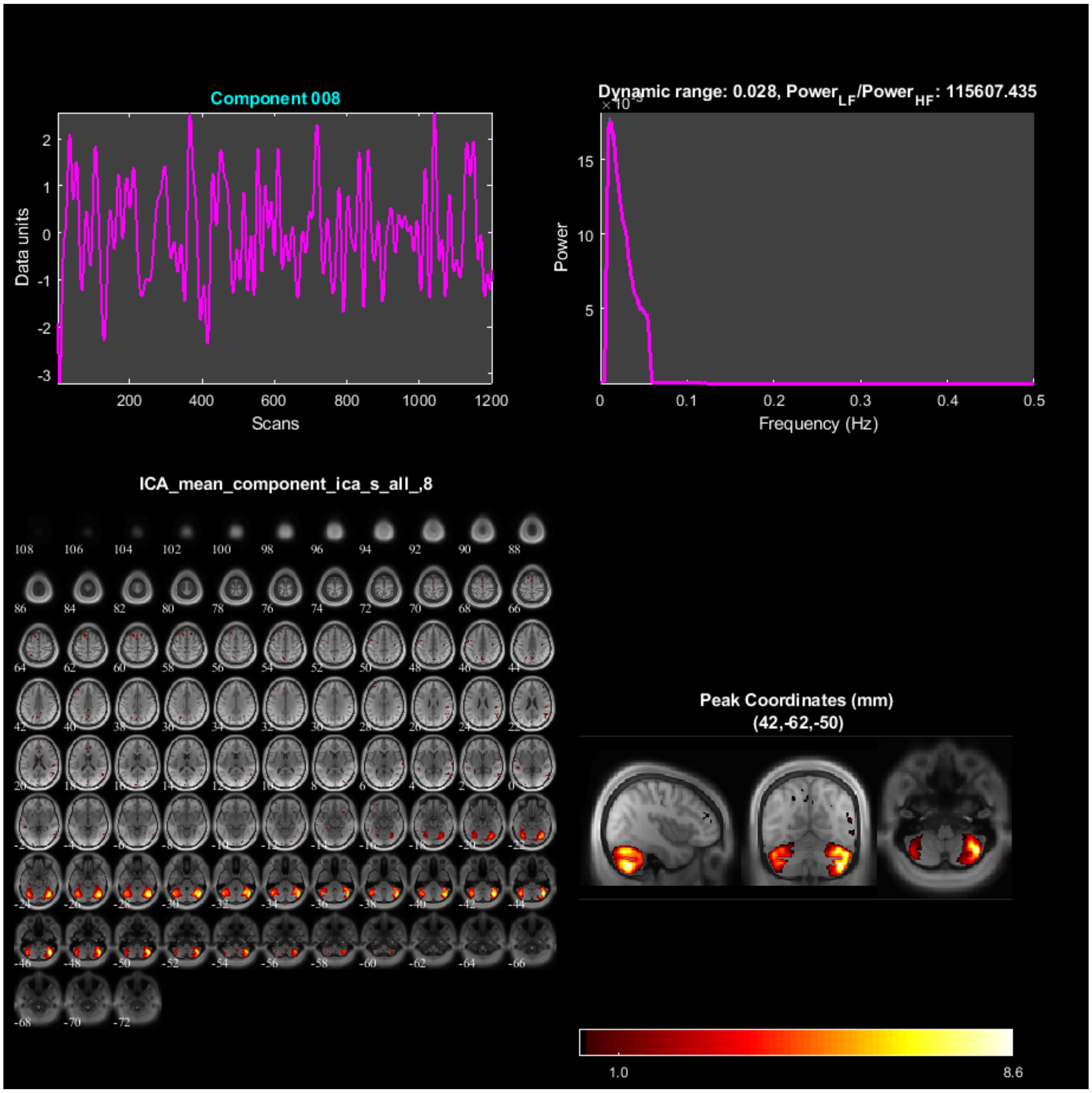

Component 008
Data units
Scans
Dynamic range: 0.028, Power$_{LF}$/Power$_{HF}$: 115607.435
Power
Frequency (Hz)
ICA_mean_component_ica_s_all_,8
Peak Coordinates (mm)
(42,-62,-50)
1.0
8.6

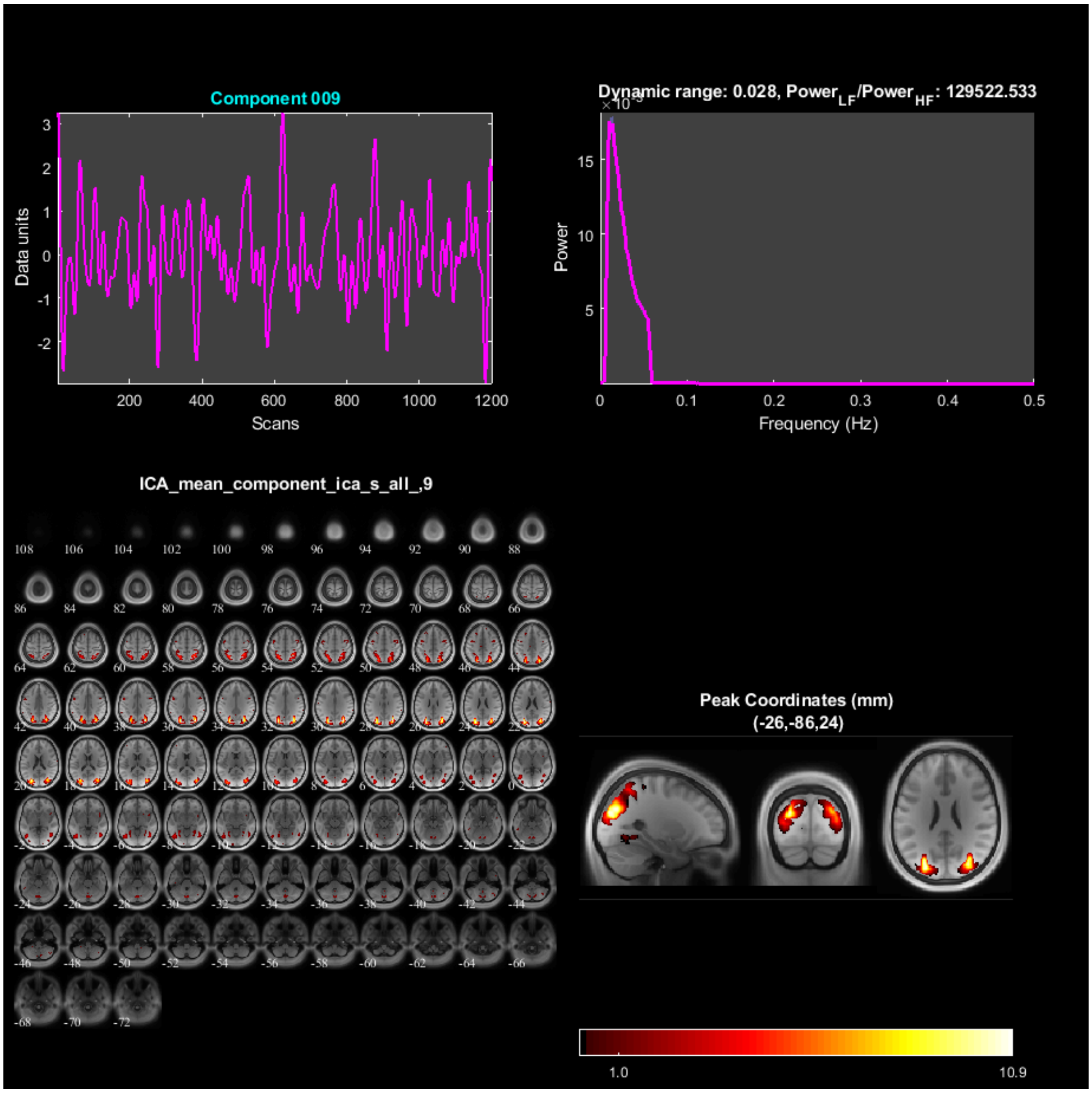
Component 009
Data units
Scans
Dynamic range: 0.028, Power_LF/Power_HF: 129522.533
Power
Frequency (Hz)
ICA_mean_component_ica_s_all_,9
Peak Coordinates (mm)
(-26,-86,24)
1.0
10.9

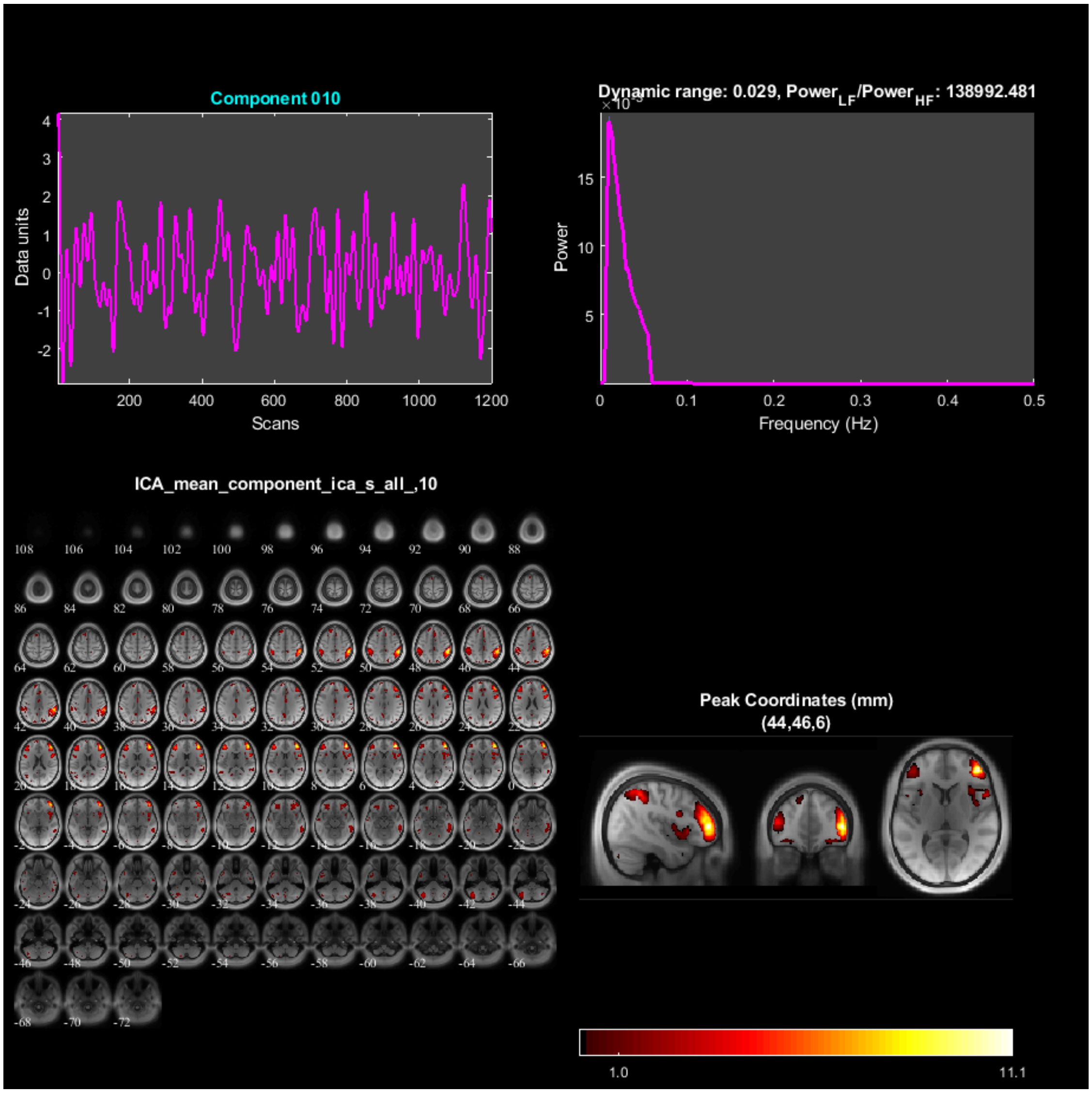

Component 010
Data units
Scans
Dynamic range: 0.029, Power$_{LF}$/Power$_{HF}$: 138992.481
Power
Frequency (Hz)
ICA_mean_component_ica_s_all_,10
Peak Coordinates (mm)
(44,46,6)
1.0
11.1

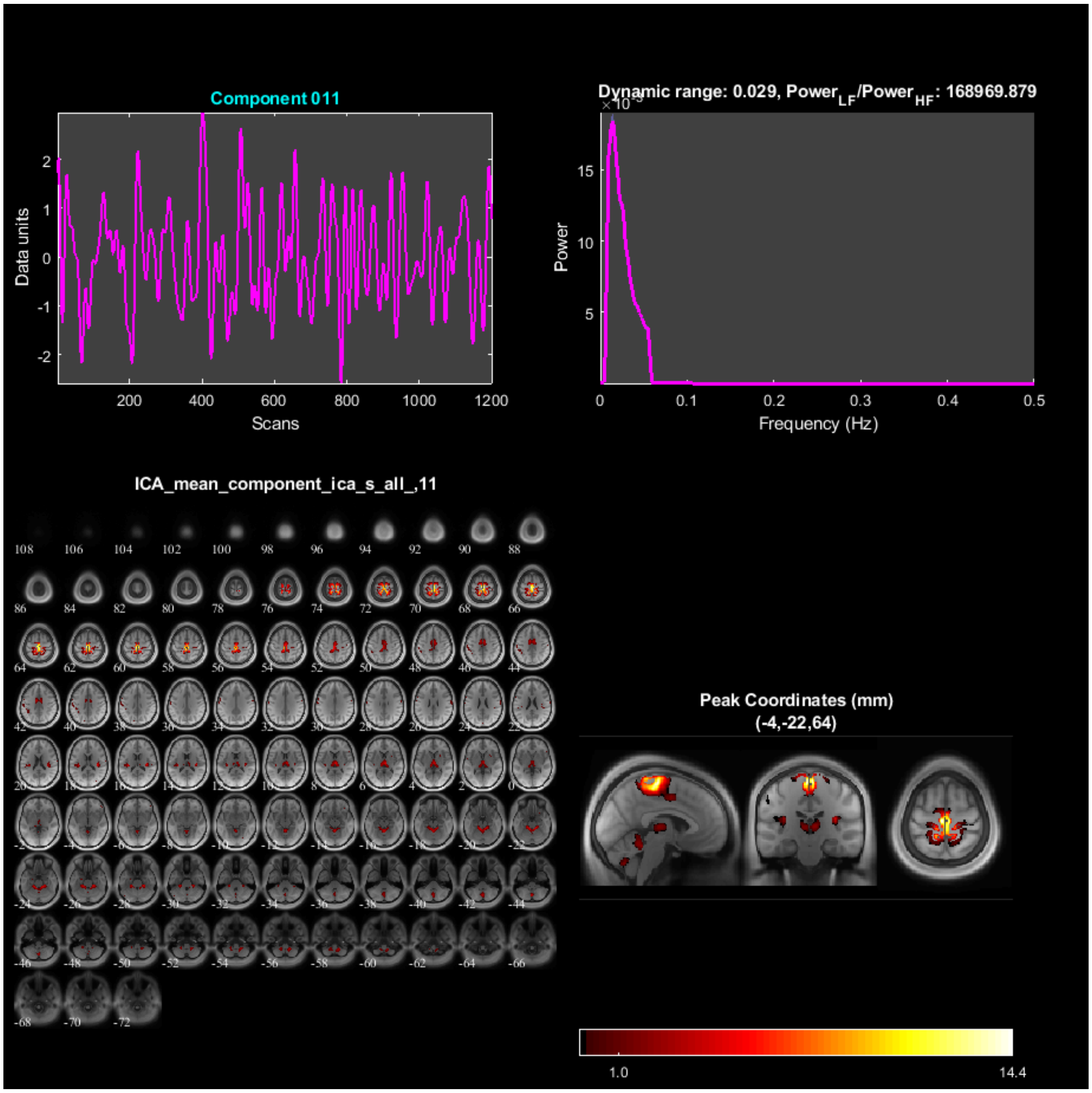

Component 011
Data units
Scans
Dynamic range: 0.029, Power_LF/Power_HF: 168969.879
Power
Frequency (Hz)
ICA_mean_component_ica_s_all_,11
Peak Coordinates (mm)
(-4,-22,64)
1.0
14.4

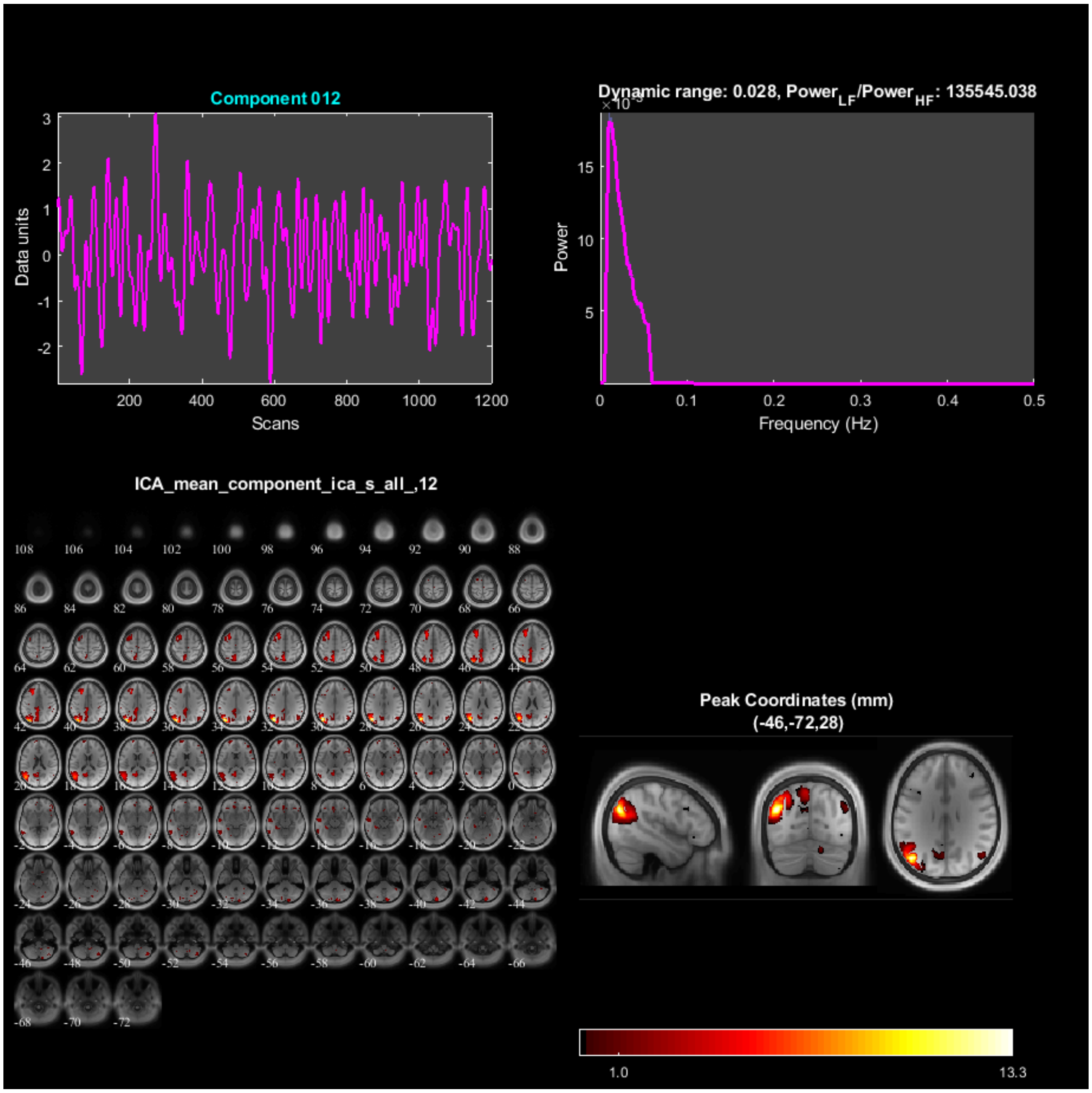
Component 012
Data units
Scans
Dynamic range: 0.028, Power$_{LF}$/Power$_{HF}$: 135545.038
Power
Frequency (Hz)
ICA_mean_component_ica_s_all_,12
Peak Coordinates (mm)
(-46,-72,28)
1.0
13.3

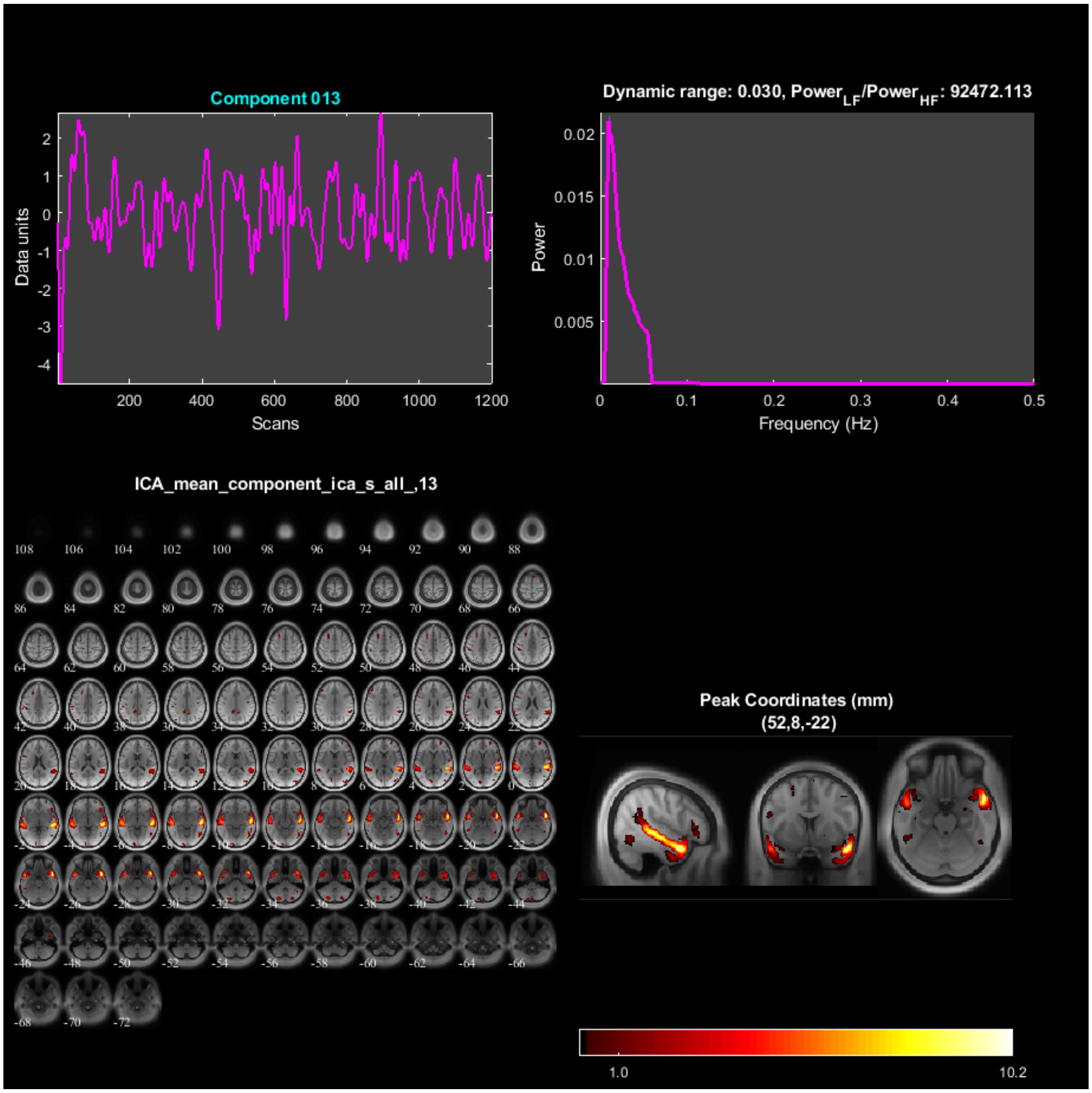
Component 013
Data units
Scans
Dynamic range: 0.030, Power$_{LF}$/Power$_{HF}$: 92472.113
Power
Frequency (Hz)
ICA_mean_component_ica_s_all_,13
Peak Coordinates (mm)
(52,8,-22)
1.0
10.2

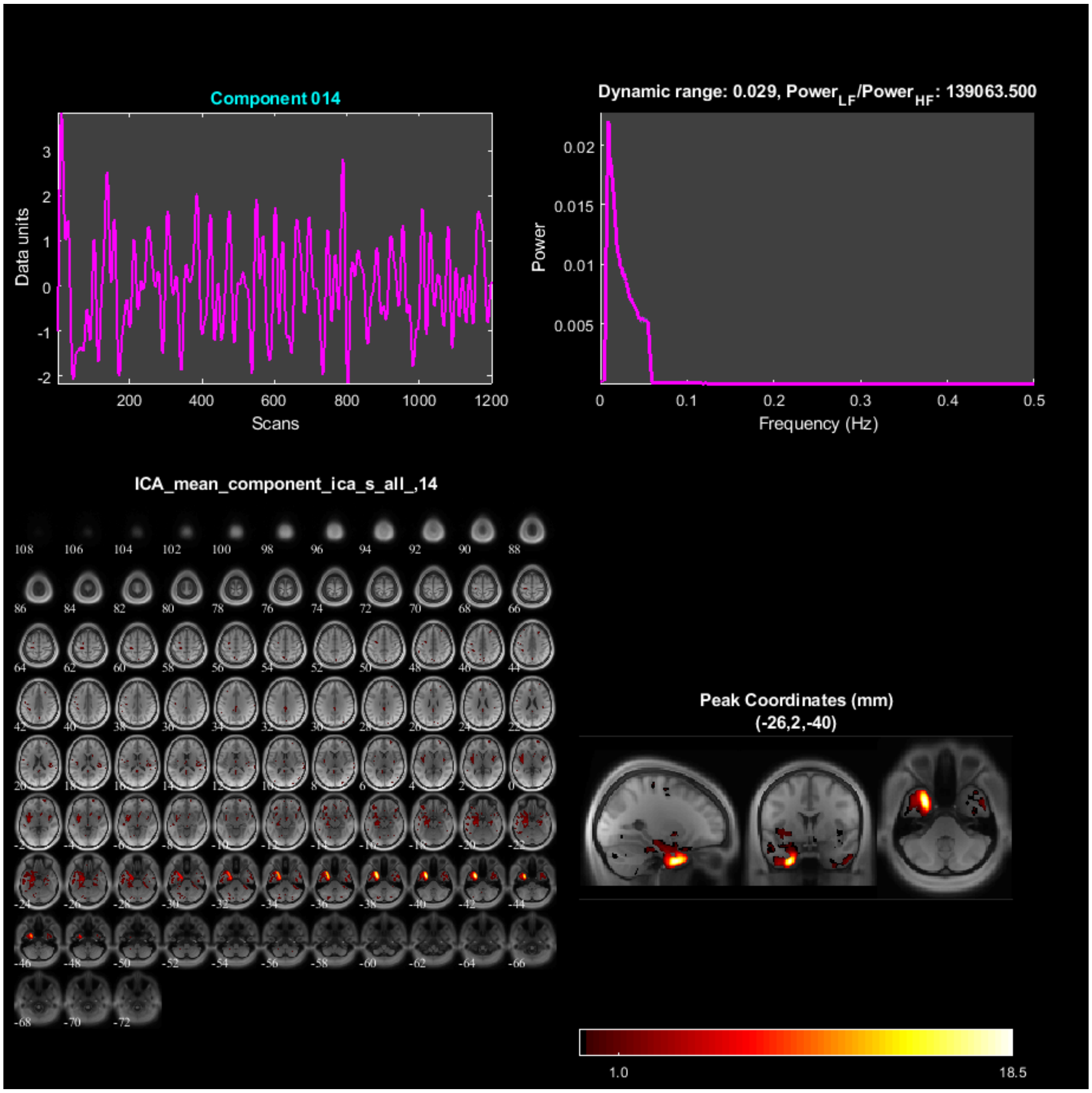
Component 014
Dynamic range: 0.029, Power$_{LF}$/Power$_{HF}$: 139063.500
Data units
Scans
Power
Frequency (Hz)
ICA_mean_component_ica_s_all_,14
Peak Coordinates (mm)
(-26,2,-40)
1.0
18.5

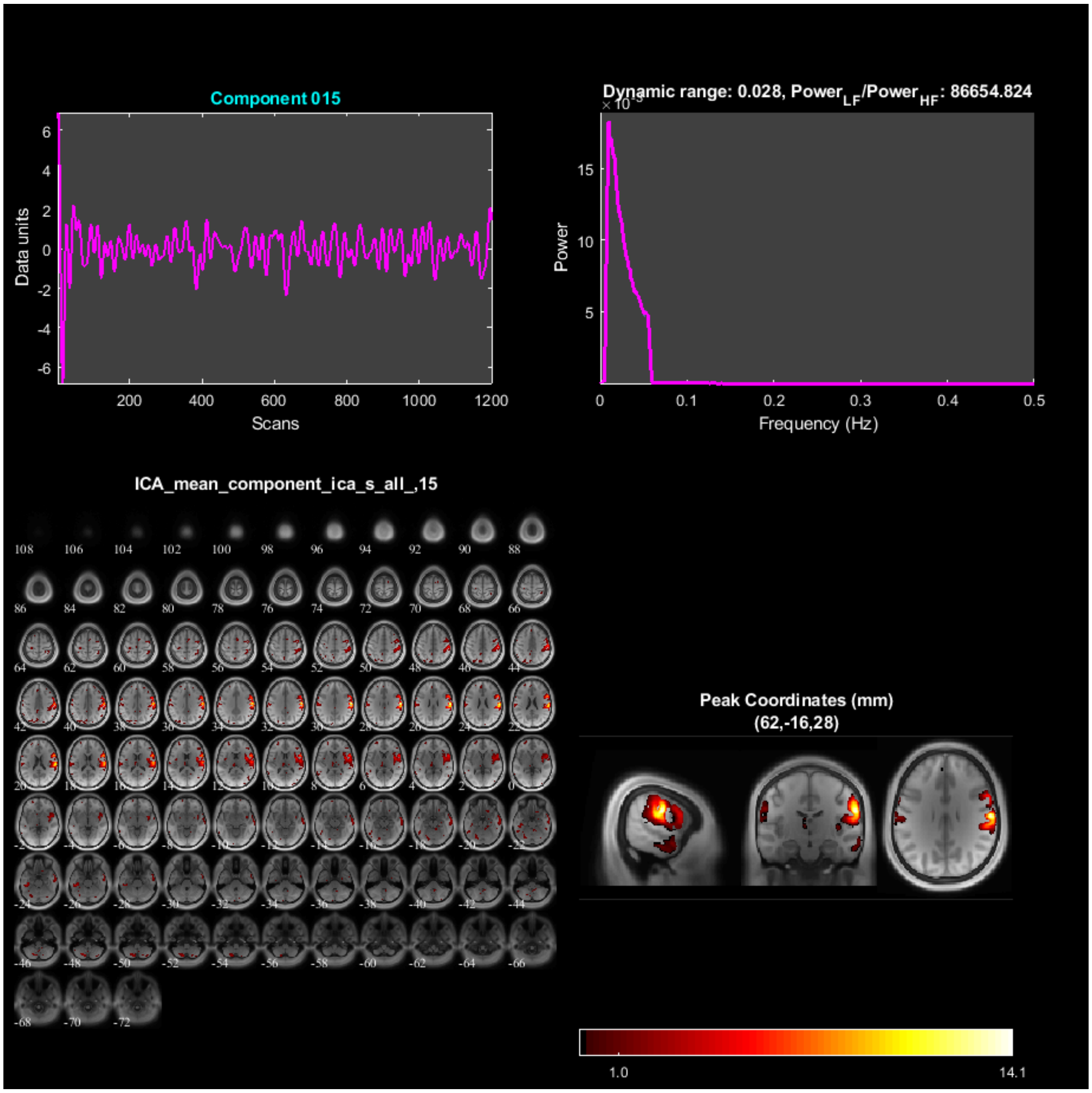
Component 015
Data units
Scans
Dynamic range: 0.028, Power_LF/Power_HF: 86654.824
Power
Frequency (Hz)
ICA_mean_component_ica_s_all_,15
Peak Coordinates (mm)
(62,-16,28)
1.0
14.1

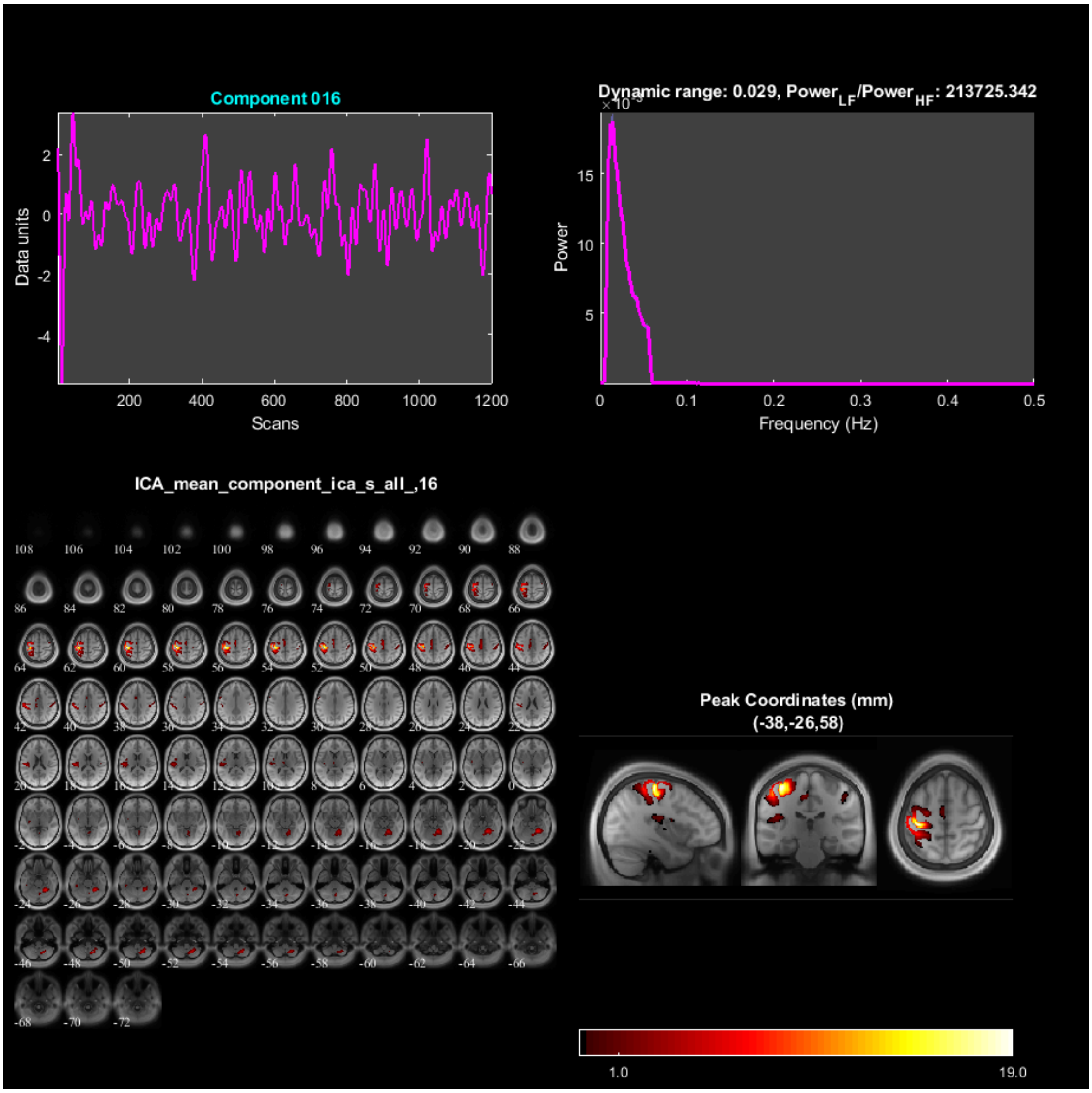
Component 016
Data units
Scans
Dynamic range: 0.029, Power$_{LF}$/Power$_{HF}$: 213725.342
Power
Frequency (Hz)
ICA_mean_component_ica_s_all_,16
Peak Coordinates (mm)
(-38,-26,58)
1.0
19.0

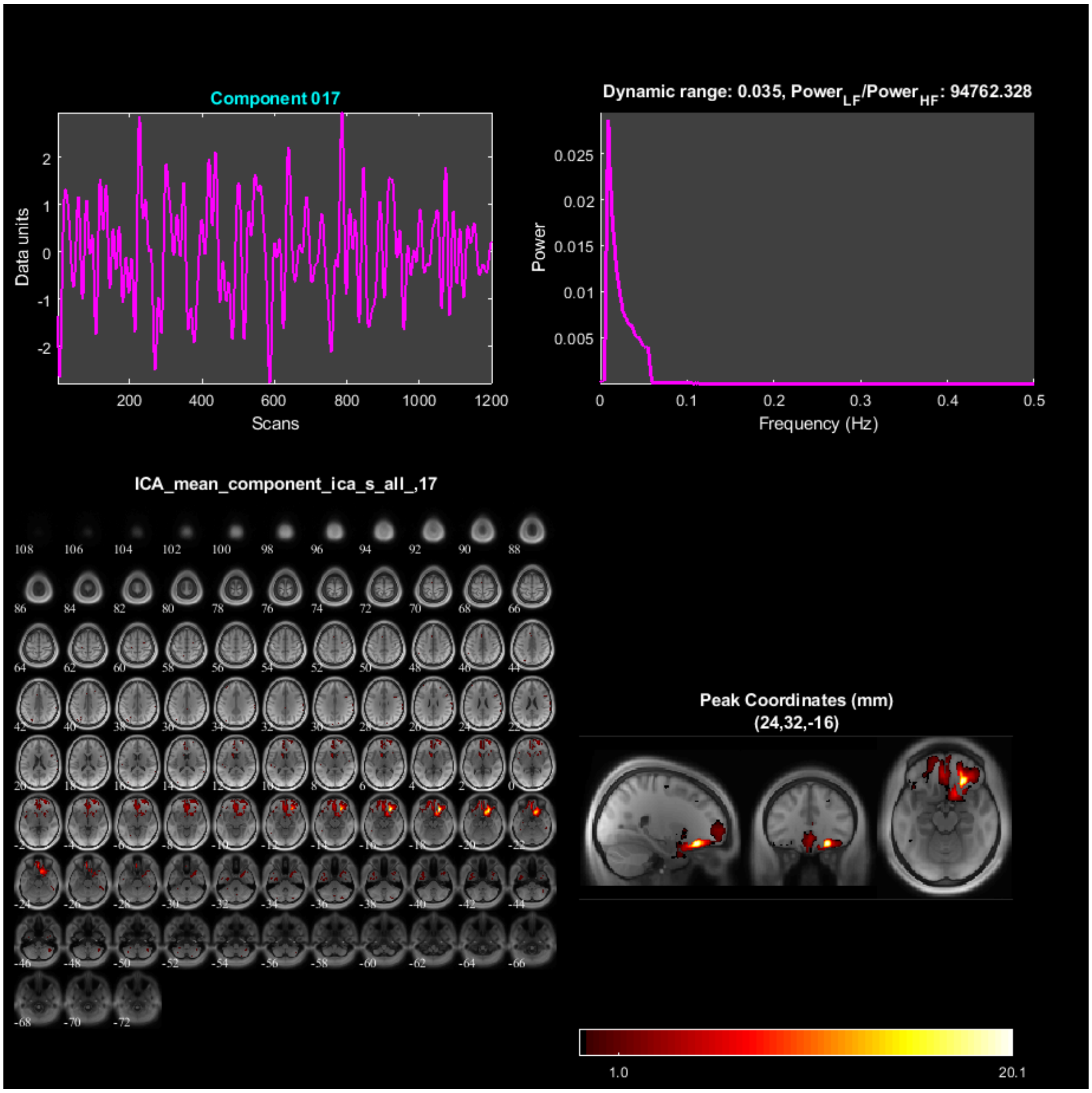
Component 017
Data units
Scans
Dynamic range: 0.035, Power$_{LF}$/Power$_{HF}$: 94762.328
Power
Frequency (Hz)
ICA_mean_component_ica_s_all_,17
Peak Coordinates (mm)
(24,32,-16)
1.0
20.1

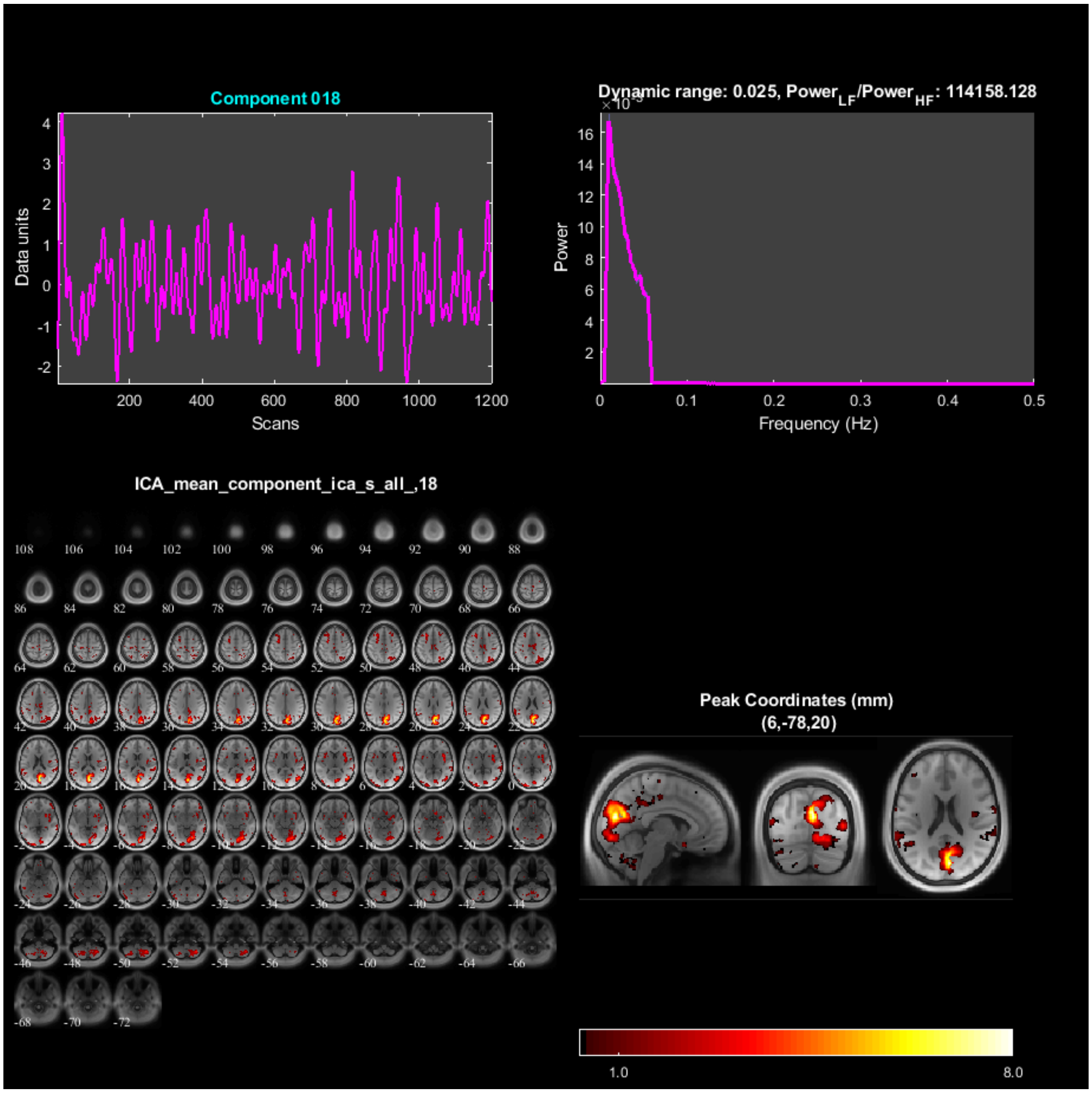

Component 018
Data units
Scans
Dynamic range: 0.025, Power_LF/Power_HF: 114158.128
Power
Frequency (Hz)
ICA_mean_component_ica_s_all_,18
Peak Coordinates (mm)
(6,-78,20)
1.0
8.0

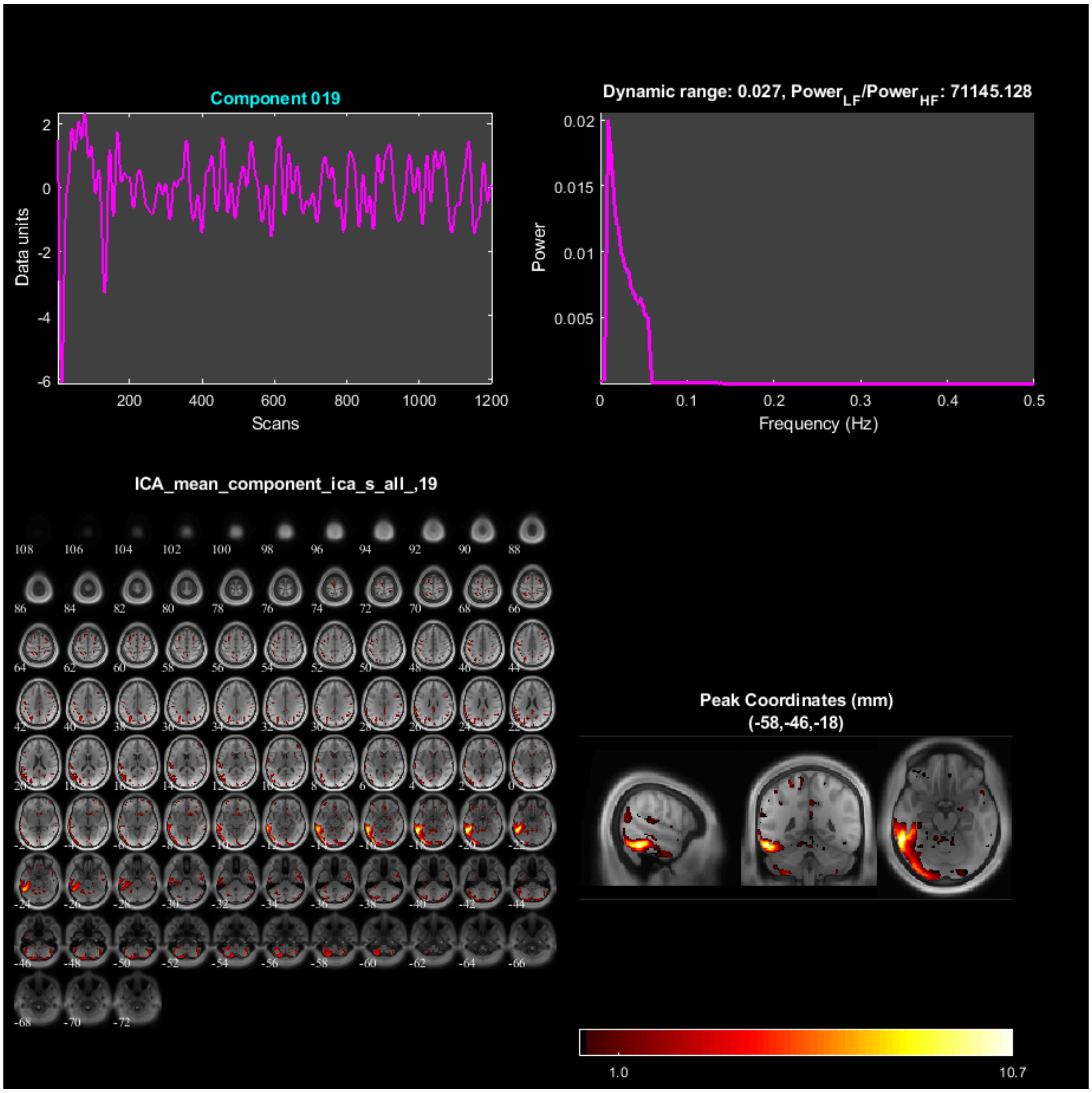
Component 019
Data units
Scans
Dynamic range: 0.027, Power$_{LF}$/Power$_{HF}$: 71145.128
Power
Frequency (Hz)
ICA_mean_component_ica_s_all_,19
Peak Coordinates (mm)
(-58,-46,-18)
1.0
10.7

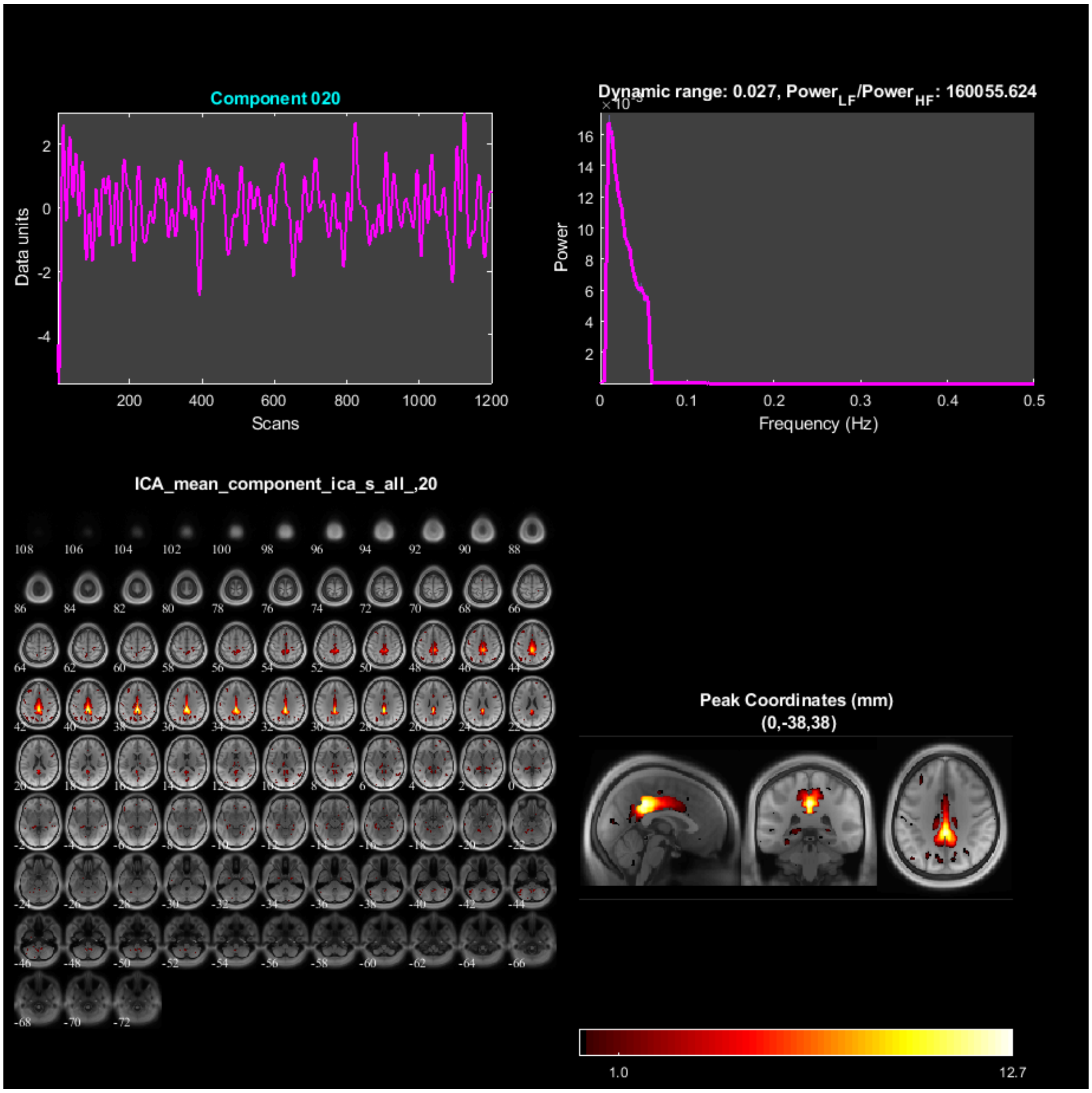
Component 020
Data units
Scans
Dynamic range: 0.027, Power$_{LF}$/Power$_{HF}$: 160055.624
Power
Frequency (Hz)
ICA_mean_component_ica_s_all_,20
Peak Coordinates (mm)
(0,-38,38)
1.0
12.7

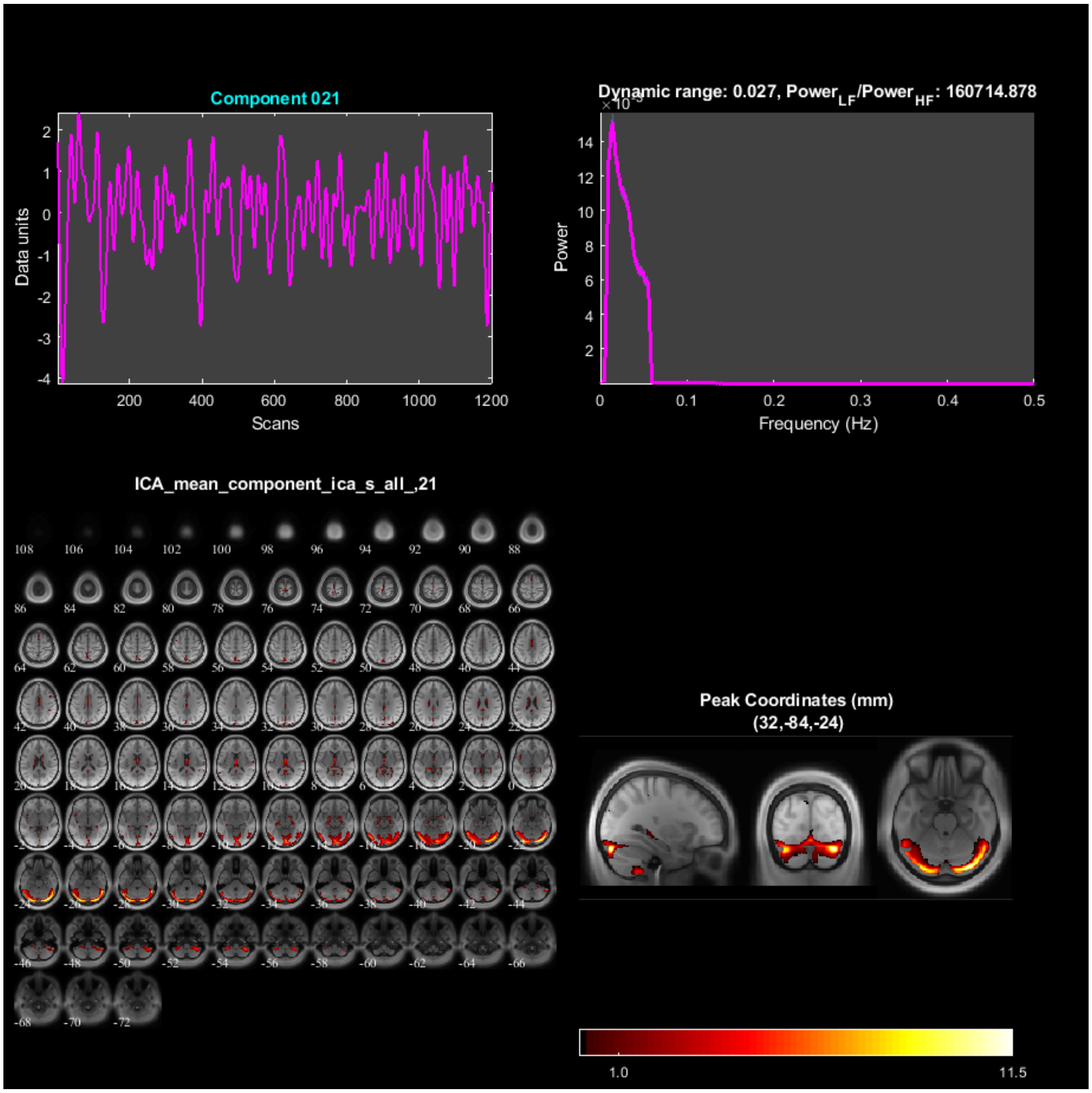
Component 021
Data units
Scans
Dynamic range: 0.027, Power$_{LF}$/Power$_{HF}$: 160714.878
Power
Frequency (Hz)
ICA_mean_component_ica_s_all_,21
Peak Coordinates (mm)
(32,-84,-24)
1.0
11.5

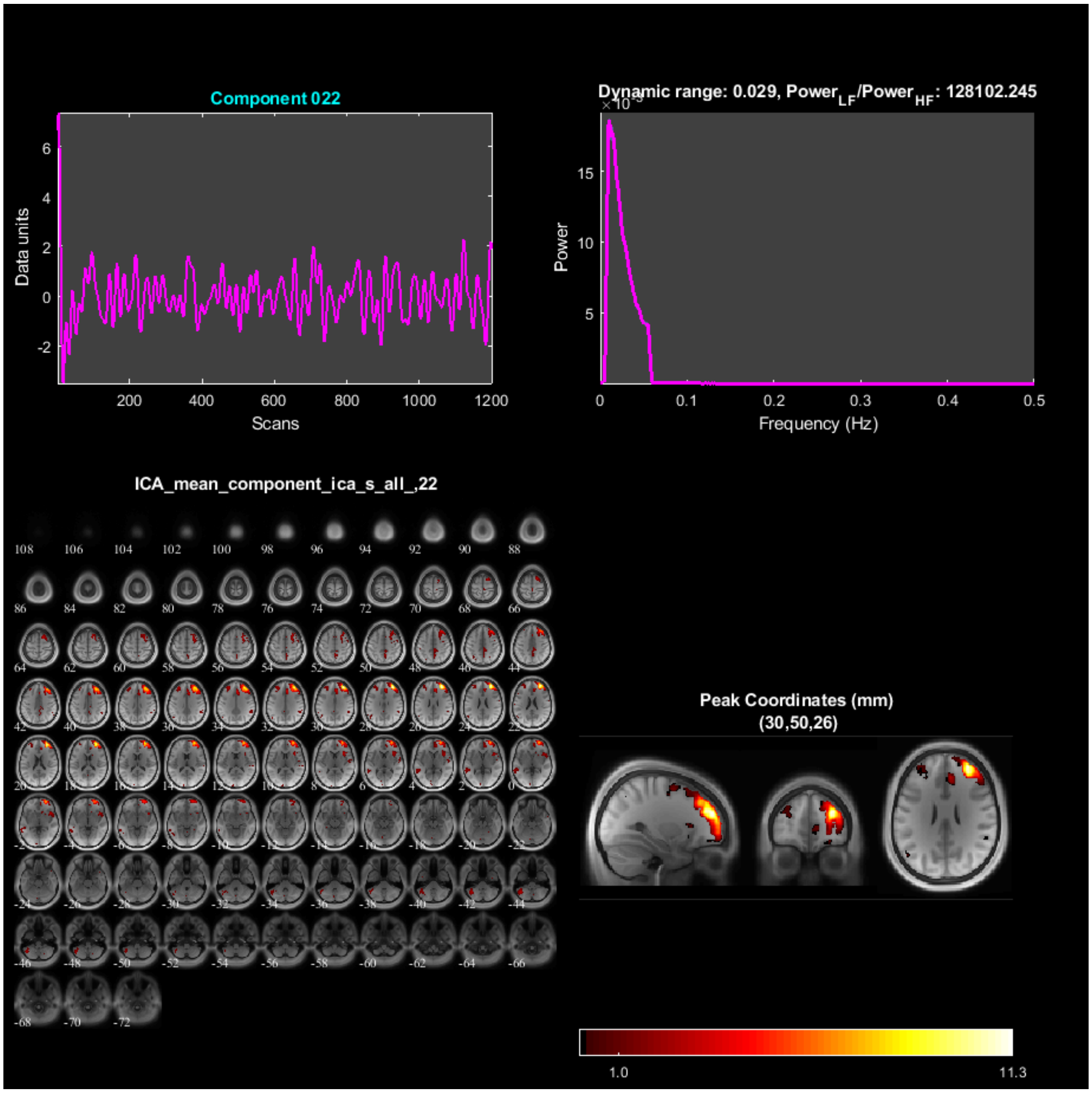

Component 022
Data units
Scans
Dynamic range: 0.029, Power$_{LF}$/Power$_{HF}$: 128102.245
Power
Frequency (Hz)
ICA_mean_component_ica_s_all_,22
Peak Coordinates (mm)
(30,50,26)
1.0
11.3

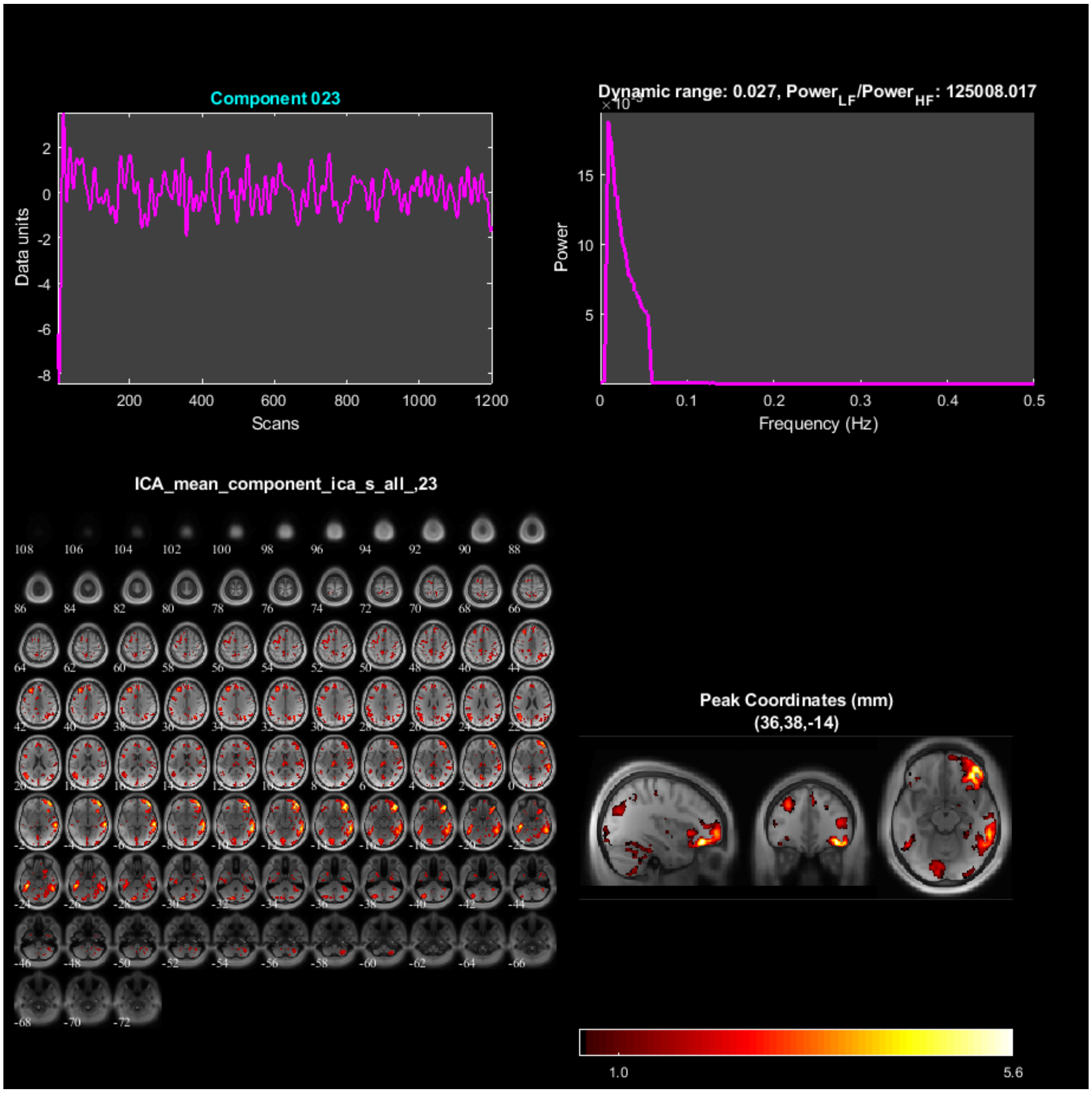
Component 023
Data units
Scans
Dynamic range: 0.027, Power$_{LF}$/Power$_{HF}$: 125008.017
Power
Frequency (Hz)
ICA_mean_component_ica_s_all_,23
Peak Coordinates (mm)
(36,38,-14)
1.0
5.6

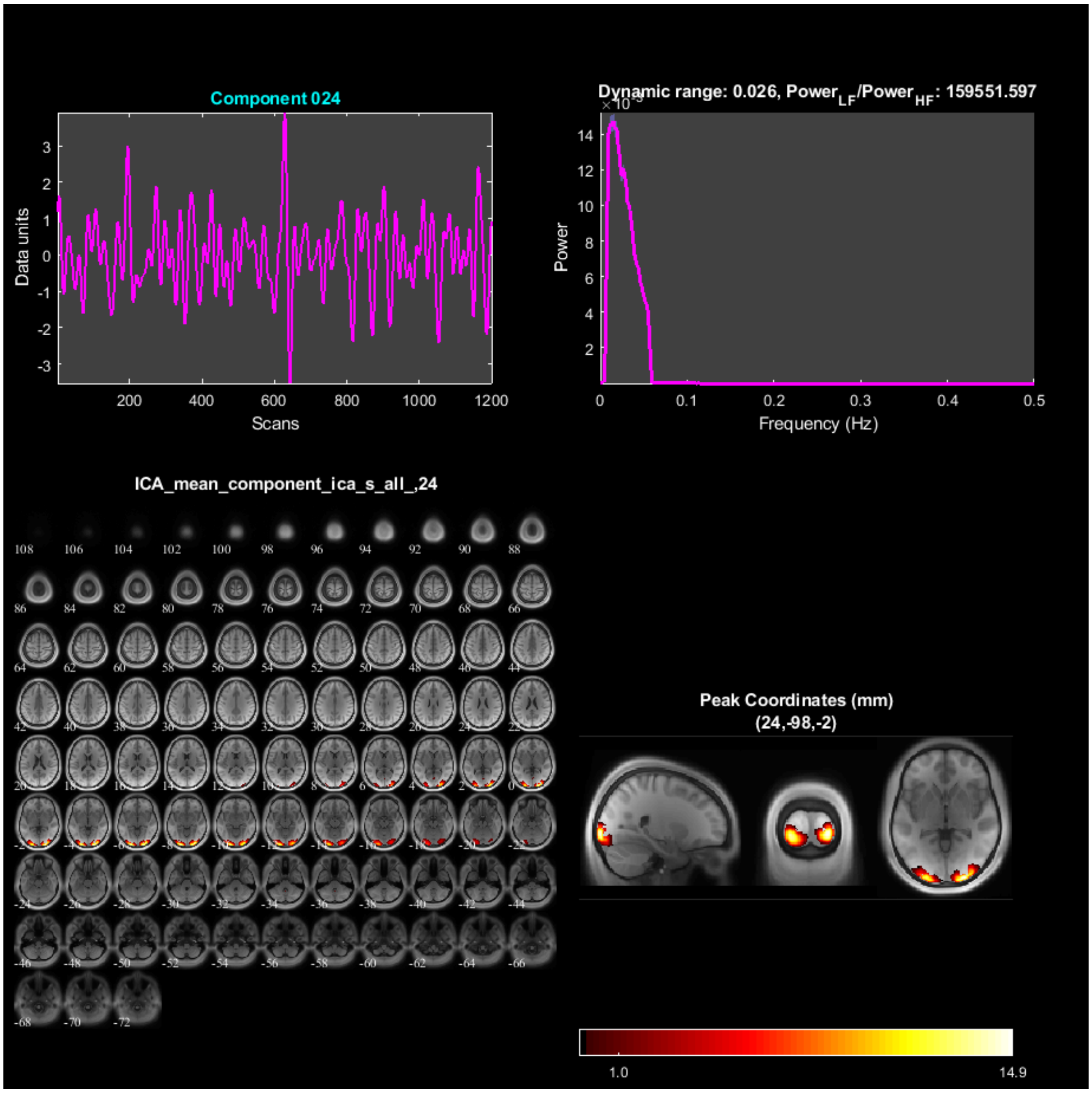
Component 024
Data units
Scans
Dynamic range: 0.026, Power$_{LF}$/Power$_{HF}$: 159551.597
Power
Frequency (Hz)
ICA_mean_component_ica_s_all_,24
Peak Coordinates (mm)
(24,-98,-2)
1.0
14.9

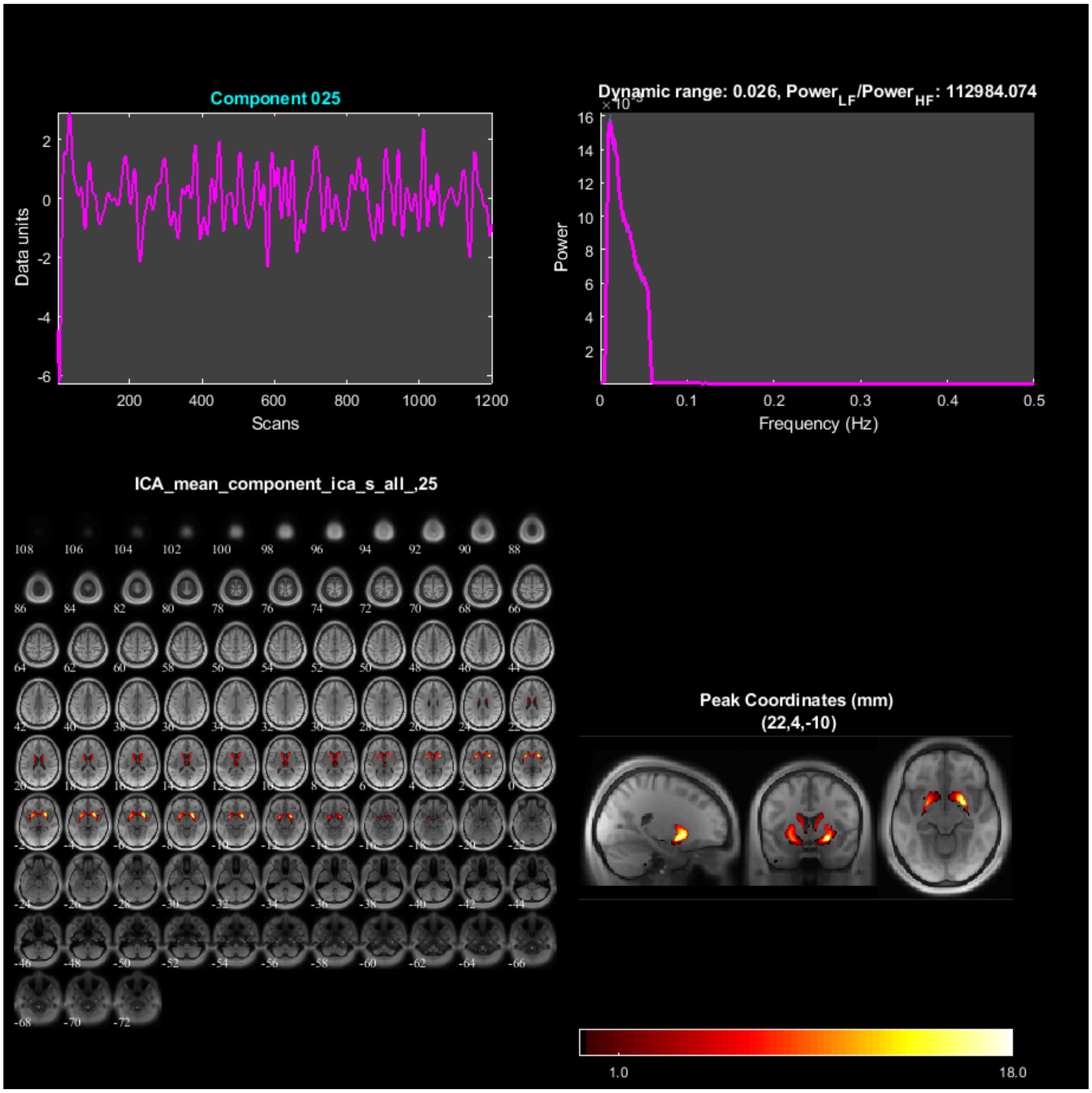
Component 025
Data units
Scans
Dynamic range: 0.026, Power_LF/Power_HF: 112984.074
Power
Frequency (Hz)
ICA_mean_component_ica_s_all_,25
Peak Coordinates (mm)
(22,4,-10)
1.0
18.0

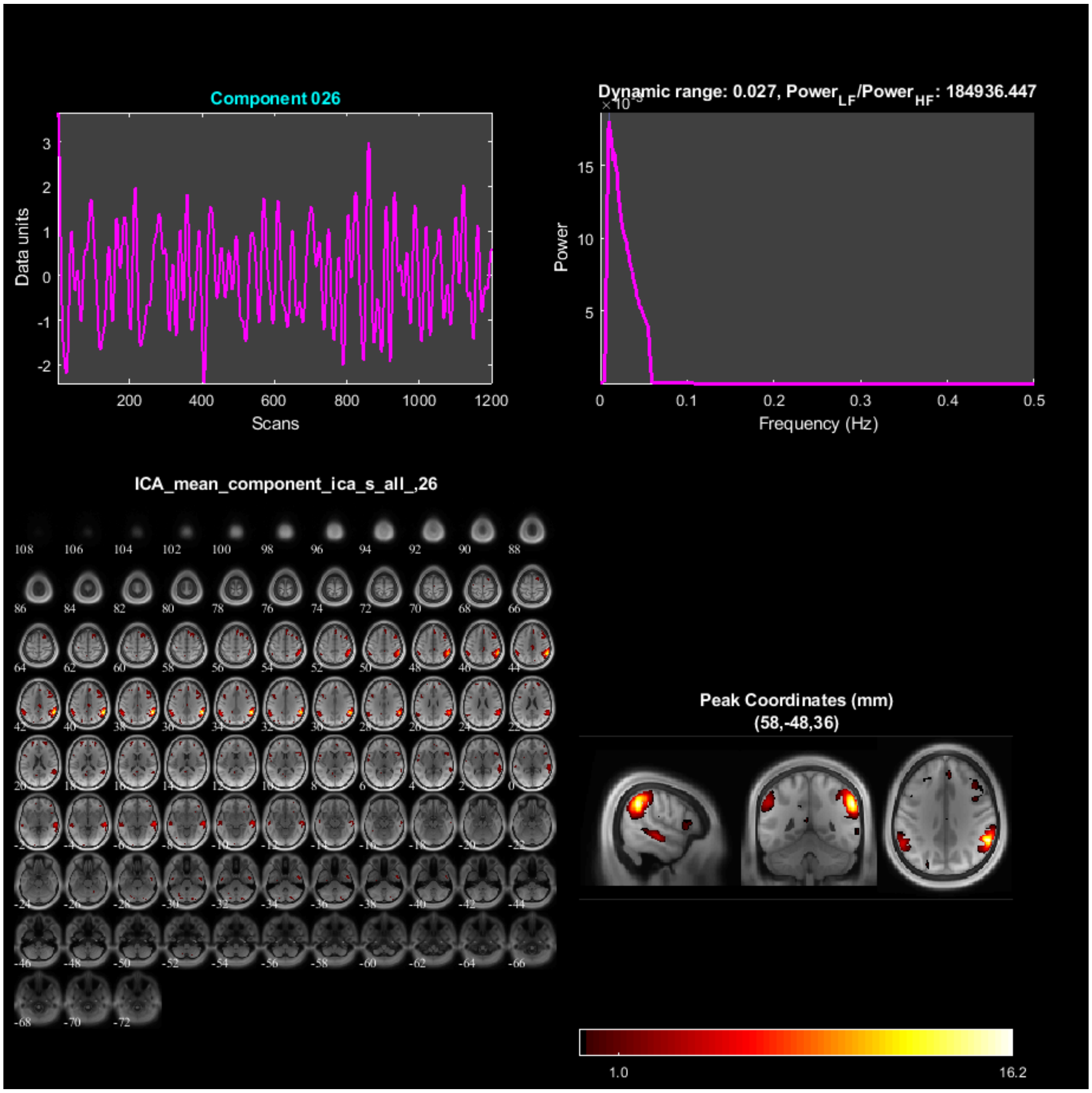
Component 026
Data units
Scans
Dynamic range: 0.027, Power$_{LF}$/Power$_{HF}$: 184936.447
Power
Frequency (Hz)
ICA_mean_component_ica_s_all_,26
Peak Coordinates (mm)
(58,-48,36)
1.0
16.2

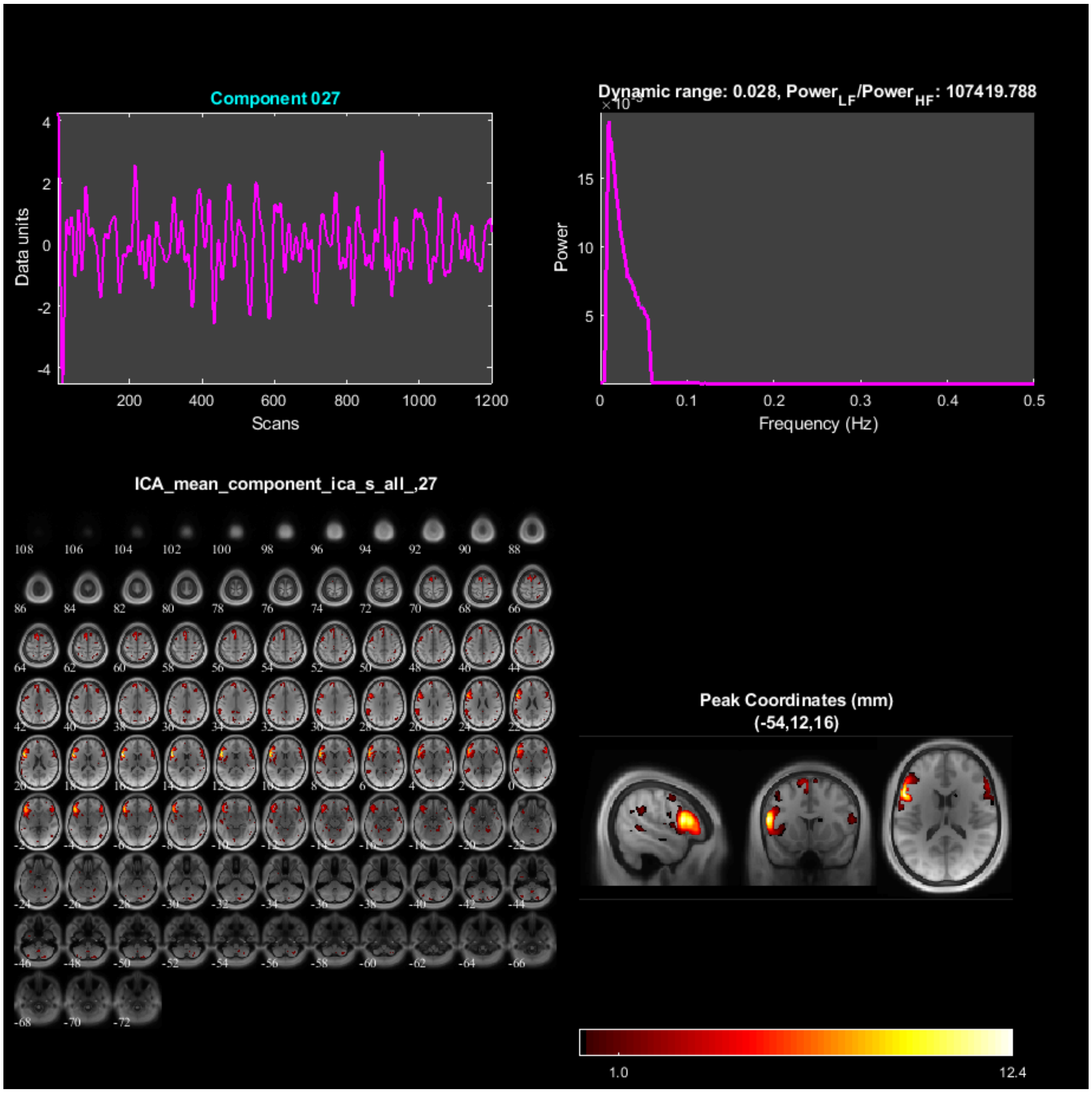

Component 027
Data units
Scans
Dynamic range: 0.028, Power$_{LF}$/Power$_{HF}$: 107419.788
Power
Frequency (Hz)
ICA_mean_component_ica_s_all_,27
Peak Coordinates (mm)
(-54,12,16)
1.0
12.4

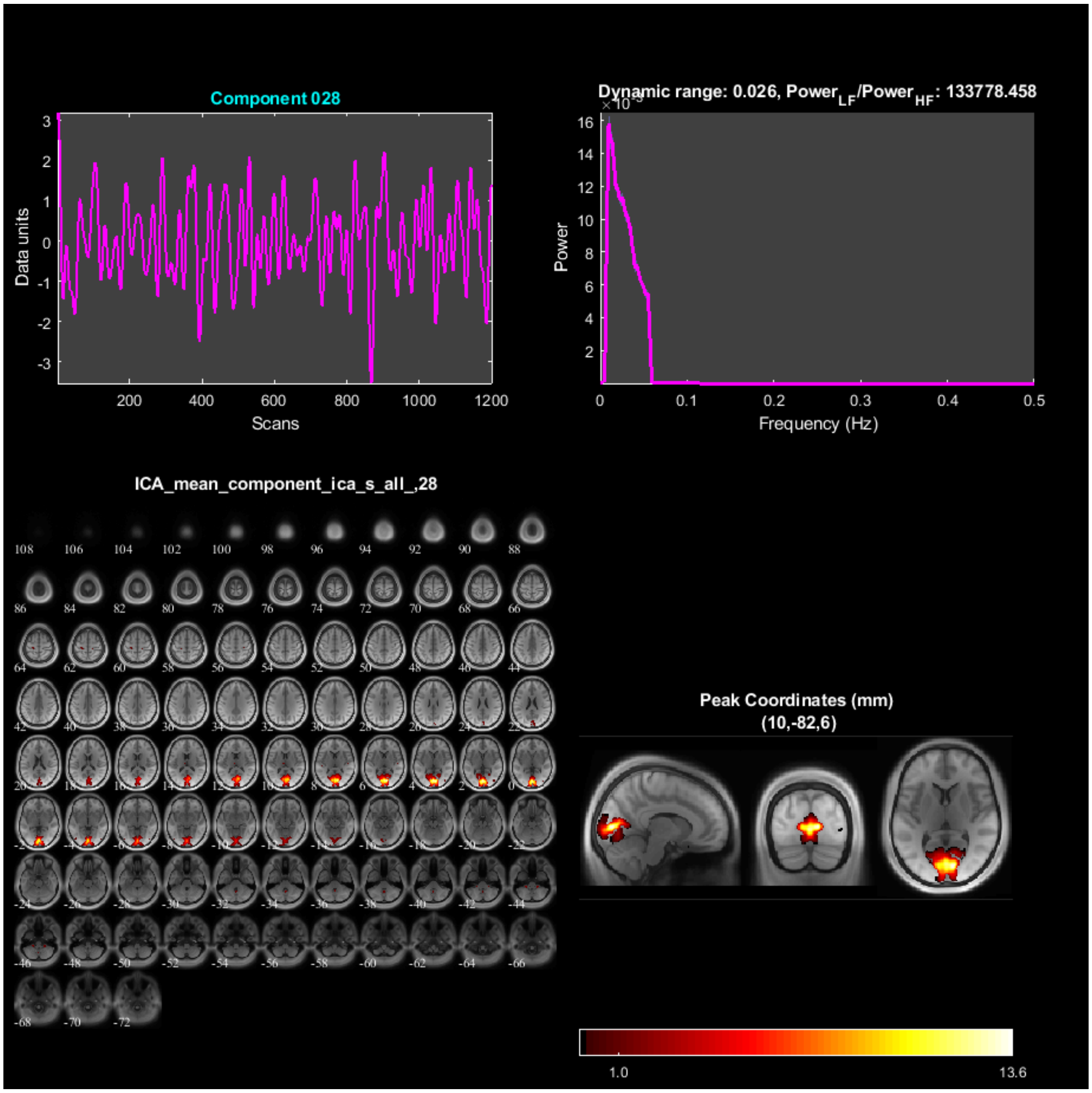

Component 028
Data units
Scans
Dynamic range: 0.026, Power_LF/Power_HF: 133778.458
Power
Frequency (Hz)
ICA_mean_component_ica_s_all_,28
Peak Coordinates (mm)
(10,-82,6)
1.0
13.6

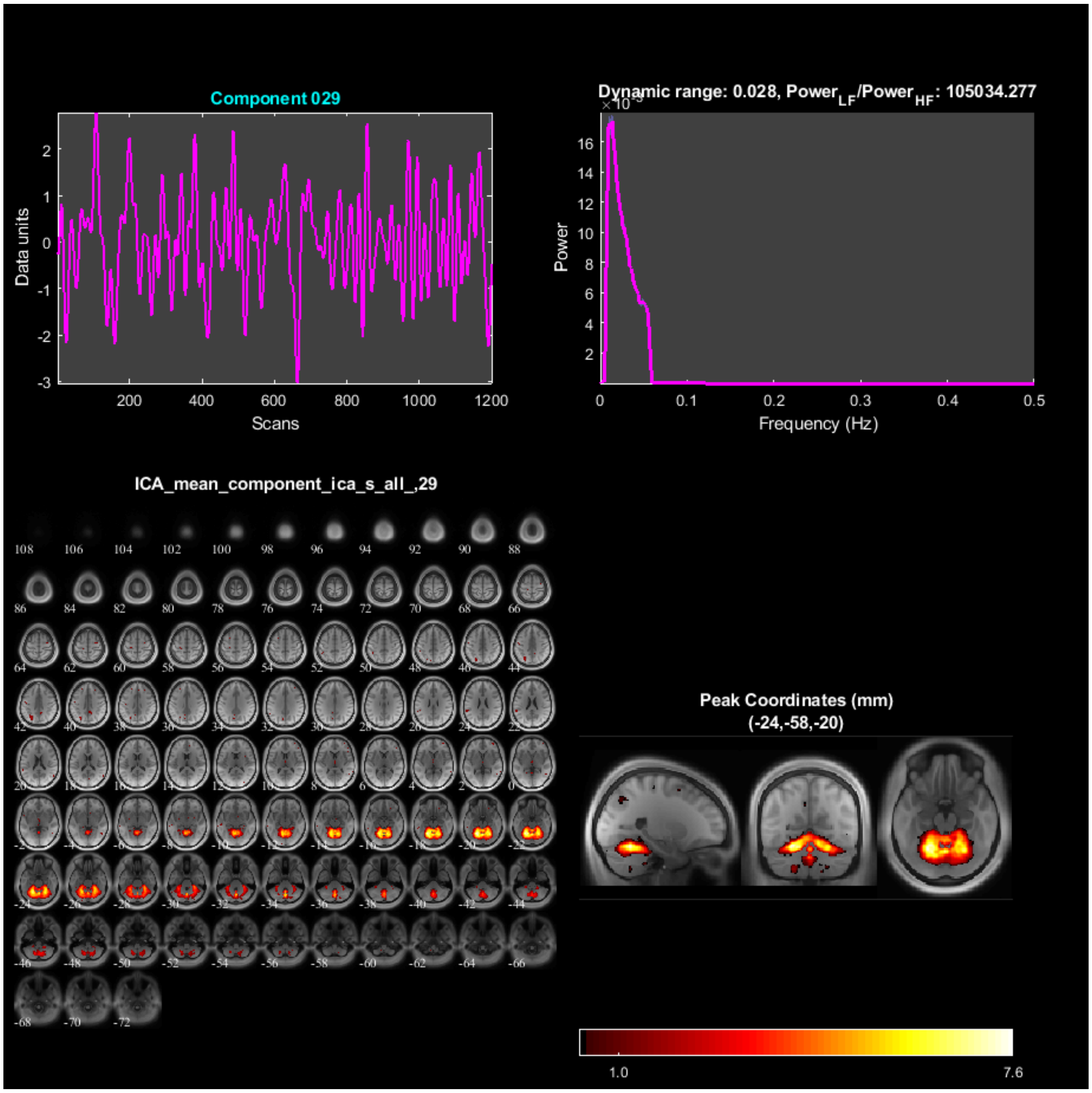

Component 029
Data units
Scans
Dynamic range: 0.028, Power$_{LF}$/Power$_{HF}$: 105034.277
Power
Frequency (Hz)
ICA_mean_component_ica_s_all_,29
Peak Coordinates (mm)
(-24,-58,-20)
1.0
7.6

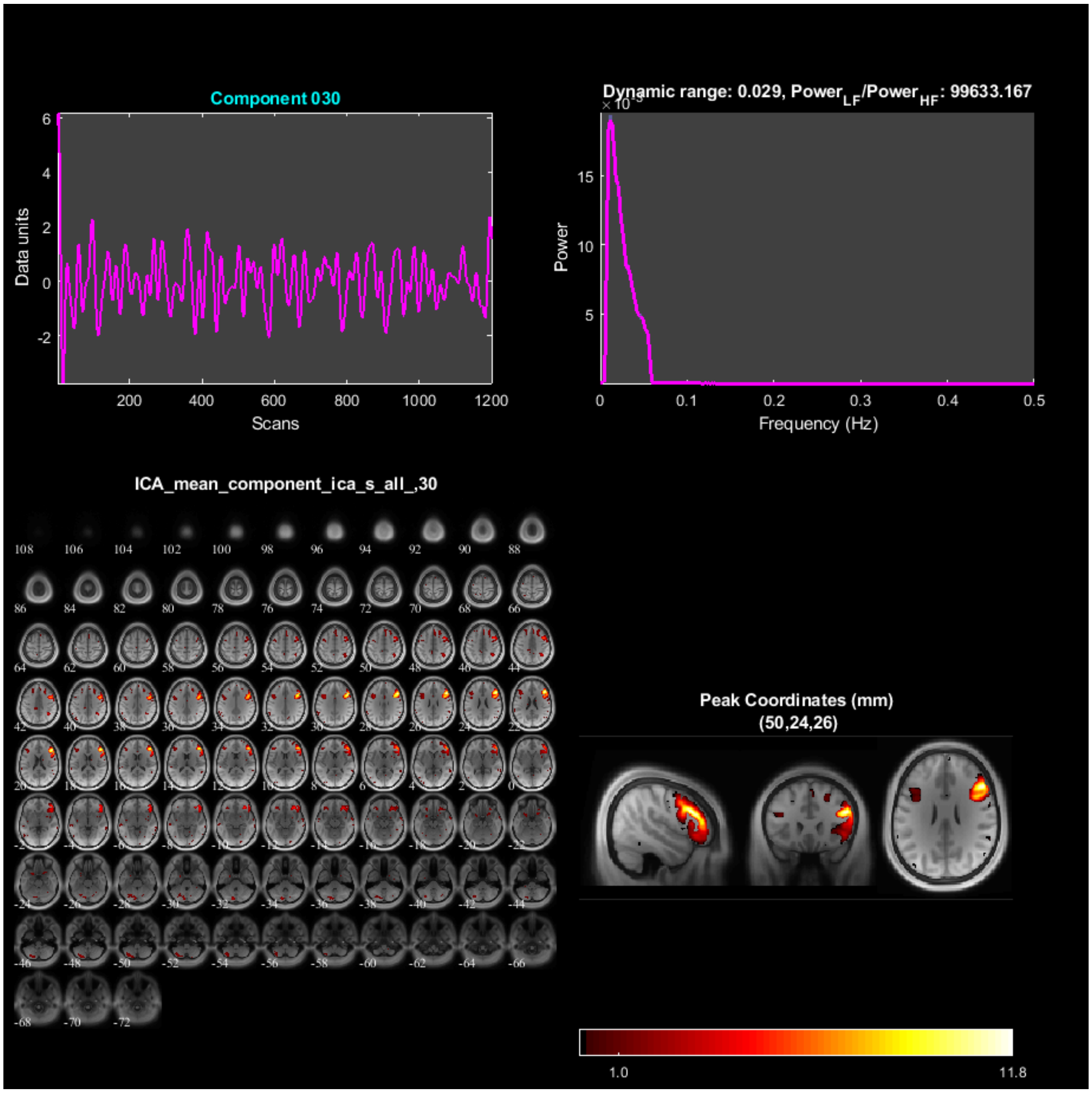
Component 030
Data units
Scans
Dynamic range: 0.029, Power$_{LF}$/Power$_{HF}$: 99633.167
Power
Frequency (Hz)
ICA_mean_component_ica_s_all_,30
Peak Coordinates (mm)
(50,24,26)
1.0
11.8

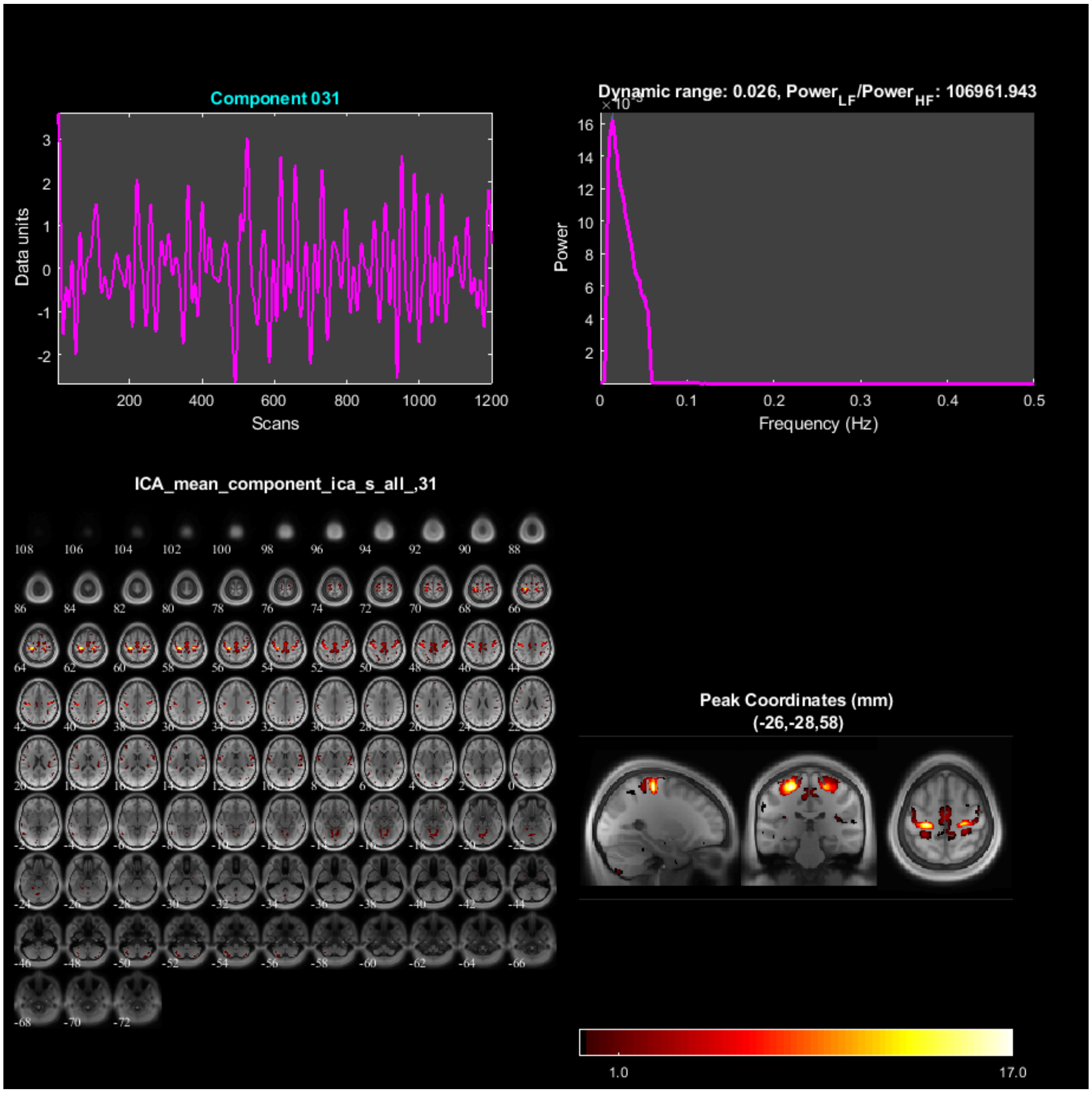

Component 031
Data units
Scans
Dynamic range: 0.026, Power$_{LF}$/Power$_{HF}$: 106961.943
Power
Frequency (Hz)
ICA_mean_component_ica_s_all_,31
Peak Coordinates (mm)
(-26,-28,58)
1.0
17.0

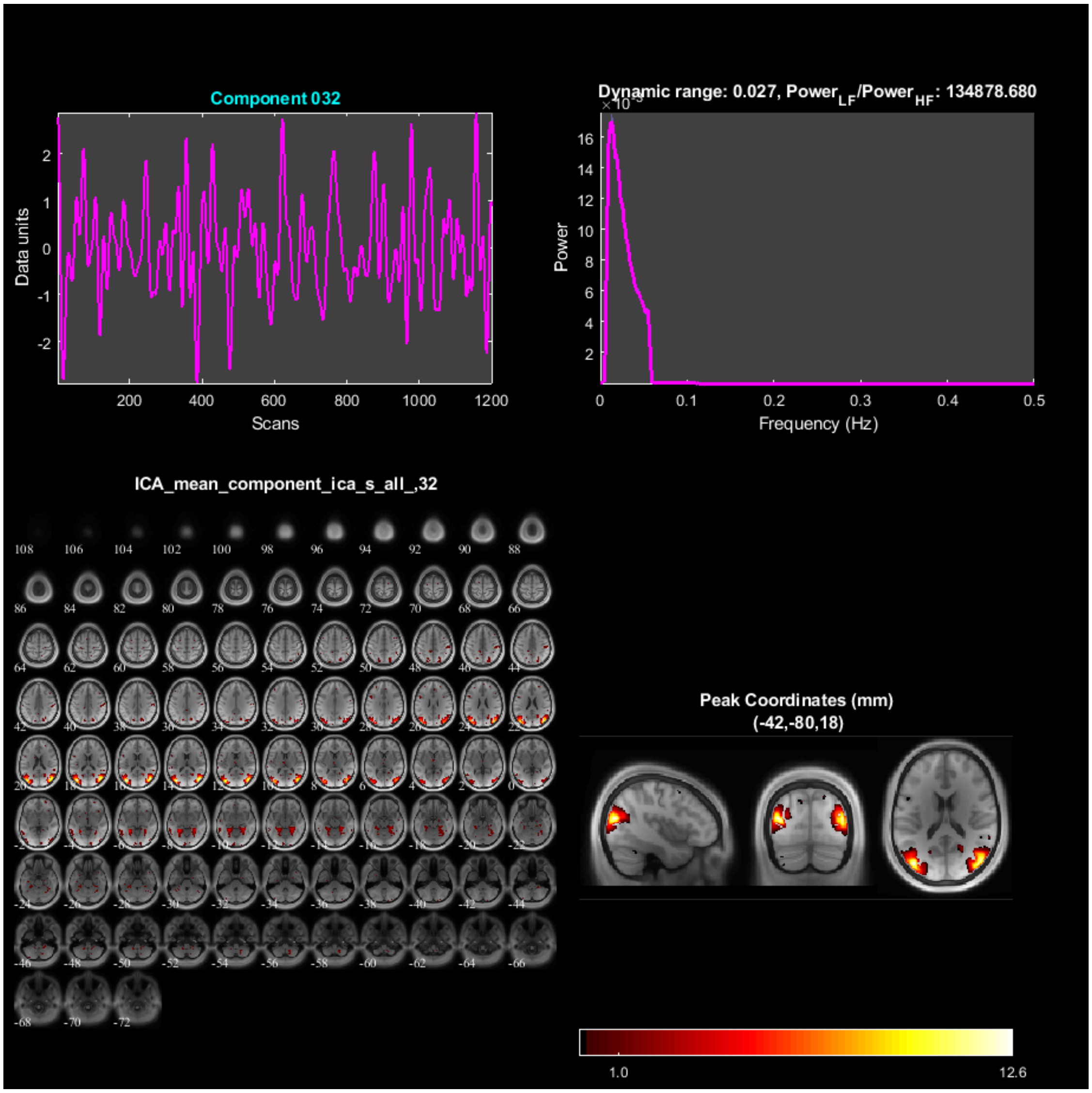
Component 032
Data units
Scans
Dynamic range: 0.027, Power_LF/Power_HF: 134878.680
Power
Frequency (Hz)
ICA_mean_component_ica_s_all_,32
Peak Coordinates (mm)
(-42,-80,18)
1.0
12.6

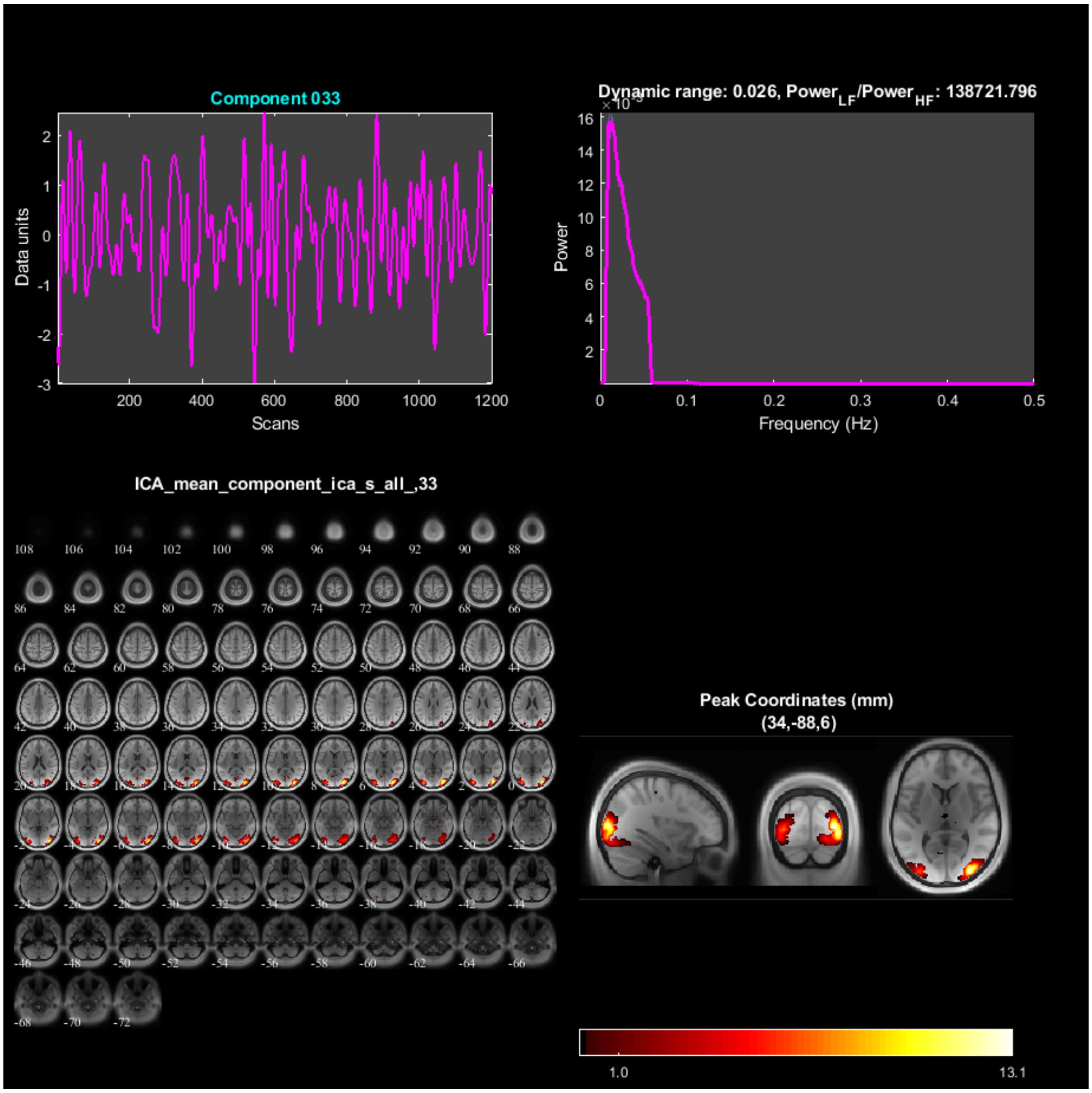

Component 033
Data units
Scans
Dynamic range: 0.026, Power$_{LF}$/Power$_{HF}$: 138721.796
Power
Frequency (Hz)
ICA_mean_component_ica_s_all_,33
Peak Coordinates (mm)
(34,-88,6)
1.0
13.1

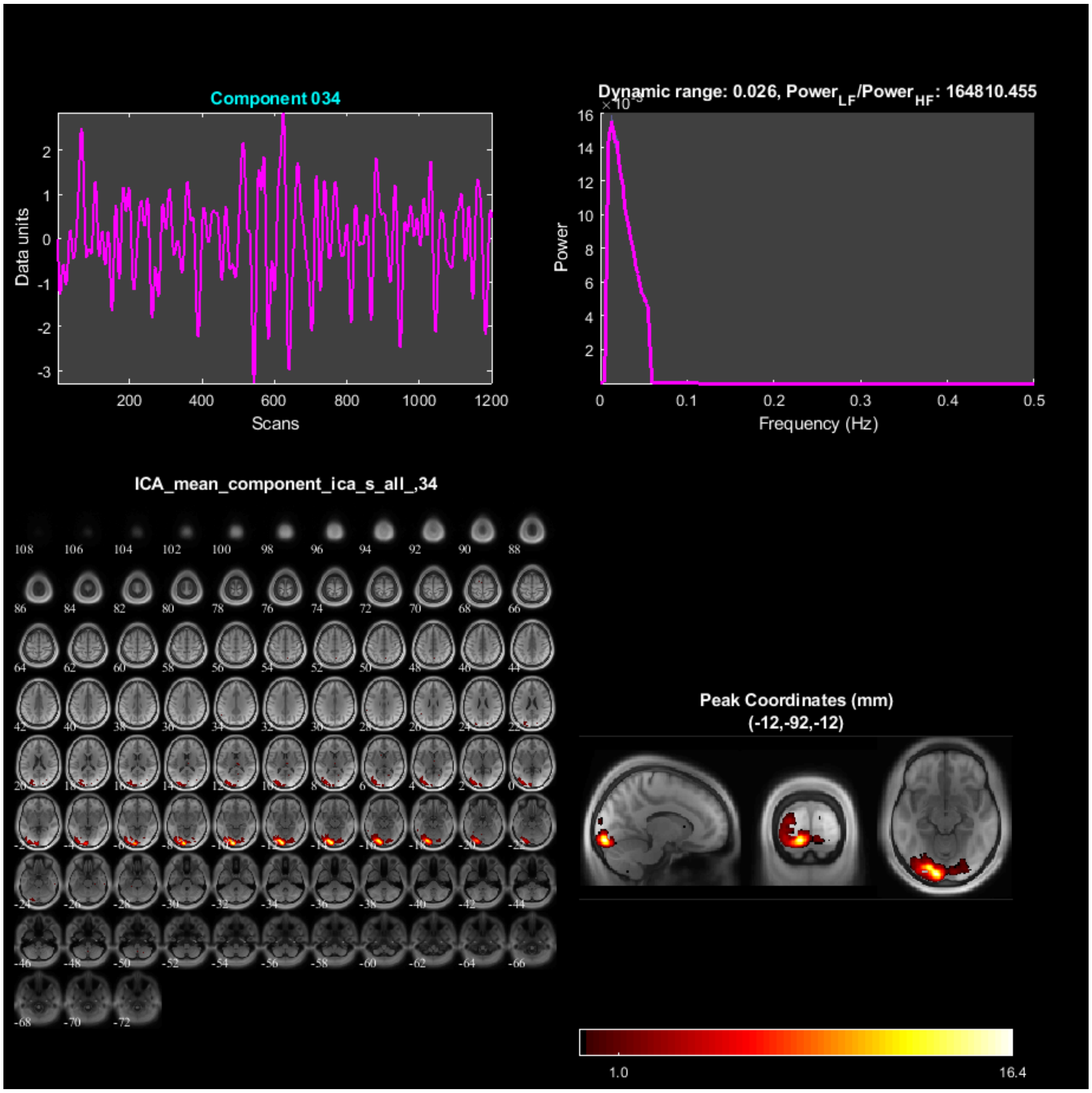

Component 034
Data units
Scans
Dynamic range: 0.026, Power_LF/Power_HF: 164810.455
Power
Frequency (Hz)
ICA_mean_component_ica_s_all_,34
Peak Coordinates (mm)
(-12,-92,-12)
1.0
16.4

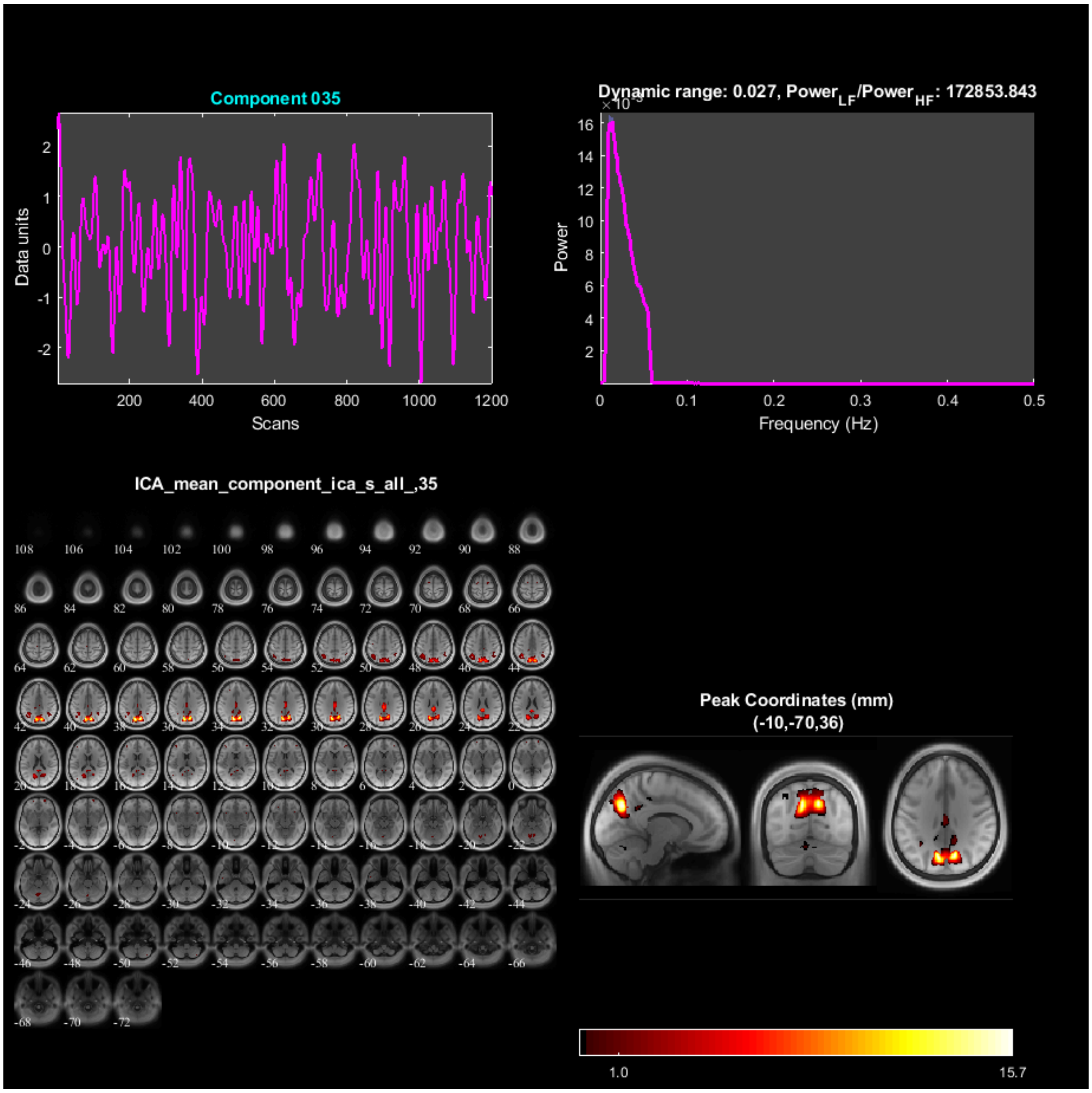
Component 035
Data units
Scans
Dynamic range: 0.027, Power$_{LF}$/Power$_{HF}$: 172853.843
Power
Frequency (Hz)
ICA_mean_component_ica_s_all_,35
Peak Coordinates (mm)
(-10,-70,36)
1.0
15.7

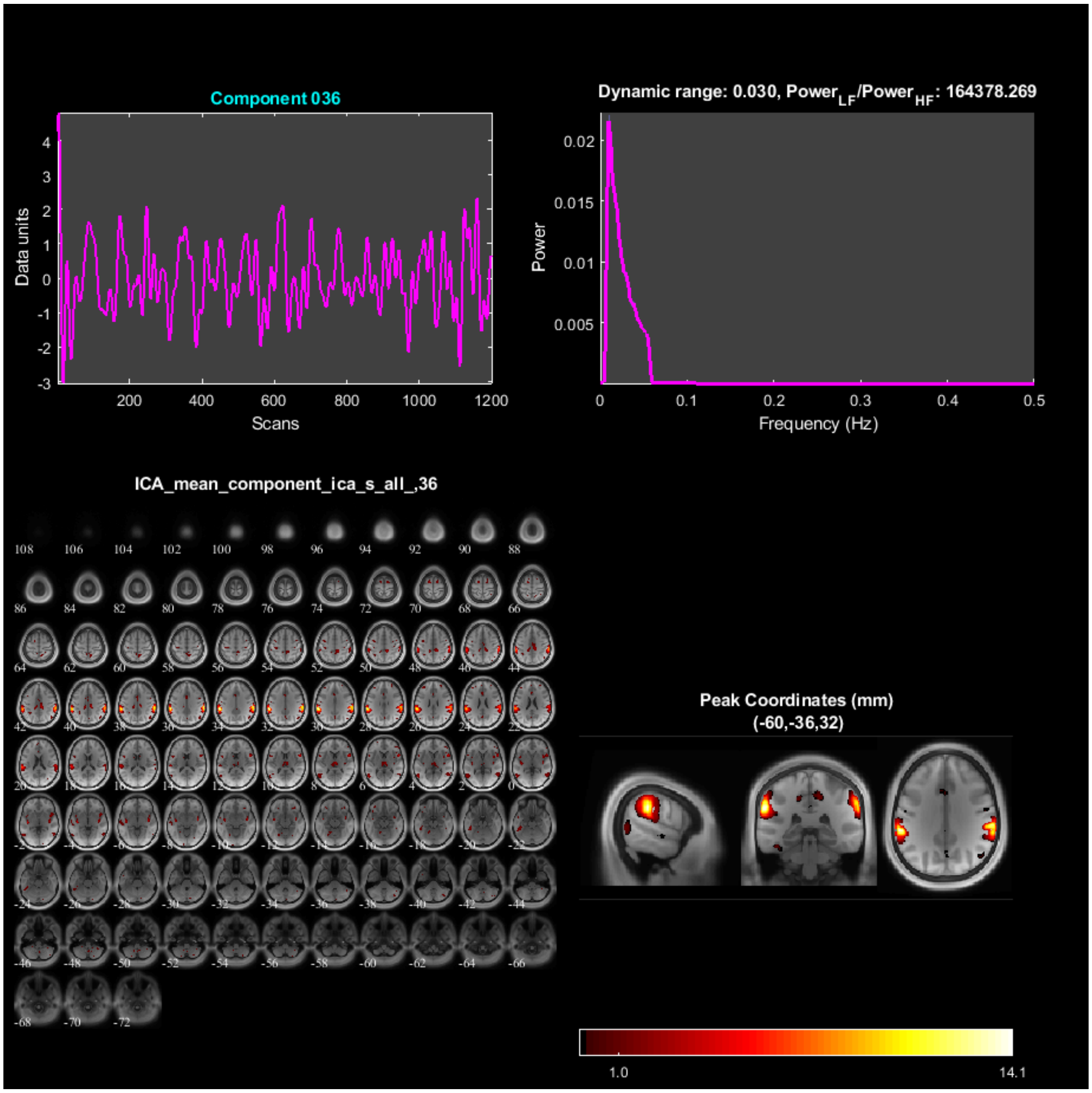
Component 036
Data units
Scans
Dynamic range: 0.030, Power$_{LF}$/Power$_{HF}$: 164378.269
Power
Frequency (Hz)
ICA_mean_component_ica_s_all_,36
Peak Coordinates (mm)
(-60,-36,32)
1.0
14.1

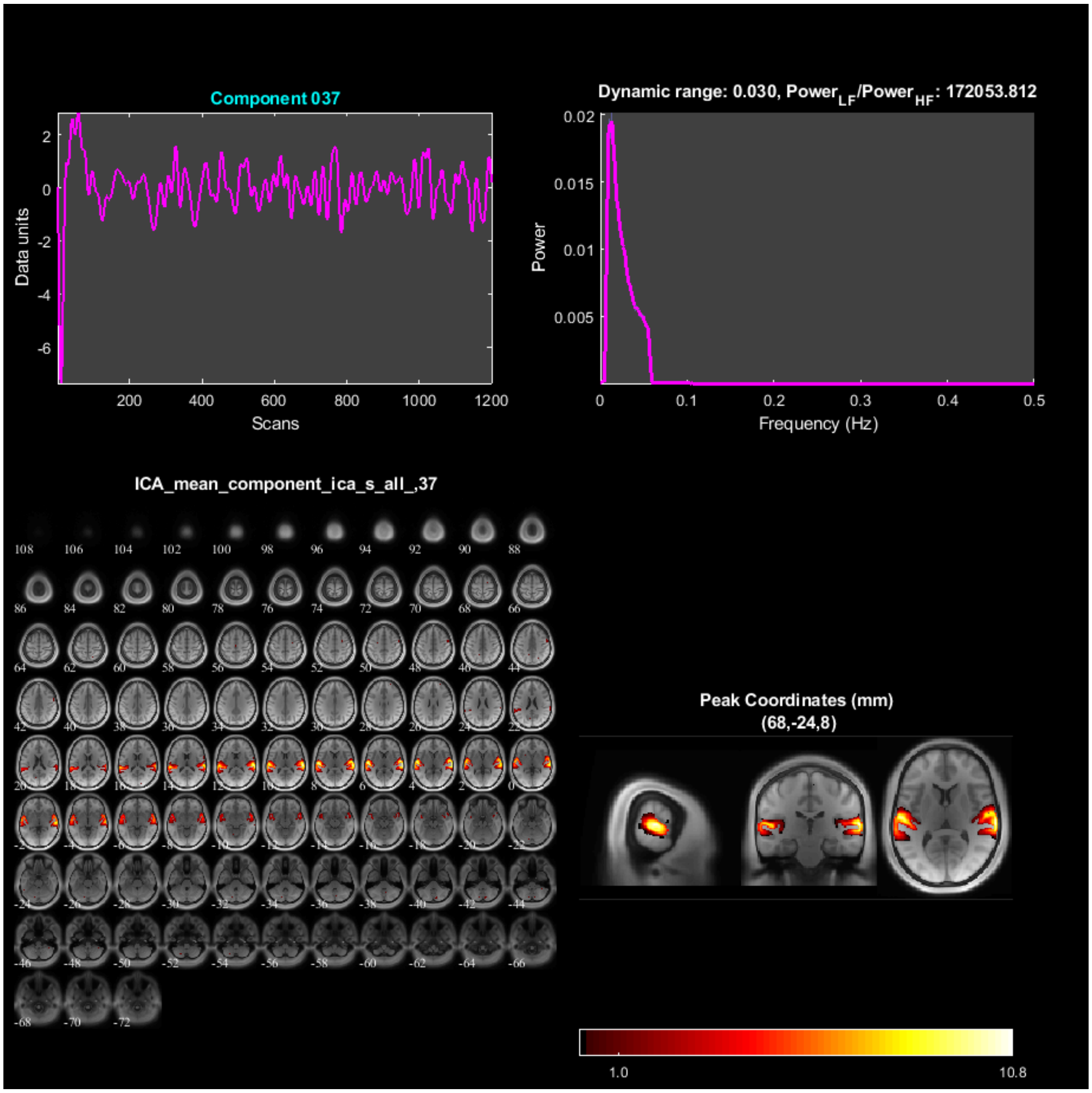

Component 037
Data units
Scans
Dynamic range: 0.030, Power$_{LF}$/Power$_{HF}$: 172053.812
Power
Frequency (Hz)
ICA_mean_component_ica_s_all_,37
Peak Coordinates (mm)
(68,-24,8)
1.0
10.8

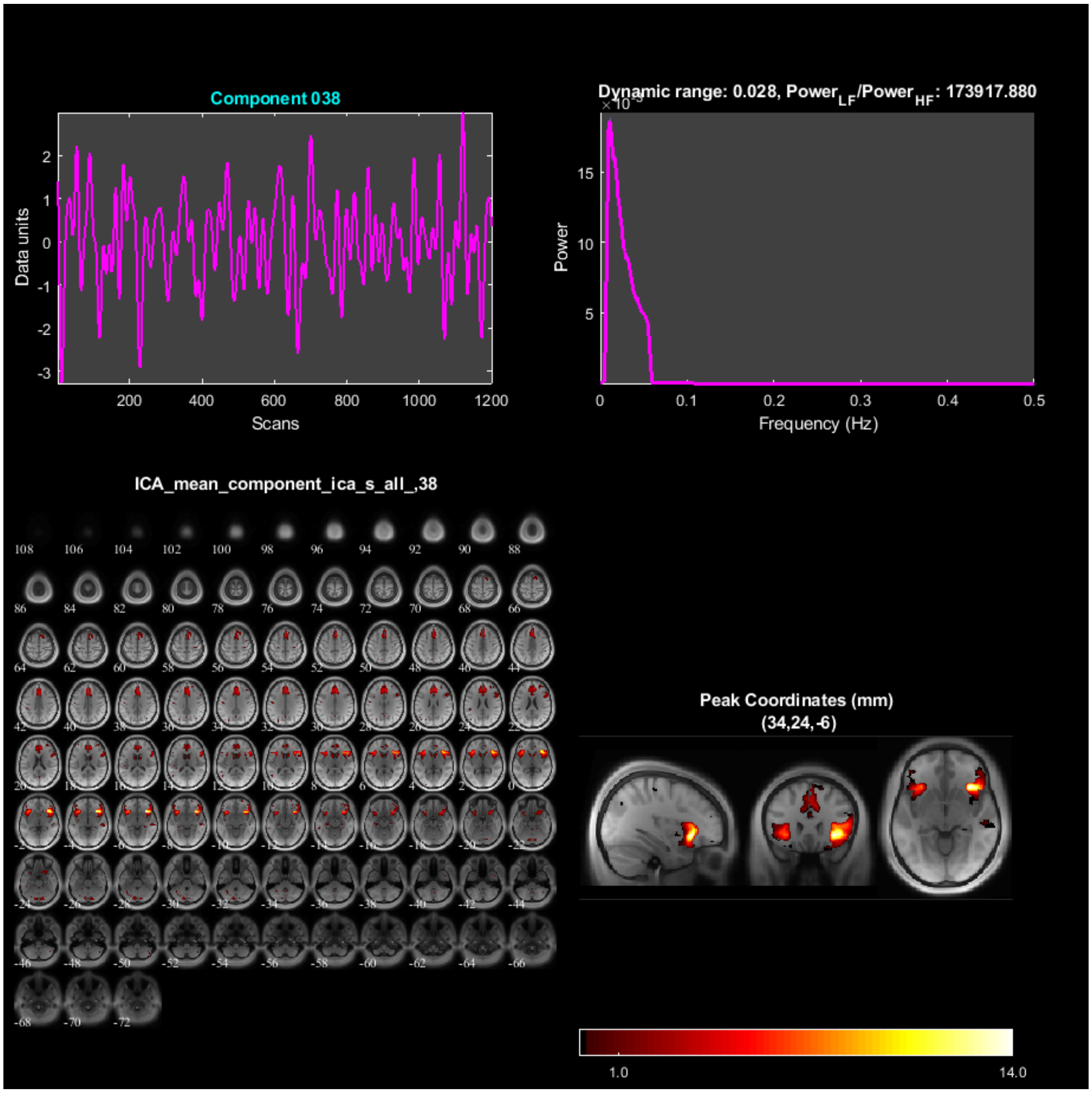

Component 038
Data units
Scans
Dynamic range: 0.028, Power$_{LF}$/Power$_{HF}$: 173917.880
Power
Frequency (Hz)
ICA_mean_component_ica_s_all_,38
Peak Coordinates (mm)
(34,24,-6)
1.0
14.0

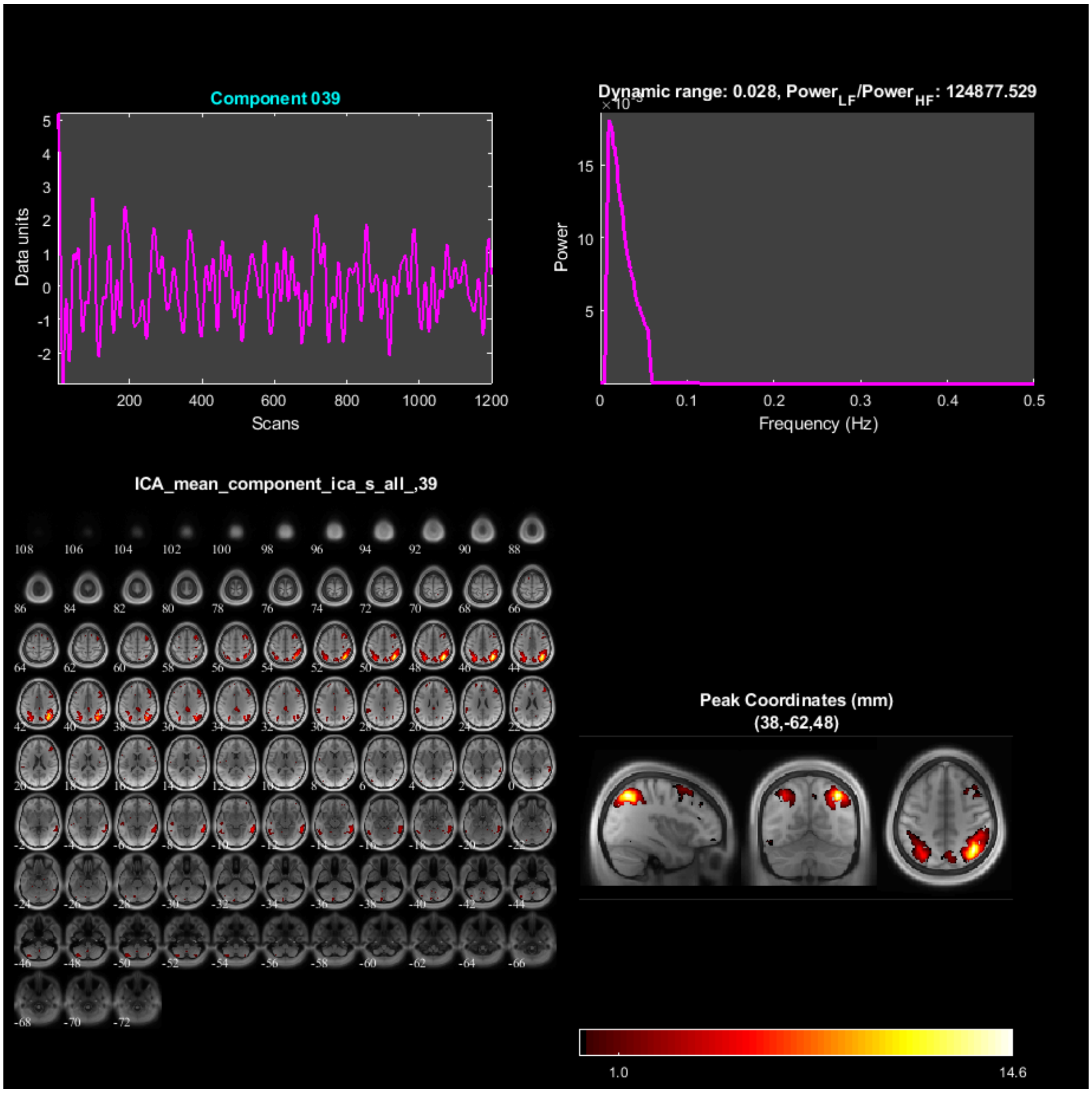

Component 039
Data units
Scans
Dynamic range: 0.028, Power$_{LF}$/Power$_{HF}$: 124877.529
Power
Frequency (Hz)
ICA_mean_component_ica_s_all_,39
Peak Coordinates (mm)
(38,-62,48)
1.0
14.6

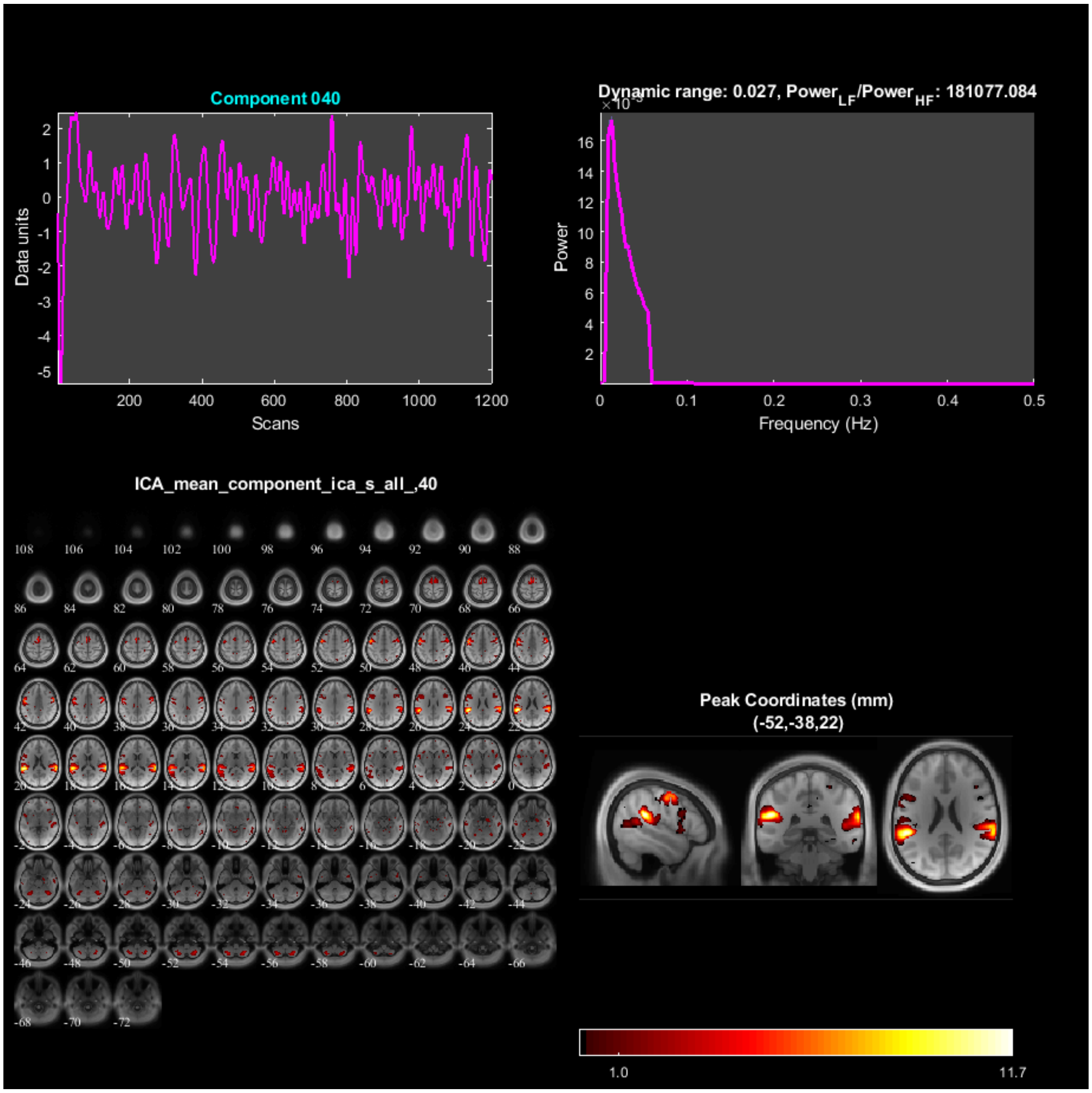
Component 040
Data units
Scans
Dynamic range: 0.027, Power$_{LF}$/Power$_{HF}$: 181077.084
Power
Frequency (Hz)
ICA_mean_component_ica_s_all_,40
Peak Coordinates (mm)
(-52,-38,22)
1.0
11.7

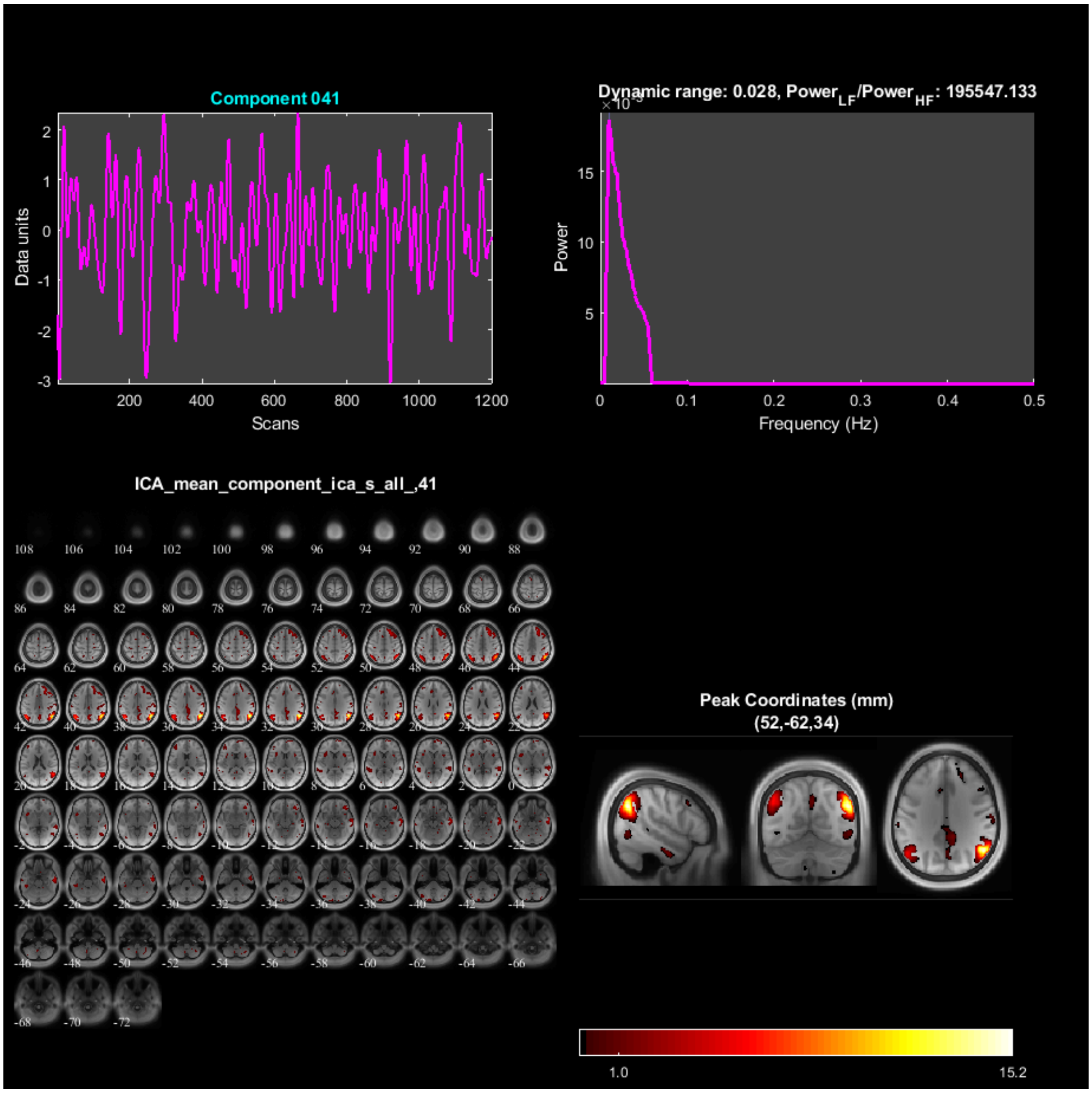

Component 041
Data units
Scans
Dynamic range: 0.028, Power$_{LF}$/Power$_{HF}$: 195547.133
Power
Frequency (Hz)
ICA_mean_component_ica_s_all_,41
Peak Coordinates (mm)
(52,-62,34)
1.0
15.2

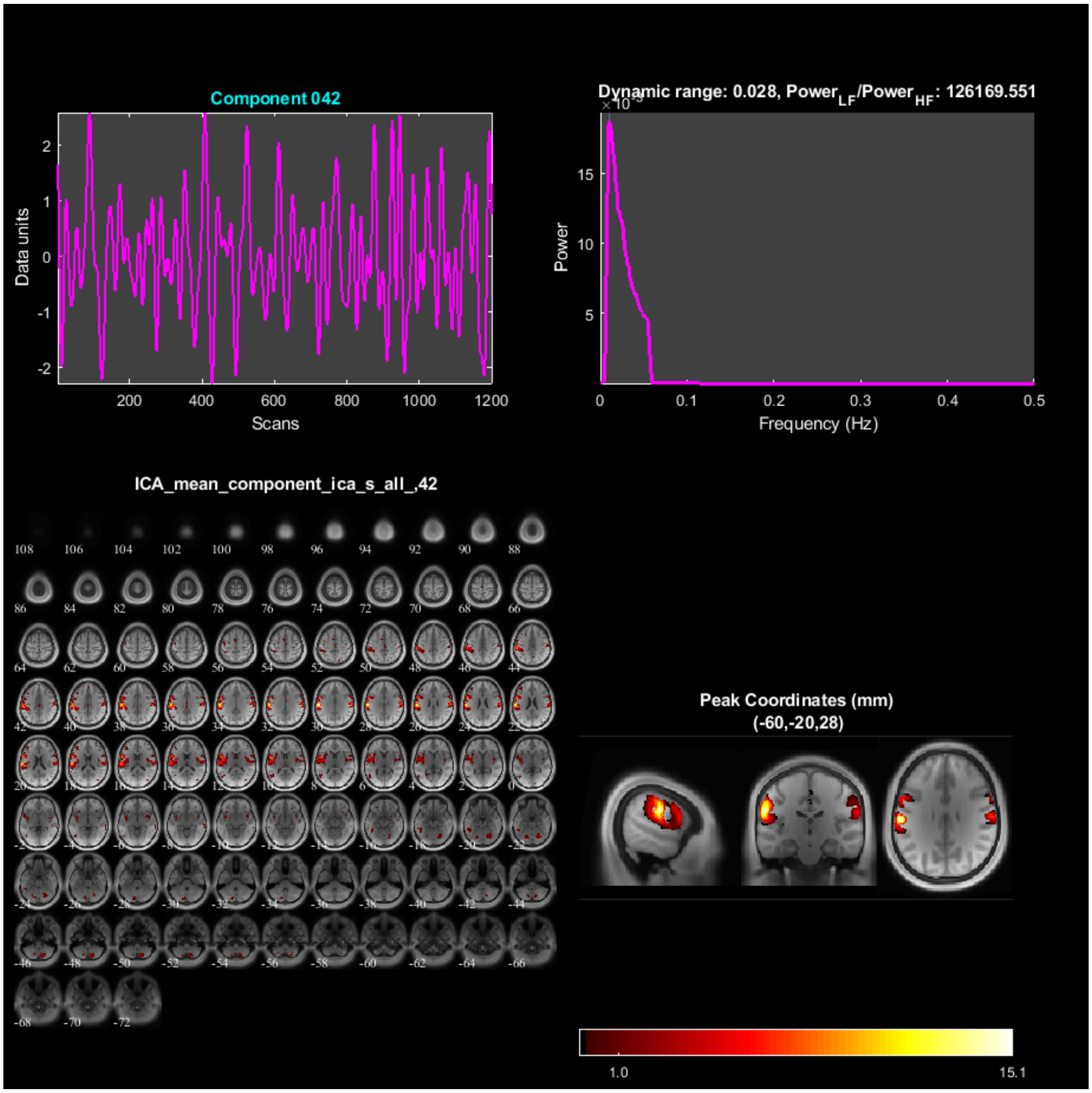

Component 042
Data units
Scans
Dynamic range: 0.028, Power$_{LF}$/Power$_{HF}$: 126169.551
Power
Frequency (Hz)
ICA_mean_component_ica_s_all_,42
Peak Coordinates (mm)
(-60,-20,28)
1.0
15.1

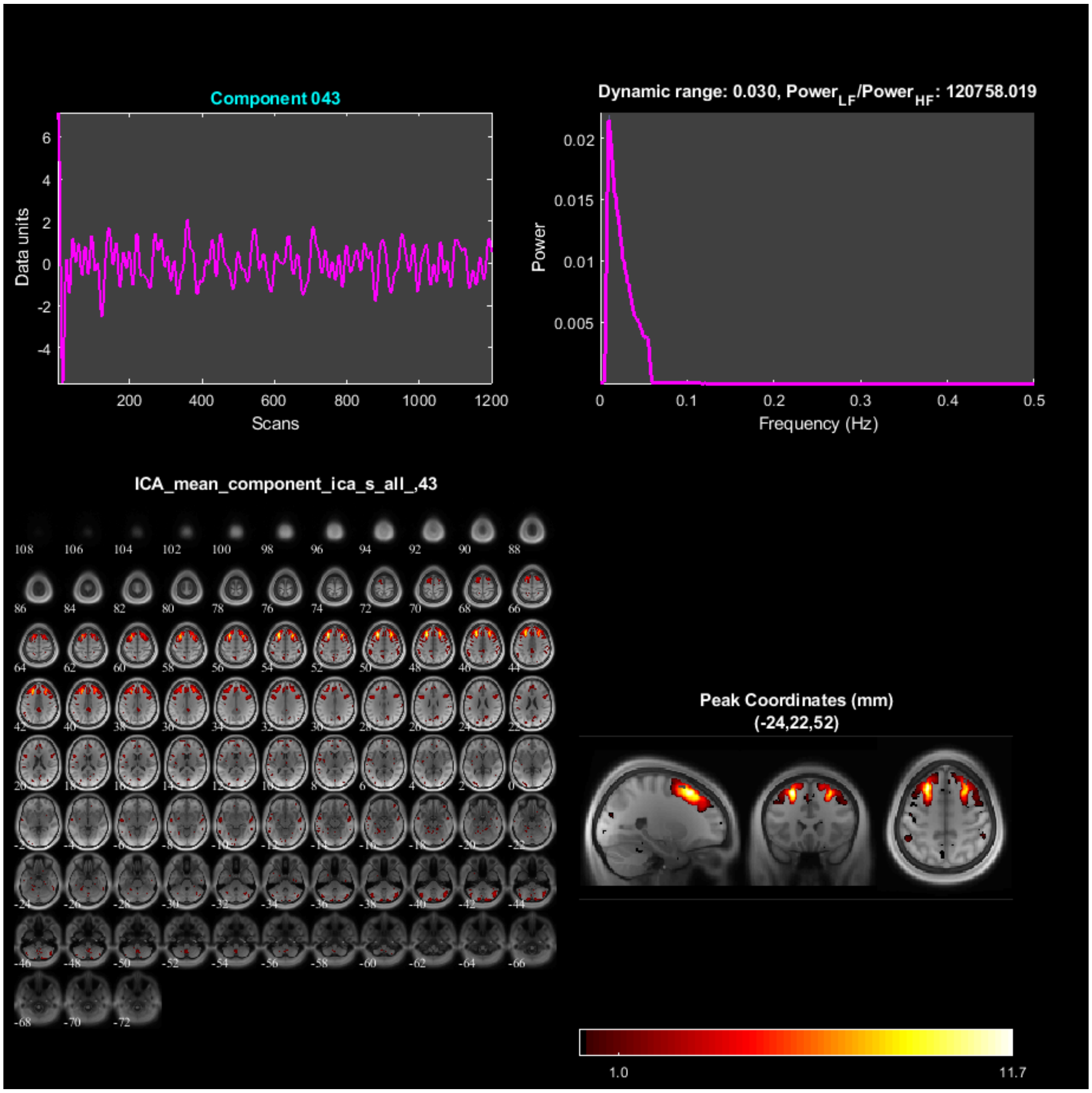
Component 043
Data units
Scans
Dynamic range: 0.030, Power$_{LF}$/Power$_{HF}$: 120758.019
Power
Frequency (Hz)
ICA_mean_component_ica_s_all_,43
Peak Coordinates (mm)
(-24,22,52)
1.0
11.7

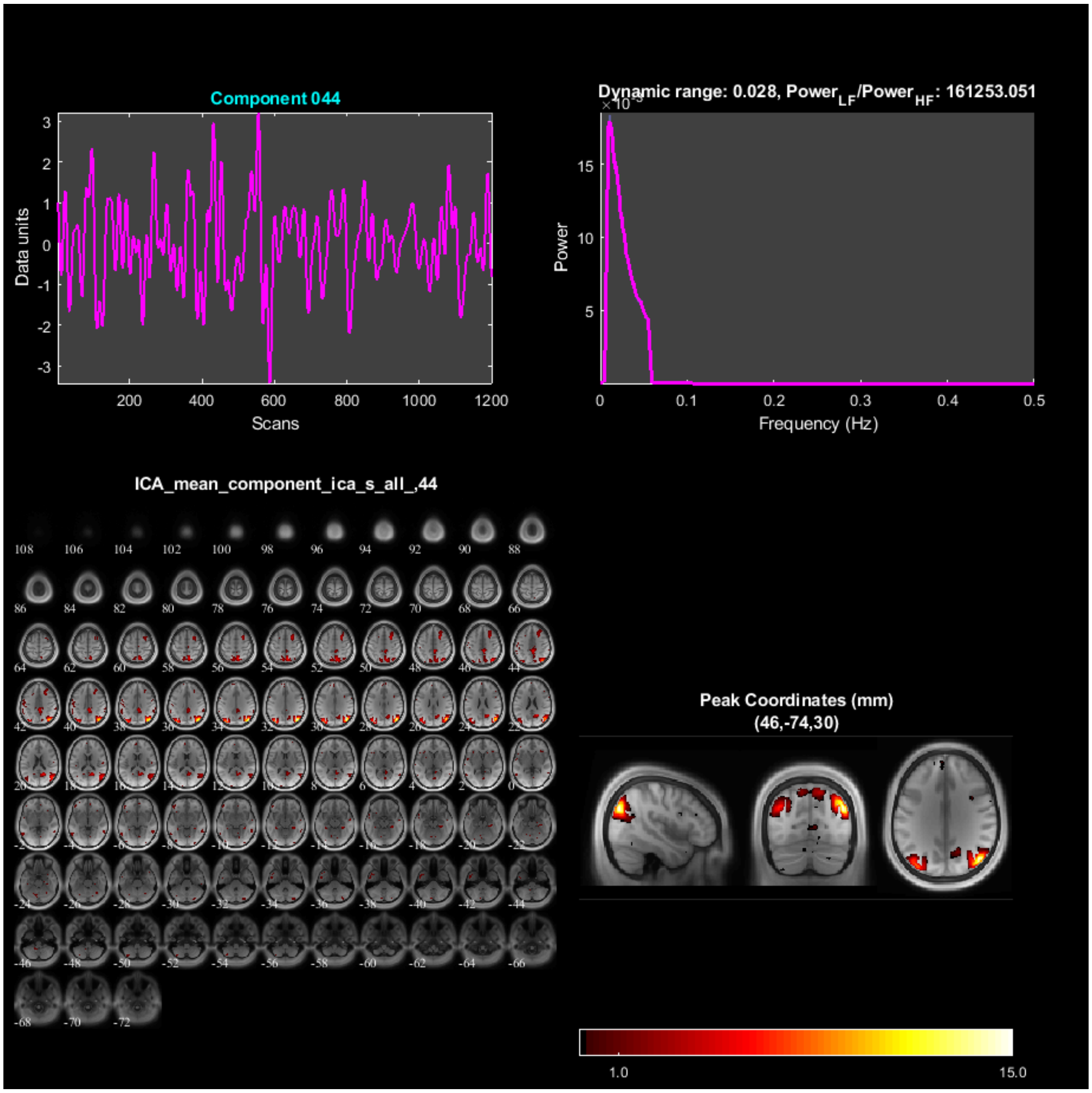
Component 044
Data units
Scans
Dynamic range: 0.028, Power$_{LF}$/Power$_{HF}$: 161253.051
Power
Frequency (Hz)
ICA_mean_component_ica_s_all_,44
Peak Coordinates (mm)
(46,-74,30)
1.0
15.0

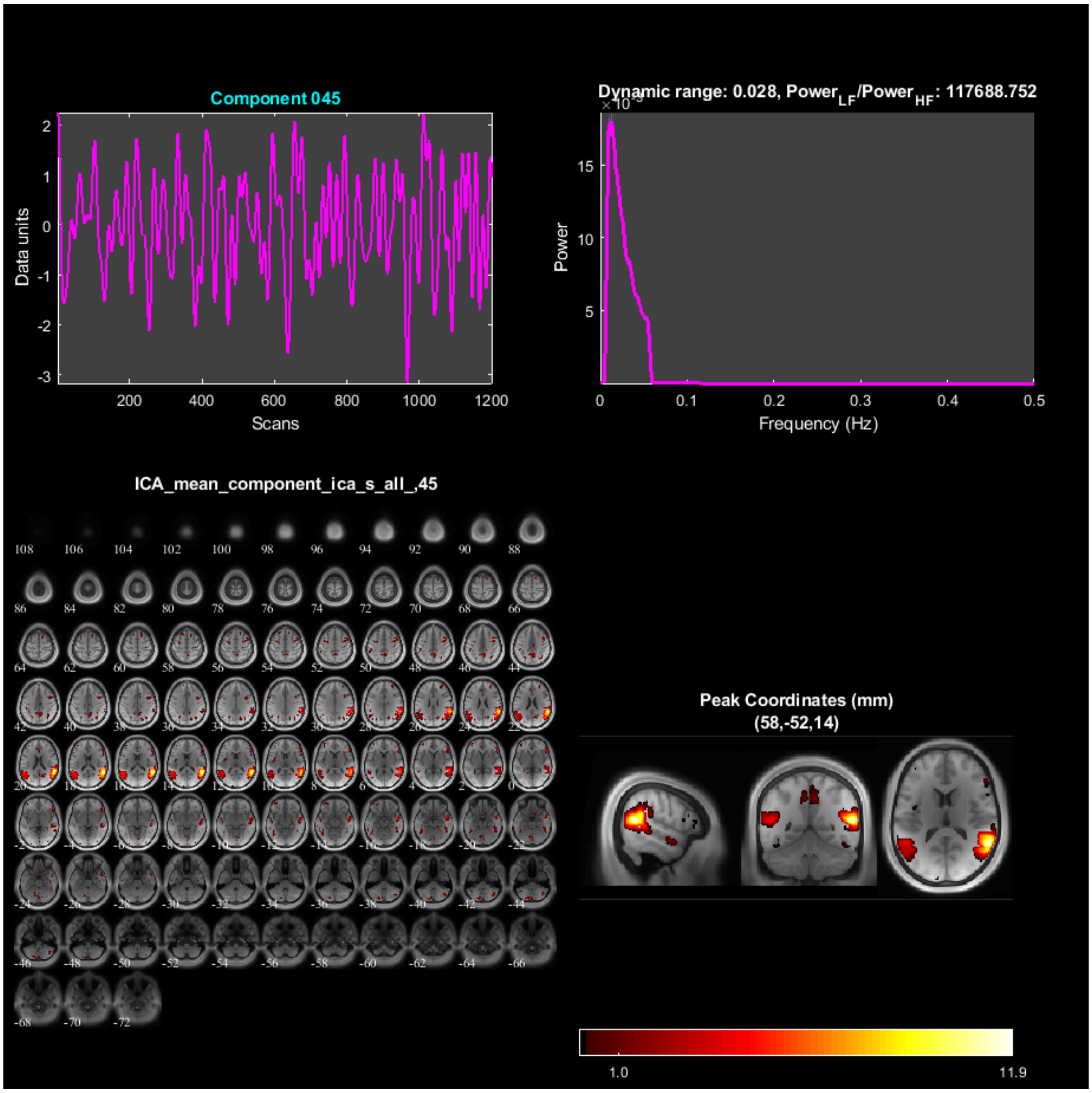
Component 045
Data units
Scans
Dynamic range: 0.028, Power$_{LF}$/Power$_{HF}$: 117688.752
Power
Frequency (Hz)
ICA_mean_component_ica_s_all_,45
Peak Coordinates (mm)
(58,-52,14)
1.0
11.9

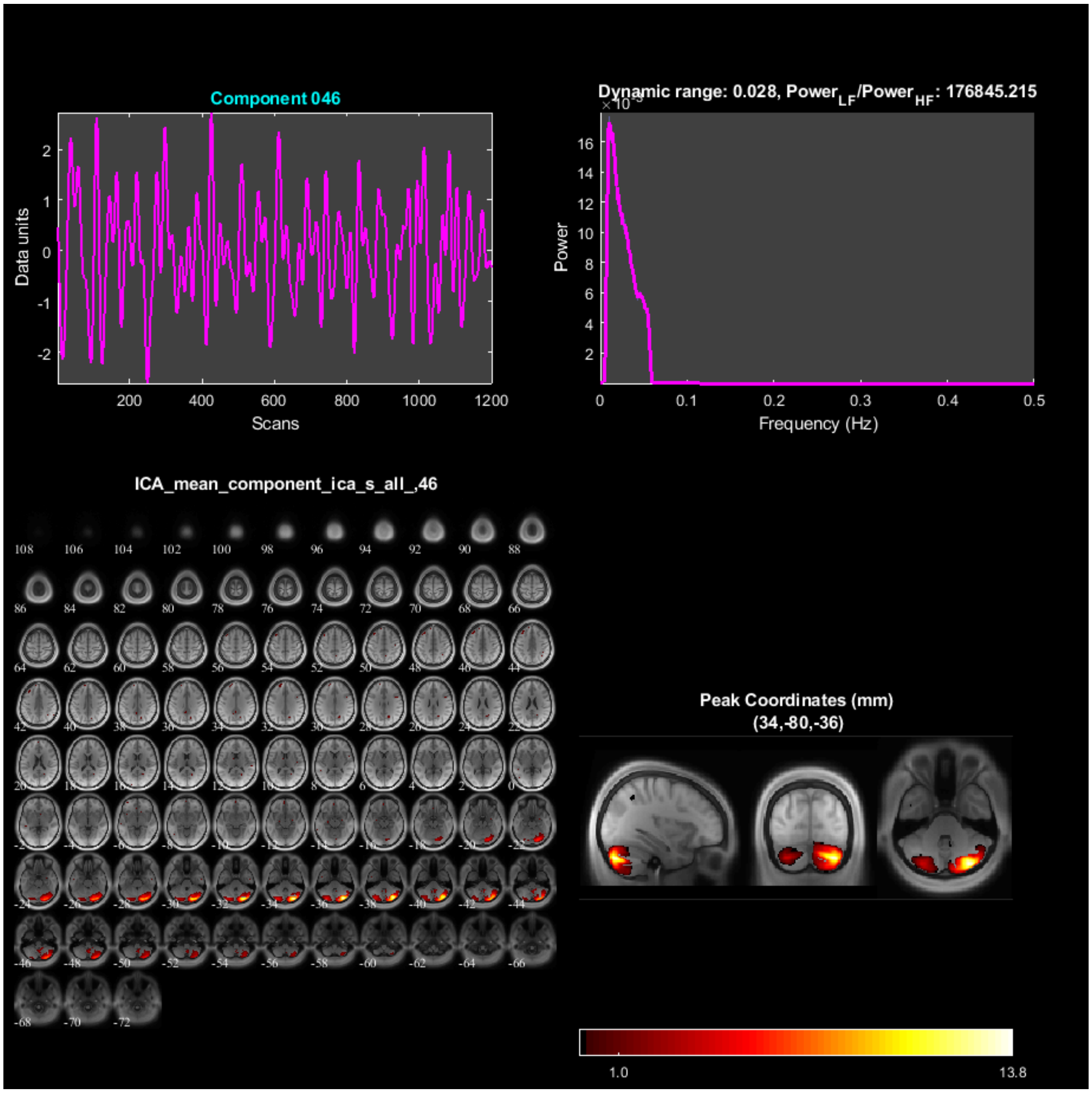
Component 046
Data units
Scans
Dynamic range: 0.028, Power_LF/Power_HF: 176845.215
Power
Frequency (Hz)
ICA_mean_component_ica_s_all_,46
Peak Coordinates (mm)
(34,-80,-36)
1.0
13.8

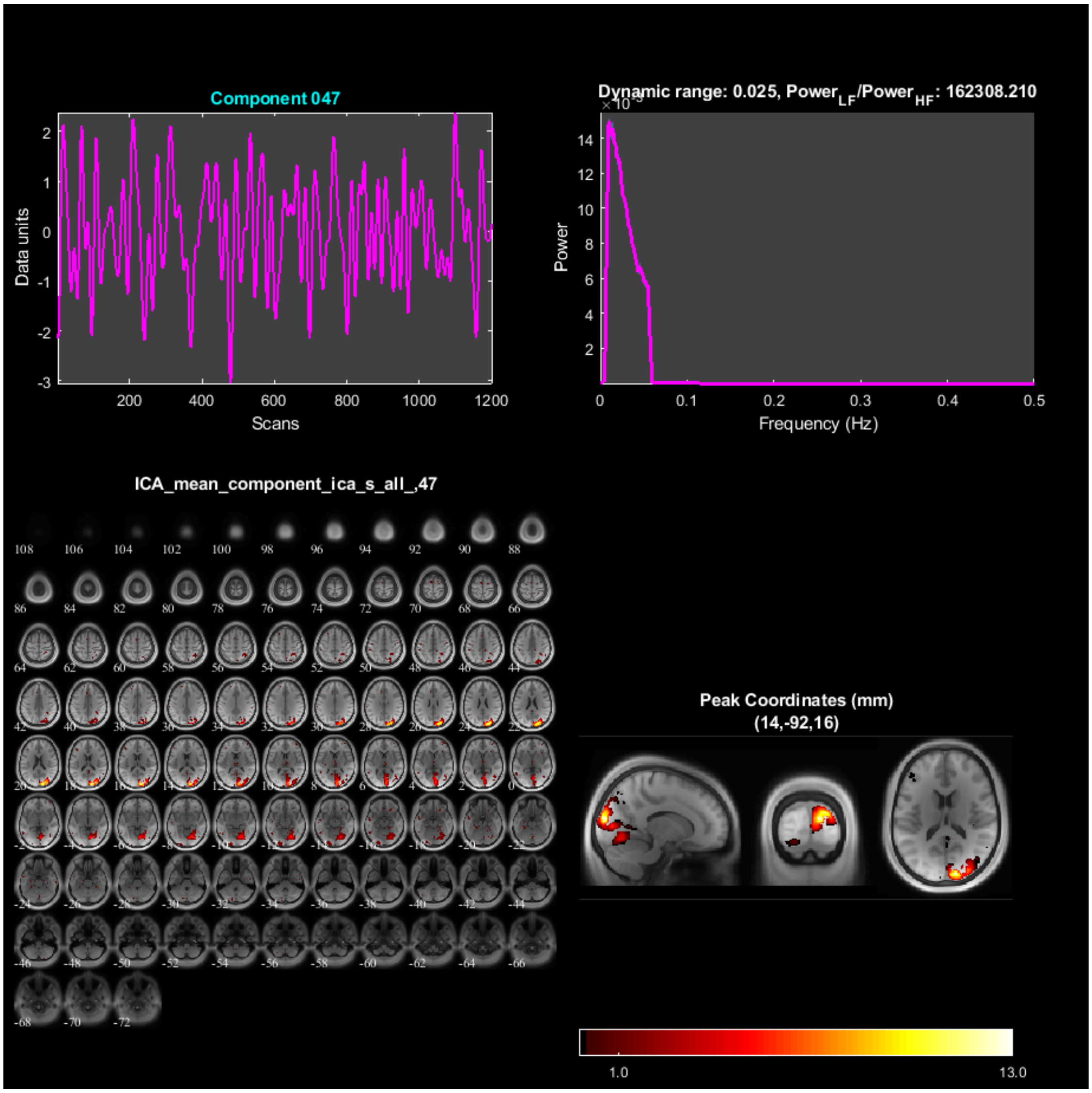
Component 047
Data units
Scans
Dynamic range: 0.025, Power$_{LF}$/Power$_{HF}$: 162308.210
Power
Frequency (Hz)
ICA_mean_component_ica_s_all_,47
Peak Coordinates (mm)
(14,-92,16)
1.0
13.0

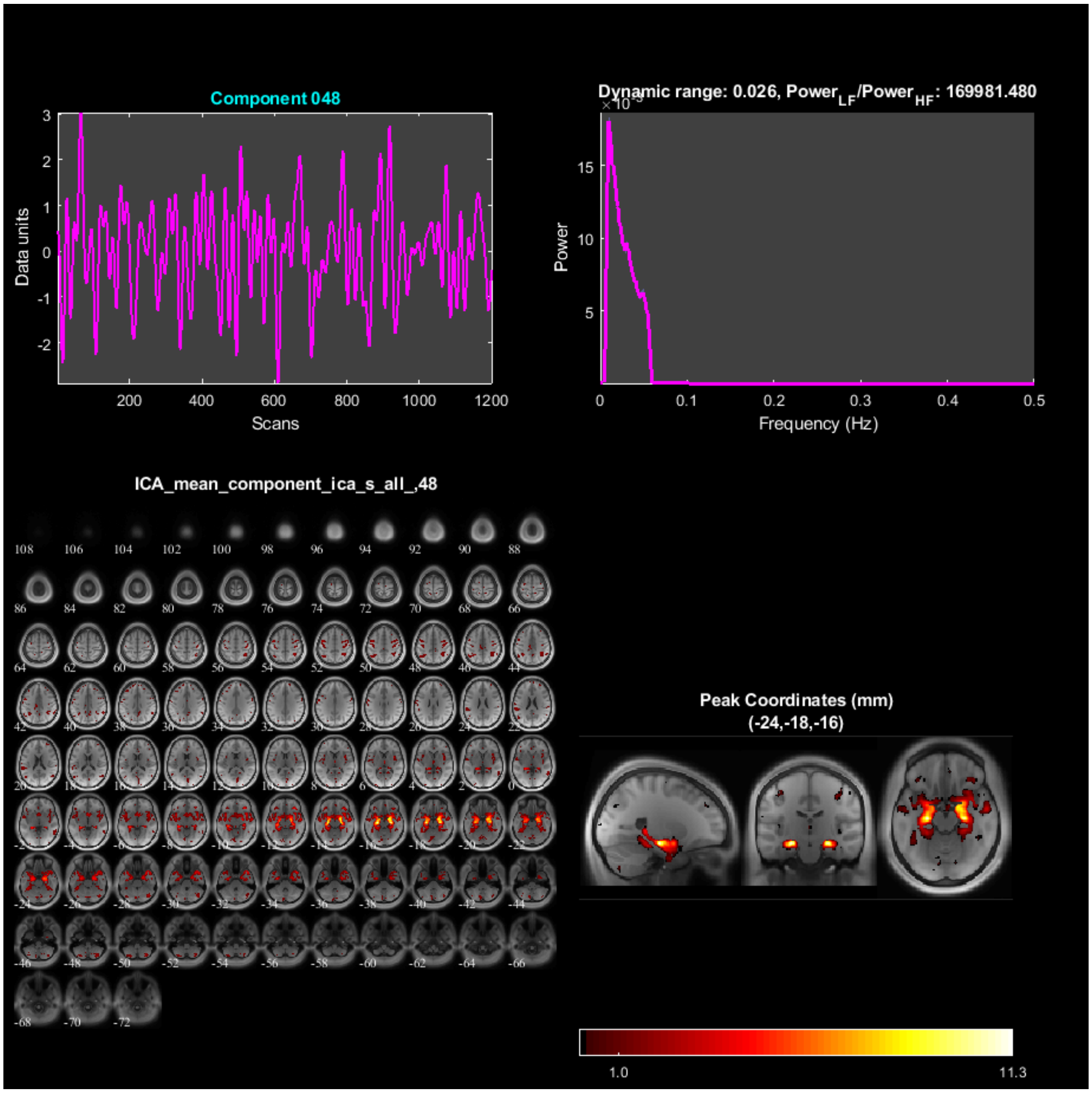
Component 048
Data units
Scans
Dynamic range: 0.026, Power_LF/Power_HF: 169981.480
Power
Frequency (Hz)
ICA_mean_component_ica_s_all_,48
Peak Coordinates (mm)
(-24,-18,-16)
1.0
11.3

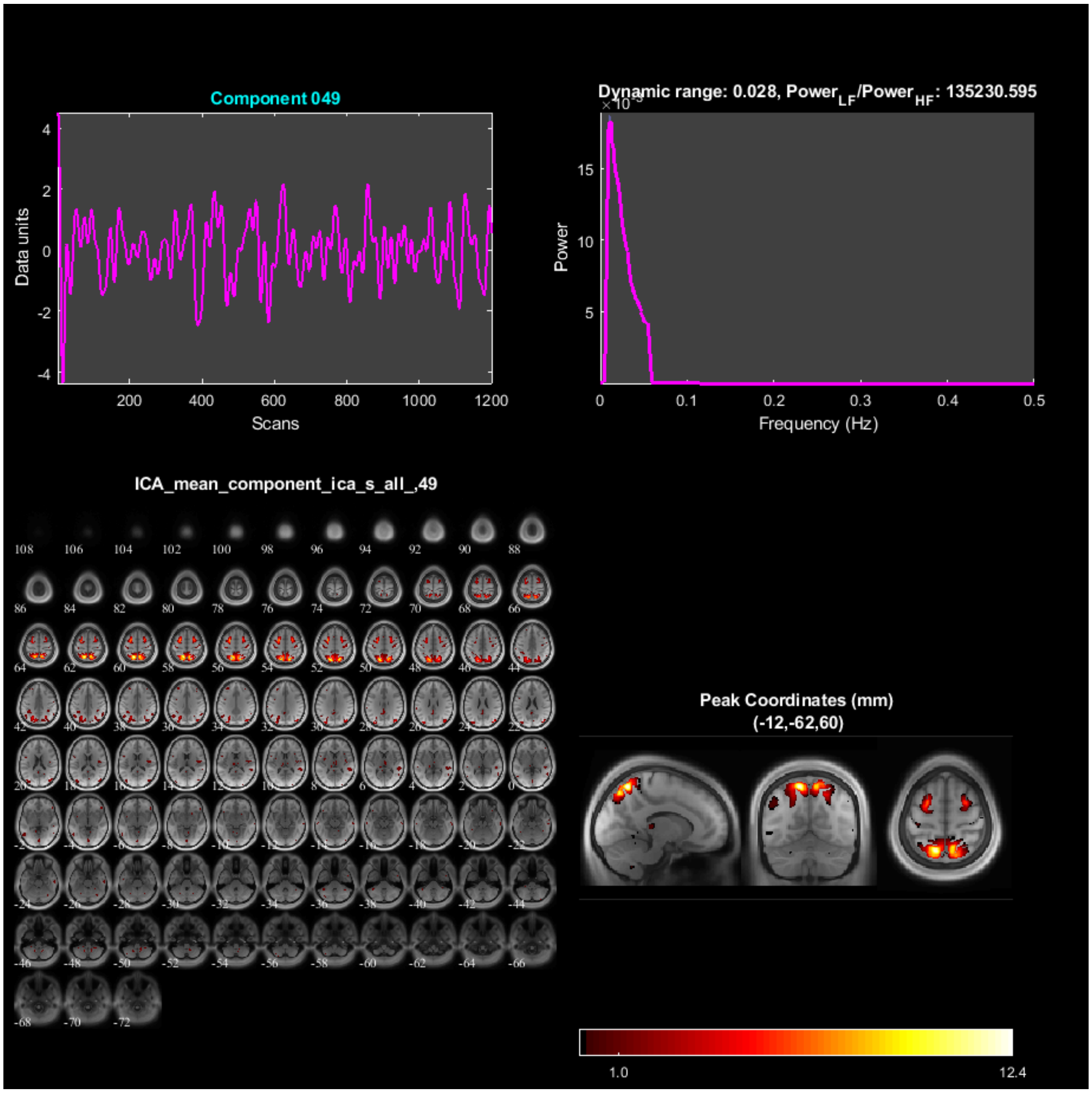

Component 049
Data units
Scans
Dynamic range: 0.028, Power_LF/Power_HF: 135230.595
Power
Frequency (Hz)
ICA_mean_component_ica_s_all_,49
Peak Coordinates (mm)
(-12,-62,60)
1.0
12.4

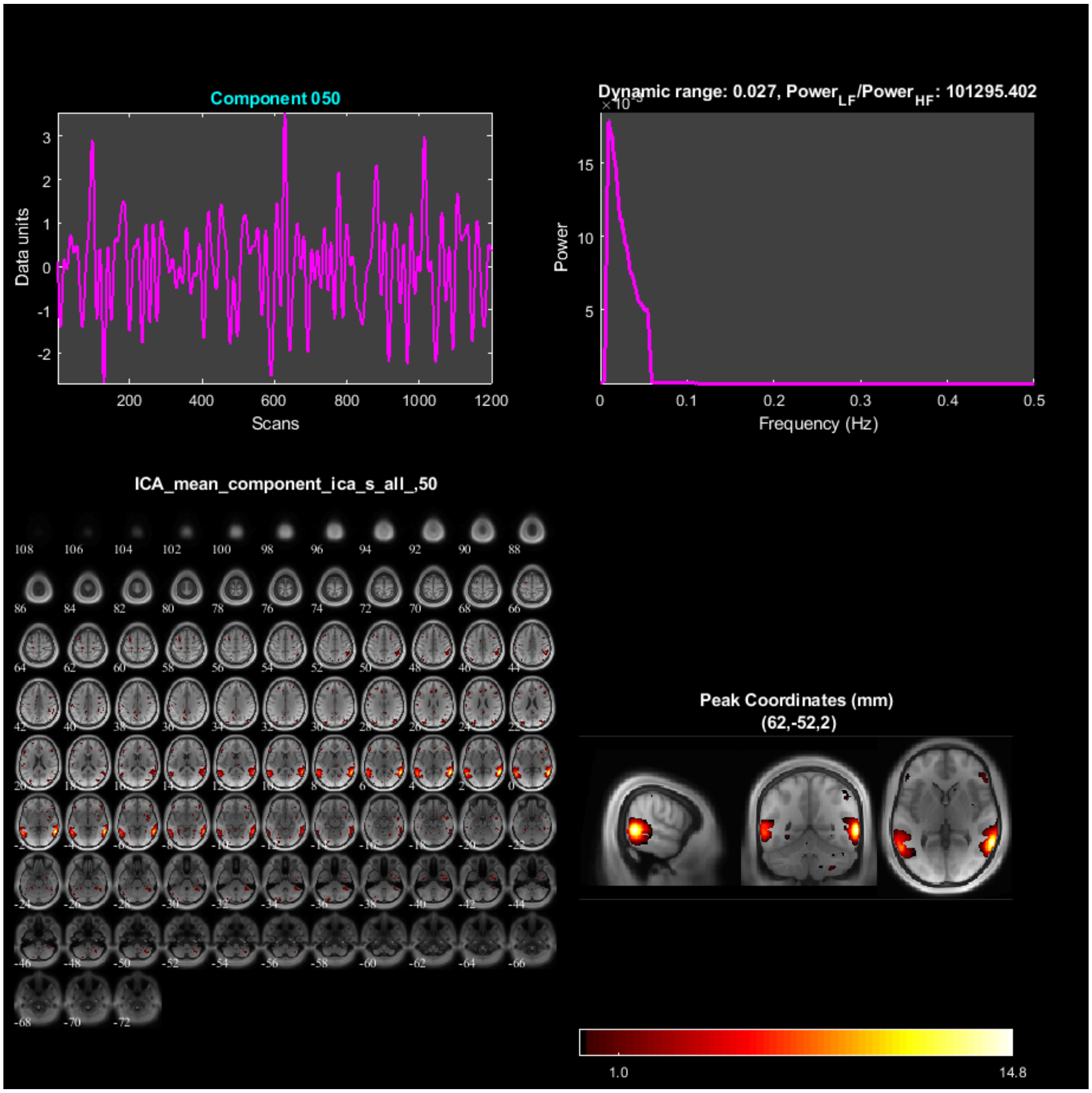

Component 050
Data units
Scans
Dynamic range: 0.027, Power$_{LF}$/Power$_{HF}$: 101295.402
Power
Frequency (Hz)
ICA_mean_component_ica_s_all_,50
Peak Coordinates (mm)
(62,-52,2)
1.0
14.8

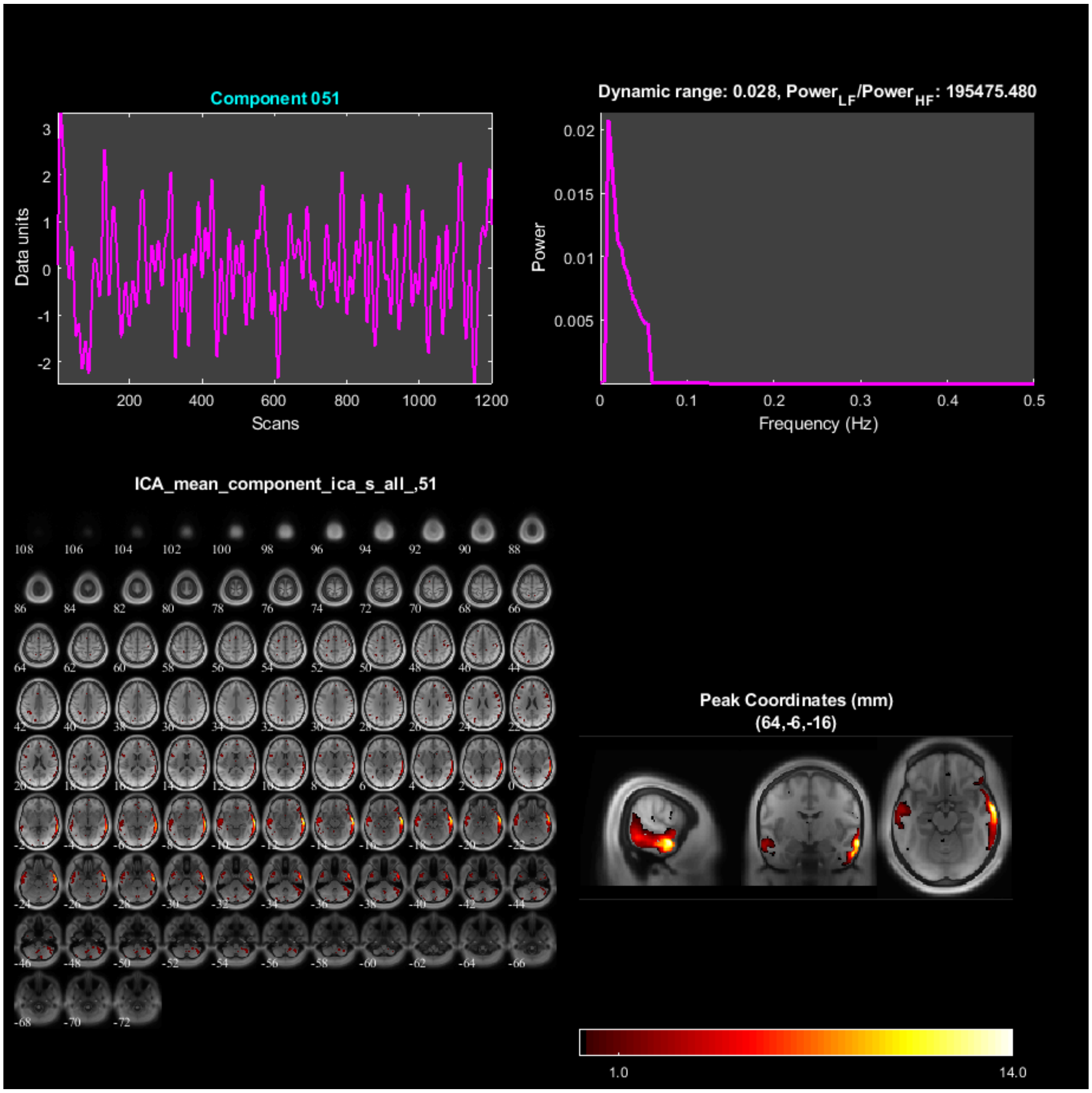

Component 051
Data units
Scans
Dynamic range: 0.028, Power$_{LF}$/Power$_{HF}$: 195475.480
Power
Frequency (Hz)
ICA_mean_component_ica_s_all_,51
Peak Coordinates (mm)
(64,-6,-16)
1.0
14.0

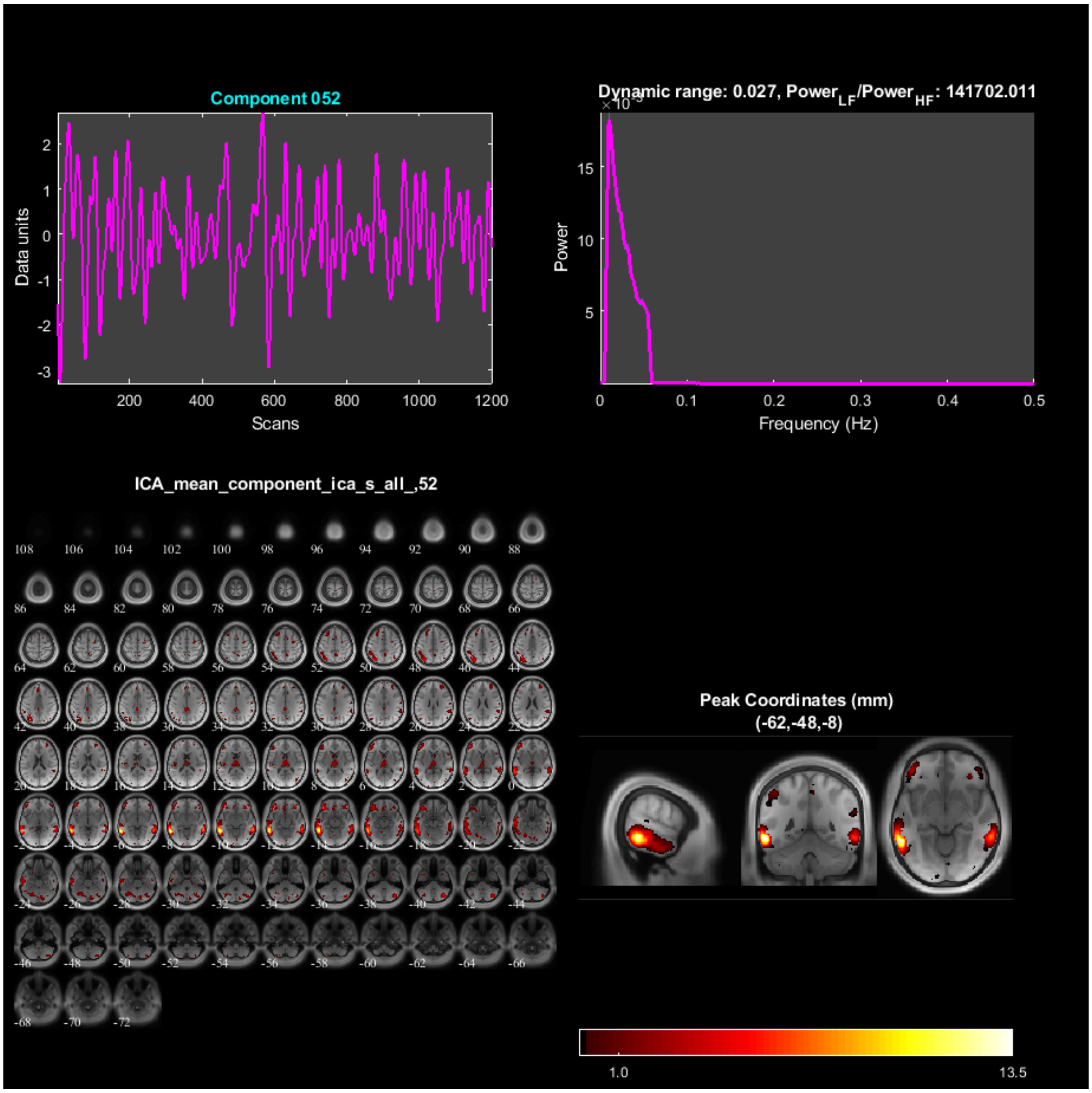
Component 052
Data units
Scans
Dynamic range: 0.027, Power_LF/Power_HF: 141702.011
Power
Frequency (Hz)
ICA_mean_component_ica_s_all_,52
Peak Coordinates (mm)
(-62,-48,-8)
1.0
13.5

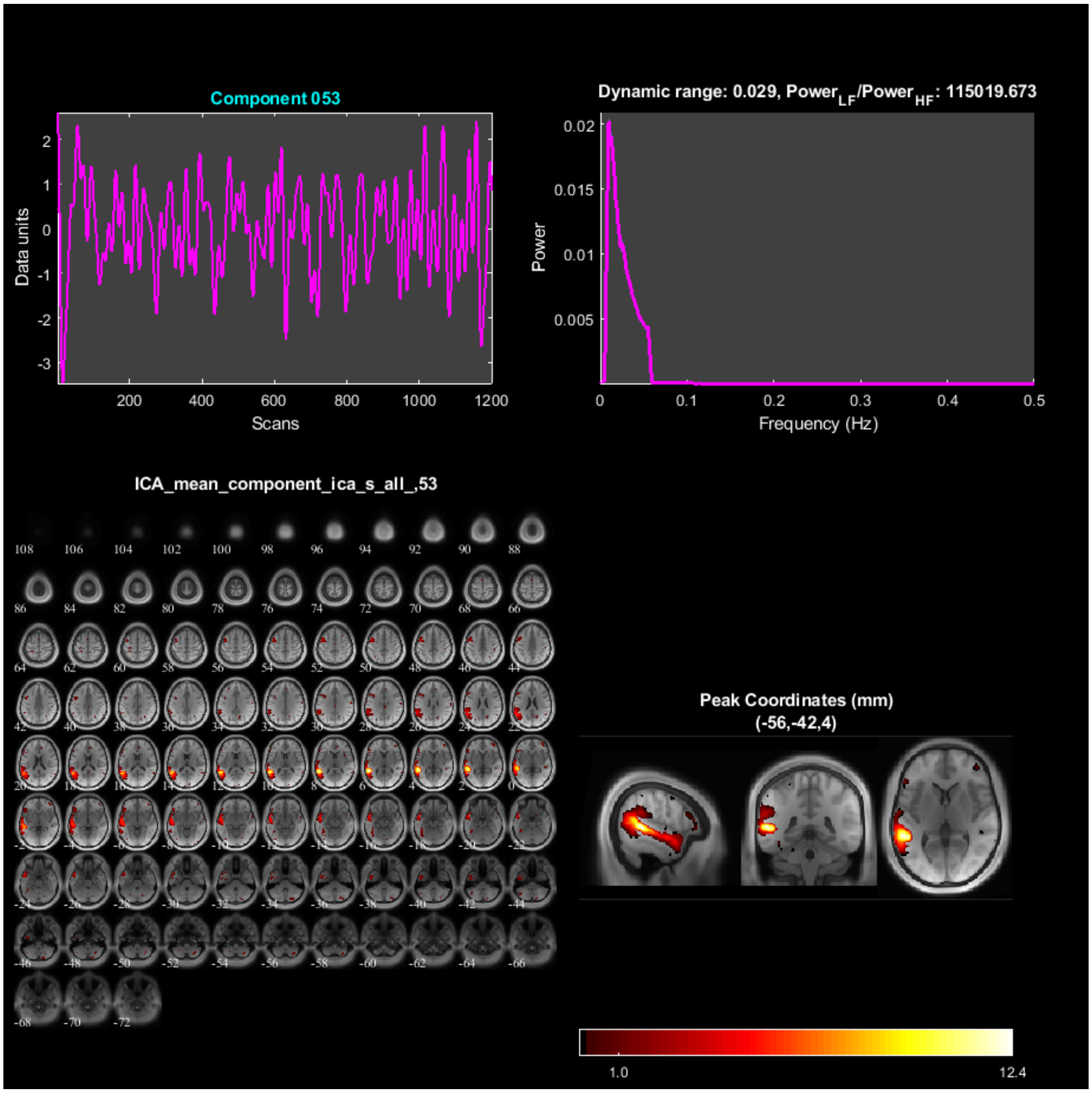

Component 053
Data units
Scans
Dynamic range: 0.029, Power$_{LF}$/Power$_{HF}$: 115019.673
Power
Frequency (Hz)
ICA_mean_component_ica_s_all_,53
Peak Coordinates (mm)
(-56,-42,4)
1.0
12.4

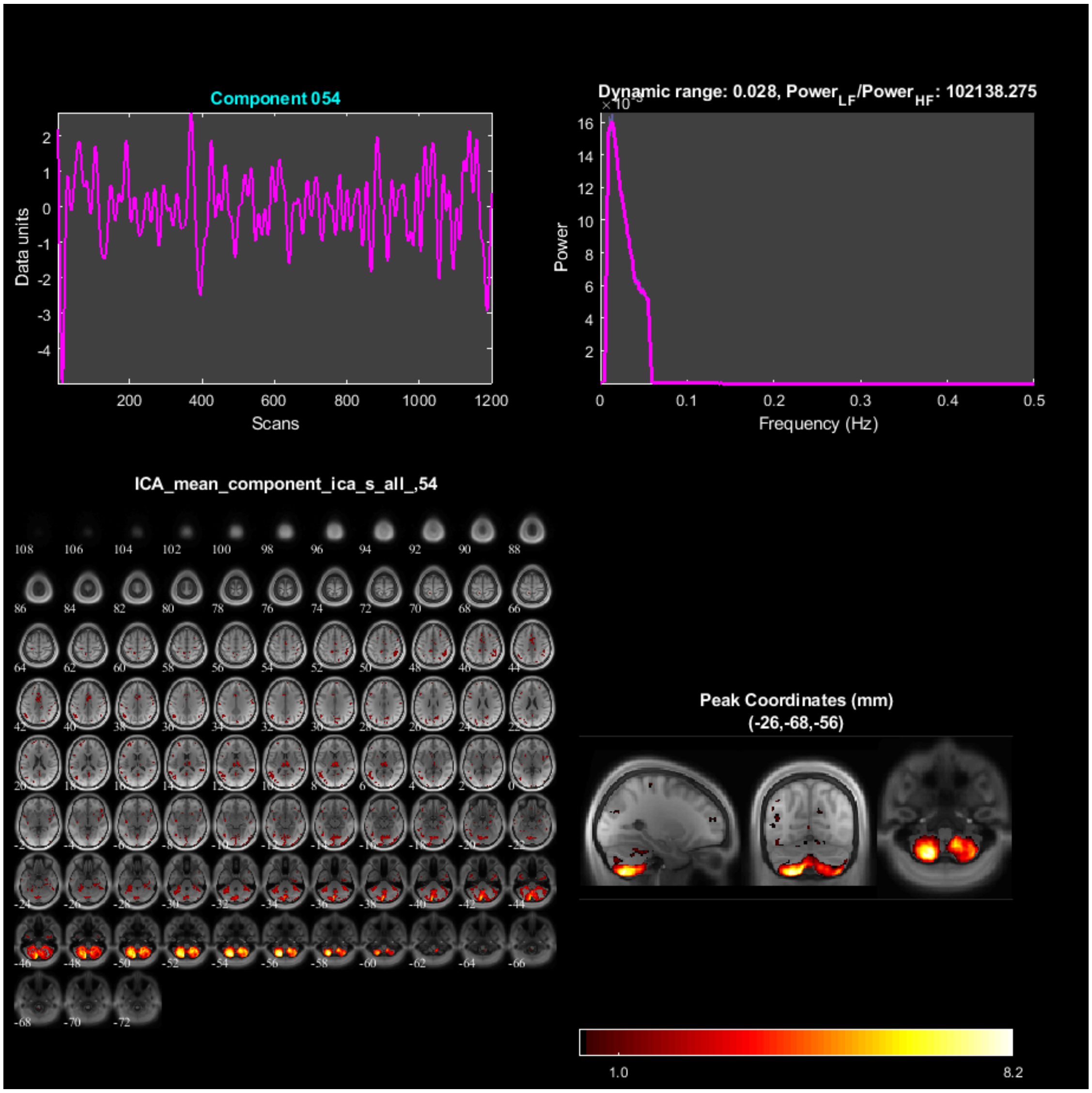
Component 054
Data units
Scans
Dynamic range: 0.028, Power$_{LF}$/Power$_{HF}$: 102138.275
Power
Frequency (Hz)
ICA_mean_component_ica_s_all_,54
Peak Coordinates (mm)
(-26,-68,-56)
1.0
8.2

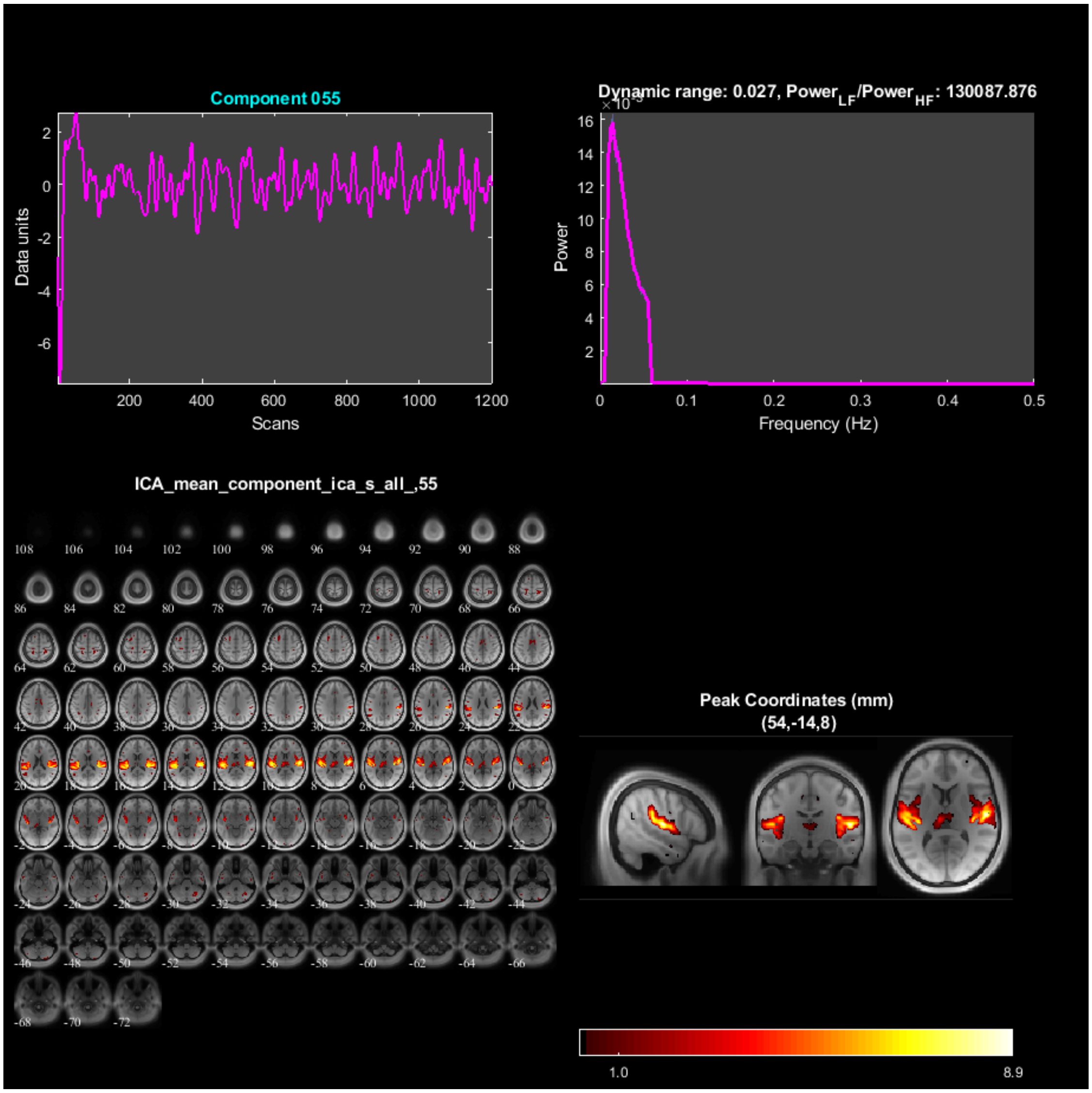

Component 055
Data units
Scans
Dynamic range: 0.027, Power_LF/Power_HF: 130087.876
Power
Frequency (Hz)
ICA_mean_component_ica_s_all_,55
Peak Coordinates (mm)
(54,-14,8)
1.0
8.9

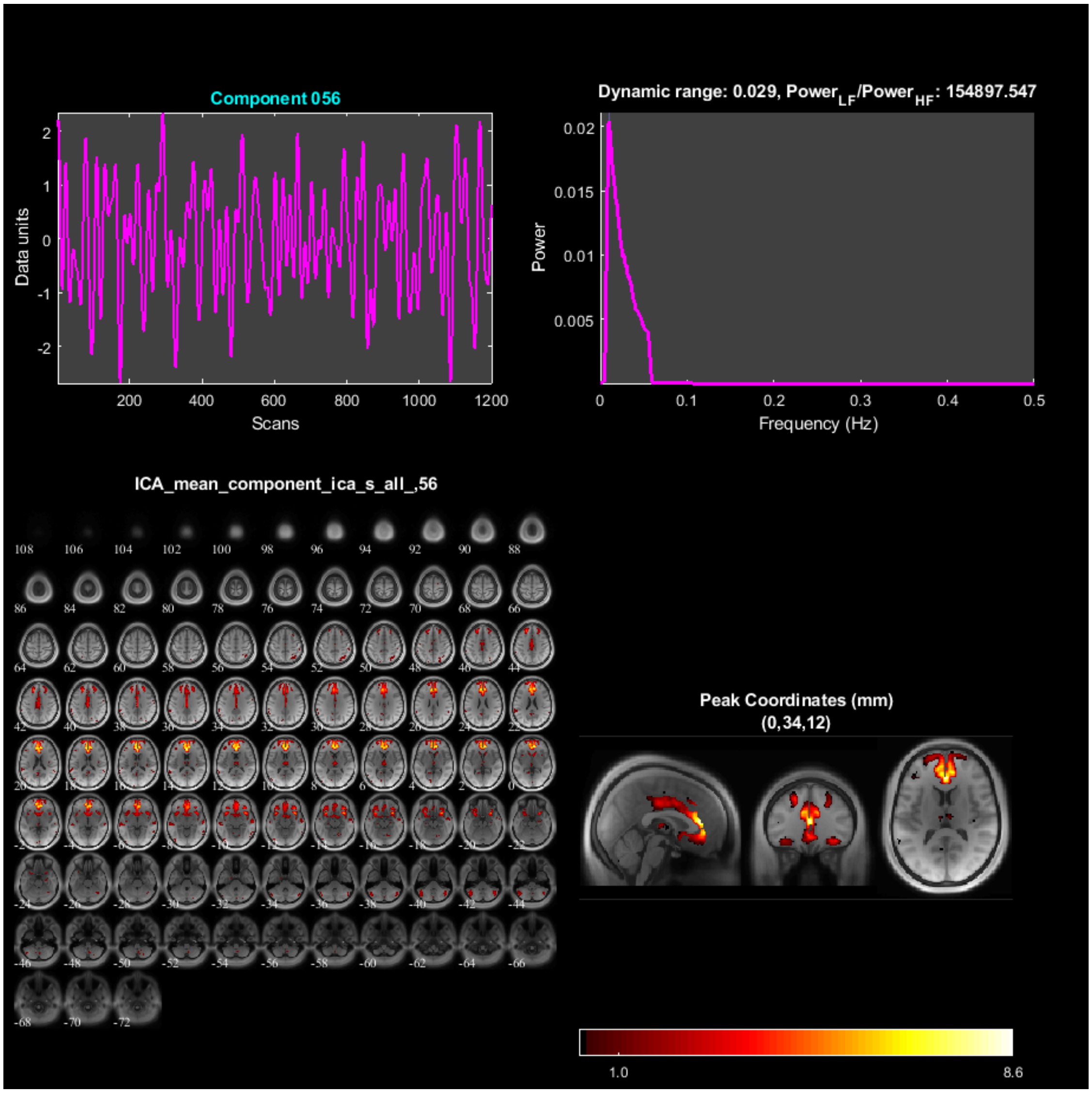

Component 056
Data units
Scans
Dynamic range: 0.029, Power$_{LF}$/Power$_{HF}$: 154897.547
Power
Frequency (Hz)
ICA_mean_component_ica_s_all_,56
Peak Coordinates (mm)
(0,34,12)
1.0
8.6

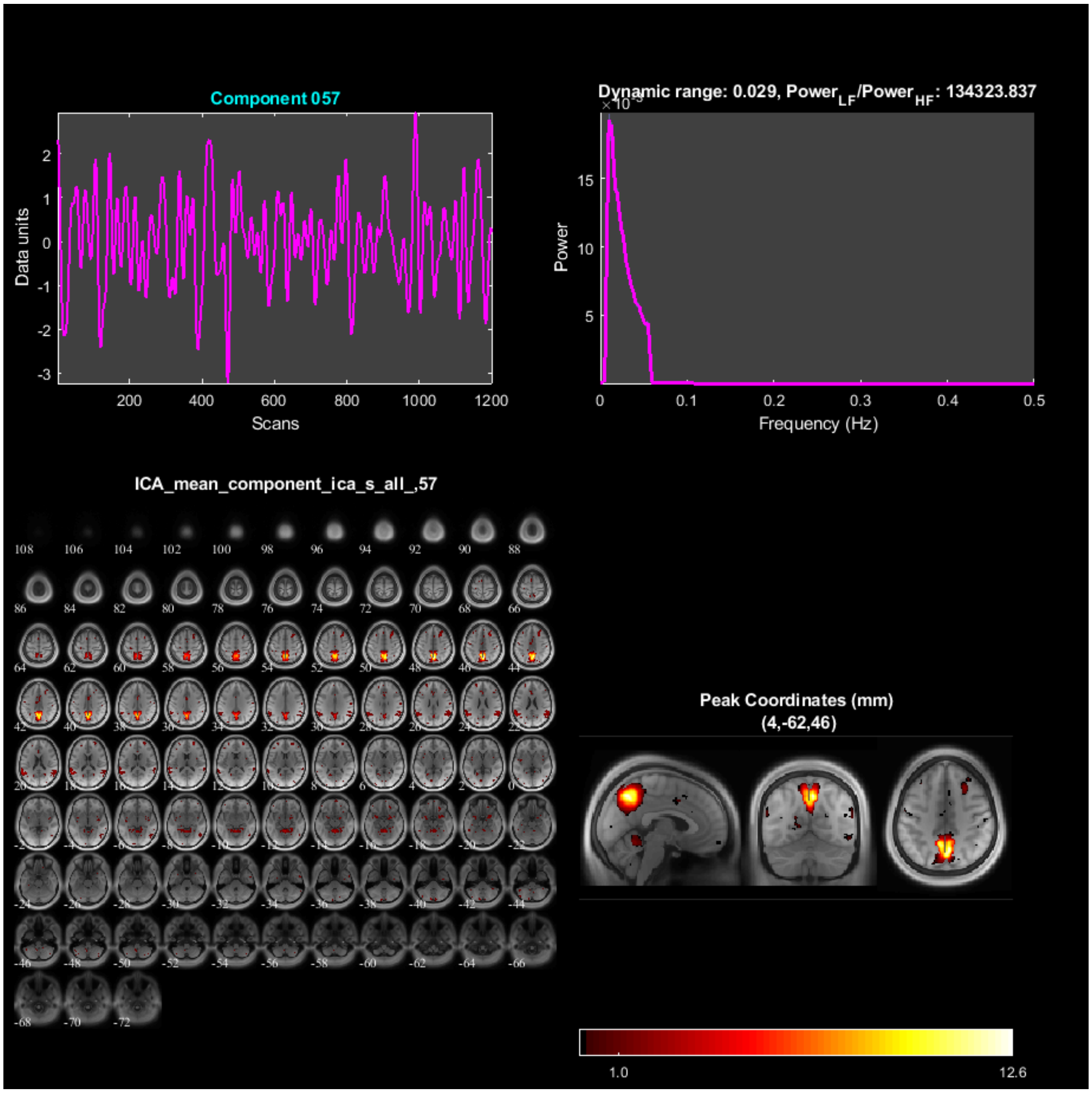
Component 057
Data units
Scans
Dynamic range: 0.029, Power$_{LF}$/Power$_{HF}$: 134323.837
Power
Frequency (Hz)
ICA_mean_component_ica_s_all_,57
Peak Coordinates (mm)
(4,-62,46)
1.0
12.6

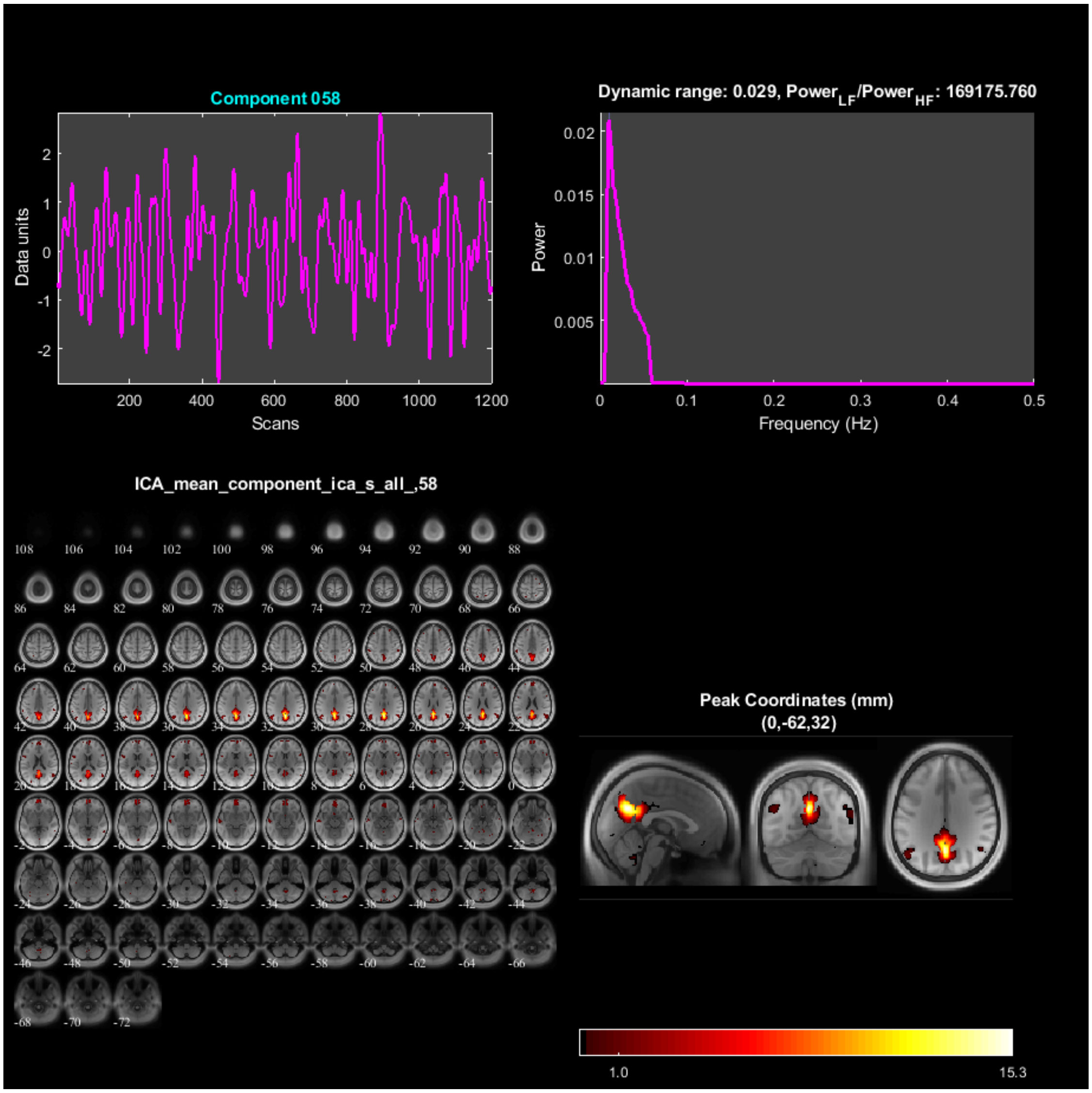
Component 058
Dynamic range: 0.029, Power$_{LF}$/Power$_{HF}$: 169175.760
Data units
Scans
Power
Frequency (Hz)
ICA_mean_component_ica_s_all_,58
Peak Coordinates (mm)
(0,-62,32)
1.0
15.3

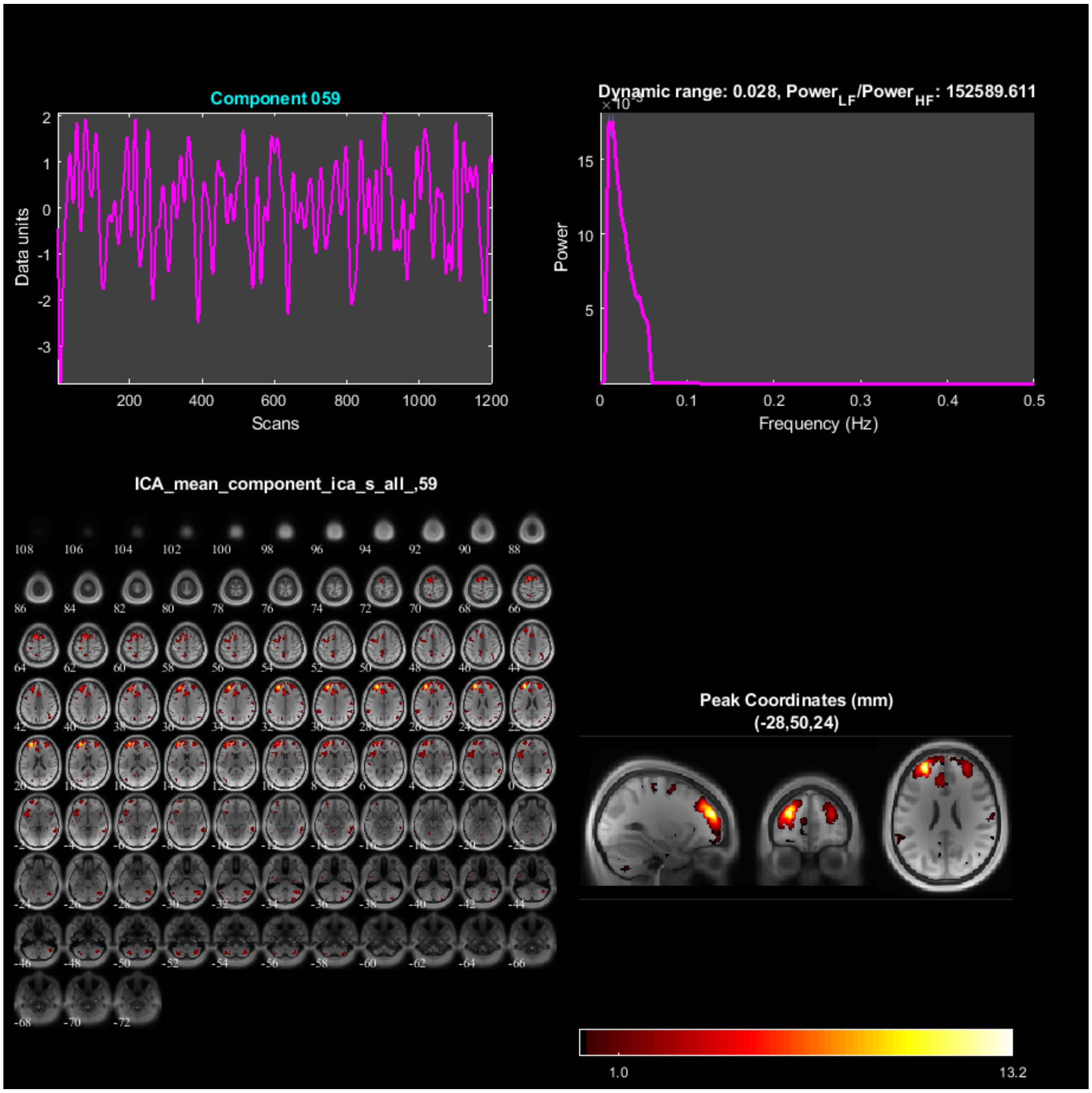
Component 059
Data units
Scans
Dynamic range: 0.028, Power_LF/Power_HF: 152589.611
Power
Frequency (Hz)
ICA_mean_component_ica_s_all_,59
Peak Coordinates (mm)
(-28,50,24)
1.0
13.2

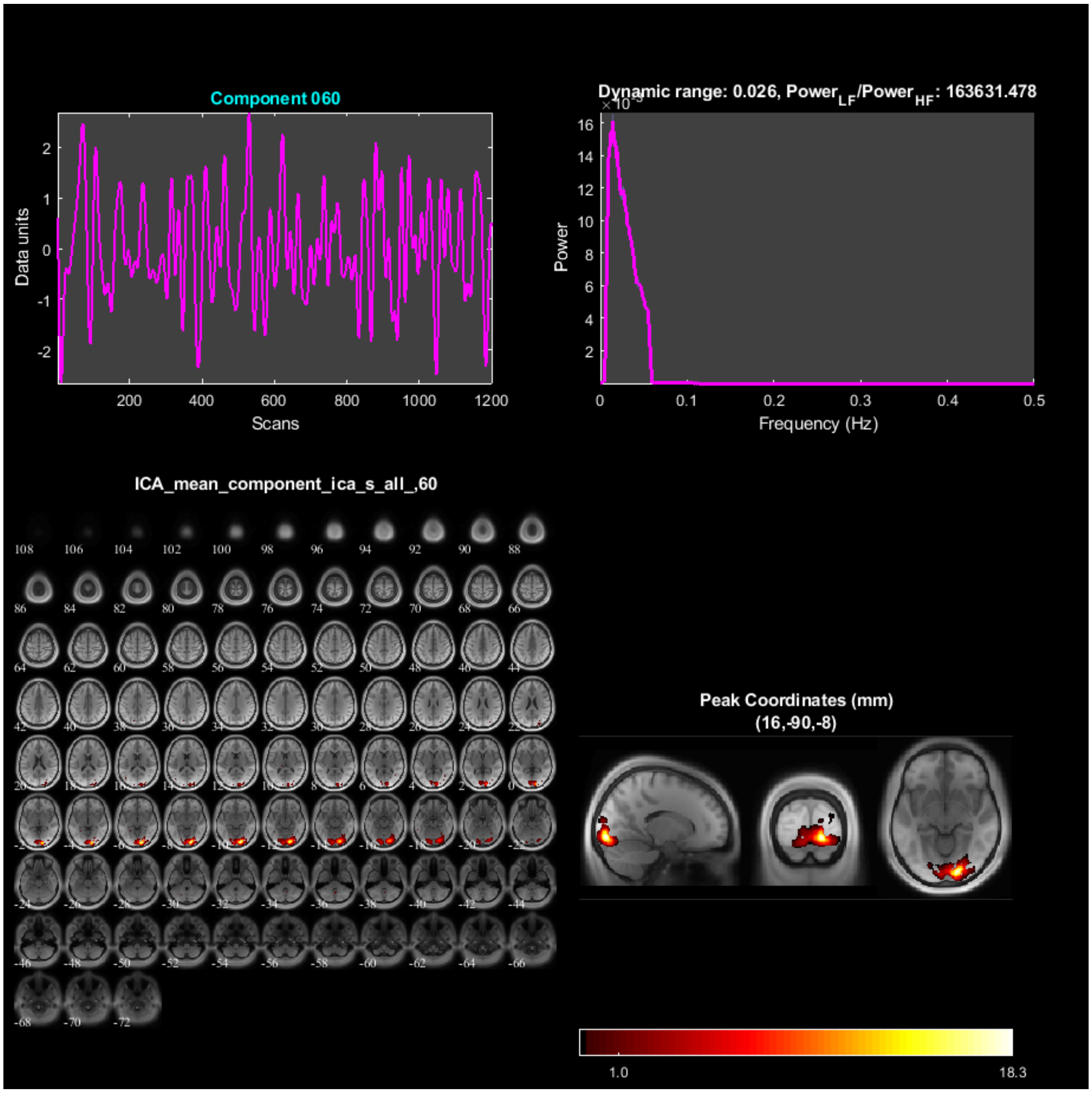

Component 060
Data units
Scans
Dynamic range: 0.026, Power$_{LF}$/Power$_{HF}$: 163631.478
Power
Frequency (Hz)
ICA_mean_component_ica_s_all_,60
Peak Coordinates (mm)
(16,-90,-8)
1.0
18.3

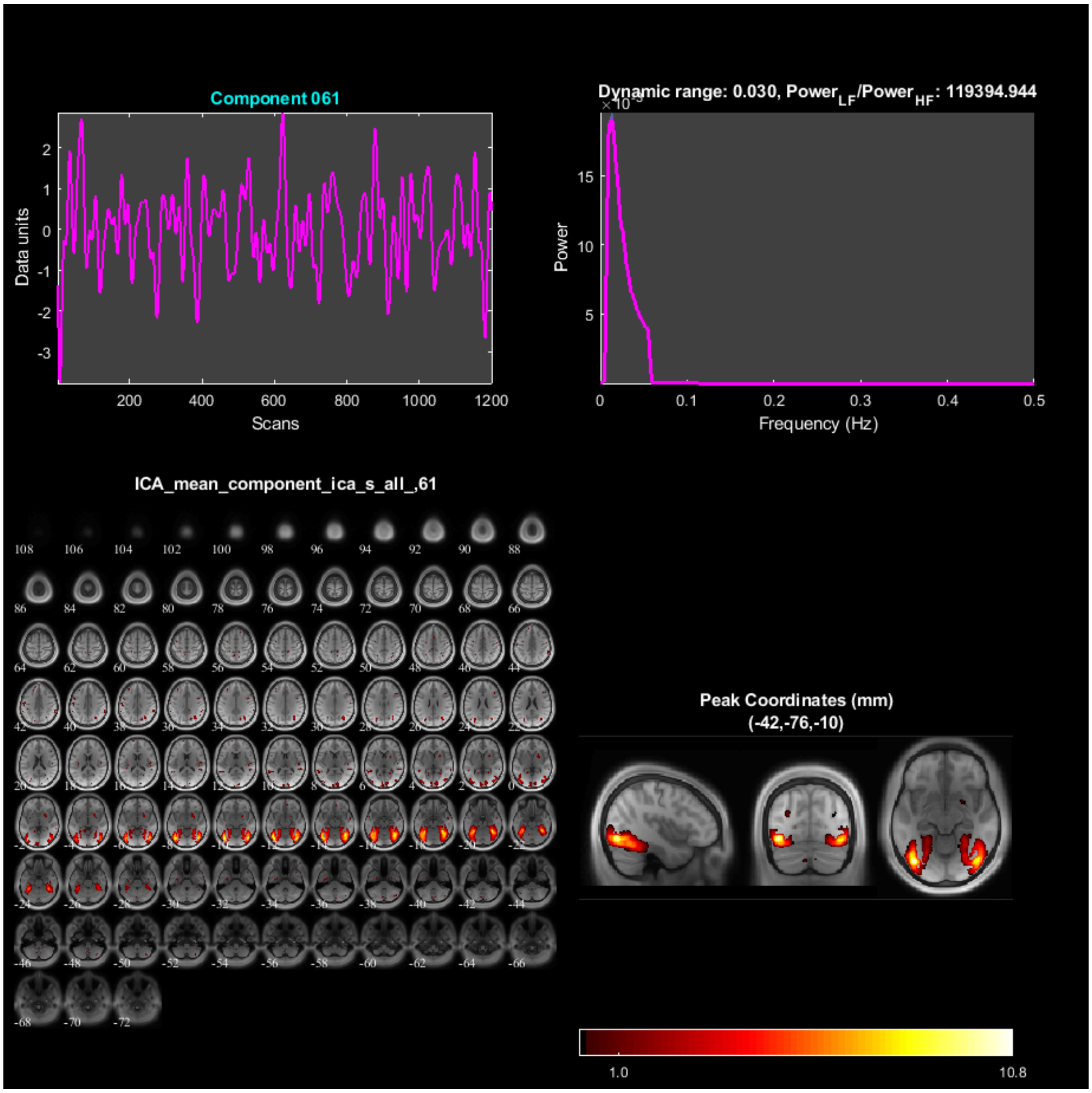
Component 061
Data units
Scans
Dynamic range: 0.030, Power$_{LF}$/Power$_{HF}$: 119394.944
Power
Frequency (Hz)
ICA_mean_component_ica_s_all_,61
Peak Coordinates (mm)
(-42,-76,-10)
1.0
10.8

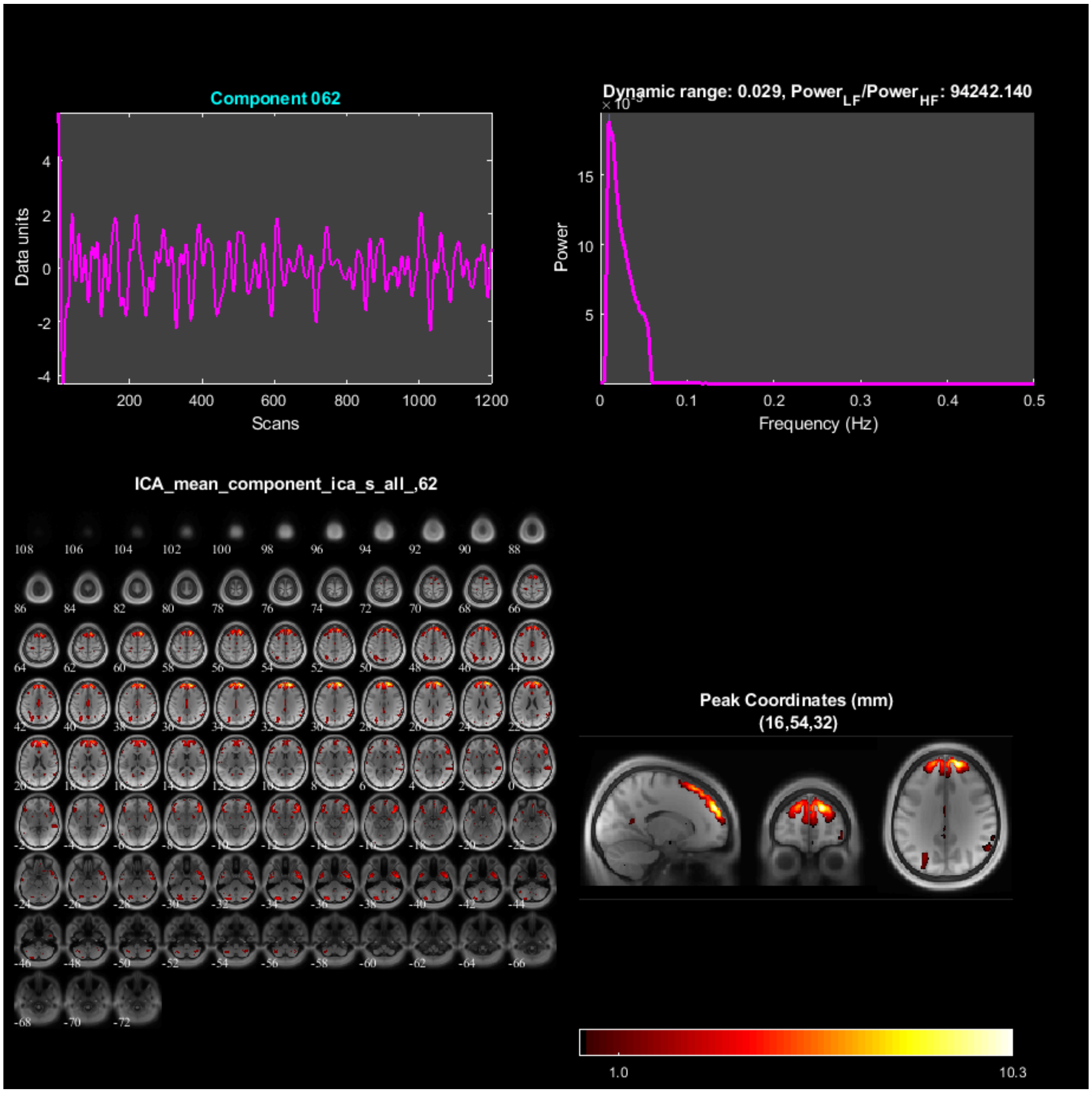
Component 062
Data units
Scans
Dynamic range: 0.029, Power$_{LF}$/Power$_{HF}$: 94242.140
Power
Frequency (Hz)
ICA_mean_component_ica_s_all_,62
Peak Coordinates (mm)
(16,54,32)
1.0
10.3

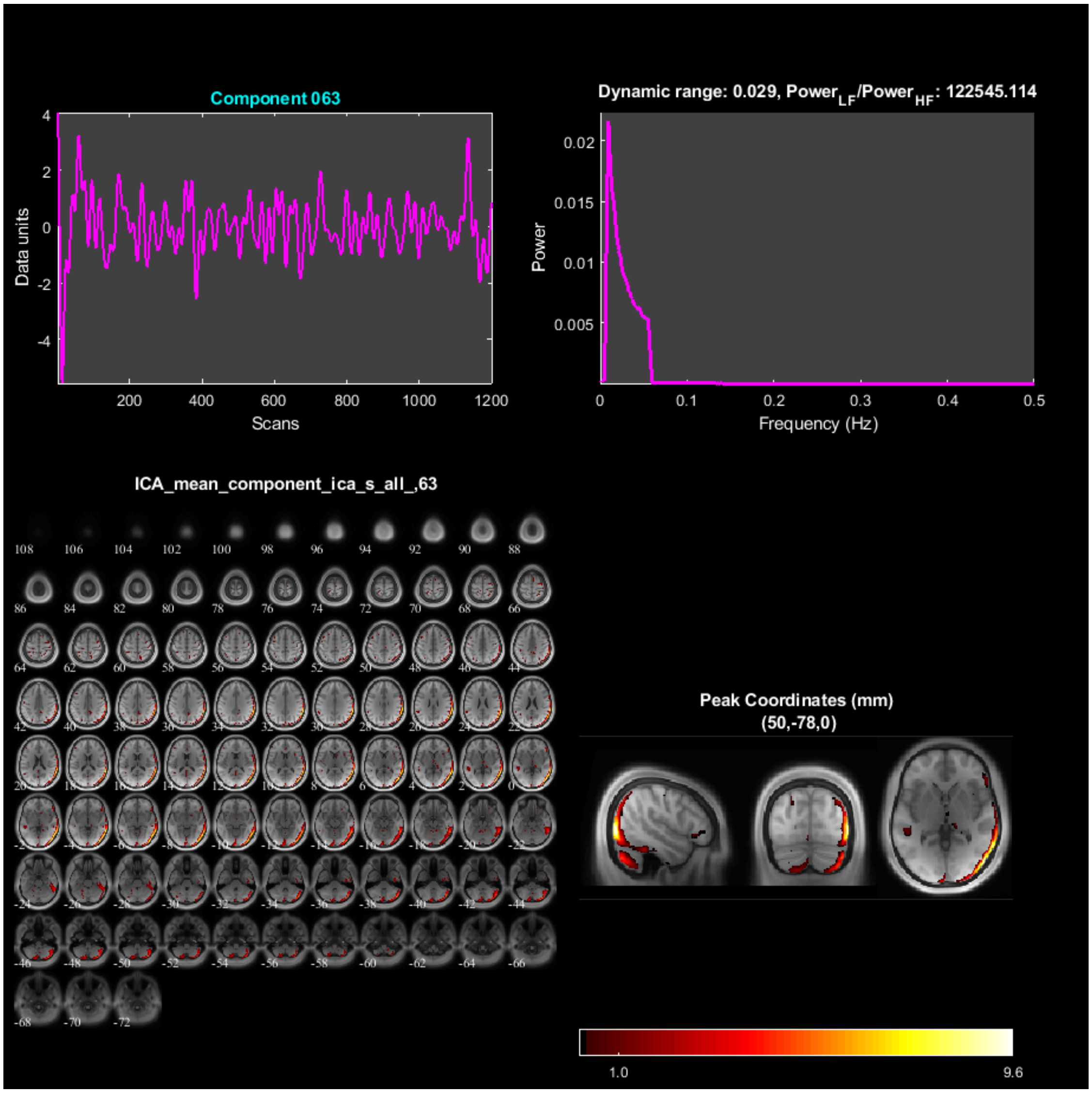
Component 063
Data units
Scans
Dynamic range: 0.029, Power_LF/Power_HF: 122545.114
Power
Frequency (Hz)
ICA_mean_component_ica_s_all_,63
Peak Coordinates (mm)
(50,-78,0)
1.0
9.6

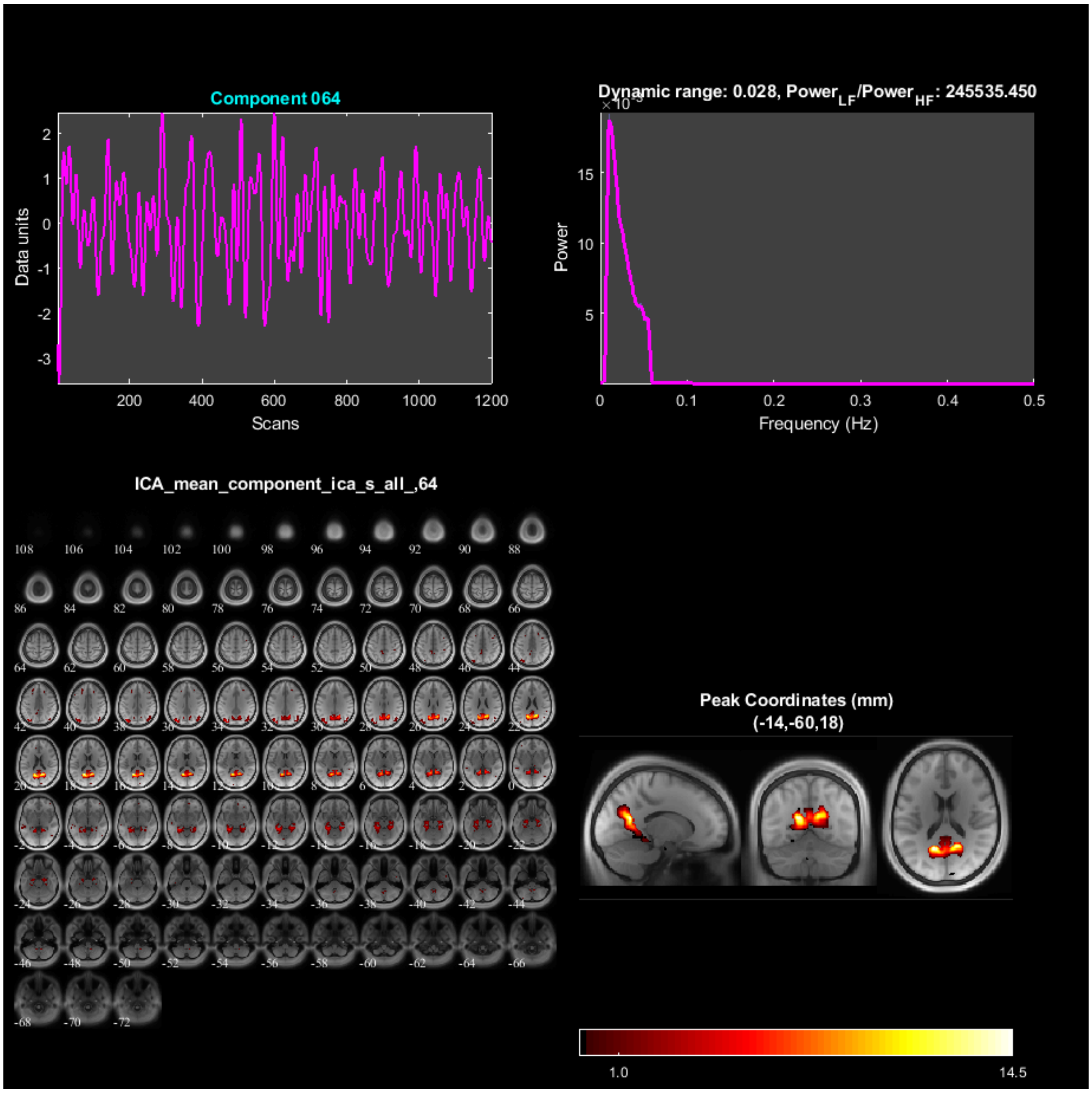
Component 064
Data units
Scans
Dynamic range: 0.028, Power$_{LF}$/Power$_{HF}$: 245535.450
Power
Frequency (Hz)
ICA_mean_component_ica_s_all_,64
Peak Coordinates (mm)
(-14,-60,18)
1.0
14.5

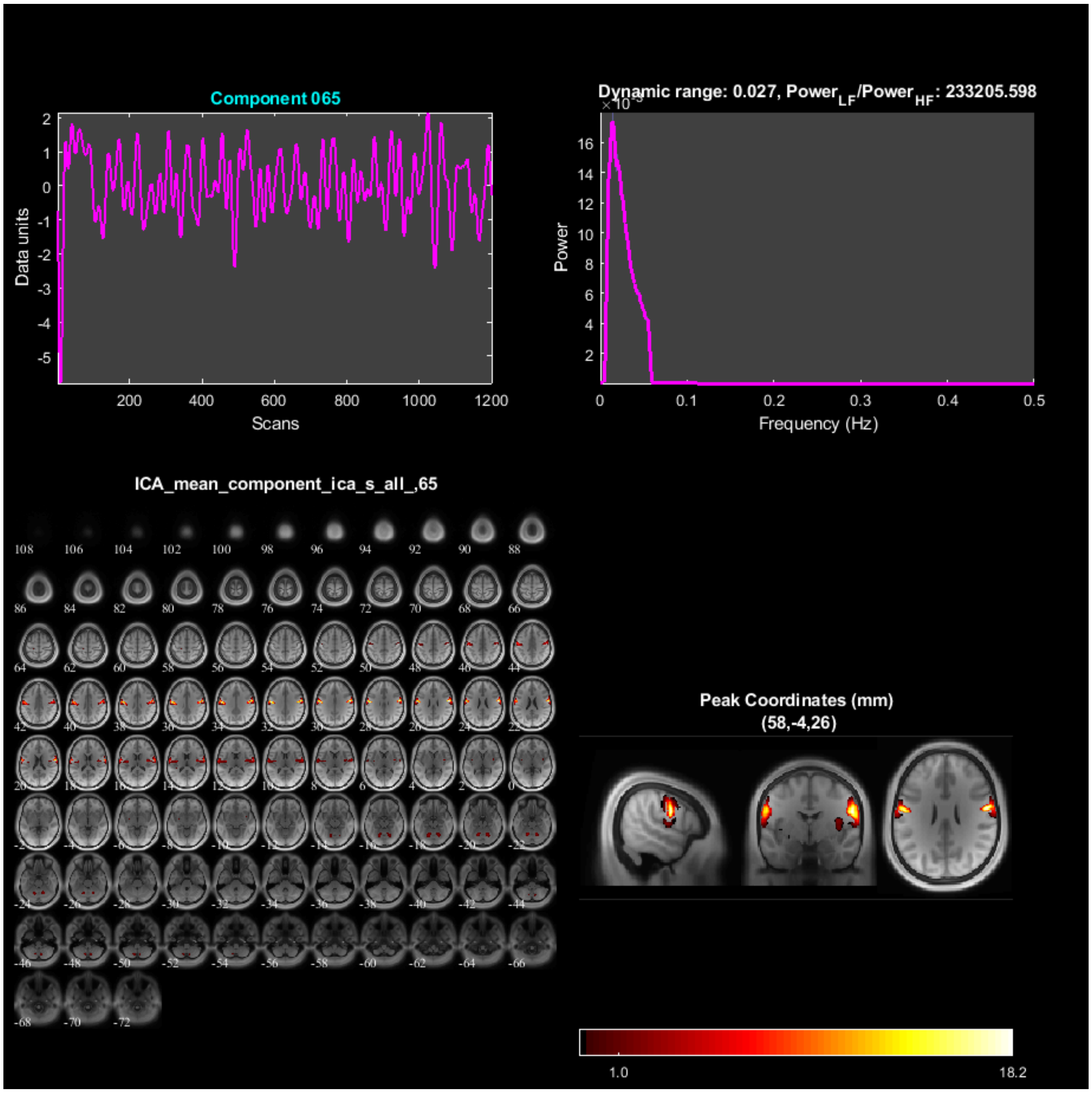
Component 065
Data units
Scans
Dynamic range: 0.027, Power$_{LF}$/Power$_{HF}$: 233205.598
Power
Frequency (Hz)
ICA_mean_component_ica_s_all_,65
Peak Coordinates (mm)
(58,-4,26)
1.0
18.2

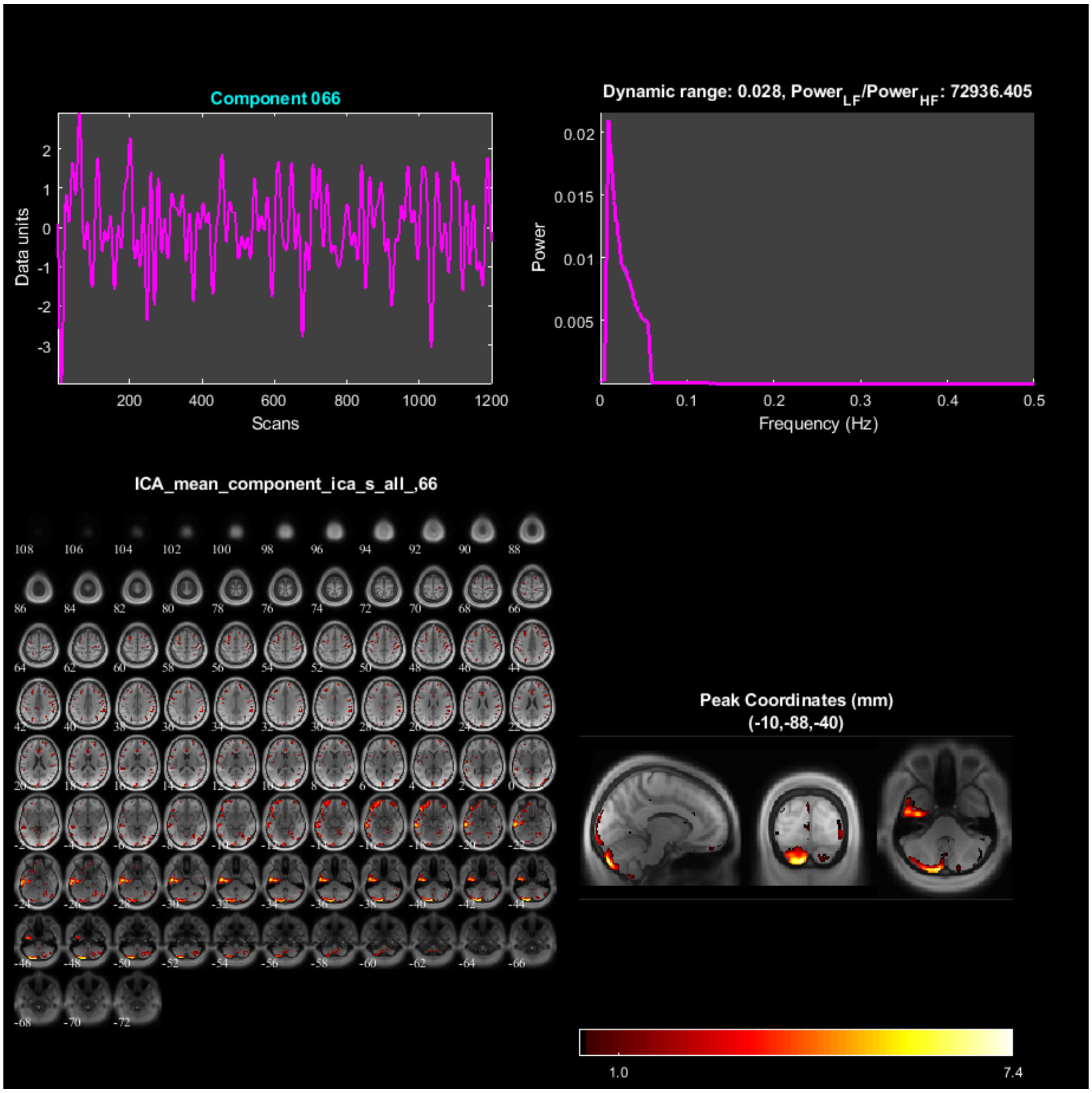

Component 066
Data units
Scans
Dynamic range: 0.028, Power$_{LF}$/Power$_{HF}$: 72936.405
Power
Frequency (Hz)
ICA_mean_component_ica_s_all_,66
Peak Coordinates (mm)
(-10,-88,-40)
1.0
7.4

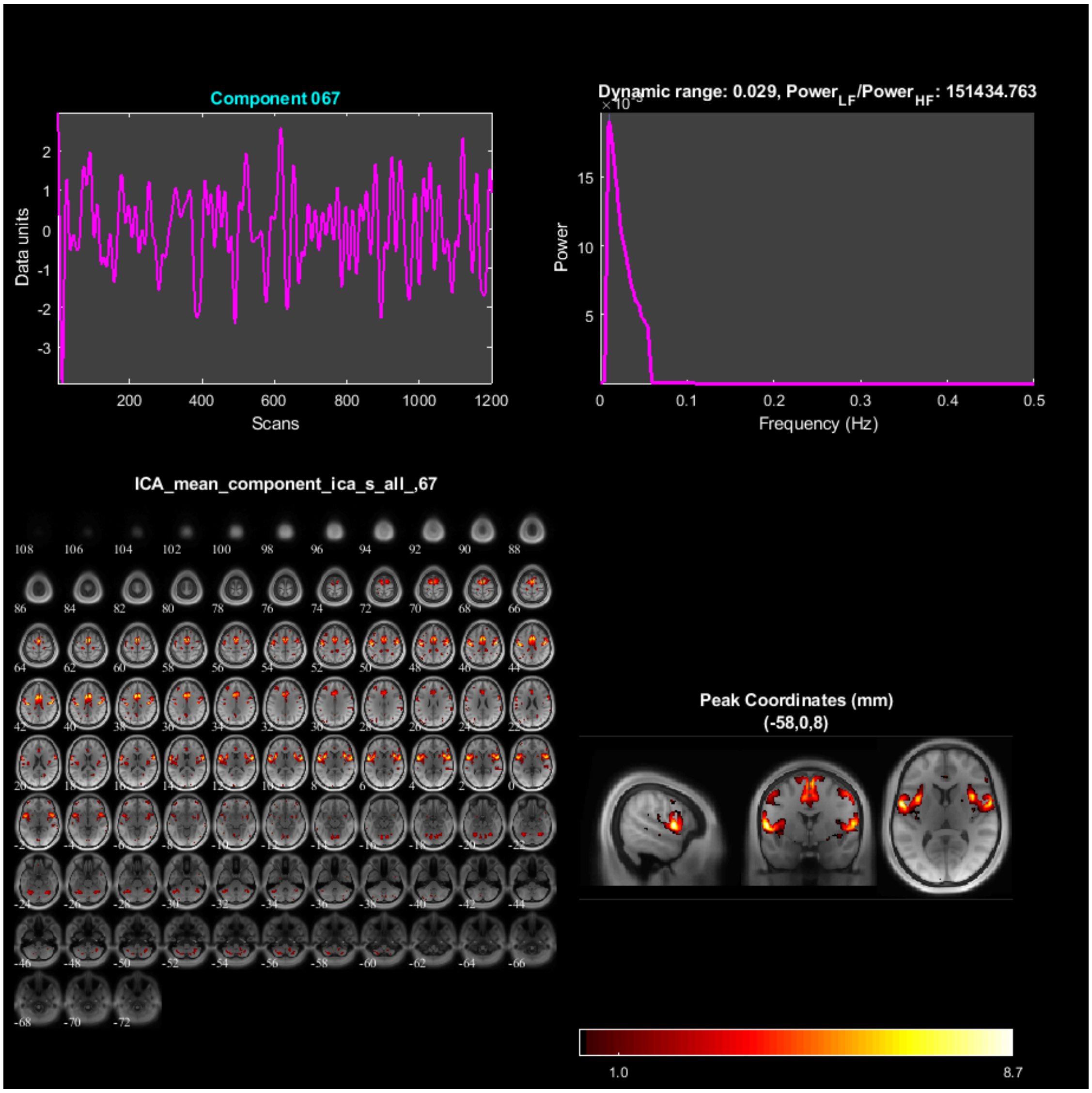
Component 067
Data units
Scans
Dynamic range: 0.029, Power$_{LF}$/Power$_{HF}$: 151434.763
Power
Frequency (Hz)
ICA_mean_component_ica_s_all_,67
Peak Coordinates (mm)
(-58,0,8)
1.0
8.7

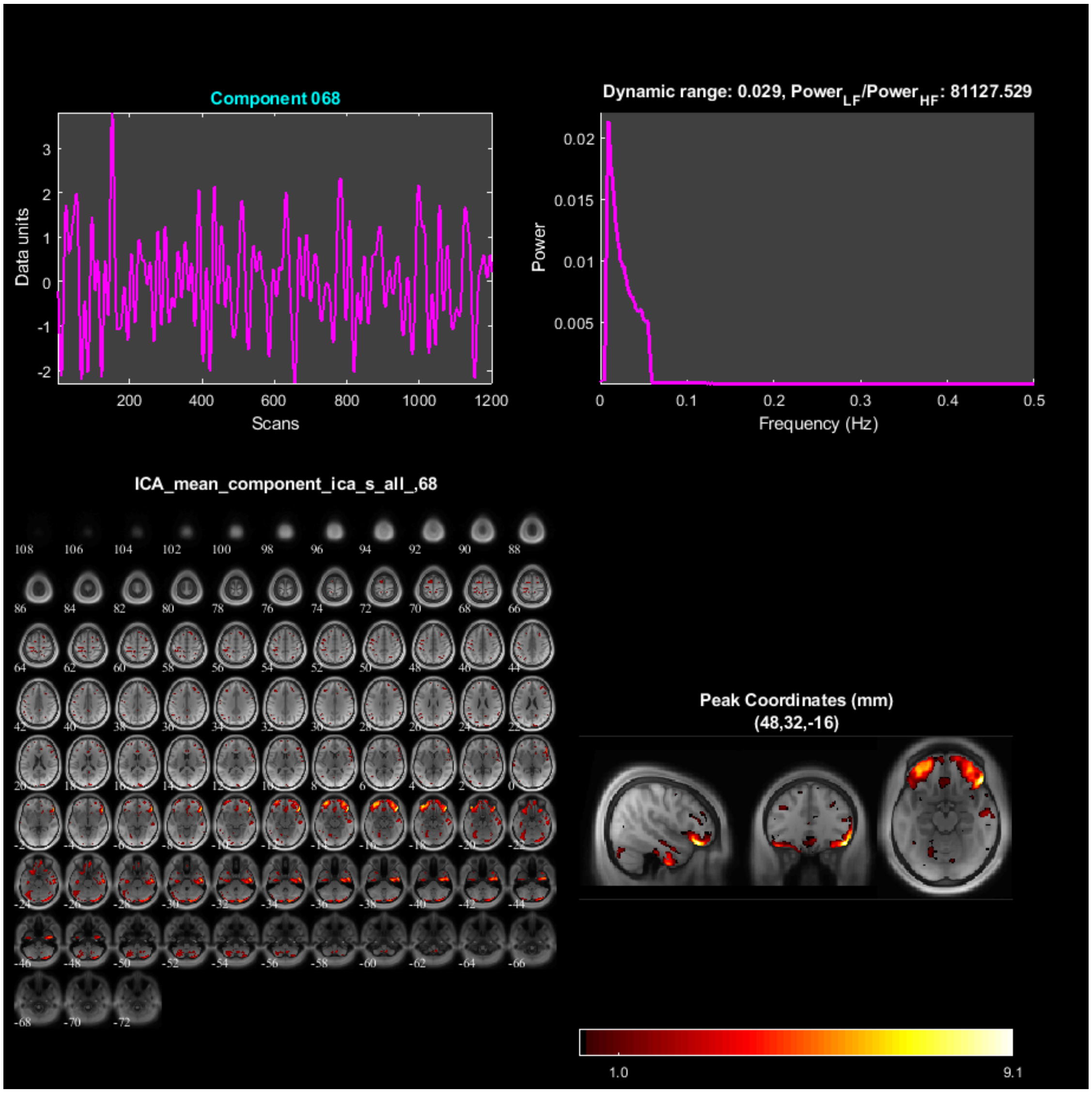
Component 068
Data units
Scans
Dynamic range: 0.029, Power_LF/Power_HF: 81127.529
Power
Frequency (Hz)
ICA_mean_component_ica_s_all_,68
Peak Coordinates (mm)
(48,32,-16)
1.0
9.1

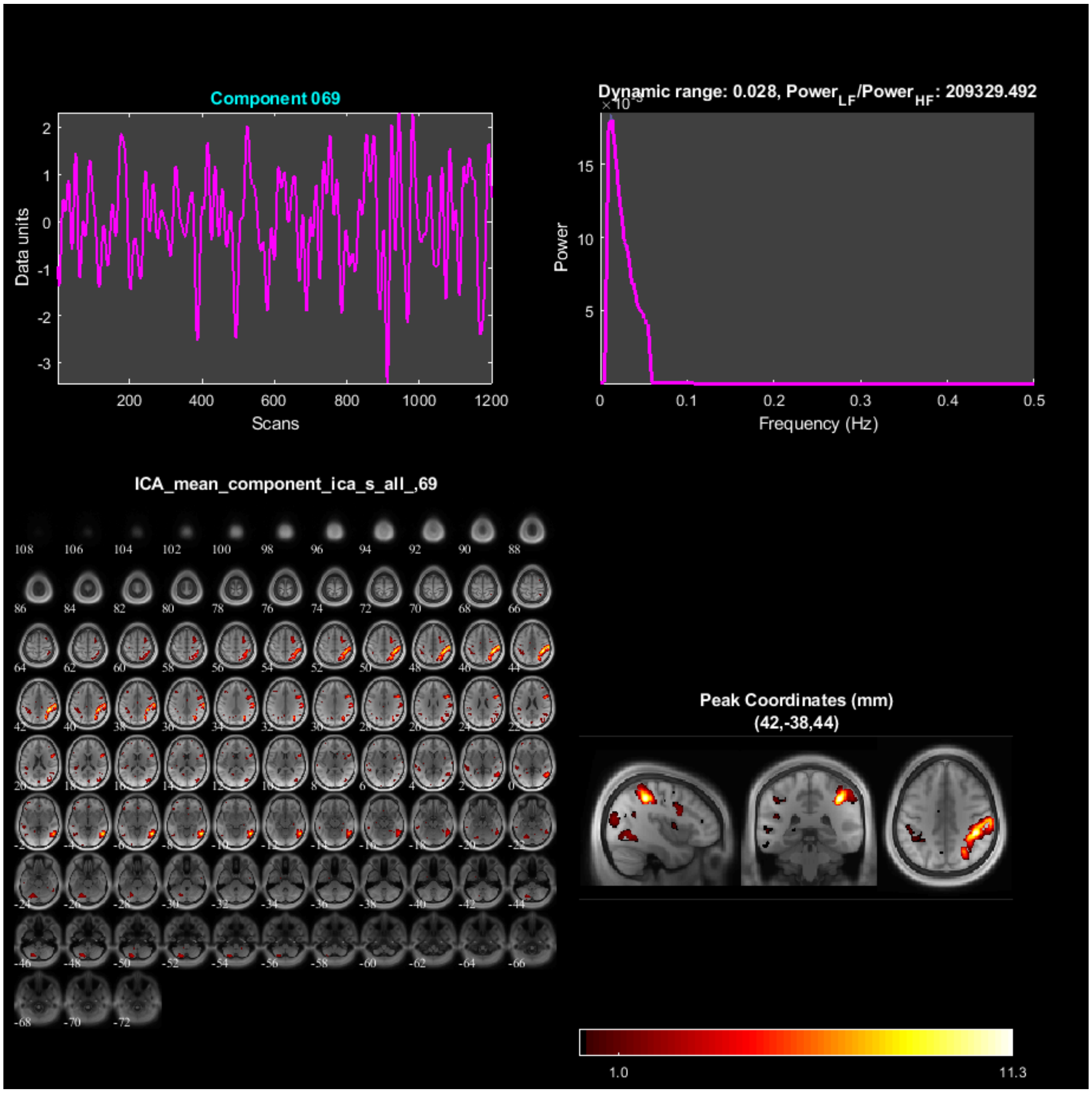
Component 069
Data units
Scans
Dynamic range: 0.028, Power$_{LF}$/Power$_{HF}$: 209329.492
Power
Frequency (Hz)
ICA_mean_component_ica_s_all_,69
Peak Coordinates (mm)
(42,-38,44)
1.0
11.3

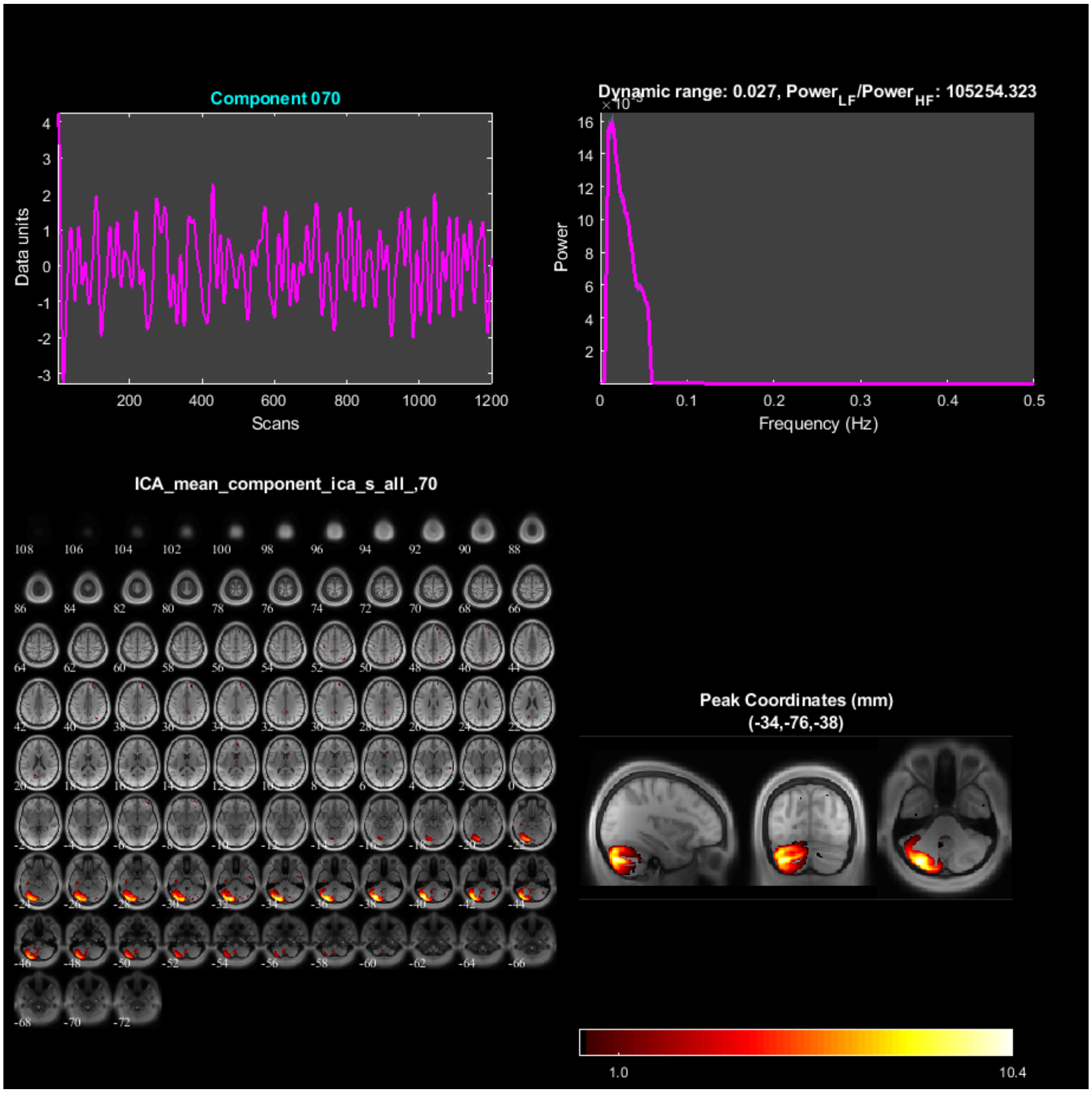
Component 070
Data units
Scans
Dynamic range: 0.027, Power$_{LF}$/Power$_{HF}$: 105254.323
Power
Frequency (Hz)
ICA_mean_component_ica_s_all_,70
Peak Coordinates (mm)
(-34,-76,-38)
1.0
10.4

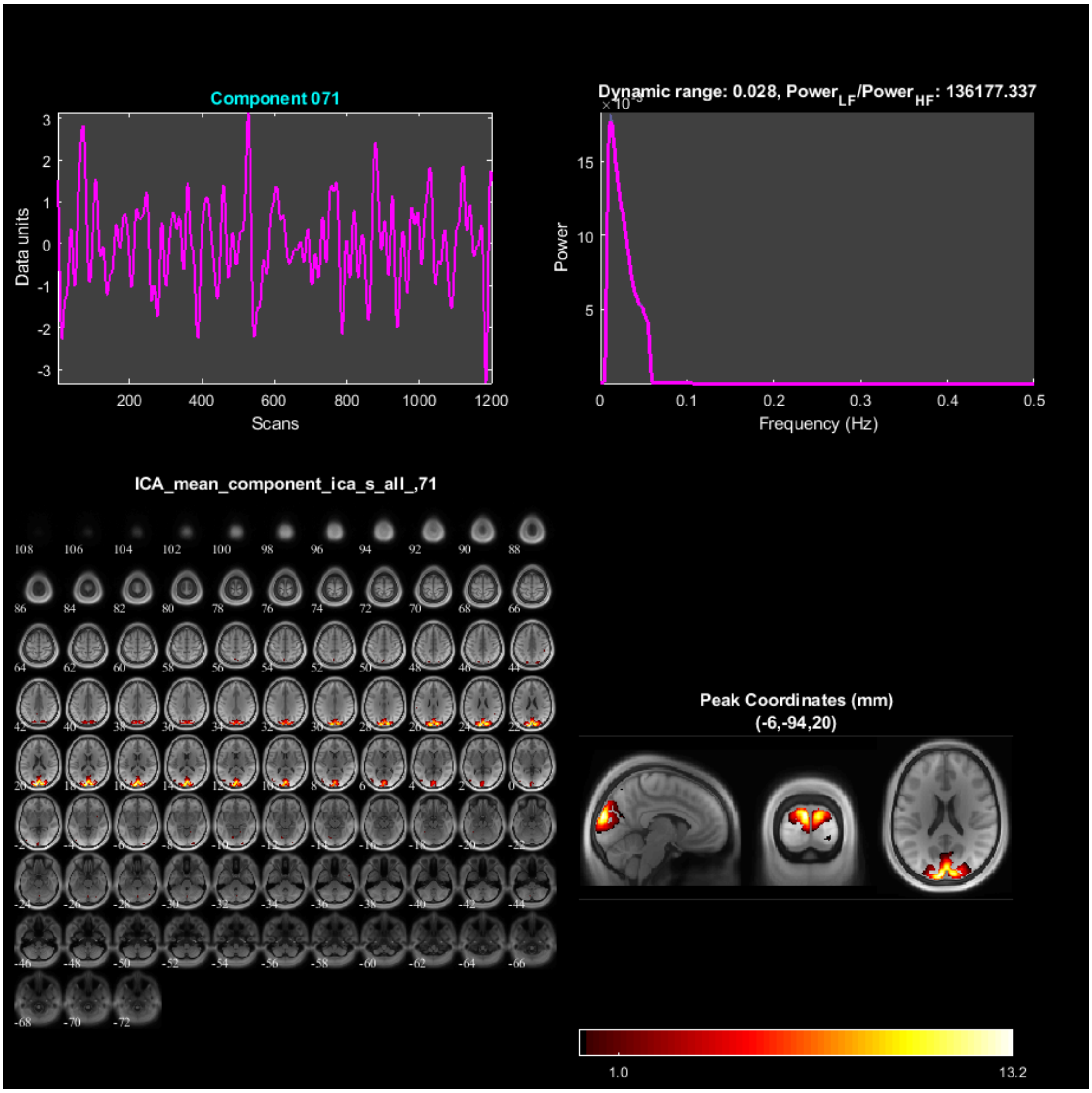
Component 071
Data units
Scans
Dynamic range: 0.028, Power_LF/Power_HF: 136177.337
Power
Frequency (Hz)
ICA_mean_component_ica_s_all_,71
Peak Coordinates (mm)
(-6,-94,20)
1.0
13.2

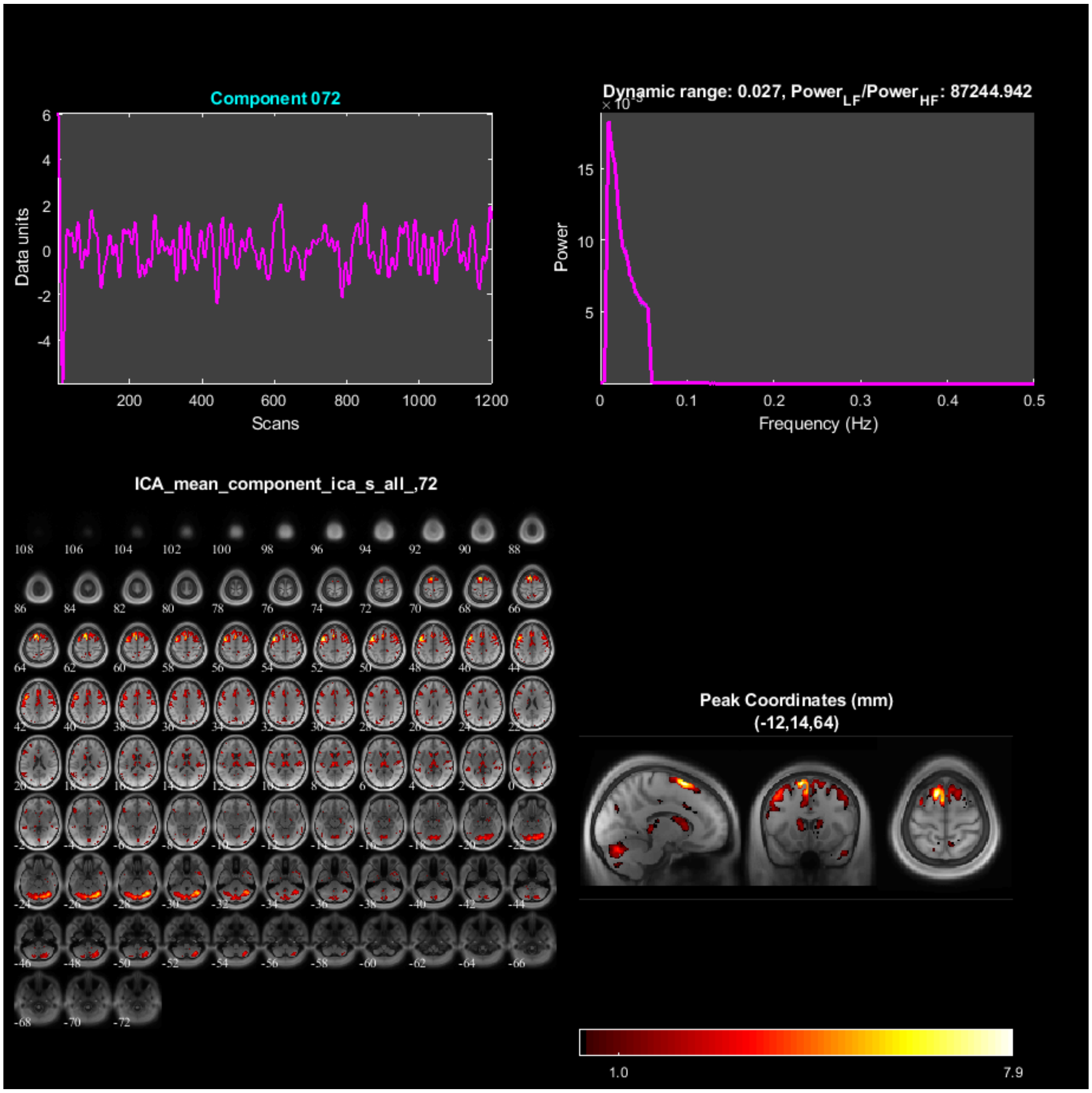
Component 072
Data units
Scans
Dynamic range: 0.027, Power_LF/Power_HF: 87244.942
Power
Frequency (Hz)
ICA_mean_component_ica_s_all_,72
Peak Coordinates (mm)
(-12,14,64)
1.0
7.9

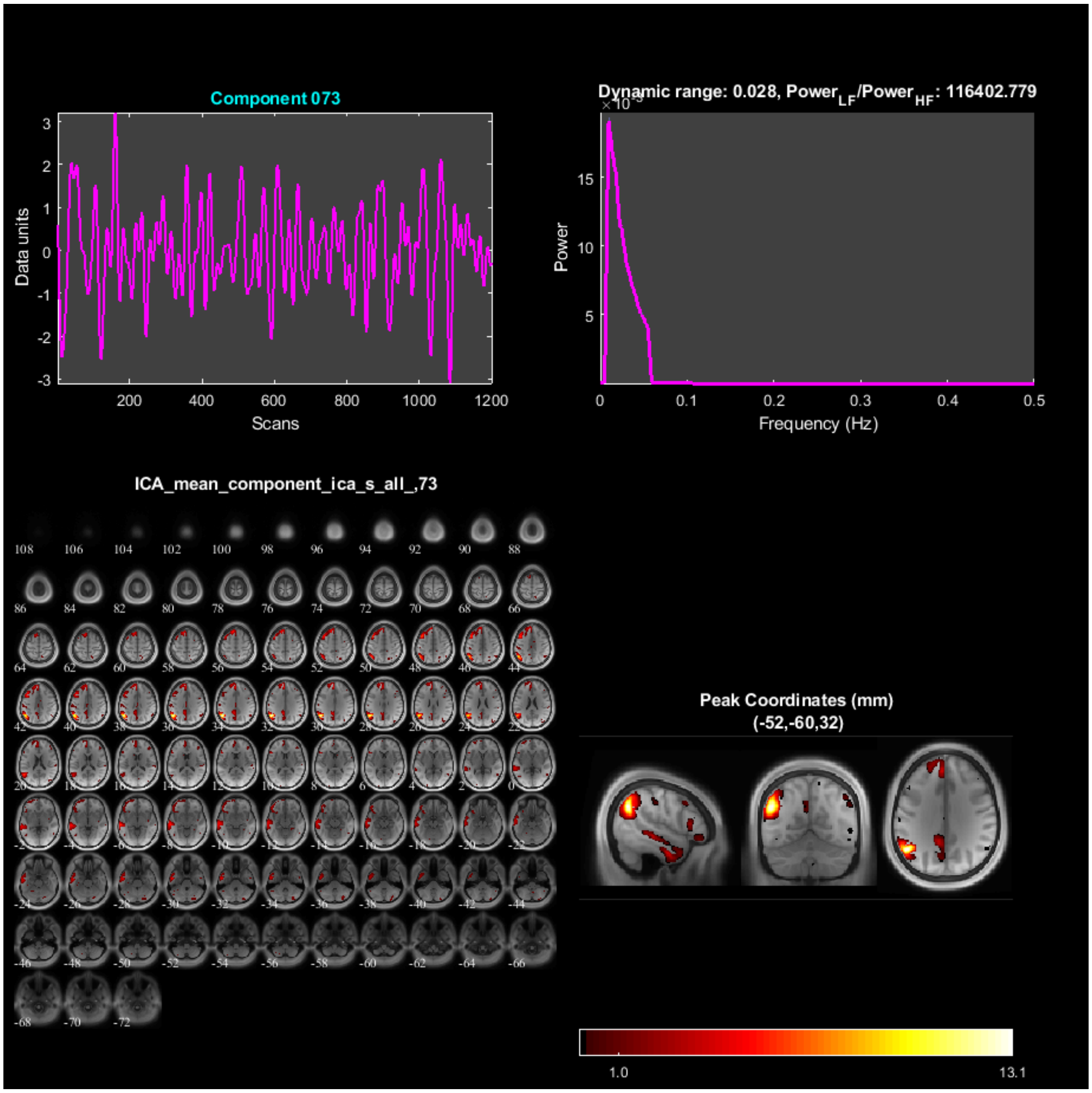
Component 073
Dynamic range: 0.028, Power$_{LF}$/Power$_{HF}$: 116402.779
Data units
Scans
Power
Frequency (Hz)
ICA_mean_component_ica_s_all_,73
Peak Coordinates (mm)
(-52,-60,32)
1.0
13.1

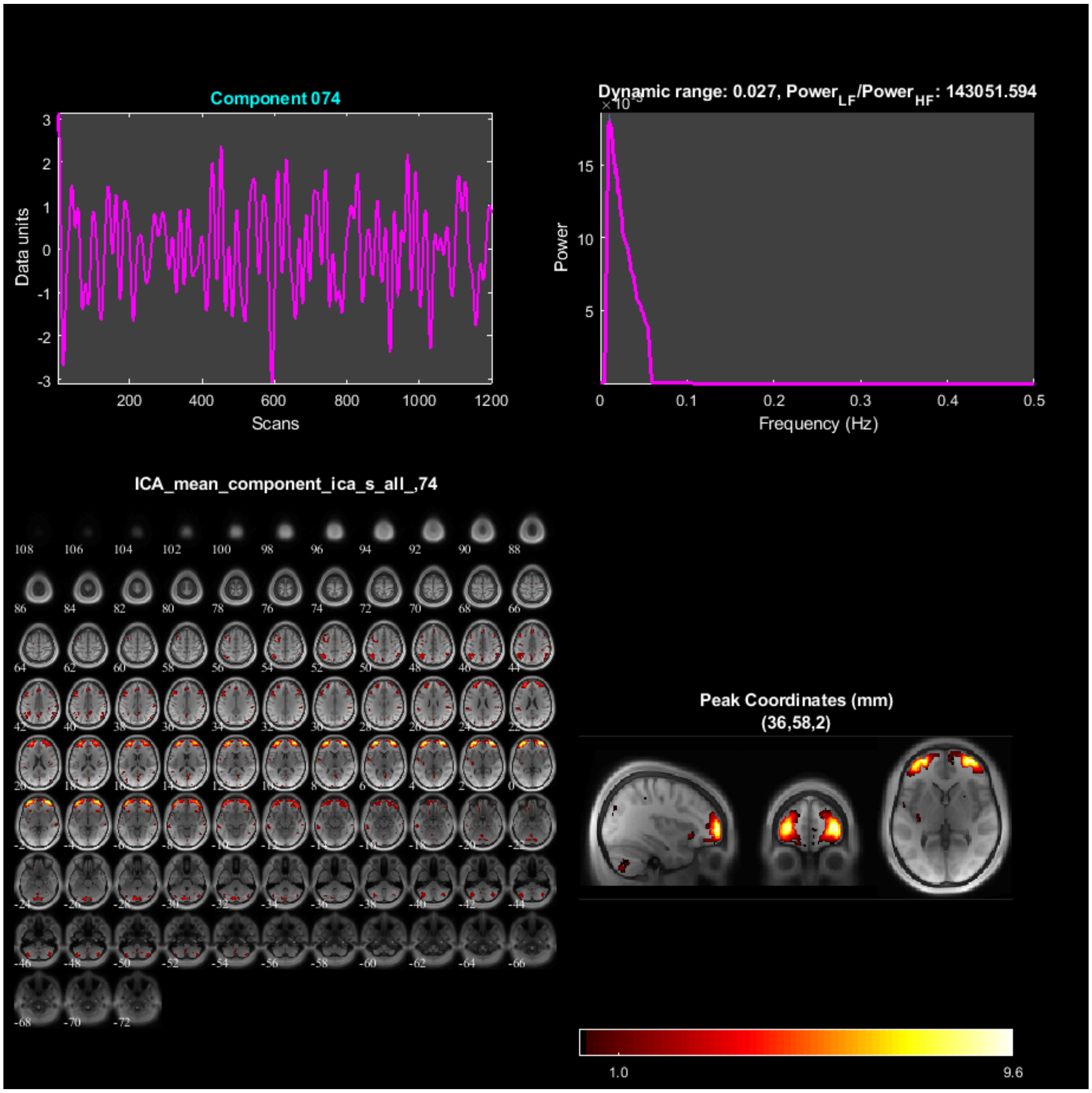

Component 074
Data units
Scans
Dynamic range: 0.027, Power$_{LF}$/Power$_{HF}$: 143051.594
Power
Frequency (Hz)
ICA_mean_component_ica_s_all_,74
Peak Coordinates (mm)
(36,58,2)
1.0
9.6

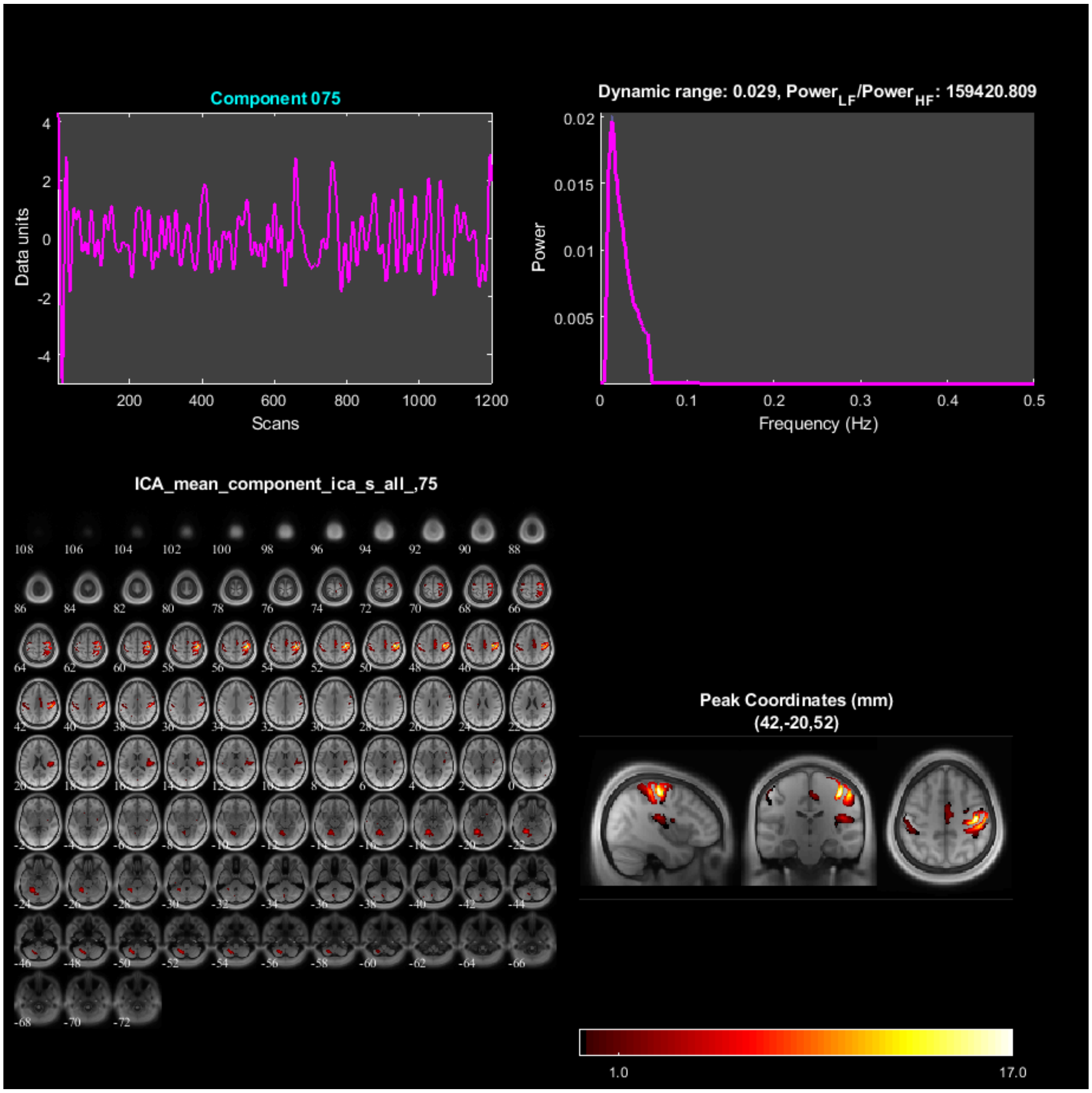

Component 075
Data units
Scans
Dynamic range: 0.029, Power_LF/Power_HF: 159420.809
Power
Frequency (Hz)
ICA_mean_component_ica_s_all_,75
Peak Coordinates (mm)
(42,-20,52)
1.0
17.0

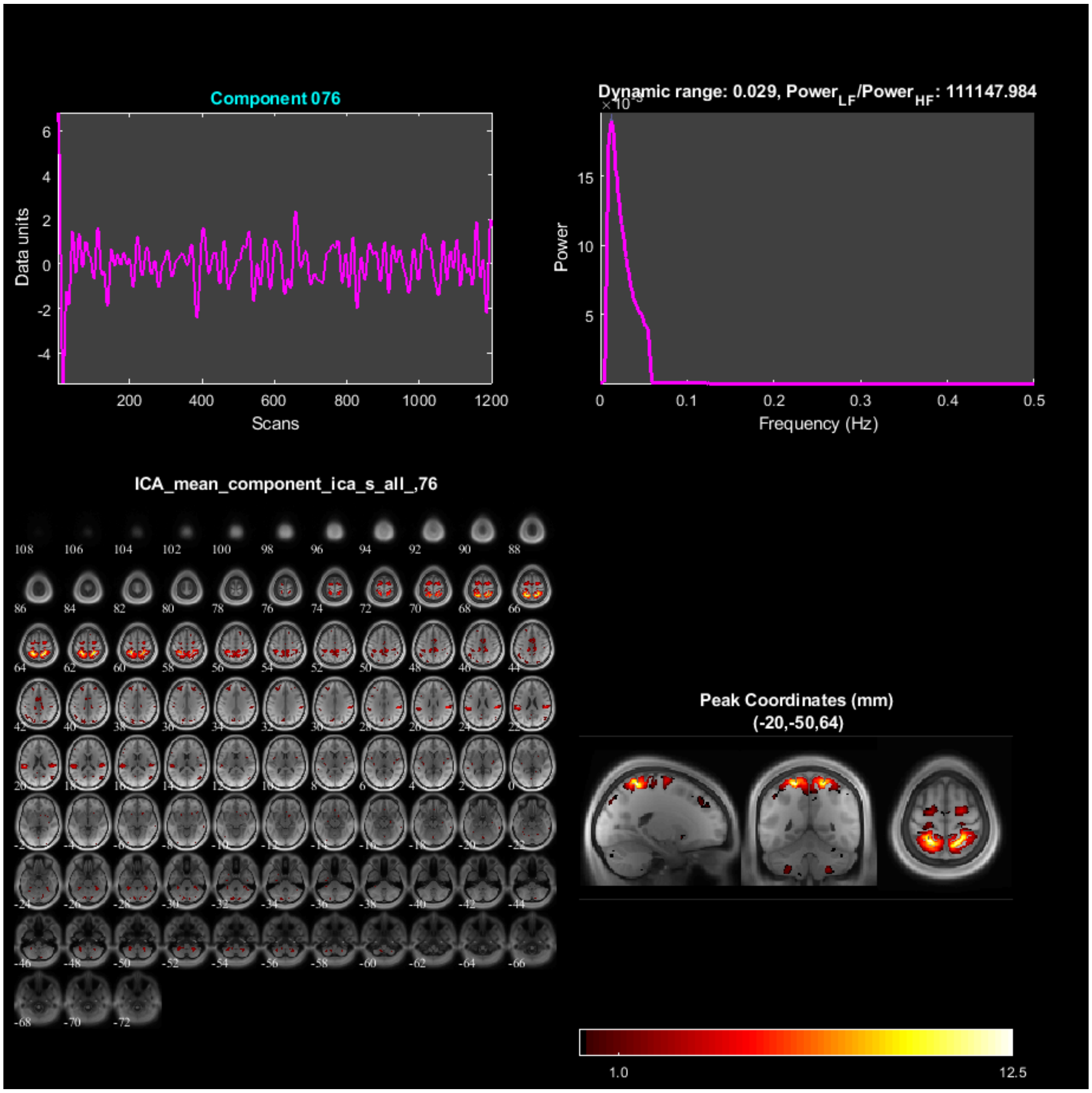

Component 076
Data units
Scans
Dynamic range: 0.029, Power$_{LF}$/Power$_{HF}$: 111147.984
Power
Frequency (Hz)
ICA_mean_component_ica_s_all_,76
Peak Coordinates (mm)
(-20,-50,64)
1.0
12.5

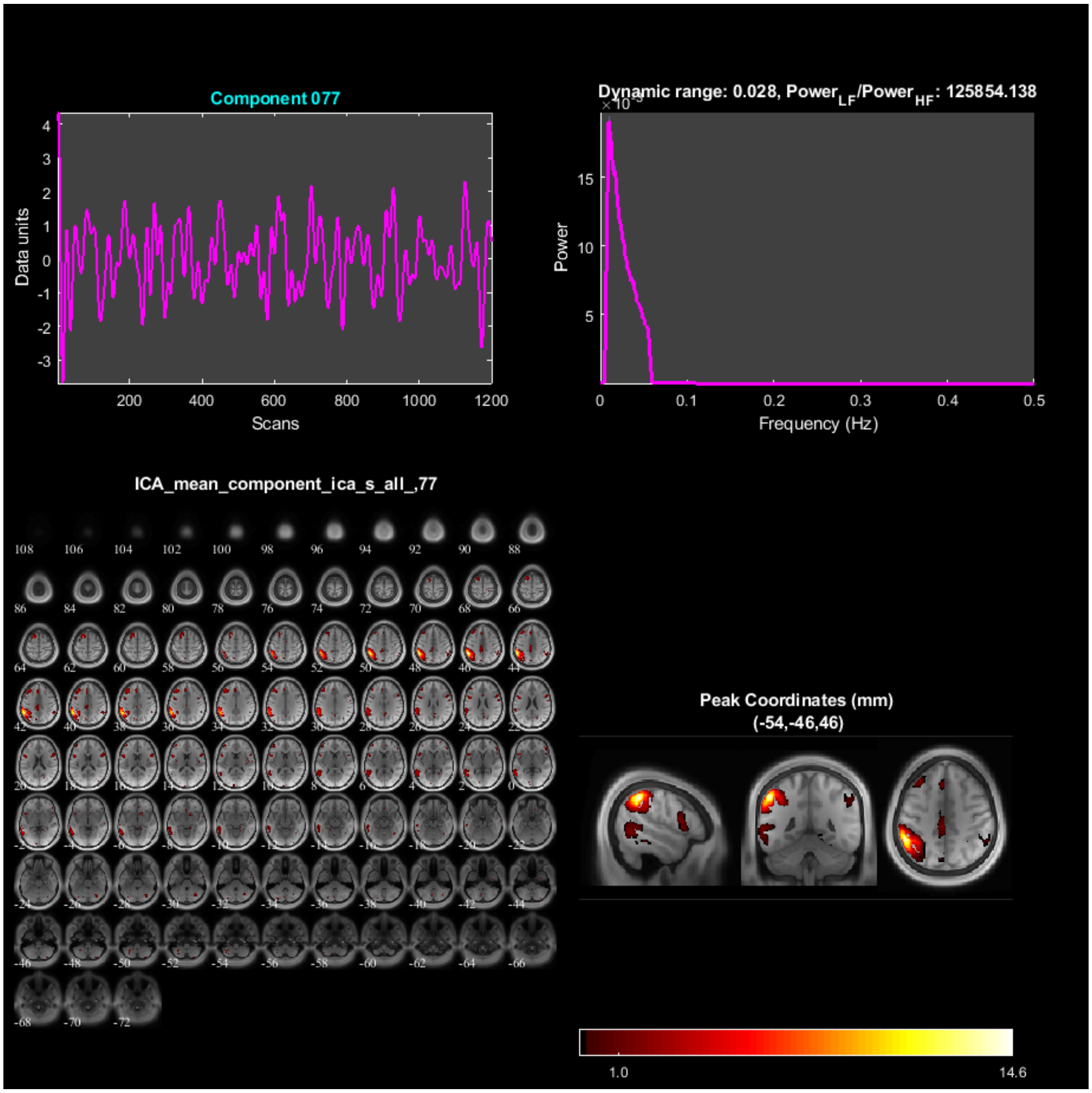

Component 077
Data units
Scans
Dynamic range: 0.028, Power$_{LF}$/Power$_{HF}$: 125854.138
Power
Frequency (Hz)
ICA_mean_component_ica_s_all_,77
Peak Coordinates (mm)
(-54,-46,46)
1.0
14.6

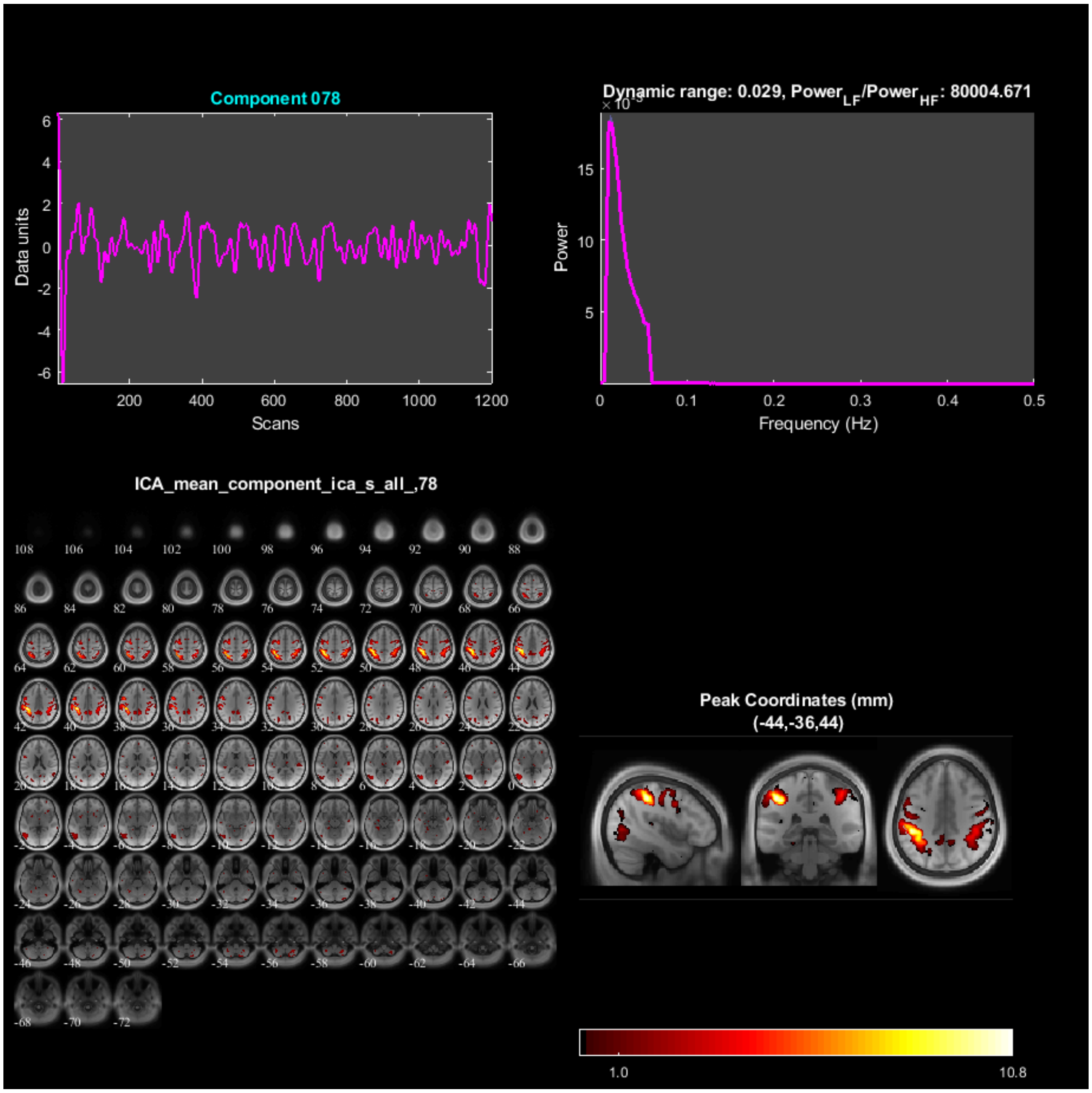
Component 078
Data units
Scans
Dynamic range: 0.029, Power_LF/Power_HF: 80004.671
Power
Frequency (Hz)
ICA_mean_component_ica_s_all_,78
Peak Coordinates (mm)
(-44,-36,44)
1.0
10.8

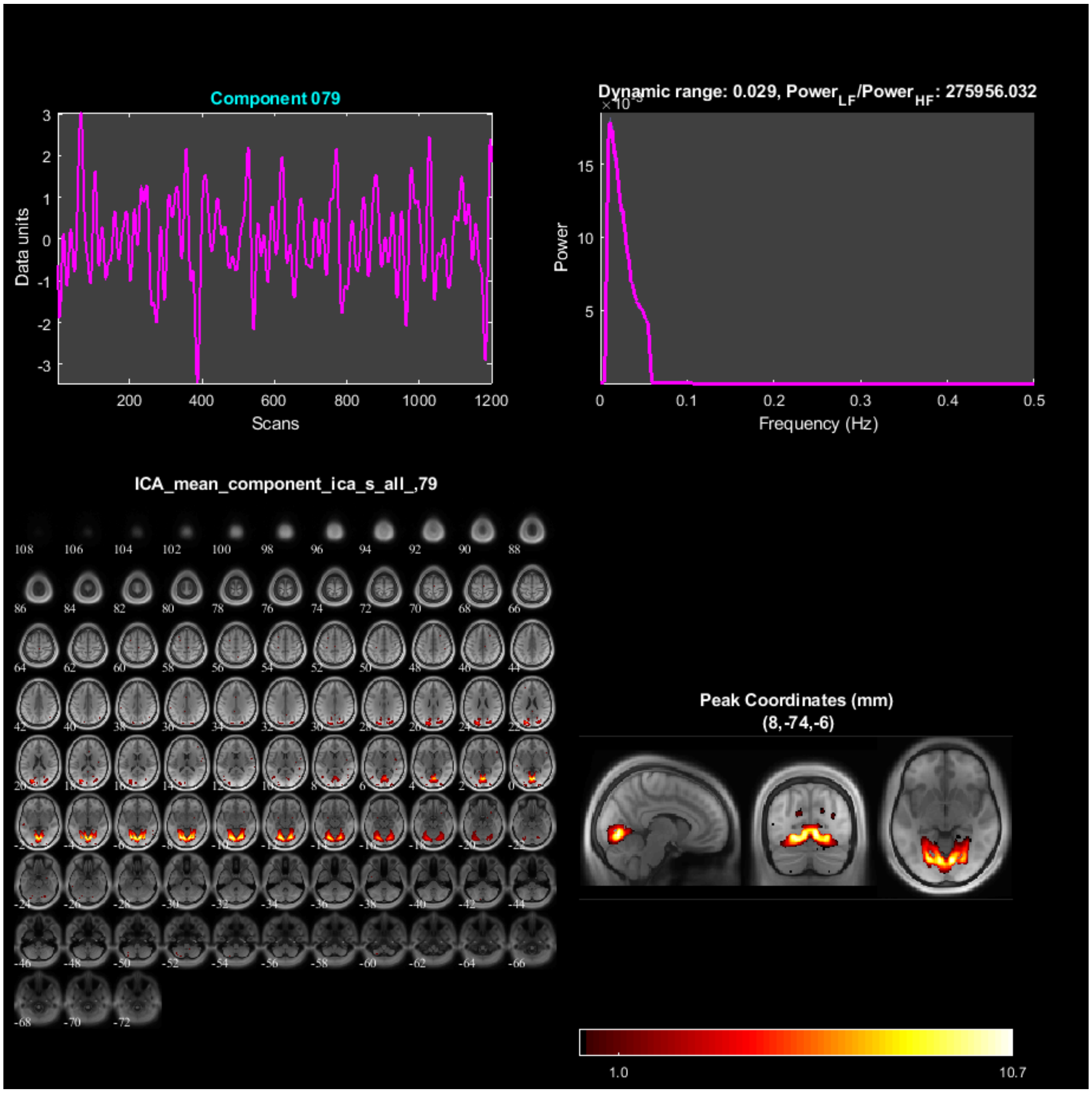

Component 079
Data units
Scans
Dynamic range: 0.029, Power$_{LF}$/Power$_{HF}$: 275956.032
Power
Frequency (Hz)
ICA_mean_component_ica_s_all_,79
Peak Coordinates (mm)
(8,-74,-6)
1.0
10.7

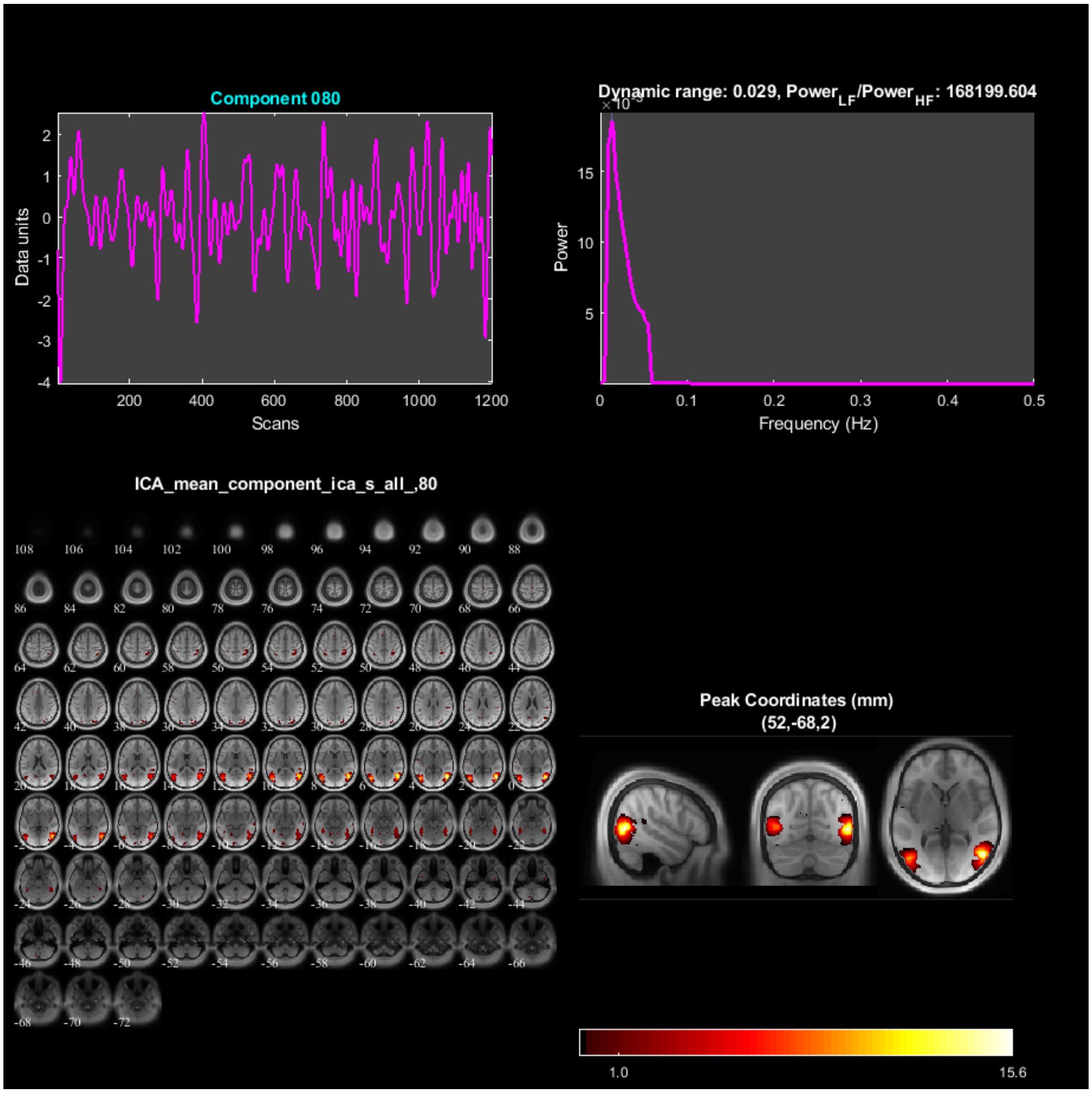

### Spectral Summary

- **a) dynamic_range** - Difference between the peak power and minimum power at frequencies to the right of the peak.
- **b) fALFF** - Low frequency to high frequency power ratio.

| ComponentNumber | DynamicRange | fALFF |
|---|---|---|
| 1 | 0.026924 | 1.1925e+05 |
| 2 | 0.02781 | 1.4812e+05 |
| 3 | 0.027293 | 2.1975e+05 |
| 4 | 0.025632 | 1.4812e+05 |
| 5 | 0.028286 | 75922 |
| 6 | 0.027835 | 78620 |
| 7 | 0.028434 | 1.3011e+05 |

| | | |
|---|---|---|
| 8 | 0.027874 | 1.1561e+05 |
| 9 | 0.02831 | 1.2952e+05 |
| 10 | 0.02914 | 1.3899e+05 |
| 11 | 0.029287 | 1.6897e+05 |
| 12 | 0.028333 | 1.3555e+05 |
| 13 | 0.029989 | 92472 |
| 14 | 0.028844 | 1.3906e+05 |
| 15 | 0.027874 | 86655 |
| 16 | 0.029031 | 2.1373e+05 |
| 17 | 0.035015 | 94762 |
| 18 | 0.025415 | 1.1416e+05 |
| 19 | 0.027181 | 71145 |
| 20 | 0.02677 | 1.6006e+05 |
| 21 | 0.026651 | 1.6071e+05 |
| 22 | 0.028703 | 1.281e+05 |
| 23 | 0.027245 | 1.2501e+05 |
| 24 | 0.02577 | 1.5955e+05 |
| 25 | 0.025673 | 1.1298e+05 |
| 26 | 0.027401 | 1.8494e+05 |
| 27 | 0.028126 | 1.0742e+05 |
| 28 | 0.026135 | 1.3378e+05 |
| 29 | 0.028189 | 1.0503e+05 |
| 30 | 0.028871 | 99633 |
| 31 | 0.025861 | 1.0696e+05 |
| 32 | 0.027004 | 1.3488e+05 |
| 33 | 0.026137 | 1.3872e+05 |
| 34 | 0.025944 | 1.6481e+05 |
| 35 | 0.026797 | 1.7285e+05 |
| 36 | 0.030338 | 1.6438e+05 |
| 37 | 0.029865 | 1.7205e+05 |
| 38 | 0.028082 | 1.7392e+05 |
| 39 | 0.027535 | 1.2488e+05 |
| 40 | 0.026508 | 1.8108e+05 |
| 41 | 0.027958 | 1.9555e+05 |
| 42 | 0.028465 | 1.2617e+05 |
| 43 | 0.029897 | 1.2076e+05 |
| 44 | 0.028038 | 1.6125e+05 |
| 45 | 0.028101 | 1.1769e+05 |
| 46 | 0.027501 | 1.7685e+05 |
| 47 | 0.025044 | 1.6231e+05 |
| 48 | 0.026378 | 1.6998e+05 |
| 49 | 0.028071 | 1.3523e+05 |
| 50 | 0.027266 | 1.013e+05 |
| 51 | 0.028139 | 1.9548e+05 |
| 52 | 0.026888 | 1.417e+05 |
| 53 | 0.029327 | 1.1502e+05 |
| 54 | 0.027712 | 1.0214e+05 |
| 55 | 0.026884 | 1.3009e+05 |
| 56 | 0.029079 | 1.549e+05 |
| 57 | 0.029063 | 1.3432e+05 |
| 58 | 0.028634 | 1.6918e+05 |
| 59 | 0.027774 | 1.5259e+05 |
| 60 | 0.0262 | 1.6363e+05 |
| 61 | 0.030022 | 1.1939e+05 |

| | | |
|---|---|---|
| 62 | 0.028916 | 94242 |
| 63 | 0.028863 | 1.2255e+05 |
| 64 | 0.028187 | 2.4554e+05 |
| 65 | 0.02731 | 2.3321e+05 |
| 66 | 0.028259 | 72936 |
| 67 | 0.028849 | 1.5143e+05 |
| 68 | 0.028706 | 81128 |
| 69 | 0.027847 | 2.0933e+05 |
| 70 | 0.027203 | 1.0525e+05 |
| 71 | 0.028024 | 1.3618e+05 |
| 72 | 0.027313 | 87245 |
| 73 | 0.027865 | 1.164e+05 |
| 74 | 0.027496 | 1.4305e+05 |
| 75 | 0.029487 | 1.5942e+05 |
| 76 | 0.028909 | 1.1115e+05 |
| 77 | 0.027906 | 1.2585e+05 |
| 78 | 0.029145 | 80005 |
| 79 | 0.028616 | 2.7596e+05 |
| 80 | 0.029267 | 1.682e+05 |

### Temporal Stats On Beta Weights

Multiple regression is done using the timecourses from SPM design matrix as model and ICA timecourses as observations. R^2 values for each component are shown in bar plot. For each component, one sample t-test results of each session and condition are shown in the bar plots.

### Kurtosis of timecourses and spatial maps

Mean across subjects is reported in table. Figure shows mean+/- SEM across subjects

| ComponentNumber | Timecourses | SpatialMaps |
|---|---|---|
| 1 | 3.6068 | 3.6011 |
| 2 | 3.2486 | 3.7042 |
| 3 | 3.3761 | 4.0481 |
| 4 | 3.4323 | 4.4485 |
| 5 | 3.6905 | 3.5249 |
| 6 | 3.1389 | 3.5391 |
| 7 | 3.0958 | 3.6872 |
| 8 | 3.1654 | 3.5374 |
| 9 | 3.2882 | 3.7555 |
| 10 | 3.1612 | 3.5757 |
| 11 | 3.4557 | 3.9303 |
| 12 | 3.0224 | 3.572 |
| 13 | 3.1846 | 3.6077 |
| 14 | 4.6573 | 4.3217 |
| 15 | 3.8009 | 3.5881 |
| 16 | 3.5724 | 4.8114 |
| 17 | 3.5657 | 4.2825 |
| 18 | 3.254 | 3.4575 |

| | | |
|---|---|---|
| 19 | 3.7032 | 3.5387 |
| 20 | 3.3406 | 3.6526 |
| 21 | 3.8238 | 4.0143 |
| 22 | 3.1118 | 3.62 |
| 23 | 3.2239 | 3.4267 |
| 24 | 3.2418 | 6.4134 |
| 25 | 3.1594 | 4.0652 |
| 26 | 3.0547 | 3.8713 |
| 27 | 3.2326 | 3.5672 |
| 28 | 3.3182 | 4.2312 |
| 29 | 4.1529 | 3.6129 |
| 30 | 3.2217 | 3.6297 |
| 31 | 3.4282 | 3.7034 |
| 32 | 3.3855 | 3.6042 |
| 33 | 3.1091 | 3.7661 |
| 34 | 3.2301 | 4.2435 |
| 35 | 3.0866 | 4.6102 |
| 36 | 3.0202 | 3.6925 |
| 37 | 3.337 | 3.8968 |
| 38 | 3.211 | 3.748 |
| 39 | 3.1313 | 3.8309 |
| 40 | 3.1953 | 3.5946 |
| 41 | 2.988 | 3.7757 |
| 42 | 3.4214 | 3.7313 |
| 43 | 3.1044 | 3.5456 |
| 44 | 3.2437 | 3.6345 |
| 45 | 3.2146 | 3.6195 |
| 46 | 3.0388 | 4.1016 |
| 47 | 3.2653 | 3.5695 |
| 48 | 3.1541 | 3.5061 |
| 49 | 3.1227 | 3.6307 |
| 50 | 3.1509 | 3.5594 |
| 51 | 4.2793 | 3.826 |
| 52 | 3.0918 | 3.5591 |
| 53 | 3.2824 | 3.6592 |
| 54 | 3.8633 | 3.5492 |
| 55 | 3.5053 | 3.5864 |
| 56 | 3.084 | 3.5225 |
| 57 | 3.2469 | 3.5951 |
| 58 | 3.0787 | 3.9887 |
| 59 | 3.0507 | 3.623 |
| 60 | 3.2589 | 4.2164 |
| 61 | 3.4196 | 3.6099 |
| 62 | 3.0059 | 3.5408 |
| 63 | 3.7834 | 3.5452 |
| 64 | 3.3187 | 3.9551 |
| 65 | 3.7793 | 8.2083 |
| 66 | 3.5607 | 3.4912 |
| 67 | 3.1439 | 3.5434 |
| 68 | 3.589 | 3.5065 |
| 69 | 3.2293 | 3.6699 |
| 70 | 3.2934 | 3.7716 |
| 71 | 3.2304 | 4.0498 |
| 72 | 3.2555 | 3.4722 |

| | | |
|---|---|---|
| 73 | 2.9456 | 3.7449 |
| 74 | 3.0636 | 3.5678 |
| 75 | 3.467 | 4.3387 |
| 76 | 3.2473 | 3.591 |
| 77 | 3.129 | 3.5964 |
| 78 | 3.2343 | 3.5539 |
| 79 | 3.361 | 3.6113 |
| 80 | 3.2687 | 4.055 |

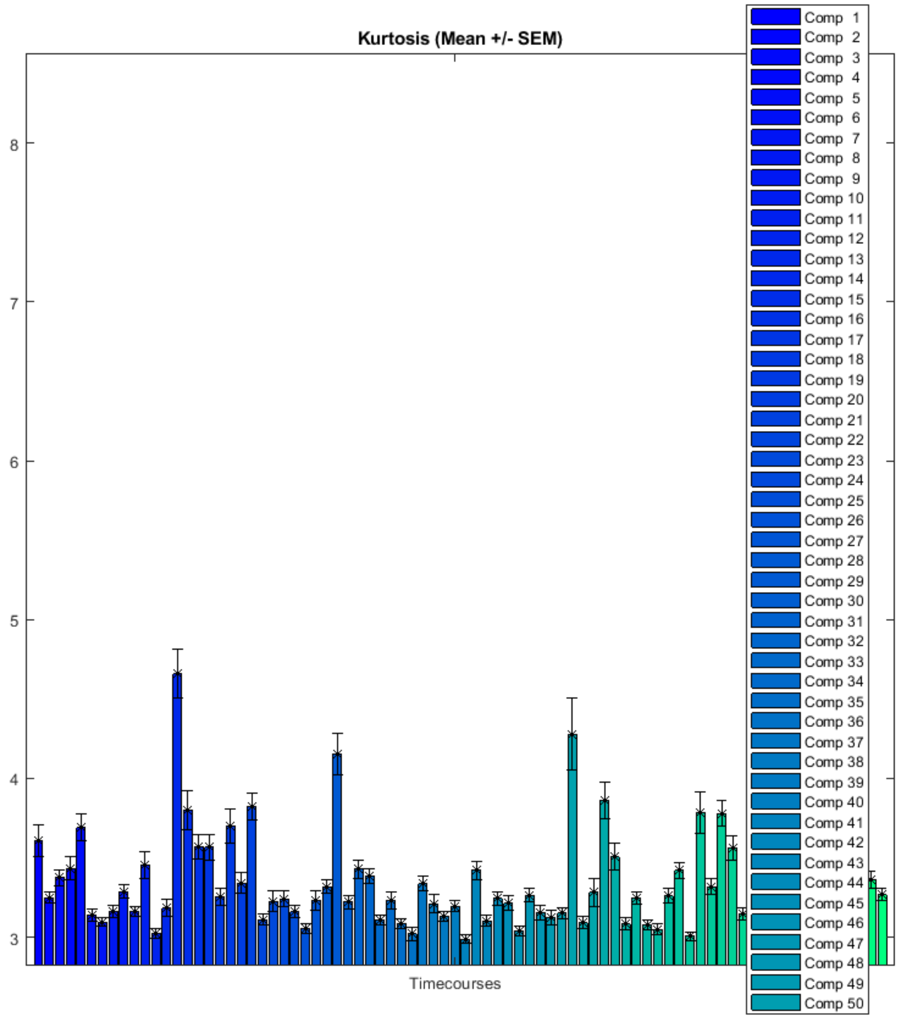

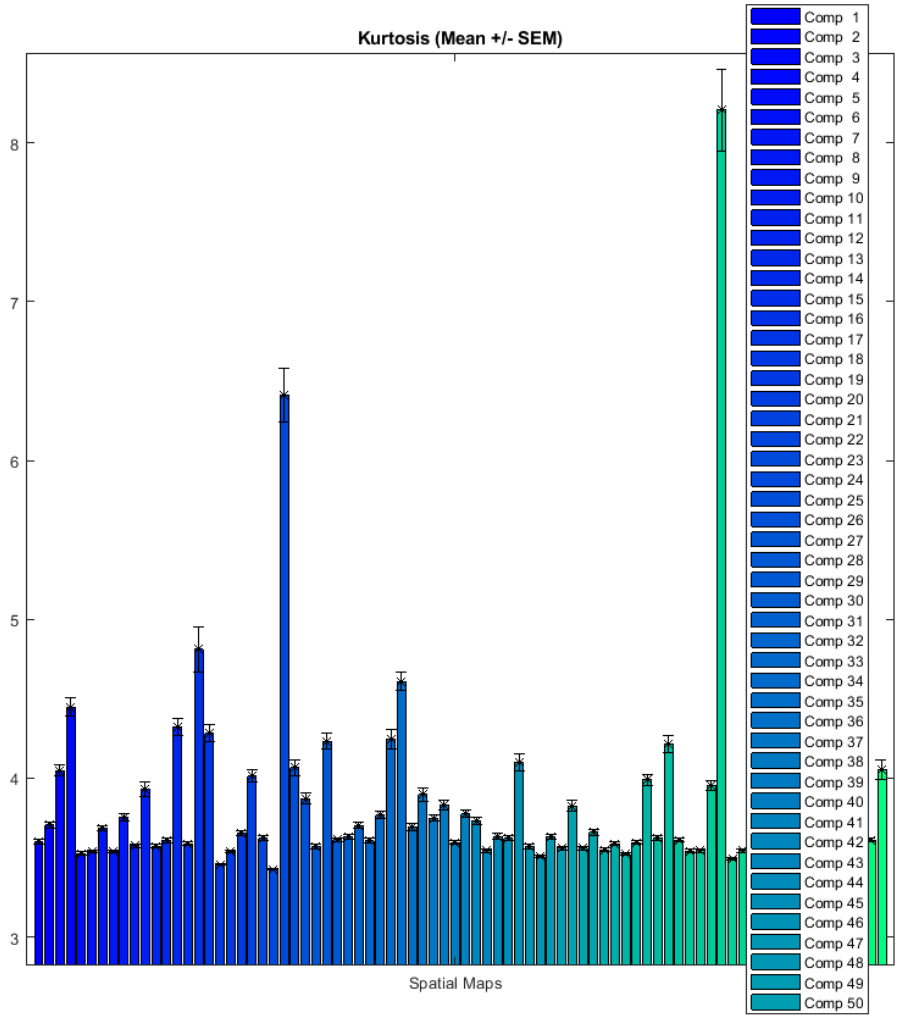

### FNC correlations

Functional network connectivity correlations are computed for each data-set and averaged across sessions.

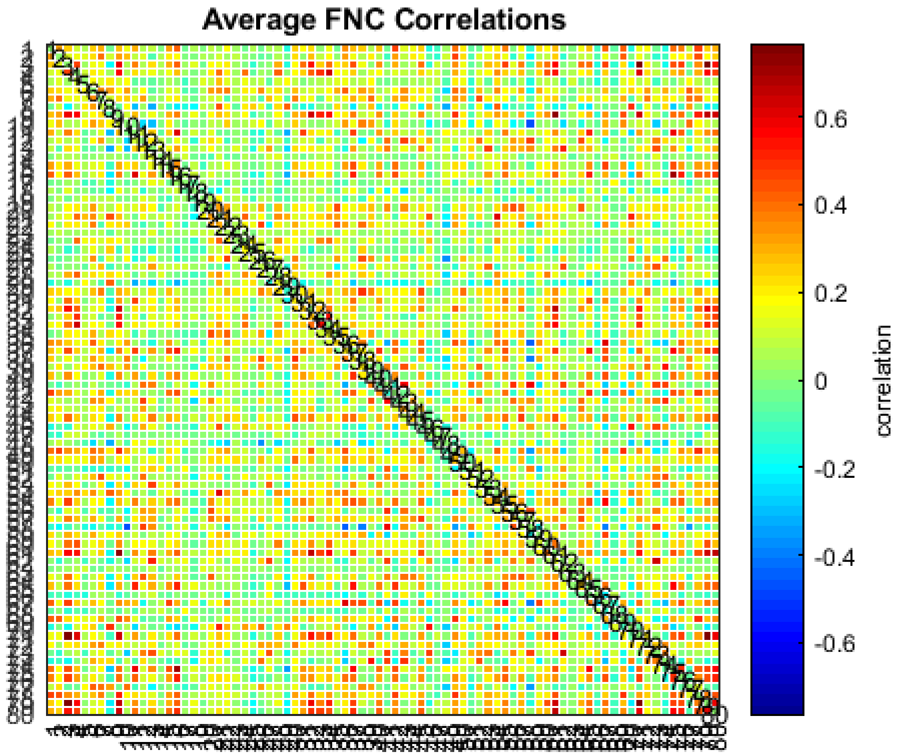

### FNC metrics of component spatial maps

Mutual information is computed between components spatially and averaged across data-sets.

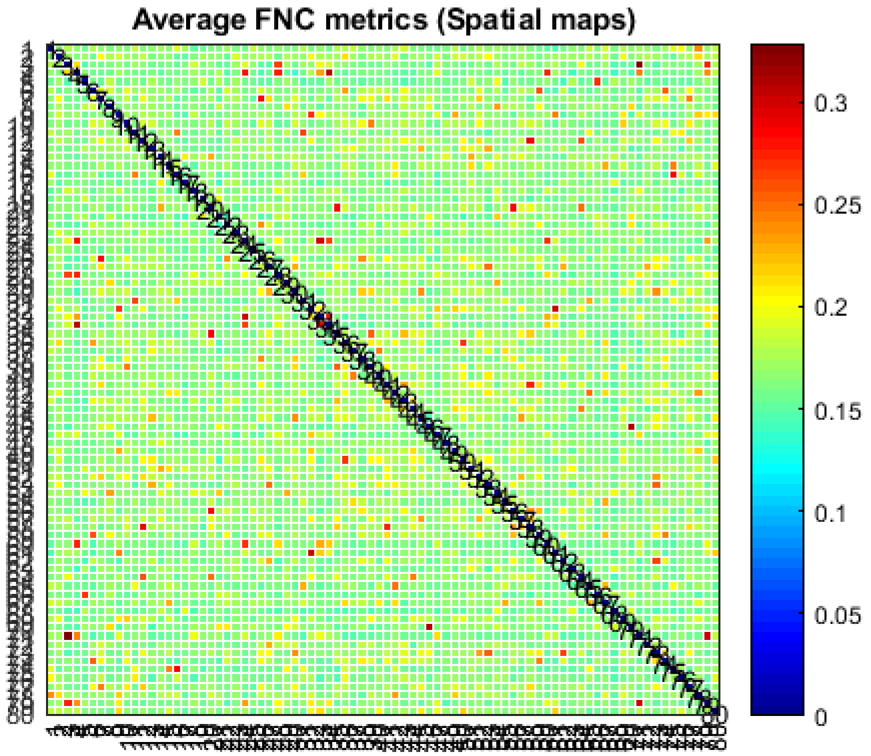

*Published with MATLAB® R2017b*
